# Temperature during *Aspergillus fumigatus* conidiophore development primes spore transcriptome for asexual, parasexual or sexual development

**DOI:** 10.64898/2026.06.11.730956

**Authors:** Justina M. Stanislaw, Michelle Momany

## Abstract

*Aspergillus fumigatus* is a thermotolerant saprobe found in soils and plant debris worldwide and an important pathogen of humans causing two million deaths annually. *A. fumigatus* makes abundant asexual spores (conidia) which are widely distributed by wind and can be inhaled from the environment. In susceptible individuals inhaled conidia break dormancy, germinate and grow in the lung leading to serious disease. Recent work has shown that conidia made at 37°C and 50°C have different morphologies and germination kinetics. While the asexual cycle is well-characterized at 37°C, much less is known about the asexual cycle at 50°C.

Here, we combine flow cytometry and transcriptomics to track morphology and gene expression in the hyphae, conidiophores and conidia of *A. fumigatus* during asexual development at 37°C or 50°C. We show that the temperature during a narrow time window in late-stage conidiophore development dictates resulting conidial morphology, transcriptional program, and germination kinetics. As expected, conidiation at 37°C resulted in upregulation of *brlA,* the master regulator of asexual development, and its downstream targets in conidiophores and conidia. Surprisingly, conidiation at 50°C resulted in upregulation of *MAT1-1*, the master regulator of sexual development and its downstream targets in conidiophores and conidia. Our findings suggest that temperature during late conidiophore development transcriptionally primes conidia for asexual, parasexual or sexual development enhancing chances of survival for progeny. Our findings are especially relevant for agricultural compost where a wide gradient of temperatures exists, abundant *A. fumigatus* has been isolated, and resistance to antifungals is thought to evolve.

**IMPORTANCE:** The human pathogen *Aspergillus fumigatus* has been found in natural and agricultural environments around the world. Disease is acquired when susceptible individuals inhale airborne asexual spores from the environment, which in agriculture generally includes proximity to compost and plant debris piles. This work shows that the environmental temperature when *A. fumigatus* spores are made determines the transcriptomes of those spores, priming them for future asexual or sexual development. The survival of asexual and sexual spores is very different at different temperatures, so these results are important for understanding how this pathogen survives in varied hostile environments. In addition, there are very few antifungal drugs with which to treat *A. fumigatus* infections, and resistance is increasing driven in part by agricultural use of fungicides. These results suggest that higher temperatures during asexual spore formation can lead to increased sexual reproduction and greater chances to evolve antifungal resistance.

## INTRODUCTION

The filamentous fungus *Aspergillus fumigatus,* a World Health Organization-designated fungal critical priority pathogen (1), is estimated to kill 2 million people annually (2). *A. fumigatus* is found in soil and leaf litter across varied environments and makes abundant dormant asexual spores (conidia) that are dispersed by wind. *A. fumigatus* disease is initiated when conidia are inhaled from the environment by a susceptible individual, break dormancy, and grow in the lungs. Although *A. fumigatus* primarily reproduces asexually, recent findings show that sexual reproduction also occurs in nature where it produces sexual spores (ascospores) that are important for generating the phenotypic diversity that allows *A. fumigatus* to survive in differing environments and to evolve antifungal resistance (3), (4).

In asexual development, the asexual reproductive structure (conidiophore) produces chains of conidia that contain genetically identical nuclei made by mitosis. Conidia are abundant and easily dispersed, but they are less genetically diverse than spores derived from other reproductive strategies. Many *Aspergillus* species use parasexual and sexual reproduction which increase diversity and chances of adapting to varied environments (5), (6). Before two unique fungal hyphae fuse (anastomose) and exchange nuclei, they must overcome barriers that prevent nonself fusion. For many filamentous fungi, including *A. fumigatus,* successful anastomosis requires that partners have specific *het* (heterokaryon incompatibility) genes at several loci ensuring that only closely related individuals fuse (7), (8). In parasexual reproduction the fusion of closely related hyphae and their nuclei creates diploids which can then haploidize (9). Though it is not clear how often parasex occurs in nature, it has been demonstrated in patients with long term *A. fumigatus* infections (9). Like asexual development, parasexual development produces chains of conidia on conidiophores by mitosis with the major difference being that parasex can give rise to variation by chromosome loss and mitotic recombination. Sexual reproduction gives much higher levels of variation via meiosis and meiotic recombination than either asexual or parasexual reproduction. Heterothallic fungi like *A. fumigatus* require partners of opposite mating types for sexual development (MAT1-1 and MAT1-2 in *A. fumigatus*). Mating partners secrete pheromones that are recognized by receptors in the opposite mating type partner and allow bypass of heterokaryon incompatibility so that unrelated individuals can fuse. In sexual development ascospores are made within specialized structures (cleistothecia) by meiosis.

Genetic regulation of asexual reproduction leading to conidia is well-defined at 37°C in standard laboratory conditions where it is known to be transcriptionally driven by a central regulatory pathway initiated by master regulator BrlA. BrlA binds to and activates expression of genes that contain BrlA response elements (BREs) (10), (11). Multiple direct and indirect downstream targets of BrlA play critical roles in conidial dormancy and maintaining conidial viability (12), including WetA and velvet regulators VeA, VosA, and VelB (11), (13). Conidiation and secondary metabolism are also tightly linked, with BrlA controlling expression of biosynthetic gene clusters leading to the production of specific secondary metabolites during conidiophore development (14), (15), (16), (17).

Genetic regulation of sexual reproduction leading to ascospores has also been defined. Since *A. fumigatus* is heterothallic, strains can only cross if they have opposite mating types, MAT1-1 and MAT1-2. These mating type genes are master regulators of sexual development that bind to downstream targets containing the corresponding DNA binding motifs. MAT targets directly or indirectly stimulate sexual development, secondary metabolism, and trigger pheromone secretion in the opposite mating type partner to initiate cleistothecia formation and ascospore production (18), (19), (20), (21), (22). Although the optimal niche for ascospore formation and germination in nature are not established, it is known that sexual spores thrive at high temperatures and require heat shocks of 65-85°C in order to break dormancy (9), (23).

In recent work, we found that conidia made at 50°C are larger and germinate faster than those made at 37°C (24). Other findings showed that conidial phenotypes and transcriptional profiles are influenced by the growth environment during the asexual lifecycle (25), (26), (27). Importantly, larger 50°C conidia have distinct cell wall compositions (26), (27), and are more virulent than those made at 37°C (24).

Although it has been established that distinct environments produce distinct conidial morphologies and transcriptomes, it is not known when this variation is determined during the asexual lifecycle of *A. fumigatus*. Here, we combine ImageStream flow cytometry and transcriptomics to investigate the impact of environmental temperature on landmark intermediates in the *A. fumigatus* asexual lifecycle.

We show that environmental temperature during late-stage conidiophore development dictates the resulting conidial phenotype and that once established, the phenotype cannot be altered. We found that transcriptomes of conidiophores made at 37°C are significantly upregulated in *brlA*, which encodes the master regulator of asexual development, and that conidia produced from these conidiophores are upregulated in downstream targets of BrlA. Surprisingly, we also found that conidiophores made at 50°C are significantly upregulated in *MAT1-1*, which encodes the master regulator of sexual development, and that conidia produced from these conidiophores are upregulated in downstream targets of MAT1-1. Our findings suggest that the growth environment during late-stage conidiophore development directs distinct regulatory pathways that prime the resulting conidia for higher fitness and impact genetic variability.

## RESULTS

### Asexual development at 37°C versus 50°C produces distinct and trackable phenotypes

*A. fumigatus* conidia are dormant and contain a single cell cycle-arrested nucleus. When exposed to carbon and water, conidia break dormancy, swell, and eventually resume nuclear division and extend germ tubes that become hyphae during vegetative growth. At the start of asexual development*, A. fumigatus* vegetative hyphae become partitioned by thick cross walls that form foot cells. Aerial hyphae emerge from foot cells and extend apically, forming conidiophore stalks. After a period of extension, stalks round at their tips to develop vesicles, and phialides emerge from the vesicles. Each phialide produces a chain of uninucleate conidia (asexual spores) which later separate and are dispersed by wind (Fig. 1A) (28). Previous work showed that *A. fumigatus* conidia produced from cultures incubated at 50°C are larger and break dormancy (germinate) faster than those produced from cultures incubated at 37°C (24). To determine whether key landmarks of the asexual lifecycle differ at 37°C and 50°C, we inoculated *A. fumigatus* CEA10 conidia at 37°C or 50°C and incubated for 4 days. We observed developing cultures every 4 hours until chains of conidia were visible (Fig. 1B and C).

**Fig 1.**
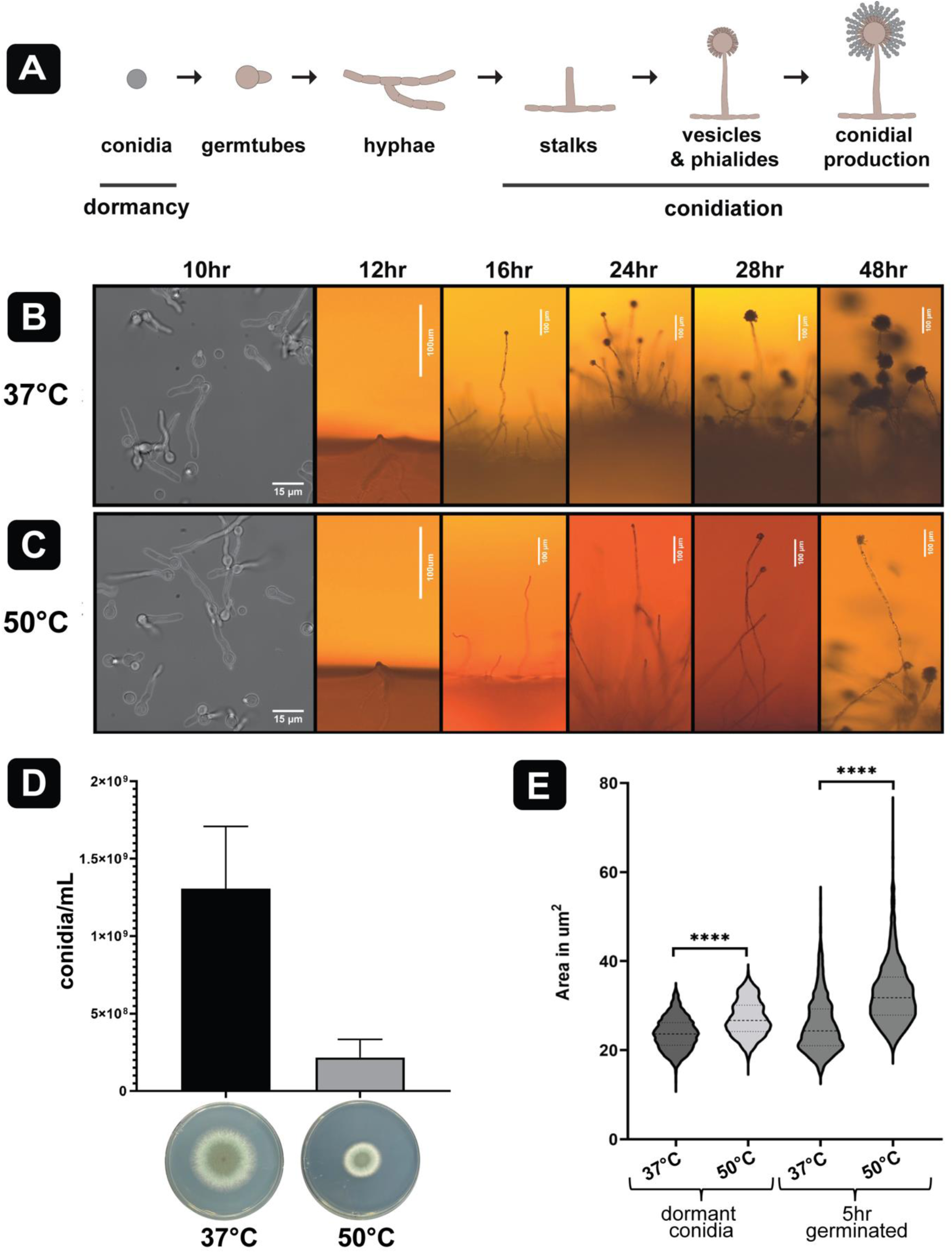
Temperature impacts conidiophore, conidia, and colony morphology. (**A**) Key milestones of *A. fumigatus* asexual lifecycle. (**B**, **C)** Asexual development at 37°C (B) or 50°C (C). Left column: Hyphae 10 hrs after inoculation. Scale bar = 15µm. Right 5 columns: 12-48 hr after inoculation. Scale bars = 100µm. **(D**) Top: Bar graph shows average number of conidia per mL produced at 37°C or 50°C from a single plate after 4d. n=3. (Bottom) colony morphologies after 3d incubation. (**E**) Image Stream flow cytometry measurements of conidia produced by incubation at 37°C or 50°C for 3 days and germinated for 5-hours at 37°C in liquid AMM. Conidial area in um^2^ (y-axis) with the middle line showing median. 2-tailed t-tests determined statistical significance (Mann-Whitney, nonparametric). P-values are indicated by stars *** p<0.001, **** p<0.0001. n=3 of 1,500-3,000 per sample.

No differences were observed between 37°C and 50°C cultures for the first 12 hours of incubation when both were making vegetative hyphae (Fig. 1B and C). Aerial hyphae emerged at 12 hours in both conditions as well. Following the emergence of aerial hyphae, growth rates and morphologies showed drastic differences between 37°C and 50°C conditions. At 37°C, we observed the end of stalk elongation and vesicle formation at 16 hours, presence of vesicles at 24 hours, and observable conidial production by 28 hours, consistent with previous reports (29). At 50°C, the end of conidiophore stalk elongation and vesicle/phialide formation for most of the population occurred at 28 hours with observable conidial production by 48 hours. Aerial hypha development was slower and less synchronous at 50°C than at 37°C, and 50°C conidiophores had longer, thinner stalks, smaller vesicles, and produced fewer conidia (Fig. 1B and C). The colony phenotypes also demonstrated striking differences, with the 50°C conditions giving rise to much smaller colonies with fewer conidia produced than at 37°C (Fig. 1D). After 4 days of growth, 37°C produced approximately 1.3x10^9^ conidia/mL, and 50°C produced roughly 2x10^8^ conidia/mL (Fig. 1E). We also observed smaller colonies with growth at 50°C compared to 37°C in *A. fumigatus* strain Af293 (Fig. S1A). Conidial phenotype was analyzed by flow cytometry to determine conidial area (Fig. 1E and S1B-D). The average area of the 50°C produced conidia was larger than 37°C produced, with a median of approximately 27 μm² and 24 μm², respectively (Fig. 1E, Table S1). Conidia produced at 37°C and 50°C showed significant differences in germination 5 hours after exposure to carbon containing medium, with the 50°C conidia establishing isotropic growth, germ tube emergence, and resuming nuclear division faster than the conidia produced at 37°C (Fig. S1C), consistent with results from previous studies (24), (25).

### The conidial phenotype is determined *during* conidiophore development and is unalterable once established

We next investigated whether there might be sensitive phases of the asexual lifecycle in which exposure to 37°C or 50°C caused the differences in conidial size and whether there might be a commitment point after which a change in temperature would not change the conidial phenotype. To detect possible sensitive phases and commitment points, we shifted cultures from 37°C to 50°C at defined developmental landmarks (Fig. 1B and C), incubated for 96 hours, and measured the resulting conidia with flow cytometry (Fig. 2A and B). Transferring plates from 37°C to 50°C during hyphal growth (8 hours) resulted in large conidia indistinguishable from conidia produced with 50°C incubation for the entire lifecycle (96 hours) (Fig. 2A). Transferring plates from 37°C to 50°C early in conidiophore development (16 hours) resulted in mostly large conidia with some smaller conidia also present.

**Fig 2.**
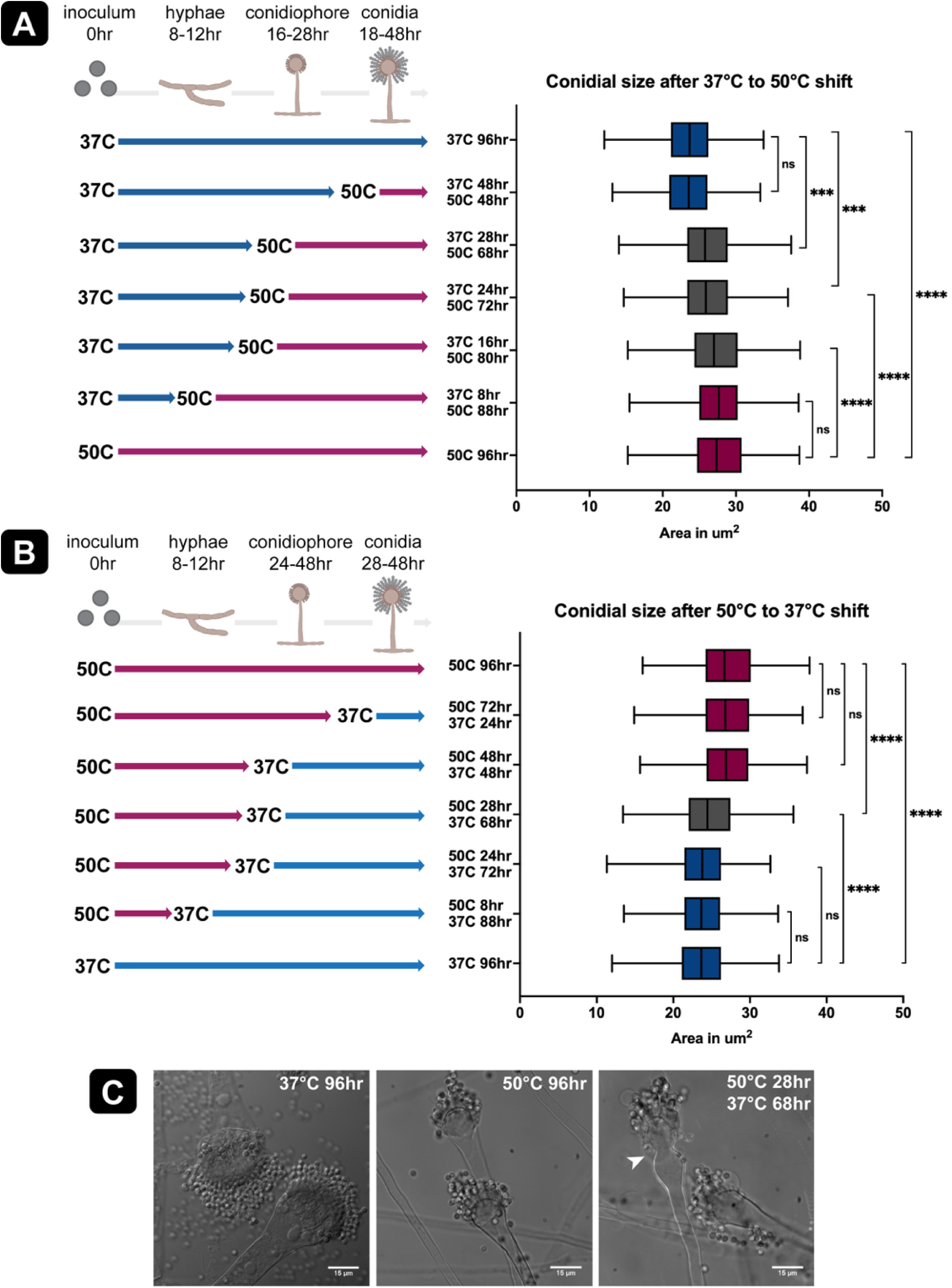
Conidial phenotype is determined during conidiophore development. Left: Temperature swaps during specified developmental milestones (**A**) 37°C to 50°C transfers. (**B**) 50°C to 37°C). Right: Conidia were collected after 96 hr incubation and cell area was measured with flow cytometry. The x-axis shows area of cells in um^2^. The middle line represents the median cell area. Statistical significance was determined by a 2-tailed t-test (Mann-Whitney, nonparametric). P-values are indicated by stars *** p<0.001, **** p<0.0001. Representative data of 3 replicates are plotted (n= 1,500-3,000 per sample). (**C**) Representative conidiophore morphologies after 96 hr incubation. White arrow indicates abnormal vesicle development.

Transferring plates from 37°C to 50°C later in conidiophore development (24-28 hours) also resulted in a mix of large and small conidia with a shift toward more small conidia. Transferring plates from 37°C to 50°C after completion of conidiophore development during conidial production (48 hours) resulted in small conidia indistinguishable from those produced entirely at 37°C (96 hours).

We also performed temperature shift experiments in the opposite direction transferring cultures from 50°C to 37°C (Fig. 2B). Considering the slower development at 50°C (Fig. 1C), our results gave an almost identical pattern as the 37°C to 50°C shift experiments (Fig. 2A). Transferring plates from 50°C to 37°C during hyphal growth (8 hours) or early in conidiophore development (24 hours) resulted in large conidia indistinguishable from conidia made by colonies incubated entirely at 50°C (96 hours).

Transferring plates from 50°C to 37°C later in conidiophore development (28 hours) showed a mix of large and small conidia. Transferring plates from 50°C to 37°C after conidiophore development was complete and conidia were being produced (48 and 72 hours) resulted in large conidia indistinguishable from conidia from colonies incubated entirely at 50°C (96 hours). Intriguingly, when the temperature was shifted during late conidiophore development, we also observed some conidiophores with abnormal vesicles and swollen phialides (Fig. 2C).

Our results suggest that environmental input during late conidiophore development (vesicle and phialide formation) determines the conidial phenotype and so defines the environmentally-sensitive phase of conidial development. Our results further suggest that the conidial phenotype is not alterable once conidia begin to emerge from phialides, defining the commitment point of conidial phenotype.

These patterns hold true whether beginning at 37°C and shifting to 50°C or beginning at 50°C and shifting to 37°C (Fig. 2B). The distinct conidiophore morphologies at 37°C versus 50°C (Fig. 1B and C) and conidial morphologies that resulted from temperature shift experiments (Fig. 2C) suggest that the conidiophores themselves also have an environmentally-sensitive phase either before or early in aerial hypha emergence.

### Hyphae, conidiophores, and conidia have distinct transcriptomes at 37°C versus 50°C

To better understand the mechanisms underlying the temperature sensitive phase and phenotypic commitment point of conidia, we developed a method for isolating conidiophores from hyphae that used aerial growth through a thin layer of water agar followed by careful shearing with a razor (Fig. S2). We then extracted RNA from isolated hyphae, conidiophores, or conidia produced at each temperature, and analyzed the transcriptomes. We performed pairwise comparisons of total genes expressed in each cell type at 37°C or 50°C (Fig. 3A, Table S2). Of 8,717 genes expressed in hyphae, most (8,003; 91.8%) were expressed both at 37°C and 50°C, with far fewer expressed only at 37°C (320; 3.6%), or only at 50°C (394; 4.5%). Of 9,549 genes expressed in conidiophores, most (8,924; 93.5%) were expressed at both 37°C and 50°C, with far fewer expressed only at 37°C (33; 0.3%), or only at 50°C (592; 6.2%). Of 9,157 genes expressed in conidia, most (8,307; 90.7%) were expressed at both 37°C and 50°C, with far fewer expressed only at 37°C (310; 3.4%), or only at 50°C (540; 5.9%). Since the largest transcriptional differences between each cell type at both temperatures were in the conidiophores, we speculated that the conidial phenotypic program determined during late conidiophore development is transcriptionally driven.

**Fig 3.**
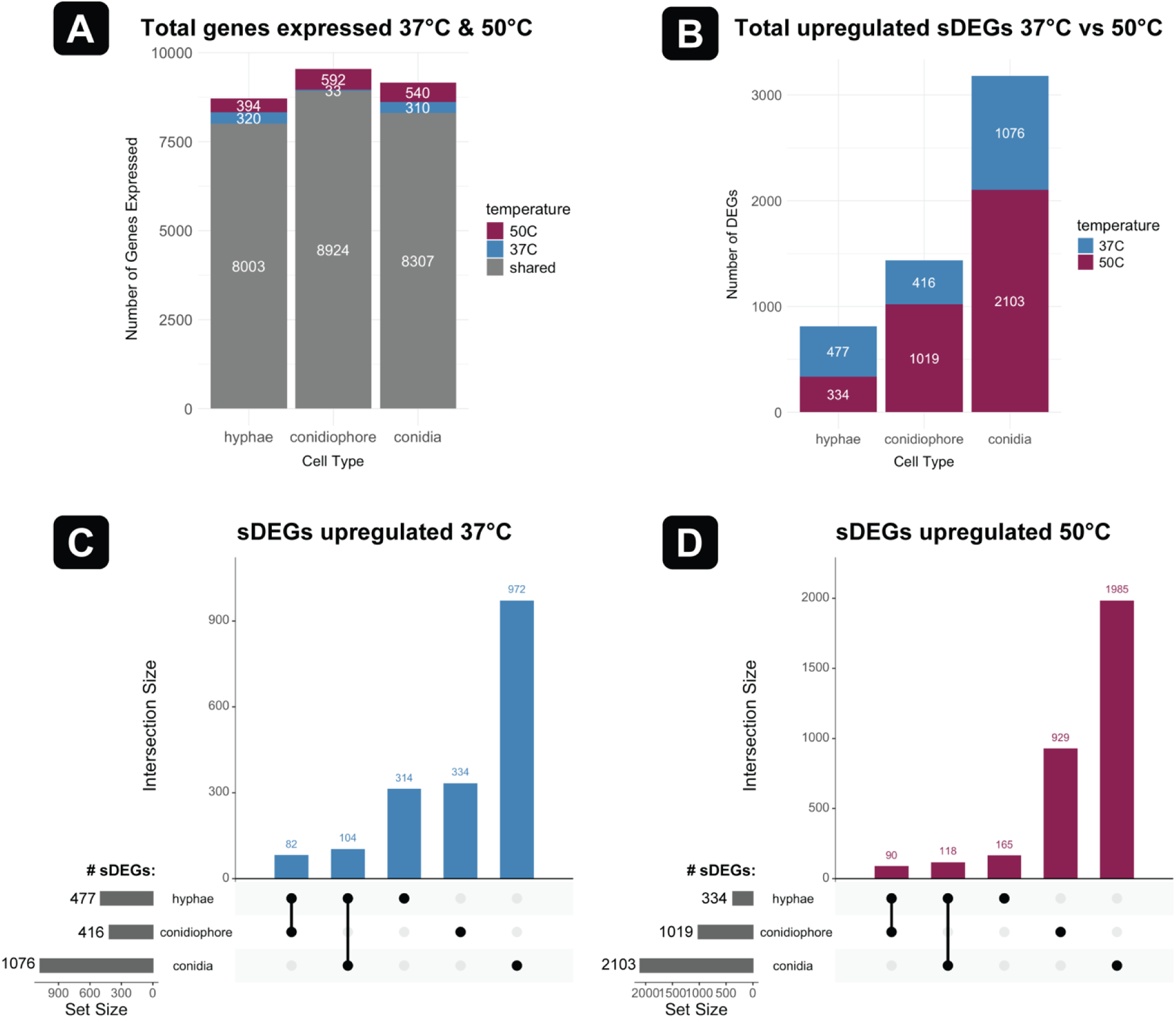
Hyphae, conidiophores, and conidia have distinct transcriptomes at 37°C versus 50°C. (**A**)Total genes expressed in hyphae, conidiophores, and conidia at 37°C and 50°C. Magenta denotes genes expressed only at 50°C. Blue denotes genes expressed only at 37°C. Grey denotes genes expressed at both temperatures. White denotes total number of genes for each category. (**B**) Significant differentially expressed genes (sDEGs) in hyphae, conidiophores, and conidia at 37°C versus 50°C. (**C**) Upregulated sDEGs at 37°C. (**D**) Upregulated sDEGs at 50°C.

We next identified genes with significant differentially expressed genes (sDEGs) between 37°C versus 50°C in hyphae, conidiophores, and conidia using DESeq2 with filtering based on an adjusted p-value (padj) of < 0.05 and Log2Fold Change > 2 (Fig. 3B-D, Table S3). Our results showed that hyphae had the least sDEGs (811), followed by conidiophores (1,435), and conidia had the most sDEGs (3,179) (Fig. 3B). Based on our results showing that shifting temperature during hyphal growth does not impact asexual development (Fig. 1B-C and 2A-B), we removed all upregulated sDEGs that were shared with hyphae for 37°C and 50°C, respectively. Removal of upregulated hyphal sDEGs yielded 334 conidiophore and 972 conidial genes significantly upregulated at 37°C (Fig. 3C), and 929 conidiophore and 1,985 conidial genes significantly upregulated at 50°C (Fig. 3D, Table S3). The resulting hyphal-removed conidiophore sDEGs and conidial sDEGs were further analyzed to determine regulation of conidial production at 37°C and 50°C.

### 37°C conidiophores are upregulated in *brlA* while 50°C conidiophores are upregulated in *MAT1-1*

To identify genes that could be important in the environmentally sensitive phase for conidial development, we analyzed significant differentially expressed genes (sDEGs) between 37°C and 50°C conidiophores. In gene ontology (GO) enrichment analysis of upregulated sDEGs (30), the top biological processes for 37°C conidiophores included genes associated with peroxisome synthesis, transmembrane transport, and alkaloid metabolism (Fig. S3A and B). The top biological processes for 50°C conidiophores included genes associated with secondary metabolism, mycotoxin production and gliotoxin biosynthesis (Fig. S3C and D). The enrichment for genes associated with fumagillin/pseurotin and gliotoxin biosynthetic gene clusters (BGCs) in 50°C conidiophores was especially surprising because these secondary metabolites have been shown to occur during sexual development, not in the asexual development that gives rise to conidiophores (21).

The unexpected enrichment of secondary metabolite transcripts associated with sexual development in 50°C conidiophores prompted us to examine the expression of known regulators of both asexual and sexual development in 37°C and 50°C conidiophores (Fig. 4B, Table S3). The upregulated genes in 37°C conidiophores included the master regulator of asexual development, *brlA* (AFUB_015960) (10), (11) (Fig. 4A and C, Tables 1 and S3). In contrast, other regulators of asexual development showed very similar expression levels in 37°C versus 50°C conidiophores (Fig. 4B). Members of the Fumigaclavine C BGC and alkaloid BGC, known downstream targets of BrlA, were also upregulated in 37°C conidiophores (Fig. 4B and C, Table 1), including *easA* (AFUB_033650), FtmPT1 (AFUB_086330), FtmPT2 (AFUB_086290), FtmP450 (AFUB_086300), FtmE (AFUB_086320), FtmD (AFUB_086340) (14), (15), (16). The upregulation of *brlA*, Fumigaclavine C BGC and alkaloid secondary metabolite pathways in 37°C conidiophores indicated that the well-characterized asexual development pathway was active.

**Fig 4.**
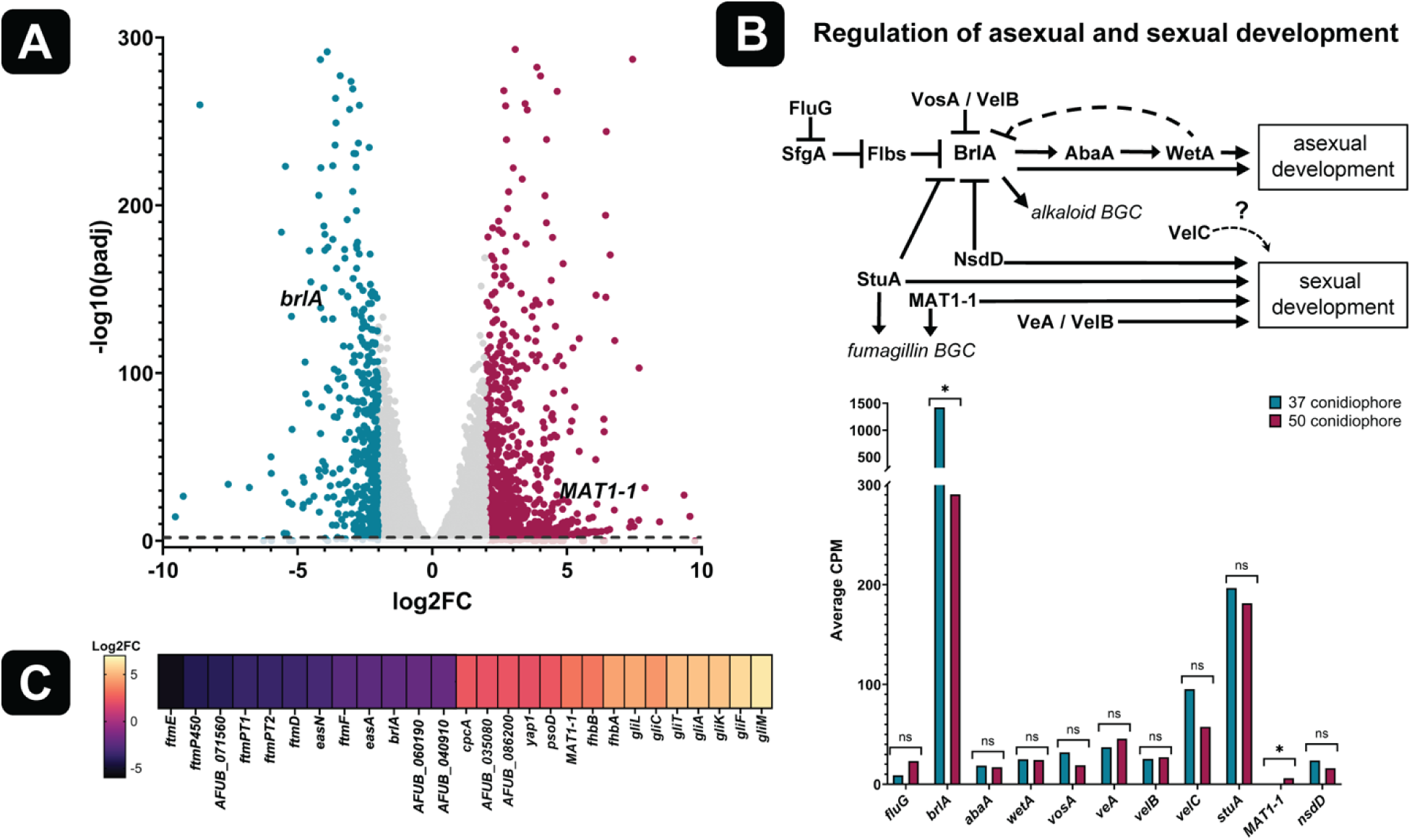
*brlA* is upregulated in 37°C conidiophores while *MAT1-1* is upregulated in 50°C conidiophores. (**A**) Volcano plot of 37°C versus 50°C conidiophore sDEGs with Log2FoldChange (Log2FC) and adjusted p-value (-Log10(padj)). Blue denotes 37°C upregulated sDEGs (416 total); magenta, 50°C upregulated sDEGs (1019 total); grey and below dashed line, not significant (filtered |Log2FC| < 5). (**B**) Top: Schematic diagram of the genetic regulation of asexual and sexual development. Bottom: transcript levels of regulators of asexual and sexual developmental (average CPM of 3 biological replicates). Blue: 37°C Magenta: 50°C. The star (*) indicates significant differential expression between 37°C and 50°C conidiophores; ns indicates not significant. (**C**) Heatmap of conidiophore sDEGs according to increasing Log2FC. Negative values (purple/black) indicate upregulation in 37°C conidiophores, positive values (orange/yellow) indicate upregulation in 50°C conidiophores. Detailed descriptions of genes in heatmap are in Table 1.

**TABLE 1.** 37°C versus 50°C conidiophore upregulated sDEGs^a^.

| Gene IDb | Gene | Descriptionc | Reference |
| --- | --- | --- | --- |
| **37°C conidiophore upregulated sDEGs** |  |  |  |
| AFUB_015960 | *brlA* | Central regulator of asexual development: activator of conidiation | (10), (11) |
| AFUB_040910 | NA | Predicted role in asexual sporulation resulting in formation of a cellular spore, establishment of cell polarity | NA |
| AFUB_060190 | NA | Predicted plasma membrane protein, predicted roles in regulation conidium formation | NA |
| AFUB_086330 | *FtmPT1* | Brevianamide F prenyltransferase FtmPT1, Fumitremorgin biosynthetic process. | (14), (15), (16) |
| AFUB_086290 | *FtmPT2* | 12-alpha,13-alpha-dihydroxyfumitremorgin C prenyltransferase FtmPT2, Fumitremorgin biosynthetic process | (14), (15), (16) |
| AFUB_086300 | *FtmP450* | FtmP450 Fumitremorgin C monooxygenase, Fumitremorgin biosynthetic process | (14), (15), (16) |
| AFUB_086340 | *ftmD* | O-methyltransferase FtmD, Fumitremorgin biosynthetic process | (14), (15), (16) |
| AFUB_086320 | *ftmE* | Cytochrome P450 FtmE, Fumitremorgin biosynthetic process. | (14), (15), (16) |
| AFUB_086310 | *ftmF* | Verruculogen synthase FtmF, Fumitremorgin biosynthetic process | (14), (15), (16) |
| AFUB_033650 | *easA* | NADH oxidase EasA, Ergot alkaloid biosynthetic process | (14), (15), (16) |
| AFUB_033710 | *easN* | Fumigaclavine B O-acetyltransferase EasN, Ergot alkaloid biosynthetic process | (14), (15), (16) |
| **50°C conidiophore upregulated sDEGs** |  |  |  |
| AFUB_042900 | *MAT1-1* | *A. fumigatus* mating-type protein MAT1-1 | (21), (19) |
| AFUB_099650 | *fhbA* | NO response FhbA, contains MAT1-1 binding motif. Regulator of sexual development | (22), (20) |
| AFUB_081400 | *fhbB* | FhbB, regulator of sexual development | (22), (46) |
| AFUB_035080 | NA | Predicted role in cleistothecium development, sexual sporulation resulting in formation of a cellular spore | NA |
| AFUB_086200 | NA | Putative beta-ketoacyl synthase. Predicted roles in pseurotin A biosynthetic process | NA |
| AFUB_086010 | *psoD* | PsoD P450 monooxygenase, Fumagillin/pseurotin biosynthesis. | (21) |
| AFUB_069420 | *cpcA* | CpcA, transcriptional activator of cross-pathway control system of amino acid biosynthesis | (61) |
| AFUB_075760 | *gliA* | GliA, gliotoxin biosynthetic process | (68) |
| AFUB_036230 | *gliC* | GliC, gliotoxin biosynthetic process | (68) |
| AFUB_075780 | *gliF* | GliF, gliotoxin biosynthetic process | (68) |
| AFUB_075750 | *gliK* | GliK, gliotoxin biosynthetic process | (68) |
| AFUB_075690 | *gliL* | GliL, gliotoxin biosynthetic process | (68) |
| AFUB_036280 | *gliM* | GliM, gliotoxin biosynthetic process | (68) |
| AFUB_075790 | *gliT* | GliT, gliotoxin biosynthetic process | (68) |
| AFUB_075990 | *yap1* | Yap1, regulator of oxidative stress. Important in sensing and producing gliotoxin | (68) |

<sup>a</sup>Upregulated significant differentially expressed genes (sDEGs) from 37°C versus 50°C conidiophore differential analyses that are listed in Fig. 4C heatmap.
<sup>b</sup> Gene ID from FungiDB for *Aspergillus fumigatus* strain A1163.
<sup>c</sup> Description simplified from FungiDB predicted protein function and literature references.

To our surprise, the upregulated genes in 50°C conidiophores included the master regulator of sexual development, MAT1-1 (AFUB_042900) (Fig. 4A and C, Tables 1 and S3) (19), (22). Transcript levels of other known regulators of sexual development showed similar expression levels in 37°C versus 50°C conidiophores (Fig. 4B). Numerous targets of MAT1-1 containing MAT1.1 DNA binding motifs including members of the Fumagillin/pseurotin and gliotoxin BGCs, and *fhbA* (AFUB_099650) and *fhbB* (AFUB_081400) (22), (20) (Fig. 4C) were upregulated in 50°C conidiophores (Fig. 4C, Table 1).

Notably, MAT1-1 was not differentially expressed between the 37°C versus 50°C conidia or hyphae, indicating that upregulation of MAT1-1 at 50°C is limited to the conidiophore. The upregulation of MAT1-1 and its downstream targets in 50°C conidiophores suggest that an alternative developmental pathway might be active at the higher temperature.

### 37°C conidia express higher levels of BrlA targets and dormancy regulators

To identify genes that could be important in the commitment point for conidial development, we analyzed conidial genes whose expression were significantly differential between 37°C versus 50°C (sDEGs). In GO term enrichment analyses (30) of sDEGs upregulated in 37°C conidia, the top biological GO processes included carbon and acetate utilization, response to light, regulation of spore-bearing structures, and the biosynthesis of several sugars including trehalose (Fig. S4A and B), all consistent with processes known to be important in making dormant asexual spores (conidia) or early conidial germination (13), (31).

Based on the upregulation of *brlA* in the 37°C conidiophores (Fig. 4A-C, Table S3), we expected that downstream targets of BrlA would be upregulated in 37°C conidia versus 50°C conidia. Indeed, numerous downstream targets of BrlA were significantly upregulated in 37°C conidia, including *wetA* (AFUB_070140) (32) and velvet regulators *vosA* (AFUB_067940) and *velB* (AFUB_002350) (10), (11), (33) (Fig. 5A and B, Tables 2 and S3). Previous studies found that WetA regulates trehalose biosynthesis and cell wall integrity, playing vital roles in conidial viability during dormancy across *Aspergillus* species (34), (13), (35). In addition to their importance during conidiation, VosA and VelB together play critical roles in conidial dormancy (33). The VosA-VelB heterodimer inhibits germination and is important in trehalose production, consistent with our GO enrichment findings (Fig. S4A and B). Other regulators of conidial viability and dormancy maintenance were upregulated in 37°C conidia (Fig. 5A-D, Tables 2 and S3) including members of the mitogen-activate protein kinase (HOG-MAPK) pathway including MAPK SakA (AFUB_012420) (36), (35) and its downstream transcription factor AtfA (AFUB_037850). Conidial dormancy is regulated by AtfA which inhibits germination and controls 63% of all conidial transcripts (4). Many downstream targets of AtfA were also upregulated in 37°C conidia, including *atfB* (AFUB_060680) (37), *atfC* (AFUB_016730) (38), *cspA* (AFUB_040120) (39), *drpA* (AFUB_101360) (40), *catA* (AFUB_094400) (41), and *con-10* (AFUB_095090) (Fig 5D, Tables 2 and S3). The transcriptional driver of conidial survival MybA (AFUB_041990) (31) was also upregulated, which functions upstream of VelB-VosA and WetA that in turn upregulates trehalose biosynthesis and cell wall genes (12), (35). 37°C conidia expressed higher levels of BrlA targets and known conidial dormancy genes, consistent with 37°C conidiophores’ upregulation of *brlA*, the master regulator of asexual development.

**Fig 5.**
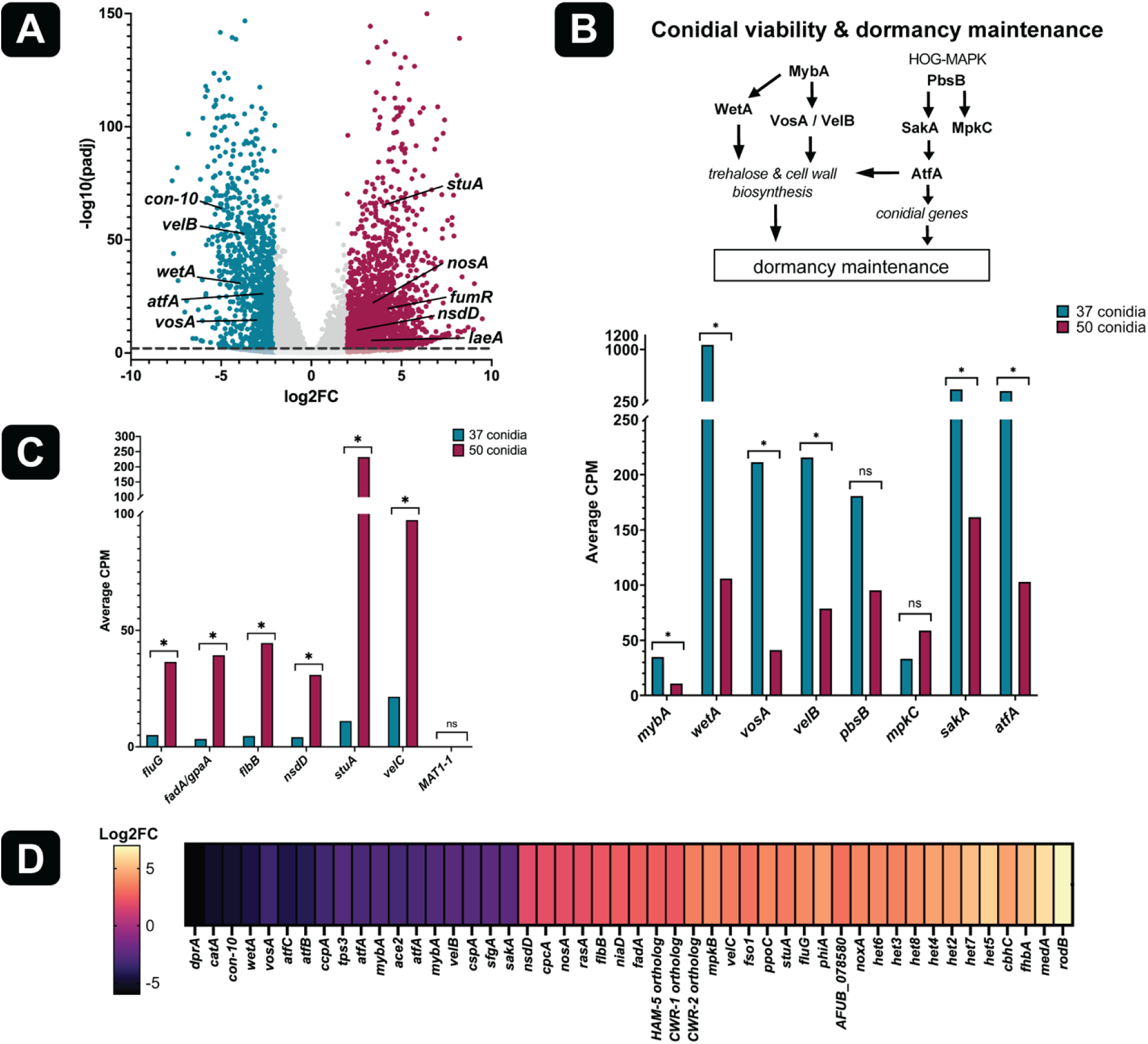
Targets of brlA and regulators of dormancy are upregulated in 37°C conidia while MAT1-1 targets and anastomosis genes are upregulated in 50°C. (**A**) Volcano plot of 37°C versus 50°C conidia sDEGs with Log2FoldChange (Log2FC) and adjusted p-value (-Log10(padj)). Blue denotes 37°C upregulated sDEGs (1,076 total); magenta, 50°C upregulated sDEGs (2,103 total); grey and below dashed line, not significant (filtered |Log2FC| < 5). (**B**) Top: Schematic diagram of the genetic regulation of viability and dormancy maintenance in conidia. Bottom: transcript levels of regulators of viability and dormancy (average CPM of 3 biological replicates). Blue: 37°C Magenta: 50°C. The star (*) indicates significant differential expression between 37°C and 50°C conidia; ns indicates not significant. (**C**) Transcript levels of negative regulators of asexual development and positive regulators of sexual development (average CPM of 3 biological replicates). Blue: 37°C Magenta: 50°C. The star (*) indicates significant differential expression between 37°C and 50°C conidia; ns indicates not significant. (**D**) Heatmap of conidial sDEGs according to increasing Log2FC. Negative values (purple/black) indicate significant upregulation in 37°C conidia, positive values (orange/yellow) indicate significant upregulation in 50°C conidia. Detailed descriptions of genes in heatmap are in Table 2.

**TABLE 2.** 37°C and 50°C conidia upregulated sDEGs involved in development^a^.

| Gene ID<sup>b</sup> | Gene | Description<sup>c</sup> | Reference |
| --- | --- | --- | --- |
| **37°C conidia upregulated sDEGs** |  |  |  |
| AFUB_051340 | *sfgA* | C6 transcription factor SgfA. Involved in regulating conidiation initiation | (69) |
| AFUB_070140 | *wetA* | Central regulator of asexual development and spore maturation WetA. Important in conidia wall formation and trehalose biosynthesis. Regulated by BrIA. | (32), (10), (33) |
| AFUB_012420 | *sakA* | Protein kinase SakA, in HOG-MAPK pathway and CWI pathway | (35), (4) |
| AFUB_037850 | *atfA* | Transcription factor AtfA. Regulator of dormancy maintenance | (4) |
| AFUB_060680 | *atfB* | Transcription factor AtfB. Conidial stress response and secondary metabolism. Regulated by AtfA | (37), (38) |
| AFUB_016730 | *atfC* | Transcription factor AtfC. Regulated by AtfA | (38) |
| AFUB_002350 | *velB* | Asexual developmental regulator and velvet protein VelB. Required for conidial maturation | (10), (11), (33) |
| AFUB_067940 | *vosA* | Asexual developmental regulator and velvet protein VosA | (11), (33) |
| AFUB_041990 | *mybA* | Transcription factor MybA. Involved in conidiation, conidial cell wall biosynthesis, trehalose biosynthesis, and conidial viability | (31), (12), (35) |
| AFUB_101360 | *dprA* | Dehydrin protein DrpA. Involved in conidial stress response and HOG-MAPK. Regulated by AtfA | (40) |
| AFUB_095090 | *con-10* | Conidiation-specific protein Con-10 (ConJ ortholog). Regulated by AtfA | (70), (4) |
| AFUB_094400 | *catA* | Conidial specific catalase; important in response to reactive oxygen species and development. Regulated by AtfA | (41), (4) |
| AFUB_040120 | *cspA* | Conidial cell wall protein CspA; important in conidial cell wall structure and trehalose biosynthesis. Regulated by MybA | (39) |
| AFUB_013160 | *ccpA* | Conidial cell wall protein A CcpA. Important in the conidial surface structure | (71) |
| AFUB_089470 | *tps3* | Predicted alpha, alpha-trehalose phosphate synthase subunit Tsp3. Involved in trehalose biosynthesis | (72) |
| AFUB_037920 | *ace2* | Transcription factor Ace2. Positive regulator of asexual sporulation and spore wall assembly. Important in conidial pigmentation and virulence | (73) |
| **50°C conidia upregulated sDEGs** |  |  |  |
| AFUB_035330 | *nsdD* | Transcription factor NsdD. Positive regulator of sexual sporulation and secondary metabolism. Regulated by MAT1-1 | (22), (46),<br>(21) |
| AFUB_031050 | *fluG* | Transcription factor FluG. Upstream negative regulator of asexual conidiation initiation | (74) |
| AFUB_031740 | *flbB* | Transcription factor FlbB. Upstream negatively regulator of asexual conidiation initiation | (74) |
| AFUB_012620 | *fadA* | Guanine nucleotide binding protein (G-protein), alpha subunit FadA (GpgA homolog). Positive regulator of vegetative growth, regulated by MAT1-1 | (22) |
| AFUB_099650 | *fhbA* | Nitric oxide (NO) response FhbA. Contains MAT1-1 binding motif. Regulator of sexual reproduction | (22) |
| AFUB_066820 | *nosA* | Transcription factor NosA. Roles in cleistothecium development and ascospore formation | (48) |
| AFUB_016640 | *rodB* | Conidial hydrophobin, RodB. Contains MAT1.1 binding motif | (22), (29) |
| AFUB_023920 | *stuA* | APSES-domain containing transcription factor StuA. Hyphal competency switch, repressor of *brlA* during conidiation initiation. Positive regulator of sexual development | (29) |
| AFUB_028890 | *medA* | Transcription factor MedA. Important in phialide formation during conidiation | (75) |
| AFUB_012300 | *niaD* | Nitrate reductase NiaD. Regulated by FhbA | (22), (46) |
| AFUB_001420 | *noxA* | NADPH oxidase NoxA. Positive regulator of cleistothecium development | (49) |
| AFUB_066880 | *velC* | Velvet regulator VelC. *Aspergillus nidulans* VelC ortholog positively regulates sexual development | (29), (76) |
| AFUB_045170 | *phiA* | Cell wall protein PhiA. Role in asexual and sexual sporulation, regulated by StuA | (29) |
| AFUB_078580 | NA | Predicted roles in cleistothecium development | NA |
| AFUB_017010 | *het2* | Heterokaryon incompatibility | (56) |
| AFUB_028340 | *het3* | Heterokaryon incompatibility | (56) |
| AFUB_046770 | *het4* | Heterokaryon incompatibility | (56) |
| AFUB_045110 | *het5* | Heterokaryon incompatibility | (56) |
| AFUB_078080 | *het6* | Heterokaryon incompatibility | (56) |
| AFUB_085570 | *het7* | Heterokaryon incompatibility | (56) |
| AFUB_080400 | *het8* | Heterokaryon incompatibility | (56) |
| AFUB_080180 | NA | Predicted heterokaryon incompatibility | NA |
| AFUB_051520 | NA | *N. crassa* CWR-1 (NCU01380) ortholog. Predicted protein of unknown function | (55) |
| AFUB_030130 | NA | *N. crassa* CWR-2 (NCU01382) ortholog. Predicted protein of unknown function. | (55) |
| AFUB_078810 | *mpkB* | Mitogen-activated protein (MAP) kinase MpkB. *N. crassa* MAK-2 (NCU02393) ortholog | (53) |
| AFUB_046510 | *cbhC* | Cellulose binding protein CbhC. *N. crassa* NCU07190 *doc* locus (self-recognition) ortholog | (53) |
| AFUB_072440 | *fso1* | Fso1, protein of unknown function. *N. crassa* SOFT (NCU02794) ortholog | (53) |
| AFUB_061660 | NA | *N. crassa* HAM-5 (NCU01789) ortholog. Predicted protein of unknown function | (53) |
| AFUB_058790 | *rasA* | RAS signaling pathway GTPase RasA; regulates development and cell wall integrity | (77), (78) |

a Upregulated significant differentially expressed genes (sDEGs) from 37°C versus 50°C conidial differential analyses listed in Fig. 5D heatmap.
b Gene ID from FungiDB for *Aspergillus fumigatus* strain A1163.
c Description simplified from FungiDB predicted protein function and literature references.

### 50°C conidia express higher levels of MAT1-1 targets, sexual developmental regulators, and anastomosis homologs

We reasoned that the 50°C conidial sDEGs would uncover clues on dormancy maintenance in 50°C conidia. The top biological GO processes (30) of sDEGs upregulated in 50°C conidia versus 37°C included cytoskeleton synthesis and organization, cell cycle, nuclear processes, and fumagillin biosynthesis (Fig. S4C and D), processes in which take part in the formation of sexual spores (ascospores) within the asci in sexual reproductive structures (cleistothecia) or have been associated with sexual reproduction (21). Previous work found that MAT1-1 regulates fumagillin BGCs (21), and that fumagillin metabolites are produced during both sexual development and cytoskeletal remodeling (42). Several genes including members of the fumagillin biosynthesis and BGC and were upregulated in 50°C conidia, including the positive regulator of fumagillin biosynthesis FumR (AFUB_086150) (43), (44) (Fig. 5A, Tables 3 and S3). Additionally, chromatin remodeler and global regulator of secondary metabolism LaeA (AFUB_014200) (45) was upregulated, suggesting that LaeA may play an important role in the production of fumagillin and gliotoxin associated transcripts observed in the upregulated 50°C conidial sDEGs (26), (27).

**TABLE 3.**
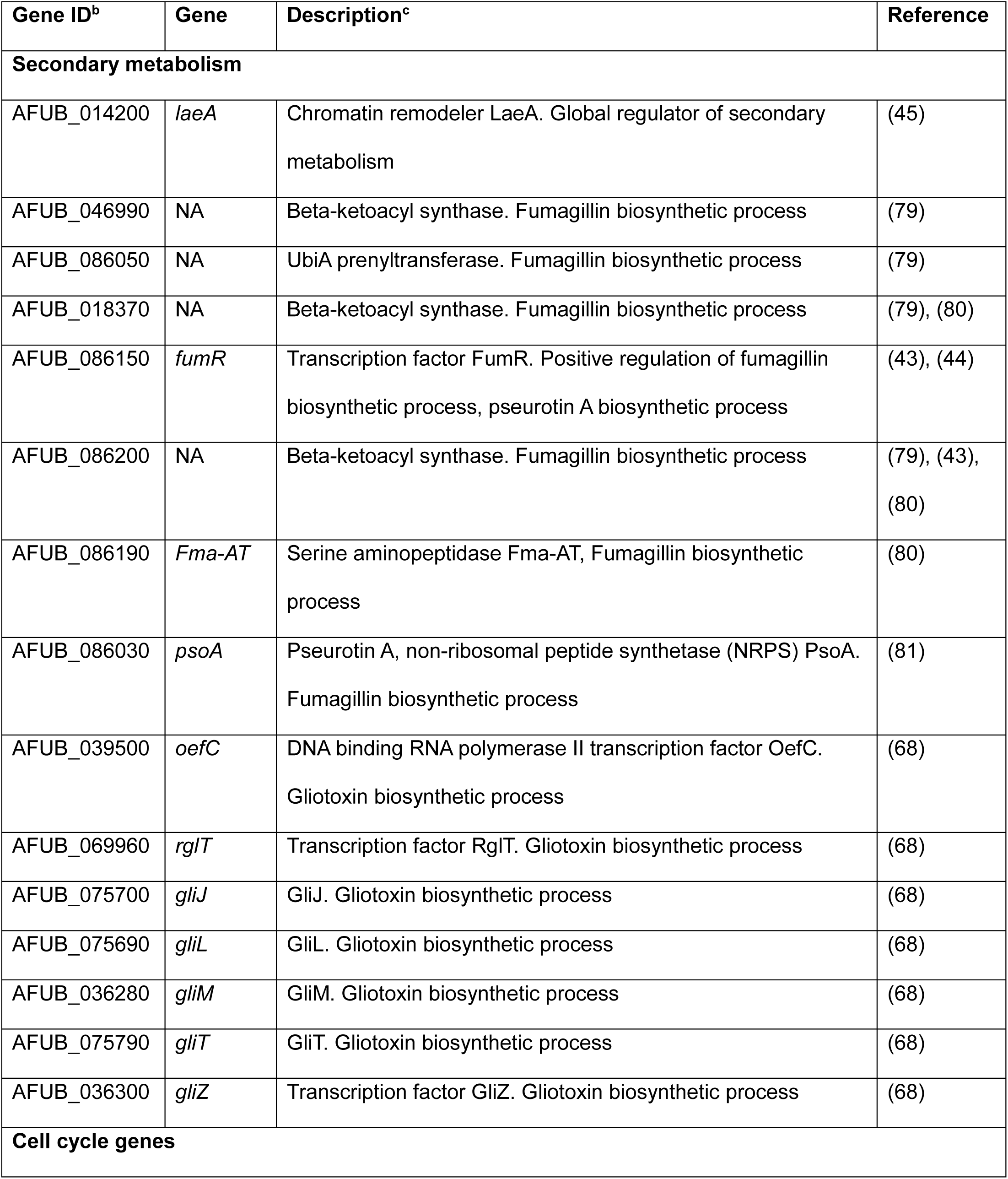
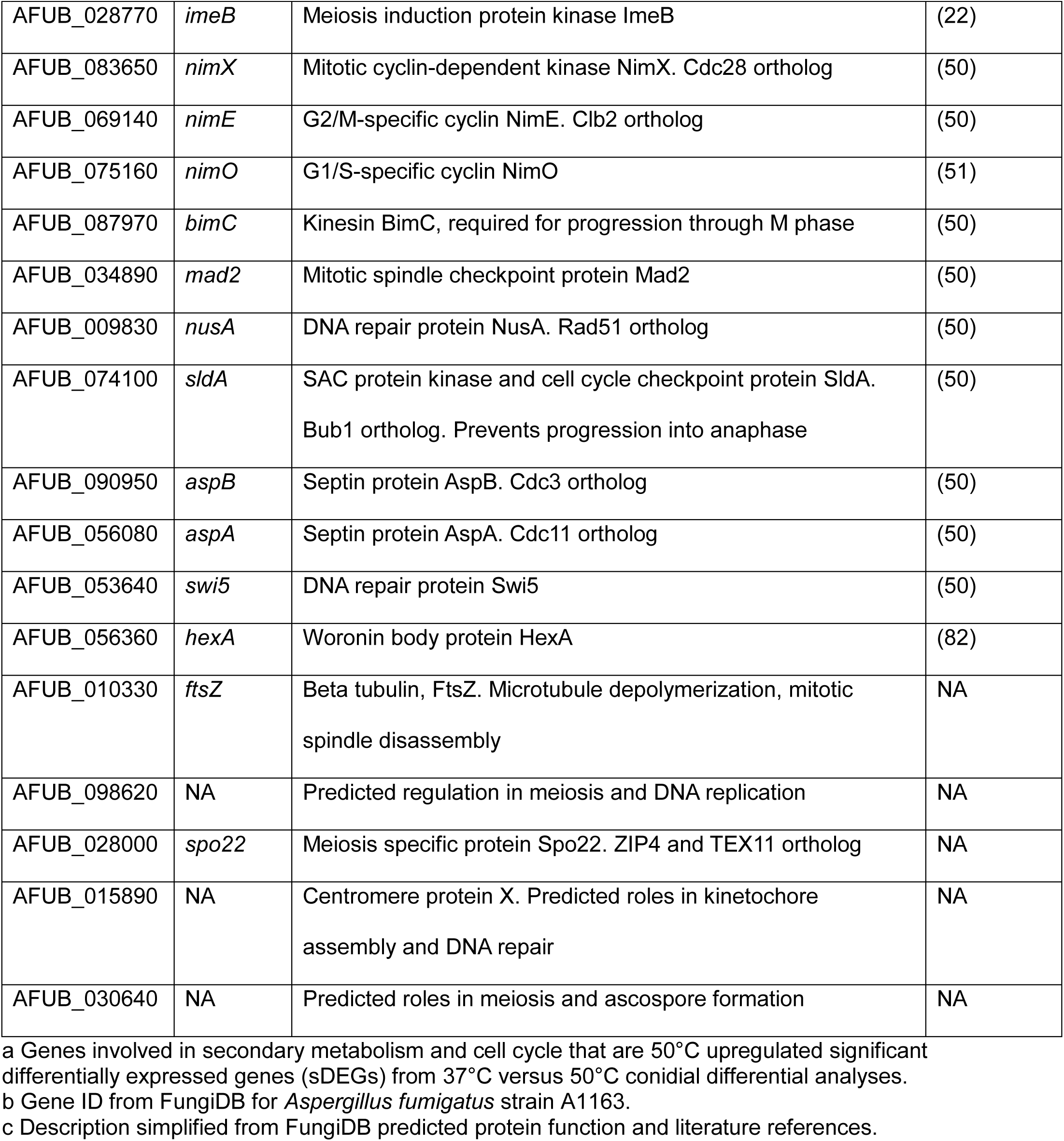
50°C conidia upregulated sDEGs important in secondary metabolism and cell cycle^a^.

Though MAT1-1 is typically involved in sexual development rather than asexual development, the upregulation of MAT1-1 in 50°C conidiophores (Fig. 4A-C, Tables 1 and S3) suggested that downstream targets of MAT1-1 might be upregulated in 50°C conidia. Indeed, many downstream targets of MAT1-1 were differentially upregulated in 50°C conidia versus 37°C conidia, including numerous genes important is sexual reproduction and cell cycle genes (Fig. 5A-D, Tables 2 and 3).

Several genes upregulated in 50°C conidia contain MAT1.1 binding motifs and are associated with sexual development including *nsdD* (AFUB_035330) (22), *fhbA (*AFUB_099650) (46), *rodB* (AFUB_016640) (18), and *fadA* (AFUB_012620) (22) (Fig. 5C and D, Table 2). Although not directly driven by MAT1-1, transcript levels of numerous positive regulators of cleistothecium formation were also upregulated in 50°C conidia, including transcription factor StuA (AFUB_023920) (29) (47), transcription factor NosA (AFUB_066820) (48) and NADPH oxidase NoxA (AFUB_001420) (49). MAT1-1 has also been reported to regulate cell cycle genes (21), and the 50°C conidial sDEGs included numerous regulators of cell cycle progression, including mitotic cyclin-dependent kinase NimX (AFUB_083650), G2/M-specific cyclin NimE (AFUB_069140), and G1/S-specific cyclin NimO (AFUB_075160) (50), (51) (Tables 3 and S3). Based on work in *A. nidulans*, it is assumed that dormant conidia in *A. fumigatus* are arrested in the G1 phase of the cell cycle (52), however the abundant upregulation of cell cycle genes suggests nuclei in 50°C conidia are not fully arrested in G1. Consistent with the idea that nuclei in 37°C versus 50°C are in different states, ImageStream flow cytometry analyses showed a nuclear fluorescent signal that was less intense in 50°C conidia, indicating that nuclei are less compact compared to nuclei of 37°C conidia (Fig. S1C and D, Table 3). Strikingly, transcripts of the meiosis induction protein kinase ImeB (AFUB_028770) (22) and meiotic regulator Spo22 (AFUB_028000) were also upregulated, suggesting that meiotic processes are stimulated in 50°C produced conidia (Tables 3 and S3). The abundance of MAT1-1 targets, including known sexual development genes and cell cycle genes, are consistent with 50°C conidiophores’ upregulation of *MAT1-1*, the master regulator of sexual development.

In addition to sexual development genes, many regulators of anastomosis (hyphal fusion) were also upregulated in 50°C conidia versus 37°C conidia (Fig. 5D, Tables 2 and S3). In fungi, cell wall remodeling and hyphal fusion are tightly linked with self/non-self recognition, however, the underlying cell-cell communication is not well defined in *A. fumigatus*. In *Neurospora crassa*, MAK-2 (NCU02393), SOFT (NCU02794) and HAM-5 (NCU01789) govern cell-cell communication by oscillating at hyphal tips before fusion (53), (54). To our surprise, the *A. fumigatus* orthologs *mpkB* (AFUB_078810), AFUB_072440, and AFUB_061660, respectively) were upregulated in 50°C conidia (Fig. 5D, Tables 2 and S3). *N. crassa* cell wall remodeling proteins CWR-1 (NCU01380), and CWR-2 (NCU01382) serve as checkpoints during cellular fusion (55). *A. fumigatus* CWR-1 orthologs AFUB_051520 and AFUB_030130 were also upregulated. We also found that 7 previously identified heterokaryon incompatibility (*het*) genes were upregulated in 50°C conidia (Fig. 5D, Tables 2 and S3). *Het* genes play roles in determining post-fusion compatibility by self-recognition (7), (8). Although anastomosis is a prerequisite for sexual reproduction, these genes are not known to be regulated by mating-type loci and appear to independently drive vegetative compatibility (21), (56). The abundance of anastomosis homologs and heterokaryon compatibility genes in the 50°C conidial sDEGs suggests that 50°C conidia may be transcriptionally predisposed for cellular fusion during germination. Furthermore, these results suggest that the difference in conidial size and dormancy breaking in conidia produced at 37°C versus 50°C are transcriptionally driven by distinct developmental pathways.

## DISCUSSION

In previous work, we showed that *A. fumigatus* asexual spores (conidia) produced at 37°C were smaller and broke dormancy more slowly than those produced at 50°C (24) and that these differences correlated with transcriptional changes (25). In this work, we investigated key landmarks in the asexual development pathway that leads to production of conidia at 37°C and 50°C. Our temperature shift experiments showed that the conidial phenotype influenced by environmental temperature is determined during a narrow developmental window in late-stage conidiophore development (Fig. 2A-C). Our RNA-seq analyses showed that incubation at 37°C during the late-conidiophore developmental window leads to increased expression of *brlA*, the master regulator of asexual development, in conidiophores and of BrlA’s downstream targets in conidia. In contrast, incubation at 50°C during the late-conidiophore developmental window leads to increased expression of *MAT1-1*, the master regulator of sexual development, in conidiophores and of MAT1-1’s downstream targets along with anastomosis genes in conidia.

Previous work in *A. nidulans* and *A. fumigatus* characterizing asexual development at 37°C defined the paradigmatic asexual development pathway driven by the BrlA transcription factor (11), (34), (13). Our finding that *brlA* and alkaloid BGC genes were upregulated in conidiophores produced at 37°C is consistent with these previous studies of asexual development (14), (15), (16). In contrast, our finding that *MAT1-1* transcripts were upregulated in conidiophores made at 50°C was unexpected since upregulation of MAT1-1 and fumagillin/pseurotin and gliotoxin BGC genes has been previously associated with sexual reproduction (12), (21). Our finding of a switch that appears to toggle between asexual, parasexual, and sexual lifecycles in response to temperature has important implications for the diversity and fitness of *A. fumigatus* in natural and agricultural environments and for the spread of antifungal resistance through populations.

Based on our results, we propose a model shown in Figure. 6 and described below in which *A. fumigatus* follows radically different developmental pathways when incubated at 37°C versus 50°C.

**Fig 6.**
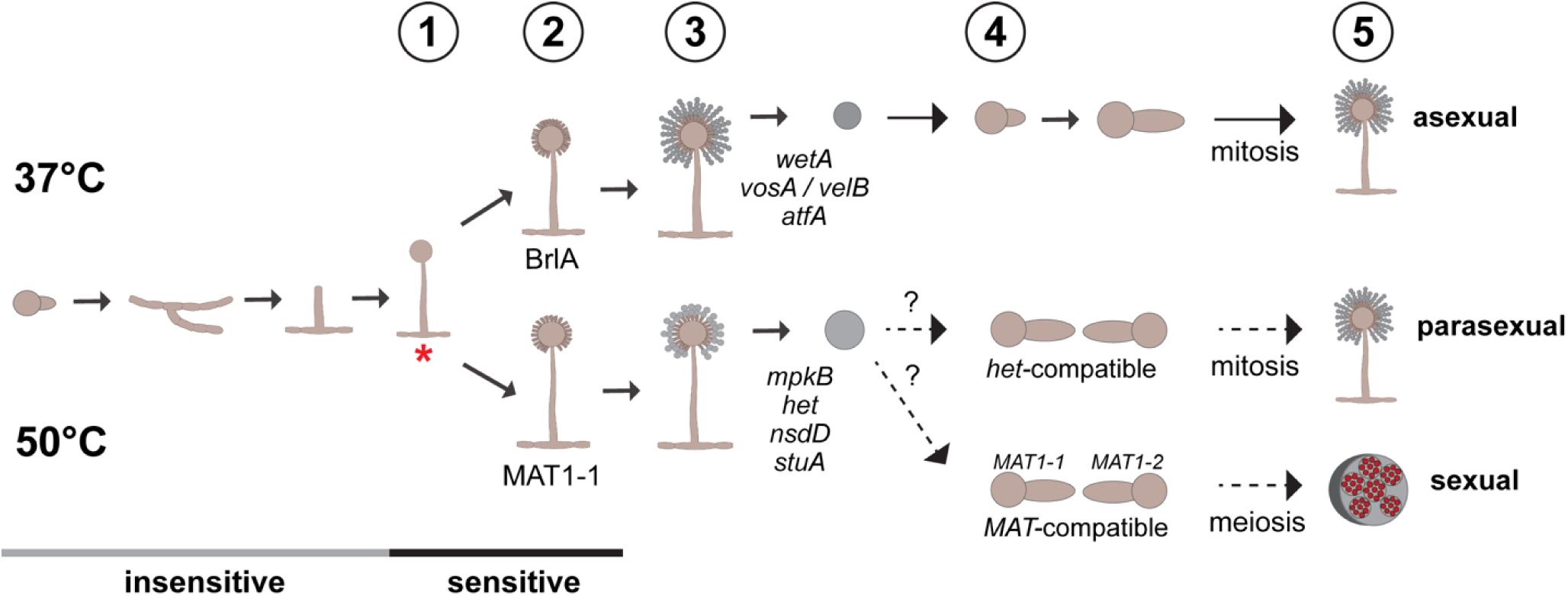
Model for impact of conidiation temperature on conidial transcriptional programs and fitness of progeny. (**1**) During conidiophore development vesicles sense the environmental temperature. (**2**) Environmental signals trigger transcription of specific developmental regulators in the conidiophore. **(3)** Developmental regulators in the conidiophores trigger transcription of their downstream targets and these transcripts are packaged in conidia. **(4)** Conidia germinate and carry out their transcriptional programs for asexual, parasexual or sexual development depending on the presence of suitable fusion partners. **(5)** Progeny of environmentally-primed conidia are better-suited for the environment that was present when the parental conidiophore formed. Grey and black line indicate stages of asexual development that are insensitive or sensitive to environmental temperature, respectively. Red asterisk indicates the phenotypic commitment point after which changes in conidiophore environmental temperature do not change the resulting conidial program. See Discussion for more detail.

**Model step 1: Late-stage conidiophores sense environmental temperature.** Through temperature shift experiments tracking morphology at defined landmarks, we found that the conidiophore is sensitive to environmental temperature only during late-stage conidiophore development (Fig. 2A and 2B, Fig. 6). Temperature shift during vegetative growth, the emergence of aerial hyphae (before vesicle formation), or the emergence of conidial chains (after phialide formation) did not result in significant morphological alterations to the conidia produced compared to controls. Our results suggest that late conidiophore development is the environmentally sensitive developmental window and the most critical time in the asexual lifecycle in determining the phenotype of future conidia. Our results also suggest that because the phenotype is not alterable once it has been established, there is a phenotypic commitment point before conidia emerge. Importantly, this pattern holds true whether beginning at 37°C and shifting to 50°C (Fig. 2A) or beginning at 50°C and shifting to 37°C (Fig. 2B), suggesting that the critical developmental period is not limited to standard 37°C laboratory grown conditions. Unfortunately, we could not perfectly synchronize conidiophore development, and so we were unable to pinpoint the environmentally sensitive phase and commitment point more exactly between vesicle formation and the emergence of conidia.

We show that environmental temperature is sensed late in conidiophore formation, during vesicle and/or phialide formation (Fig 6, step 1), and speculate that late-stage conidiophores are sensitive to environmental input across a range of temperatures and conditions. Future studies will address whether other temperatures or environmental conditions during late-stage conidiophore production can similarly impact the conidia produced

**Model step 2: Regulators that direct distinct developmental pathways are upregulated in conidiophores in response to environmental temperature.** Our RNA-seq experiments showed that incubation at 37°C versus 50°C induces upregulation of transcription factors in conidiophores that are known to direct development toward asexual reproduction (37°C, BrlA) or sexual reproduction (50°C, MAT1-1) (Fig. 4, Fig 6, step 2). Based on conidial transcripts described in model step 3 below, we speculate that there might be other developmental regulators upregulated in conidiophores in response to temperature. We also speculate that environmental input across a range of temperatures and conditions might induce upregulation of other regulators to direct other developmental pathways.

**Model step 3: Downstream targets of conidiophore developmental regulators are upregulated in conidia.** Analysis of conidial transcriptomes produced at 37°C versus 50°C was consistent with BrlA regulation or MAT1-1 regulation, respectively (Fig. 4 and 5; Tables 2 and S3). Conidia produced at 37°C showed significant upregulation of downstream targets of BrlA, including regulators of conidial viability WetA, MybA, VelB, and VosA, as well as trehalose biosynthesis and cell wall synthesis (12), (35). Essential players of the mitogen-activate protein kinase (HOG-MAPK) pathway were also upregulated in 37°C conidia, including the MAPK SakA (AFUB_012420) (36), (35). Conidial dormancy is regulated by AtfA which inhibits germination and controls 63% of all conidial transcripts (4). Importantly, *atfA* and numerous downstream targets of AtfA were upregulated in conidia produced at 37°C, including *atfB* (AFUB_060680) (37), *atfC* (AFUB_016730) (38), *cspA* (AFUB_040120) (39), *drpA* (AFUB_101360) (40), *catA* (AFUB_094400) (41), and *con-10* (AFUB_095090). Previous studies showed that conidia produced at higher temperatures have an altered cell wall composition (26), (27), consistent with the idea that conidial transcripts direct production of conidia with different cell wall composition when produced at 37°C versus 50°C.

Conidia produced at 50°C displayed upregulation of known MAT1-1 targets, including fumagillin/pseurotin and gliotoxin BGCs, sexual reproduction genes, and cell cycle genes. Dormant conidia are presumed to be arrested in the G1 phase of the cell cycle (52). The striking abundance of cell cycle genes upregulated in 50°C conidia (Fig. 5B) suggests that the cell cycle might be more active compared to 37°C conidia. Indeed, the nuclei in 50°C conidia appeared to be more diffuse and completed mitosis more rapidly than 37°C conidia (Fig. S1C). This difference in cell cycle could also be in preparation for meiosis as the meiosis induction protein kinase ImeB (22) was upregulated in 50°C conidia.

In addition to known targets of MAT1-1, we also found that multiple orthologs of genes involved in cell fusion (anastomosis) not known to be regulated by mating type loci were upregulated in conidia produced at 50°C (21), (56). The early steps in anastomosis have not been determined in *A. fumigatus*, but in *N. crassa* MAK-2, SOFT, and HAM-5 govern cell-cell communication by oscillating at hyphal tips before fusion (53), (54), (9). During cellular fusion in *N. crassa*, cell wall remodeling proteins CWR-1 (NCU01380), and CWR-2 (NCU01382) serve as fusion checkpoints at cellular contact (55). In *A. fumigatus* 50°C conidia, orthologs of MAK-2, SOFT, HAM-5, CWR-1 and CWR-2 are all upregulated (Fig. 5D, Tables 2 and S3). Also in conidia produced at 50°C, we observed significant upregulation of 7 putative heterokaryon incompatibility (*het*) genes (Fig. 5D, Tables 2 and S3) (7), (8). We speculate that these anastomosis and heterokaryon incompatibility genes might be important for parasexual development We show that conidia derived from 37°C conidiophores are upregulated in targets of BrlA, while conidia of 50°C conidiophores are upregulated in targets of MAT1-1 and genes important for anastomosis and heterokaryon incompatibility (Fig. 6, step 3). We speculate that conidia that emerge from conidiophores produced under other environmental conditions might show upregulation of other distinct developmental pathways.

**Model step 4: Environmental temperatures during conidiophore formation transcriptionally prime conidia for future asexual, sexual, or parasexual reproduction.** We show that conidiophore formation at 37°C gives rise to conidia that germinate more slowly and are upregulated in transcripts important for maintaining conidial dormancy and viability, as well as asexual development. Conversely, conidiation at 50°C leads to larger conidia that germinate rapidly and are upregulated in transcripts important for cell cycle progression, sexual reproduction and/or possibly parasexual development.

We propose that conidia are transcriptionally primed to maintain dormancy and germinate optimally in the environment to which they were exposed during late-stage conidiophore development. Specifically, we propose that conidia produced from 37°C conidiophores are primed for asexual development, and that conidia produced from 50°C conidiophores are primed for sexual or possibly parasexual development (Fig. 6, step 4). Although our results strongly suggest that conidia produced at 37°C or 50°C are primed for asexual or sexual development, respectively, we have only performed these experiments with the MAT1-1 wildtype strain A1163. To prove that 50°C conidiation primes for sexual reproduction we must repeat the experiments with mating compatible strains of opposite mating types. These experiments are underway.

**Model step 5: The progeny of transcriptionally primed conidia are more fit and adaptable in the environments that primed them.** *A. fumigatus* is a ubiquitous saprophyte found in soil and plant debris in the environment and is thermotolerant growing at temperatures up to 55°C. This thermotolerance allows *A. fumigatus* to grow well at human body temperature (37°C) and even in intermediate and outer layers of compost where temperatures routinely reach 50°C or higher (Fischer et al 1998) (57). *A. fumigatus* produces both conidia and ascospores at temperatures ranging from 25°C to 37°C (23). Conidia break dormancy and germinate optimally at 30-37°C. Ascospores require a heat shock of 60-80°C for efficient germination, while conidia are killed at such high temperatures (23).

We propose that the rapid germination observed in conidia produced from 50°C conidiophores (Fig. 1E) enables more rapid growth through the surrounding substrate where compatible partners might be encountered that have the opposite mating type (sexual) or compatible *het* loci (parasexual). We speculate that upon germ tube extension, if the hyphae with the opposite mating type are encountered, hyphal fusion and sexual reproduction will occur, resulting in the production of ascospores within a cleistothecium. Over time, the cleistothecium will break down releasing ascospores. If ascospores later encounter 60-80°C temperatures they germinate giving rise to genetically varied progeny. If the germ tube encounters a hypha that is not the opposite mating type, but is compatible at *het* loci, hyphal and nuclear fusion and parasexual development will occur resulting in the production of conidia.

### Concluding thoughts

As others have noted, the environmental conditions in compost are similar to conditions that favor *A. fumigatus* sexual reproduction and the temperature gradient in compost piles (20-70°C) includes internal regions hot enough to furnish the required heat shock for germination of any ascospores produced (23), (9), (5), (6). Recent work has uncovered extremely high levels of meiotic recombination in *A. fumigatus* and this recombination is thought to have assisted in the generation and spread of alleles causing resistance to azole antifungals (56). Azole resistant *A. fumigatus* has been found frequently in agricultural environments including in soil, compost, and plant debris (58). Our results strongly support the previously proposed idea that compost might be the environmental niche for *A. fumigatus* sexual reproduction (9). Because a temperature above 50°C kills conidia and most competing microbes, but not ascospores, sexual development at 50°C would give *A. fumigatus* a major survival advantage as well as the potential for better adaptation of progeny because of the genetic diversity of ascospores.

Our results strongly suggest that the temperature-induced programs determined during late-stage conidiophore development are transcriptionally regulated; in addition, our results suggest that post-transcriptional regulation might also be important. Other sDEGs upregulated in 50°C conidiophores included an abundance of both non-coding RNAs (ncRNAs) and ribosomal biogenesis genes, including the 18S rRNA subunit (AFUB_090410), translation initiation factors IF6 (AFUB_053550), eIF-1A (AFUB_059560), and ribosome biogenesis proteins NOP2/RsmB (AFUB_050290), and Slx9 (AFUB_002540) (Table S3). Although ribosome biogenesis utilizes approximately 80% of cellular energy (59), increased ribosomal production drives cellular growth, stress response, and enhances environmental adaptation (60), consistent with altering growth regulation. Importantly, previous studies demonstrated that BrlA suppresses ribosome biogenesis, which is presumed to be useful in metabolic preservation during conidial production (17). Additionally, 50°C conidiophores showed upregulation of *cpcA* (AFUB_069420) (Fig 4C, Tables 1 and S3), whose protein product drives the cross-pathway control system (61) which shifts metabolic activities as well as amino acid-production in response to environmental stress. These results suggest that post-transcriptional processes might also be used during conidiophore development in response to environmental temperature to regulate distinct developmental. Future work will investigate the range of environmental temperatures and conditions that impact conidial development and the importance of non-transcriptional regulation in this process.

## MATERIALS AND METHODS

### Strains and conidial preparation

The *Aspergillus fumigatus* wild-type strain CEA10 was utilized for all analyses (62), (63). Conidia were inoculated to Aspergillus minimal media (AMM) (64), incubated for 3 days at 37°C, and harvested in 10 mL of DEPC treated water (Invitrogen). Isolated conidia were filtered 4x through 22-25 µm Miracloth (Sigma) and washed by centrifuging for 1 minute at 7,000 x g at 4°C and resuspended in 1 mL DEPC-treated water. Conidia were quantified via hemocytometer prior to all downstream analyses. Conidia were diluted to a common concentration of 1x10^8^ conidia/mL for all experiments.

### Radial growth analyses

Colony growth was observed by inoculating 1 µL of 1x10^8^ conidia/mL onto the middle of petri dishes containing AMM and incubating at 37°C or 50°C for 3-days. Colony diameter was measured every 24 hours. Three biological replicates were performed for each condition. Data were plotted and analyzed using GraphPad Prism software version 10.

### Defining developmental milestones

Conidia were inoculated onto pre-warmed solid AMM petri plates at a final concentration of 1x10^7^ conidia/plate and incubated at 37°C or 50°C. Growth was monitored every 4 hours via standard light microscopy. Wet mounts were analyzed for hyphal cells, and aerial conidiophore development was analyzed by slicing and removing a 0.5 x 2 cm rectangle cubes from the petri plates, placing them onto microscope slides, and analyzing the side-view of the aerial conidiophore growth at various timepoints. Three biological replicates were carried out with a minimum of 10 conidiophores observed per replicate. All three replicates gave very similar patterns.

### Temperature shift experiments

Conidia were inoculated onto pre-warmed solid AMM petri plates at a final concentration of 1x10^7^ conidia/plate and incubated at 37°C or 50°C. Growing cultures were transferred at defined milestones from 37°C to 50°C, or vice versa, and resulting conidia produced after 96 hours were isolated and size was analyzed using flow cytometry. Three biological replicates were carried out per experiment.

### Isolation of hyphae

200 µL of 1x10^8^ conidia/mL *Aspergillus fumigatus* CEA10 conidia were inoculated into 500mL Erlenmeyer flasks containing 200 mL liquid AMM. Cultures were incubated for 12 hours at either 37°C or 50°C with shaking at approximately 150 rpm. Hyphae were filtered through 22–25 µm Miracloth, washed with DEPC-treated water, and RNA extraction was immediately carried out.

### Isolation of conidiophores

200 µL of 1x10^8^ conidia/mL *Aspergillus fumigatus* CEA10 conidia were inoculated into 40 mL melted AMM (cooled to 50°C) and poured into 60 mm x 15 mm petri dishes (ThermoFisher Scientific). The conidia/media mixture was allowed to harden at room temperature for 15 minutes, and then 4 mL of melted H20 agar was layered directly on top to physically isolate developing conidiophores from the underlying hyphae. After 28 and 38 hours of incubation at 37°C or 50°C respectively, the conidiophores were isolated by gently shaving of the top conidiophore layer using a razor blade. Resulting isolated conidiophores were gently scraped from the plate and viewed microscopically to ensure purity. The resulting conidiophore shavings were transferred into a 15 mL falcon tube for RNA extractions.

### RNA-extractions

Isolated cells were immediately lysed by resuspending cells in 100 µL 1xTE buffer (Invitrogen) in a 15 mL falcon tube. Four 2.33 mm beads (BioSpec Products) and 0.1mL (100 mg) of 425-600 µM beads (Sigma) were added and vortexed on the highest setting for 2 minutes. RNA extractions were carried out utilizing TRIzol according to the manufacturer’s protocol (Invitrogen).

### RNA-sequencing

RNA-sequencing was performed by Plasmidsaurus on RNA extracted from isolated conidia, hyphae and conidiophores produced at 37°C or 50°C (Fig. S2). Three biological replicates of each condition were sequenced. Quality of the fastq files was assessed using FastQC v0.12.1. Reads were then quality filtered using fastp v0.24.0 with poly-X tail trimming, 3’ quality-based tail trimming, a minimum Phred quality score of 15, and a minimum length requirement of 50 bp. Quality-filtered reads were aligned to the reference genome using STAR aligner v2.7.11 with non-canonical splice junction removal and output of unmapped reads, followed by coordinate sorting using samtools v1.22.1. PCR and optical duplicates were removed using UMI-based deduplication with UMIcollapse v1.1.0.

Alignment quality metrics, strand specificity, and read distribution across genomic features were assessed using RSeQC v5.0.4 and Qualimap v2.3, with results aggregated into a comprehensive quality control report using MultiQC v1.32. Gene-level expression quantification was performed using featureCounts (subread package v2.1.1) with strand-specific counting, multi-mapping read fractional assignment, exons and three prime UTR as the feature identifiers, and grouped by gene_id. Final gene counts were annotated with gene biotype and other metadata extracted from the reference GTF file. Sample-sample correlations for sample-sample heatmap and PCA were calculated on normalized counts (TMM, trimmed mean of M-values) using Pearson correlation. Differential analyses of RNA-seq data were carried out using DESeq2 v.1.50.2 (BiocManager v1.30.27) in R (65). Low read counts were filtered out (< 5). All parameters were performed according to standard methods. Resulting differentially expressed genes (DEGs) were filtered based upon adjusted p-value (padj < 0.05) and |Log2FoldChange| > 2.

### Gene ontology analyses

Gene ontology (GO) enrichment analyses were performed using FungiDB (30). Fisher’s exact tests were utilized for all GO analyses (p-value cutoff of 0.05), ranked from highest to lowest p-value. All graphs and plots were generated using RStudio version 2025.05.0+496 and Prism version 10 (GraphPad Software). BiocManager v1.30.27, clusterProfiler v4.18.4, dplyr v.1.1.4, ggplot2 v.4.0.1, tidyr v.1.3.2 were utilized for top 10 GO enrichment plots.

### RNA-seq data interpretation

All graphs and plots for RNA-seq and gene ontology were generated using RStudio version 2025.05.0+496 and Prism version 10 (GraphPad Software). UpSet plots were created using ComplexHeatmap v2.26.0 (BiocManager v1.30.27) and UpSetR v1.4.0. For UpSet plot construction, the resulting normalized counts from the differential analyses after filtering out low reads were used. To get the total numbers of genes expressed by each cell type (hyphae, conidiophores, conidia), individual samples of each cell type were run through DESeq2 v.1.50.2 (BiocManager v1.30.27) in R (65) to filter out low reads (<5).

### Flow cytometry analyses

Flow cytometry was performed at the Center for Tropical and Emerging Global Diseases Cytometry Shared Resource Laboratory at the University of Georgia. The Cytek® Amnis® ImageStream®X Mk II Imaging Flow Cytometer (Cytek Bioscience) was used for all flow cytometry experiments as previously described (66), (67) with slight modifications. Two gates were created using 2-15 µm standards (ThermoFisher Scientific) to filter out non-focused cells and doublets or debris, respectively (Fig. S1B-D). Gates were further optimized for *A. fumigatus* using the IDEAS analysis program (version 6.2). *A. fumigatus* ungerminated conidia (in water) or 5-hour germinated conidia (AMM, 37°C) were normalized to 1x10^8^ conidia/mL for flow cytometry experiments. Microscope images and data were collected on 10,000 events per sample, and 3 biological replicates were pooled per sample. The area data in approximate µm^2^ resulting from the approximate 1,500-3,000 events were further analyzed using GraphPad Prism software version 10. Statistical analyses on area or fluorescence data from gated events were performed using Mann-Whitney nonparametric t tests to compare the distribution amongst sample groups (two-tailed). P-values are indicated by stars *** p<0.001, **** p<0.0001.

## ACKNOWLEDGEMENTS

This work was supported by National Institutes of Health award 1R03AI176262-01 to MM.

---

**Fig S1.**
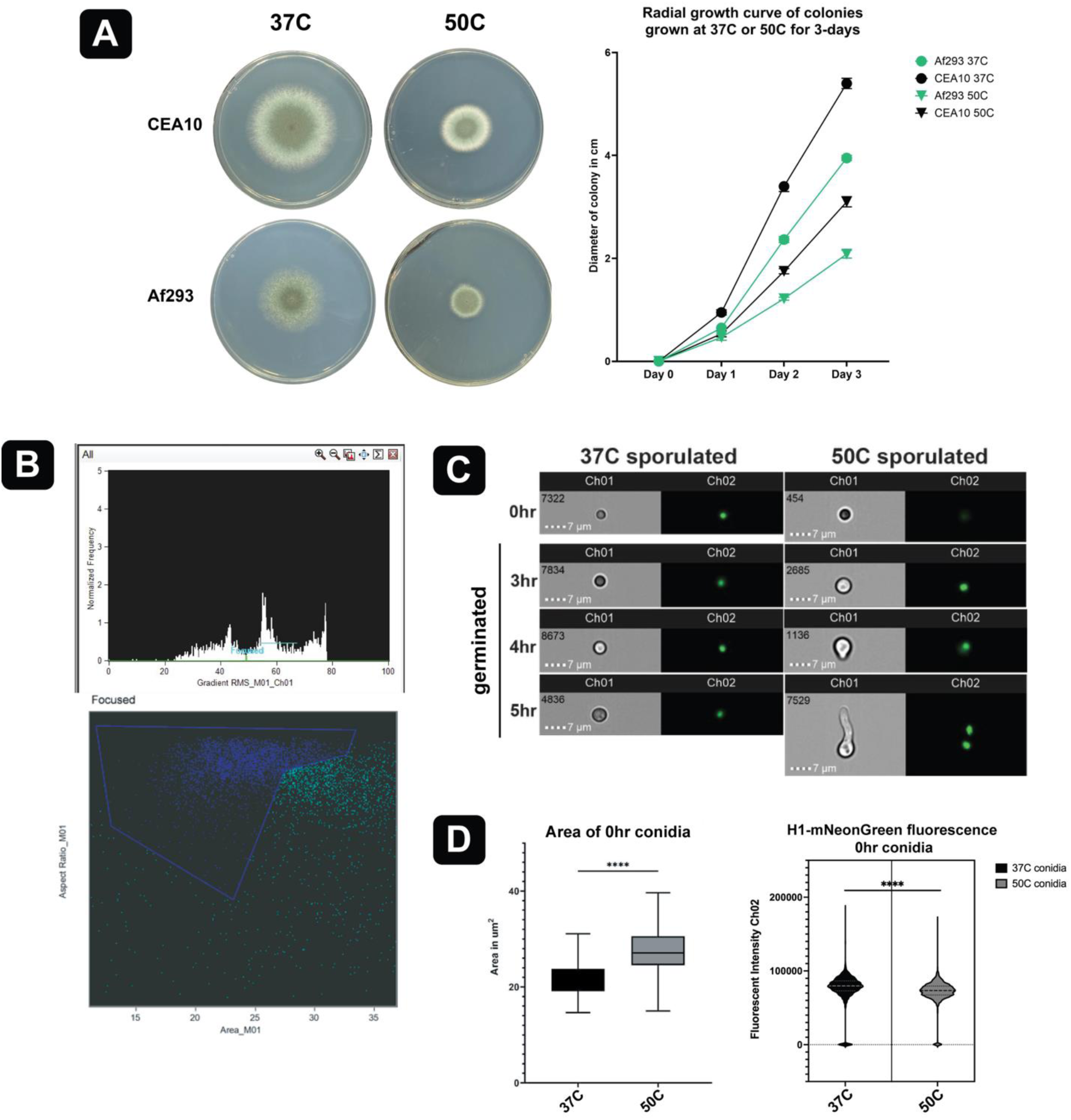
(**A**) 3-day colony growth analyses for strains CEA10 and Af293 grown at 37°C (green) and 50°C (black). n=3. **B**-**D**. Flow cytometry experiments utilizing the ImageStream®X Mk II Imaging Flow Cytometer (Cytek® Amnis®). (**B**) Gates created based upon imaging data of the fully detailed events using the IDEAS software program. IDEAS screenshots of flow cytometry gating parameters for *A. fumigatus* conidia (in water). Ch01 shows brightfield, Ch02 displays histone H1-mNeonGreen (nuclear) fluorescence. (**C**) ImageStream images of dormant (0 hr) conidia produced at 37°C (black) or 50°C (grey) or 3-5 hr germinated cells at 37°C, respectively. Images are representative of the largest, in-focus cells of the population. (**D**) Box plot (left) indicates the area range of dormant conidia produced at 37°C and 50°C (n = 1,500-3,000 events per sample) in μm^2^. The middle line represents the median. Mann-Whitney nonparametric test resulting p-value of <0.0001 indicated by stars ****. Violin plots (right) indicate fluorescence intensities of dormant conidia produced at 37°C or 50°C (n = approximately 1,500-3,000 events per sample, Ch02, mNeonGreen). Mann-Whitney nonparametric test resulting p-value of <0.0001 indicated by stars ****.

**Fig S2.**
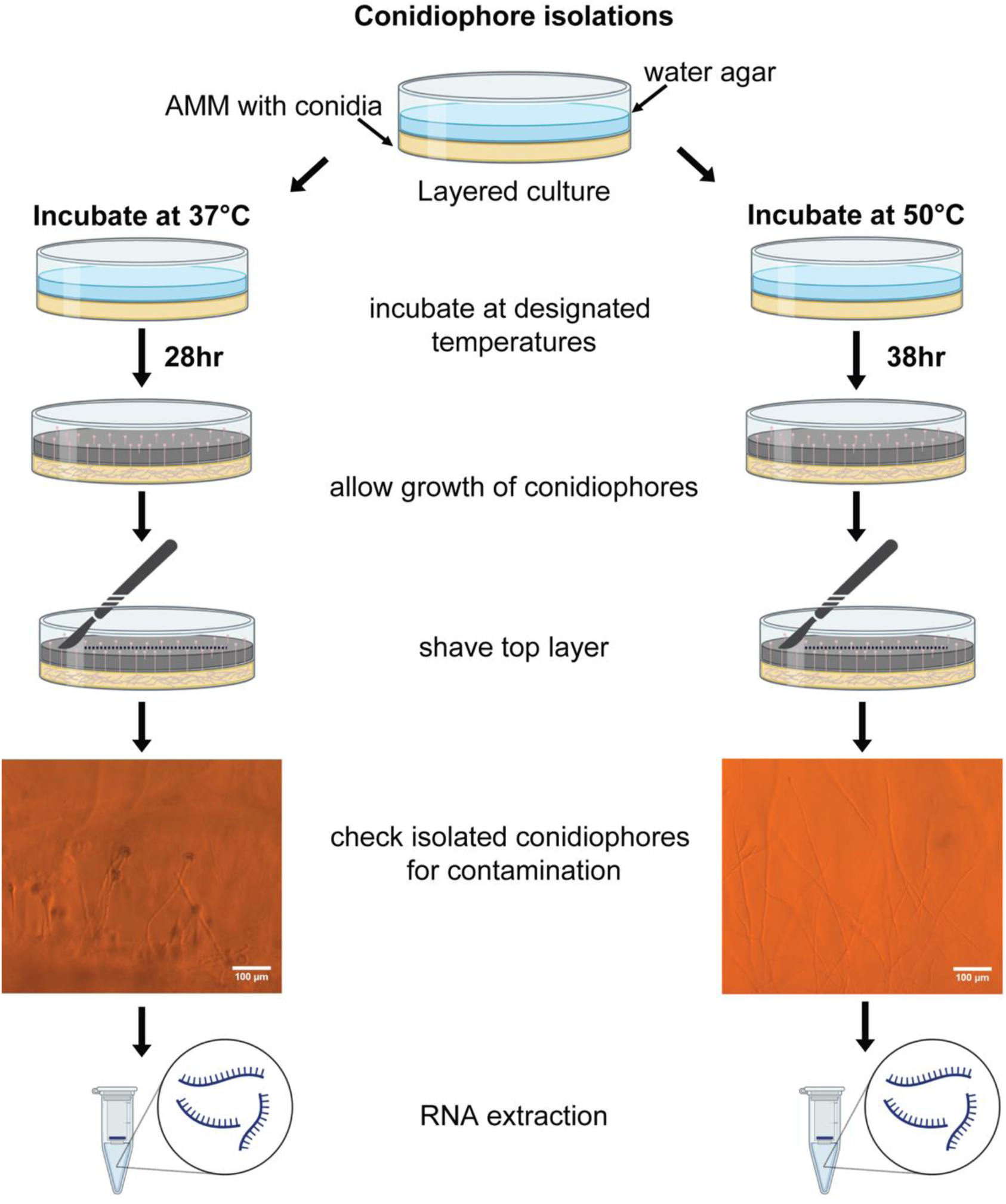
Conidiophore isolation from hyphae and conidia protocol overview. Conidiophores were physically isolated by inoculating a mixture of 2x10^7^ conidia/mL in 40mL melted AMM, mixed and poured directly into 60x15 mm petri dishes. After solidification, 4mL water agar (0.7%) was layered on top and allowed to solidify. Petri plates were incubated at 37°C or 50°C until conidiophores grew through the top agar at the specified time points: 37°C conidiophores were isolated after 28-hours, and 50°C conidiophores were isolated 38-hours after germination. Wet mounts were examined microscopically for each condition to ensure conidiophore purity prior to RNA extractions.

**Fig S3.**
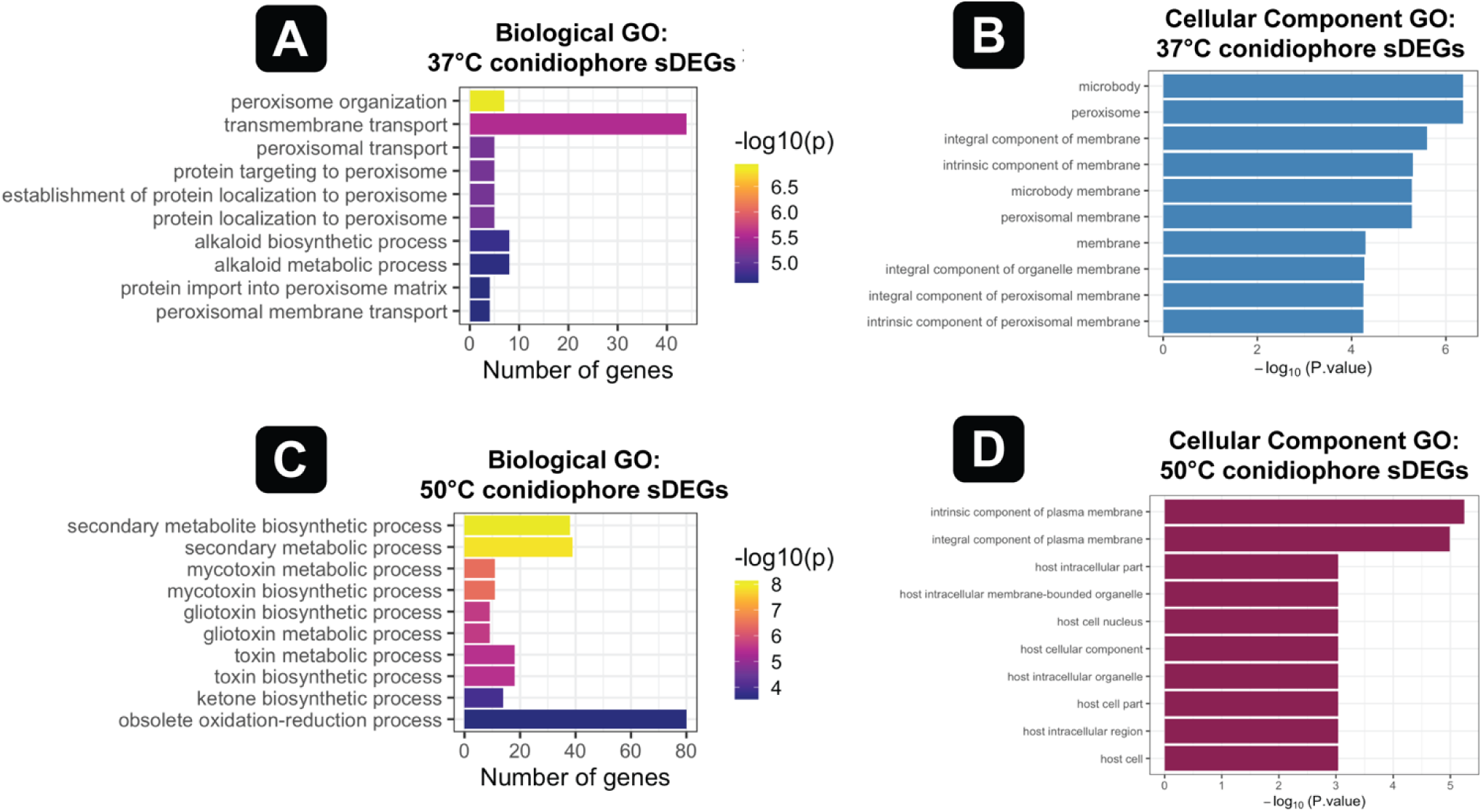
GO enrichment analyses of conidiophore sDEGs. Biological Gene Ontology (GO) enrichment (**A** and **C**) and Cellular Components GO enrichment (**B** and **D**) from the 37°C versus 50°C conidiophore differential expression analysis, with any shared upregulated hyphal DEGs removed. GO analyses were carried out using FungiDB (Fisher’s exact tests, p-value cutoff of 0.05). Bar graphs display the top 10 most significant GO terms ranked from highest to lowest p-values. (**A**) Biological GO analysis results for 37°C conidiophore sDEGs (n=334). (**B**) Cellular component GO enrichment for 37°C conidiophore sDEGs (n=334). (**C**) Biological GO analysis results for 50°C conidiophore sDEGs (n=929). (**D**) Cellular component GO enrichment for 50°C conidiophore sDEGs (n=929).

**Fig S4.**
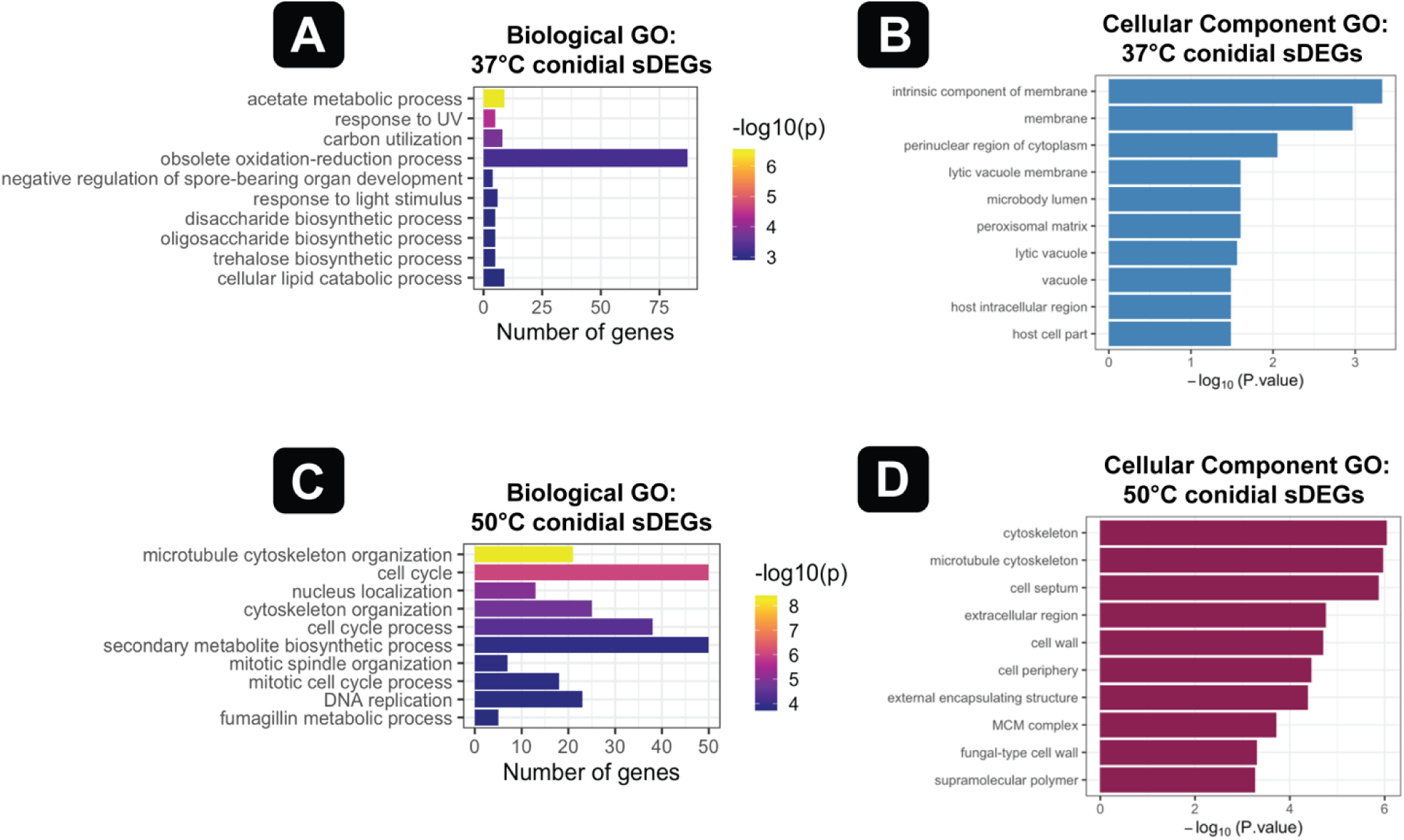
GO enrichment analyses of conidial sDEGs. Biological Gene Ontology (GO) enrichment (**A** and **C**) and Cellular Components GO enrichment (**B** and **D**) from the 37°C versus 50°C conidial differential expression analysis, with any shared upregulated hyphal DEGs removed. GO analyses were carried out using FungiDB (Fisher’s exact tests, p-value cutoff of 0.05). Bar graphs display the top 10 most significant GO terms ranked from highest to lowest p-values. (**A**) Biological GO analysis results for 37°C conidial sDEGs (n=972). (**B**) Cellular component GO enrichment for 37°C conidial sDEGs (n=972). (**C**) Biological GO analysis results for 50°C conidial sDEGs (n=1,985). (**D**) Cellular component GO enrichment for 50°C conidial sDEGs (n=1,985).

