## Supplemental Tables for "Temperature during *Aspergillus fumigatus* conidiophore development primes spore transcriptome for asexual, parasexual or sexual development"

Table S1

| 37°C hyphae vs 37°C conidiophores differential analysis | | 37°C hyphae vs 37°C conidia differential analysis | | 37°C conidia vs 37°C conidiophores differential analysis | |
|---|---|---|---|---|---|
| Total DEGs | 2,054 | Total DEGs | 3,554 | Total DEGs | 3,559 |
| Upregulated in hyphae | 1,063 | Upregulated in hyphae | 1,834 | Upregulated in conidia | 1,669 |
| Upregulated in conidiophores | 991 | Upregulated in conidia | 1,720 | Upregulated in conidiophores | 1,890 |
| **50°C hyphae vs 50°C conidiophores differential analysis** | | **50°C hyphae vs 50°C conidia differential analysis** | | **50°C conidia vs 50°C conidiophores differential analysis** | |
| Total DEGs | 2,649 | Total DEGs | 2,262 | Total DEGs | 1,523 |
| Upregulated in hyphae | 1,004 | Upregulated in hyphae | 781 | Upregulated in conidia | 818 |
| Upregulated in conidiophores | 1,645 | Upregulated in conidia | 1,481 | Upregulated in conidiophores | 705 |
| **37°C hyphae vs 50°C hyphae** | | **37°C conidiophores vs 50°C conidiophores** | | **37°C conidia vs 50°C conidia** | |
| Total DEGs | 811 | Total DEGs | 1,435 | Total DEGs | 3,179 |
| Upregulated in 37C | 477 | Upregulated in 37C | 416 | Upregulated in 37C | 1,076 |
| Upregulated in 50C | 334 | Upregulated in 50C | 1,019 | Upregulated in 50C | 2,103 |
| **37°C hyphal unique DEGs Upregulated only in hyphae 37°C** | | **37°C conidiophore unique DEGs Upregulated only in conidiophores 37°C** | | **37°C conidia unique DEGs Upregulated only in conidia 37°C** | |
| Upregulated in hyphae vs conidiophores | 1,063 | Upregulated in conidiophores vs hyphae | 991 | Upregulated in conidia vs hyphae | 1,720 |
| Upregulated in hyphae vs conidia | 1,834 | Upregulated in conidiophores vs conidia | 1,890 | Upregulated in conidia vs conidiophores | 1,669 |

Table S1

| | | | | | |
|---|---|---|---|---|---|
| venndiagram: upreg in hyphae vs conidiophore only | 557 | venndiagram: upreg in conidiophore vs hyphae only | 451 | venndiagram: upreg in conidia vs hyphae only | 650 |
| venndiagram: upreg in hyphae vs conidia only | 1328 | venndiagram: upreg in conidiophore vs conidia only | 1350 | venndiagram: upreg in conidia vs conidiophore only | 599 |
| venndiagram: Upregulated in both (shared) | 506 | venndiagram: Upregulated in both (shared) | 540 | venndiagram: Upregulated in both (shared) | 1,070 |
| venndiagram: Total upreg hyphal DEGs | 2,391 | venndiagram: Total conidiophore upreg DEGs | 2,341 | venndiagram: Total conidia upreg DEGs | 2,319 |
| **50°C hyphal unique DEGs Upregulated only in hyphae 50°C** | | **50°C conidiophore unique DEGs Upregulated only in conidiophores 50°C** | | **50°C conidial unique DEGs Upregulated only in conidia 50°C** | |
| Upregulated in hyphae vs conidiophores | 1,004 | Upregulated in conidiophores vs hyphae | 1,645 | Upregulated in conidia vs hyphae | 1,481 |
| Upregulated in hyphae vs conidia | 781 | Upregulated in conidiophores vs conidia | 705 | Upregulated in conidia vs conidiophores | 818 |
| venndiagram: upreg in hyphae vs conidiophore only | 536 | venndiagram: upreg in conidiophore vs hyphae only | 1,229 | venndiagram: upreg in conidia vs hyphae only | 1,060 |
| venndiagram: upreg in hyphae vs conidia only | 313 | venndiagram: upreg in conidiophore vs conidia only | 289 | venndiagram: upreg in conidia vs conidiophore only | 397 |
| venndiagram: Upregulated in both (shared) | 468 | venndiagram: Upregulated in both (shared) | 416 | venndiagram: Upregulated in both (shared) | 421 |
| venndiagram: Total hyphal DEGs | 1,317 | venndiagram: Total conidiophore upreg DEGs | 1,934 | venndiagram: Total conidia upreg DEGs | 1,878 |

### Table S2

| **Total genes expressed in each tissue type after filtering out low reads ( < 5)** | | | | | | **Cconidiophores 37 & 50** | | |
|---|---|---|---|---|---|---|---|---|
| 37 hyphae (8,323) | 37 conidiophores (8,957) | 37 conidia (8,617) | 50 hyphae (8,397) | 50 conidiophores (9,516) | 50 conidia (8,847) | 50 conidiophore only n=592 | 37 conidiophore only n=33 | shared 37&50 conidiophore n=8924 |
| AFUB_000010 | AFUB_000010 | AFUB_000020 | AFUB_000020 | AFUB_000010 | AFUB_000010 | AFUB_000060 | AFUB_000190 | AFUB_000010 |
| AFUB_000020 | AFUB_000020 | AFUB_000040 | AFUB_000040 | AFUB_000020 | AFUB_000020 | AFUB_000070 | AFUB_000860 | AFUB_000020 |
| AFUB_000040 | AFUB_000040 | AFUB_000080 | AFUB_000080 | AFUB_000040 | AFUB_000040 | AFUB_000090 | AFUB_002900 | AFUB_000040 |
| AFUB_000080 | AFUB_000050 | AFUB_000110 | AFUB_000100 | AFUB_000050 | AFUB_000050 | AFUB_000100 | AFUB_015220 | AFUB_000050 |
| AFUB_000100 | AFUB_000080 | AFUB_000120 | AFUB_000110 | AFUB_000060 | AFUB_000110 | AFUB_000120 | AFUB_016750 | AFUB_000080 |
| AFUB_000110 | AFUB_000110 | AFUB_000130 | AFUB_000240 | AFUB_000070 | AFUB_000140 | AFUB_000170 | AFUB_019890 | AFUB_000110 |
| AFUB_000240 | AFUB_000130 | AFUB_000140 | AFUB_000260 | AFUB_000080 | AFUB_000150 | AFUB_000180 | AFUB_025190 | AFUB_000130 |
| AFUB_000260 | AFUB_000140 | AFUB_000150 | AFUB_000280 | AFUB_000090 | AFUB_000190 | AFUB_000200 | AFUB_029250 | AFUB_000140 |
| AFUB_000300 | AFUB_000150 | AFUB_000190 | AFUB_000300 | AFUB_000100 | AFUB_000200 | AFUB_000210 | AFUB_033380 | AFUB_000150 |
| AFUB_000310 | AFUB_000160 | AFUB_000240 | AFUB_000310 | AFUB_000110 | AFUB_000210 | AFUB_000270 | AFUB_033510 | AFUB_000160 |
| AFUB_000340 | AFUB_000190 | AFUB_000260 | AFUB_000320 | AFUB_000120 | AFUB_000240 | AFUB_000370 | AFUB_033630 | AFUB_000240 |
| AFUB_000350 | AFUB_000240 | AFUB_000270 | AFUB_000330 | AFUB_000130 | AFUB_000260 | AFUB_000950 | AFUB_033930 | AFUB_000260 |
| AFUB_000360 | AFUB_000260 | AFUB_000280 | AFUB_000340 | AFUB_000140 | AFUB_000270 | AFUB_001020 | AFUB_034440 | AFUB_000280 |
| AFUB_000370 | AFUB_000280 | AFUB_000290 | AFUB_000350 | AFUB_000150 | AFUB_000280 | AFUB_001380 | AFUB_039510 | AFUB_000290 |
| AFUB_000380 | AFUB_000290 | AFUB_000300 | AFUB_000360 | AFUB_000160 | AFUB_000290 | AFUB_001390 | AFUB_041110 | AFUB_000300 |
| AFUB_000390 | AFUB_000300 | AFUB_000310 | AFUB_000370 | AFUB_000170 | AFUB_000300 | AFUB_002180 | AFUB_041660 | AFUB_000310 |
| AFUB_000400 | AFUB_000310 | AFUB_000320 | AFUB_000380 | AFUB_000180 | AFUB_000310 | AFUB_002220 | AFUB_044240 | AFUB_000320 |
| AFUB_000410 | AFUB_000320 | AFUB_000330 | AFUB_000390 | AFUB_000200 | AFUB_000320 | AFUB_002600 | AFUB_044790 | AFUB_000330 |
| AFUB_000420 | AFUB_000330 | AFUB_000340 | AFUB_000400 | AFUB_000210 | AFUB_000330 | AFUB_002630 | AFUB_044840 | AFUB_000340 |
| AFUB_000430 | AFUB_000340 | AFUB_000360 | AFUB_000410 | AFUB_000240 | AFUB_000340 | AFUB_003020 | AFUB_046640 | AFUB_000350 |
| AFUB_000440 | AFUB_000350 | AFUB_000380 | AFUB_000420 | AFUB_000260 | AFUB_000360 | AFUB_003180 | AFUB_046940 | AFUB_000360 |
| AFUB_000450 | AFUB_000360 | AFUB_000390 | AFUB_000430 | AFUB_000270 | AFUB_000380 | AFUB_003340 | AFUB_054510 | AFUB_000380 |
| AFUB_000460 | AFUB_000380 | AFUB_000400 | AFUB_000440 | AFUB_000280 | AFUB_000390 | AFUB_003590 | AFUB_059100 | AFUB_000390 |
| AFUB_000470 | AFUB_000390 | AFUB_000410 | AFUB_000450 | AFUB_000290 | AFUB_000400 | AFUB_003710 | AFUB_062310 | AFUB_000400 |
| AFUB_000480 | AFUB_000400 | AFUB_000420 | AFUB_000460 | AFUB_000300 | AFUB_000410 | AFUB_003730 | AFUB_065330 | AFUB_000410 |
| AFUB_000490 | AFUB_000410 | AFUB_000430 | AFUB_000470 | AFUB_000310 | AFUB_000420 | AFUB_003740 | AFUB_071569 | AFUB_000420 |
| AFUB_000500 | AFUB_000420 | AFUB_000440 | AFUB_000480 | AFUB_000320 | AFUB_000430 | AFUB_003890 | AFUB_081160 | AFUB_000430 |
| AFUB_000510 | AFUB_000430 | AFUB_000450 | AFUB_000490 | AFUB_000330 | AFUB_000440 | AFUB_004260 | AFUB_082510 | AFUB_000440 |
| AFUB_000520 | AFUB_000440 | AFUB_000460 | AFUB_000500 | AFUB_000340 | AFUB_000450 | AFUB_004550 | AFUB_084420 | AFUB_000450 |
| AFUB_000530 | AFUB_000450 | AFUB_000470 | AFUB_000510 | AFUB_000350 | AFUB_000460 | AFUB_005470 | AFUB_085020 | AFUB_000460 |
| AFUB_000540 | AFUB_000460 | AFUB_000480 | AFUB_000520 | AFUB_000360 | AFUB_000470 | AFUB_007040 | AFUB_087600 | AFUB_000470 |
| AFUB_000550 | AFUB_000470 | AFUB_000490 | AFUB_000530 | AFUB_000370 | AFUB_000480 | AFUB_007250 | AFUB_096860 | AFUB_000480 |
| AFUB_000560 | AFUB_000480 | AFUB_000500 | AFUB_000540 | AFUB_000380 | AFUB_000490 | AFUB_007400 | AFUB_097340 | AFUB_000490 |
| AFUB_000570 | AFUB_000490 | AFUB_000510 | AFUB_000550 | AFUB_000390 | AFUB_000500 | AFUB_007870 | | AFUB_000500 |
| AFUB_000580 | AFUB_000500 | AFUB_000520 | AFUB_000560 | AFUB_000400 | AFUB_000510 | AFUB_009590 | | AFUB_000510 |
| AFUB_000590 | AFUB_000510 | AFUB_000530 | AFUB_000570 | AFUB_000410 | AFUB_000520 | AFUB_009650 | | AFUB_000520 |
| AFUB_000600 | AFUB_000520 | AFUB_000540 | AFUB_000580 | AFUB_000420 | AFUB_000530 | AFUB_009890 | | AFUB_000530 |
| AFUB_000610 | AFUB_000530 | AFUB_000550 | AFUB_000590 | AFUB_000430 | AFUB_000540 | AFUB_010480 | | AFUB_000540 |
| AFUB_000620 | AFUB_000540 | AFUB_000560 | AFUB_000600 | AFUB_000440 | AFUB_000550 | AFUB_010670 | | AFUB_000550 |
| AFUB_000630 | AFUB_000550 | AFUB_000570 | AFUB_000610 | AFUB_000450 | AFUB_000560 | AFUB_010770 | | AFUB_000560 |
| AFUB_000640 | AFUB_000560 | AFUB_000580 | AFUB_000620 | AFUB_000460 | AFUB_000570 | AFUB_010780 | | AFUB_000570 |
| AFUB_000650 | AFUB_000570 | AFUB_000590 | AFUB_000630 | AFUB_000470 | AFUB_000580 | AFUB_011190 | | AFUB_000580 |
| AFUB_000660 | AFUB_000580 | AFUB_000600 | AFUB_000640 | AFUB_000480 | AFUB_000590 | AFUB_011880 | | AFUB_000590 |
| AFUB_000670 | AFUB_000590 | AFUB_000610 | AFUB_000650 | AFUB_000490 | AFUB_000600 | AFUB_013130 | | AFUB_000600 |
| AFUB_000690 | AFUB_000600 | AFUB_000620 | AFUB_000660 | AFUB_000500 | AFUB_000610 | AFUB_013300 | | AFUB_000610 |
| AFUB_000700 | AFUB_000610 | AFUB_000630 | AFUB_000670 | AFUB_000510 | AFUB_000620 | AFUB_013320 | | AFUB_000620 |
| AFUB_000740 | AFUB_000620 | AFUB_000640 | AFUB_000680 | AFUB_000520 | AFUB_000630 | AFUB_013780 | | AFUB_000630 |
| AFUB_000750 | AFUB_000630 | AFUB_000650 | AFUB_000690 | AFUB_000530 | AFUB_000640 | AFUB_014010 | | AFUB_000640 |
| AFUB_000760 | AFUB_000640 | AFUB_000660 | AFUB_000700 | AFUB_000540 | AFUB_000650 | AFUB_014180 | | AFUB_000650 |

### Table S3

| **Total genes expressed in each tissue type after filtering out low reads ( < 5)** | | | | | | **Cconidiophores 37 & 50** | | |
|---|---|---|---|---|---|---|---|---|
| 37 hyphae (8,323) | 37 conidiophores (8,957) | 37 conidia (8,617) | 50 hyphae (8,397) | 50 conidiophores (9,516) | 50 conidia (8,847) | 50 conidiophore only n=592 | 37 conidiophore only n=33 | shared 37&50 conidiophore n=8924 |
| AFUB_000010 | AFUB_000010 | AFUB_000020 | AFUB_000020 | AFUB_000010 | AFUB_000010 | AFUB_000060 | AFUB_000190 | AFUB_000010 |
| AFUB_000020 | AFUB_000020 | AFUB_000040 | AFUB_000040 | AFUB_000020 | AFUB_000020 | AFUB_000070 | AFUB_000860 | AFUB_000020 |
| AFUB_000040 | AFUB_000040 | AFUB_000080 | AFUB_000080 | AFUB_000040 | AFUB_000040 | AFUB_000090 | AFUB_002900 | AFUB_000040 |
| AFUB_000080 | AFUB_000050 | AFUB_000110 | AFUB_000100 | AFUB_000050 | AFUB_000050 | AFUB_000100 | AFUB_015220 | AFUB_000050 |
| AFUB_000100 | AFUB_000080 | AFUB_000120 | AFUB_000110 | AFUB_000060 | AFUB_000110 | AFUB_000120 | AFUB_016750 | AFUB_000080 |
| AFUB_000110 | AFUB_000110 | AFUB_000130 | AFUB_000240 | AFUB_000070 | AFUB_000140 | AFUB_000170 | AFUB_019890 | AFUB_000110 |
| AFUB_000240 | AFUB_000130 | AFUB_000140 | AFUB_000260 | AFUB_000080 | AFUB_000150 | AFUB_000180 | AFUB_025190 | AFUB_000130 |
| AFUB_000260 | AFUB_000140 | AFUB_000150 | AFUB_000280 | AFUB_000090 | AFUB_000190 | AFUB_000200 | AFUB_029250 | AFUB_000140 |
| AFUB_000300 | AFUB_000150 | AFUB_000190 | AFUB_000300 | AFUB_000100 | AFUB_000200 | AFUB_000210 | AFUB_033380 | AFUB_000150 |
| AFUB_000310 | AFUB_000160 | AFUB_000240 | AFUB_000310 | AFUB_000110 | AFUB_000210 | AFUB_000270 | AFUB_033510 | AFUB_000160 |
| AFUB_000340 | AFUB_000190 | AFUB_000260 | AFUB_000320 | AFUB_000120 | AFUB_000240 | AFUB_000370 | AFUB_033630 | AFUB_000240 |
| AFUB_000350 | AFUB_000240 | AFUB_000270 | AFUB_000330 | AFUB_000130 | AFUB_000260 | AFUB_000950 | AFUB_033930 | AFUB_000260 |
| AFUB_000360 | AFUB_000260 | AFUB_000280 | AFUB_000340 | AFUB_000140 | AFUB_000270 | AFUB_001020 | AFUB_034440 | AFUB_000280 |
| AFUB_000370 | AFUB_000280 | AFUB_000290 | AFUB_000350 | AFUB_000150 | AFUB_000280 | AFUB_001380 | AFUB_039510 | AFUB_000290 |
| AFUB_000380 | AFUB_000290 | AFUB_000300 | AFUB_000360 | AFUB_000160 | AFUB_000290 | AFUB_001390 | AFUB_041110 | AFUB_000300 |
| AFUB_000390 | AFUB_000300 | AFUB_000310 | AFUB_000370 | AFUB_000170 | AFUB_000300 | AFUB_002180 | AFUB_041660 | AFUB_000310 |
| AFUB_000400 | AFUB_000310 | AFUB_000320 | AFUB_000380 | AFUB_000180 | AFUB_000310 | AFUB_002220 | AFUB_044240 | AFUB_000320 |
| AFUB_000410 | AFUB_000320 | AFUB_000330 | AFUB_000390 | AFUB_000200 | AFUB_000320 | AFUB_002600 | AFUB_044790 | AFUB_000330 |
| AFUB_000420 | AFUB_000330 | AFUB_000340 | AFUB_000400 | AFUB_000210 | AFUB_000330 | AFUB_002630 | AFUB_044840 | AFUB_000340 |
| AFUB_000430 | AFUB_000340 | AFUB_000360 | AFUB_000410 | AFUB_000240 | AFUB_000340 | AFUB_003020 | AFUB_046640 | AFUB_000350 |
| AFUB_000440 | AFUB_000350 | AFUB_000380 | AFUB_000420 | AFUB_000260 | AFUB_000360 | AFUB_003180 | AFUB_046940 | AFUB_000360 |
| AFUB_000450 | AFUB_000360 | AFUB_000390 | AFUB_000430 | AFUB_000270 | AFUB_000380 | AFUB_003340 | AFUB_054510 | AFUB_000380 |
| AFUB_000460 | AFUB_000380 | AFUB_000400 | AFUB_000440 | AFUB_000280 | AFUB_000390 | AFUB_003590 | AFUB_059100 | AFUB_000390 |
| AFUB_000470 | AFUB_000390 | AFUB_000410 | AFUB_000450 | AFUB_000290 | AFUB_000400 | AFUB_003710 | AFUB_062310 | AFUB_000400 |
| AFUB_000480 | AFUB_000400 | AFUB_000420 | AFUB_000460 | AFUB_000300 | AFUB_000410 | AFUB_003730 | AFUB_065330 | AFUB_000410 |
| AFUB_000490 | AFUB_000410 | AFUB_000430 | AFUB_000470 | AFUB_000310 | AFUB_000420 | AFUB_003740 | AFUB_071569 | AFUB_000420 |
| AFUB_000500 | AFUB_000420 | AFUB_000440 | AFUB_000480 | AFUB_000320 | AFUB_000430 | AFUB_003890 | AFUB_081160 | AFUB_000430 |
| AFUB_000510 | AFUB_000430 | AFUB_000450 | AFUB_000490 | AFUB_000330 | AFUB_000440 | AFUB_004260 | AFUB_082510 | AFUB_000440 |
| AFUB_000520 | AFUB_000440 | AFUB_000460 | AFUB_000500 | AFUB_000340 | AFUB_000450 | AFUB_004550 | AFUB_084420 | AFUB_000450 |
| AFUB_000530 | AFUB_000450 | AFUB_000470 | AFUB_000510 | AFUB_000350 | AFUB_000460 | AFUB_005470 | AFUB_085020 | AFUB_000460 |
| AFUB_000540 | AFUB_000460 | AFUB_000480 | AFUB_000520 | AFUB_000360 | AFUB_000470 | AFUB_007040 | AFUB_087600 | AFUB_000470 |
| AFUB_000550 | AFUB_000470 | AFUB_000490 | AFUB_000530 | AFUB_000370 | AFUB_000480 | AFUB_007250 | AFUB_096860 | AFUB_000480 |
| AFUB_000560 | AFUB_000480 | AFUB_000500 | AFUB_000540 | AFUB_000380 | AFUB_000490 | AFUB_007400 | AFUB_097340 | AFUB_000490 |
| AFUB_000570 | AFUB_000490 | AFUB_000510 | AFUB_000550 | AFUB_000390 | AFUB_000500 | AFUB_007870 | | AFUB_000500 |
| AFUB_000580 | AFUB_000500 | AFUB_000520 | AFUB_000560 | AFUB_000400 | AFUB_000510 | AFUB_009590 | | AFUB_000510 |
| AFUB_000590 | AFUB_000510 | AFUB_000530 | AFUB_000570 | AFUB_000410 | AFUB_000520 | AFUB_009650 | | AFUB_000520 |
| AFUB_000600 | AFUB_000520 | AFUB_000540 | AFUB_000580 | AFUB_000420 | AFUB_000530 | AFUB_009890 | | AFUB_000530 |
| AFUB_000610 | AFUB_000530 | AFUB_000550 | AFUB_000590 | AFUB_000430 | AFUB_000540 | AFUB_010480 | | AFUB_000540 |
| AFUB_000620 | AFUB_000540 | AFUB_000560 | AFUB_000600 | AFUB_000440 | AFUB_000550 | AFUB_010670 | | AFUB_000550 |
| AFUB_000630 | AFUB_000550 | AFUB_000570 | AFUB_000610 | AFUB_000450 | AFUB_000560 | AFUB_010770 | | AFUB_000560 |
| AFUB_000640 | AFUB_000560 | AFUB_000580 | AFUB_000620 | AFUB_000460 | AFUB_000570 | AFUB_010780 | | AFUB_000570 |
| AFUB_000650 | AFUB_000570 | AFUB_000590 | AFUB_000630 | AFUB_000470 | AFUB_000580 | AFUB_011190 | | AFUB_000580 |
| AFUB_000660 | AFUB_000580 | AFUB_000600 | AFUB_000640 | AFUB_000480 | AFUB_000590 | AFUB_011880 | | AFUB_000590 |
| AFUB_000670 | AFUB_000590 | AFUB_000610 | AFUB_000650 | AFUB_000490 | AFUB_000600 | AFUB_013130 | | AFUB_000600 |
| AFUB_000690 | AFUB_000600 | AFUB_000620 | AFUB_000660 | AFUB_000500 | AFUB_000610 | AFUB_013300 | | AFUB_000610 |
| AFUB_000700 | AFUB_000610 | AFUB_000630 | AFUB_000670 | AFUB_000510 | AFUB_000620 | AFUB_013320 | | AFUB_000620 |
| AFUB_000740 | AFUB_000620 | AFUB_000640 | AFUB_000680 | AFUB_000520 | AFUB_000630 | AFUB_013780 | | AFUB_000630 |
| AFUB_000750 | AFUB_000630 | AFUB_000650 | AFUB_000690 | AFUB_000530 | AFUB_000640 | AFUB_014010 | | AFUB_000640 |
| AFUB_000760 | AFUB_000640 | AFUB_000660 | AFUB_000700 | AFUB_000540 | AFUB_000650 | AFUB_014180 | | AFUB_000650 |

### Table S4

| Tissue-unique DEGs (venndiagram filtering of differential analyses) | | | | Tissue-unique DEGs (venndiagram filtering of differential analyses) | | | | Venndiatgram presence/absence comparison: Tissue-unique DEGs 37°C vs 50°C hyphae | | | Core conidiophore DEGs: Venndiagram comparison of 37°C vs 50°C conidiophore DEGs | Core conidial DEGs: Venndiagram comparison of 37°C vs 50°C conidia DEGs |
|---|---|---|---|---|---|---|---|---|---|---|---|---|
| 37C hyphae upregulated and unique | | 37C conidiophore upregulated | 37C conidia upregulated | 50C hyphae upregulated and unique | | 50C conidiophore upregulated | 50C conidia upregulated | 37°C hyphae vs 50°C hyphae: hyphal-specific DEGs total = 739 | | | 37°C vs 50°C conidiophores: conidiophore-specific DEGs total = 822 | 37°C vs 50°C conidia: conidia-specific DEGs total = 1,302 |
| **Total upreg DEGs = 2,391** | **37 hyphae upreg shared = 506** | **conidiophore DEGs shared = 540** | **conidia DEGs shared = 1070** | **Total upreg DEGs = 1,317** | **50C hyphae upreg shared = 468** | **50 conidiophore DEGs shared = 416** | **50C conidia DEGs shared = 421** | **Shared hyphal DEGs = 197** | **37°C hyphae only = 271** | **50°C hyphae only = 271** | **Core conidiophore DEGs = 134** | **Shared conidia DEGs = 189** |
| AFUB_003120 | AFUB_097500 | AFUB_061980 | AFUB_064910 | AFUB_066060 | AFUB_044840 | AFUB_037040 | AFUB_013610 | AFUB_062350 | AFUB_097500 | AFUB_096730 | AFUB_079770 | AFUB_080540 |
| AFUB_044500 | AFUB_062350 | AFUB_079770 | AFUB_056570 | AFUB_017500 | AFUB_054460 | AFUB_011850 | AFUB_096020 | AFUB_034560 | AFUB_079180 | AFUB_057580 | AFUB_066820 | AFUB_007700 |
| AFUB_044470 | AFUB_079180 | AFUB_020550 | AFUB_020530 | AFUB_062500 | AFUB_062350 | AFUB_015200 | AFUB_012830 | AFUB_027270 | AFUB_033500 | AFUB_100990 | AFUB_066810 | AFUB_049320 |
| AFUB_052260 | AFUB_034560 | AFUB_021610 | AFUB_036040 | AFUB_047950 | AFUB_081960 | AFUB_061170 | AFUB_080380 | AFUB_070710 | AFUB_033520 | AFUB_078220 | AFUB_088270 | AFUB_060500 |
| AFUB_044490 | AFUB_027270 | AFUB_088550 | AFUB_021800 | AFUB_015790 | AFUB_053780 | AFUB_019410 | AFUB_082620 | AFUB_012310 | AFUB_033510 | AFUB_049740 | AFUB_062250 | AFUB_056720 |
| AFUB_006180 | AFUB_033500 | AFUB_014360 | AFUB_048620 | AFUB_062510 | AFUB_062480 | AFUB_048260 | AFUB_047170 | AFUB_044850 | AFUB_047340 | AFUB_066370 | AFUB_094300 | AFUB_051030 |
| AFUB_096890 | AFUB_070710 | AFUB_072920 | AFUB_071920 | AFUB_061480 | AFUB_044850 | AFUB_095590 | AFUB_063540 | AFUB_004020 | AFUB_003760 | AFUB_057850 | AFUB_091130 | AFUB_086760 |
| AFUB_085740 | AFUB_012310 | AFUB_038860 | AFUB_098390 | AFUB_074280 | AFUB_038070 | AFUB_012000 | AFUB_025840 | AFUB_084150 | AFUB_080970 | AFUB_099290 | AFUB_078130 | AFUB_062310 |
| AFUB_044920 | AFUB_033520 | AFUB_090010 | AFUB_019100 | AFUB_022840 | AFUB_096730 | AFUB_101880 | AFUB_054390 | AFUB_044860 | AFUB_075790 | AFUB_062460 | AFUB_072000 | AFUB_025840 |
| AFUB_081940 | AFUB_044850 | AFUB_019370 | AFUB_058360 | AFUB_084900 | AFUB_034560 | AFUB_021340 | AFUB_030150 | AFUB_081960 | AFUB_097510 | AFUB_048250 | AFUB_054040 | AFUB_054390 |
| AFUB_096950 | AFUB_004020 | AFUB_066820 | AFUB_090390 | AFUB_020570 | AFUB_061490 | AFUB_053550 | AFUB_086940 | AFUB_044840 | AFUB_052270 | AFUB_057590 | AFUB_001910 | AFUB_099570 |
| AFUB_049170 | AFUB_084150 | AFUB_010810 | AFUB_090220 | AFUB_081290 | AFUB_090640 | AFUB_025700 | AFUB_056690 | AFUB_054460 | AFUB_047190 | AFUB_062520 | AFUB_091610 | AFUB_048190 |
| AFUB_070810 | AFUB_033510 | AFUB_004770 | AFUB_091420 | AFUB_026700 | AFUB_057580 | AFUB_025950 | AFUB_082980 | AFUB_071410 | AFUB_094850 | AFUB_088770 | AFUB_013770 | AFUB_088000 |
| AFUB_057450 | AFUB_047340 | AFUB_093240 | AFUB_082090 | AFUB_097010 | AFUB_100990 | AFUB_077720 | AFUB_044450 | AFUB_071390 | AFUB_101630 | AFUB_011680 | AFUB_012030 | AFUB_082620 |
| AFUB_010350 | AFUB_044860 | AFUB_078820 | AFUB_031280 | AFUB_054390 | AFUB_078220 | AFUB_043340 | AFUB_080640 | AFUB_099650 | AFUB_057610 | AFUB_096750 | AFUB_044550 | AFUB_007060 |
| AFUB_097210 | AFUB_081960 | AFUB_038570 | AFUB_031110 | AFUB_081280 | AFUB_049740 | AFUB_090250 | AFUB_007700 | AFUB_044810 | AFUB_035570 | AFUB_033900 | AFUB_071610 | AFUB_092150 |
| AFUB_036080 | AFUB_044840 | AFUB_101120 | AFUB_077800 | AFUB_079620 | AFUB_081530 | AFUB_030830 | AFUB_080190 | AFUB_056870 | AFUB_064390 | AFUB_001440 | AFUB_049450 | AFUB_070890 |
| AFUB_002680 | AFUB_054460 | AFUB_082870 | AFUB_042060 | AFUB_098830 | AFUB_076150 | AFUB_028610 | AFUB_087070 | AFUB_084100 | AFUB_071450 | AFUB_044240 | AFUB_044330 | AFUB_086930 |
| AFUB_041210 | AFUB_071410 | AFUB_011690 | AFUB_076000 | AFUB_060970 | AFUB_071390 | AFUB_011860 | AFUB_083030 | AFUB_041000 | AFUB_044960 | AFUB_101340 | AFUB_015200 | AFUB_101650 |
| AFUB_056360 | AFUB_071390 | AFUB_077450 | AFUB_059390 | AFUB_047580 | AFUB_066370 | AFUB_050990 | AFUB_074660 | AFUB_001260 | AFUB_048310 | AFUB_057900 | AFUB_033720 | AFUB_038740 |
| AFUB_049470 | AFUB_099650 | AFUB_066810 | AFUB_010090 | AFUB_036410 | AFUB_057850 | AFUB_062360 | AFUB_099510 | AFUB_101160 | AFUB_099660 | AFUB_081270 | AFUB_062530 | AFUB_013020 |
| AFUB_080890 | AFUB_044810 | AFUB_062130 | AFUB_058040 | AFUB_000470 | AFUB_099290 | AFUB_047340 | AFUB_102050 | AFUB_084410 | AFUB_094840 | AFUB_088980 | AFUB_062550 | AFUB_100820 |
| AFUB_058540 | AFUB_003760 | AFUB_082880 | AFUB_068320 | AFUB_068040 | AFUB_094880 | AFUB_092510 | AFUB_099490 | AFUB_044820 | AFUB_022090 | AFUB_084110 | AFUB_047720 | AFUB_049630 |
| AFUB_091700 | AFUB_056870 | AFUB_088270 | AFUB_077440 | AFUB_006250 | AFUB_062460 | AFUB_080700 | AFUB_045170 | AFUB_012300 | AFUB_081229 | AFUB_084920 | AFUB_080030 | AFUB_079300 |
| AFUB_091030 | AFUB_080970 | AFUB_056830 | AFUB_037120 | AFUB_090490 | AFUB_071860 | AFUB_044890 | AFUB_015060 | AFUB_000930 | AFUB_071250 | AFUB_034970 | AFUB_000760 | AFUB_081730 |
| AFUB_048140 | AFUB_075790 | AFUB_039970 | AFUB_022550 | AFUB_048770 | AFUB_091160 | AFUB_072280 | AFUB_052640 | AFUB_071680 | AFUB_049190 | AFUB_038380 | AFUB_000910 | AFUB_016740 |
| AFUB_071180 | AFUB_097510 | AFUB_015970 | AFUB_098450 | AFUB_061870 | AFUB_048250 | AFUB_047720 | AFUB_047730 | AFUB_062330 | AFUB_086740 | AFUB_096090 | AFUB_003720 | AFUB_081660 |
| AFUB_033340 | AFUB_084100 | AFUB_021510 | AFUB_014740 | AFUB_068150 | AFUB_057590 | AFUB_100020 | AFUB_071420 | AFUB_044830 | AFUB_000350 | AFUB_101320 | AFUB_004690 | AFUB_062290 |
| AFUB_086810 | AFUB_052270 | AFUB_062250 | AFUB_061160 | AFUB_013300 | AFUB_062520 | AFUB_048500 | AFUB_080400 | AFUB_099590 | AFUB_033620 | AFUB_037080 | AFUB_009670 | AFUB_004460 |
| AFUB_001840 | AFUB_041000 | AFUB_031910 | AFUB_082350 | AFUB_019980 | AFUB_036990 | AFUB_041890 | AFUB_100970 | AFUB_044590 | AFUB_087030 | AFUB_071840 | AFUB_010310 | AFUB_100330 |
| AFUB_071170 | AFUB_001260 | AFUB_066900 | AFUB_083510 | AFUB_088360 | AFUB_088770 | AFUB_044350 | AFUB_034540 | AFUB_044430 | AFUB_090600 | AFUB_058670 | AFUB_011080 | AFUB_084890 |
| AFUB_091470 | AFUB_101160 | AFUB_019530 | AFUB_017690 | AFUB_039080 | AFUB_011680 | AFUB_028810 | AFUB_086760 | AFUB_033350 | AFUB_071220 | AFUB_013290 | AFUB_013170 | AFUB_079290 |
| AFUB_087440 | AFUB_047190 | AFUB_100960 | AFUB_014300 | AFUB_084140 | AFUB_096750 | AFUB_048050 | AFUB_038220 | AFUB_046830 | AFUB_061470 | AFUB_011190 | AFUB_013310 | AFUB_090630 |
| AFUB_088560 | AFUB_084410 | AFUB_001890 | AFUB_051840 | AFUB_042050 | AFUB_051370 | AFUB_017850 | AFUB_087920 | AFUB_062380 | AFUB_092790 | AFUB_017510 | AFUB_017580 | AFUB_081780 |
| AFUB_017820 | AFUB_094850 | AFUB_046490 | AFUB_030270 | AFUB_017870 | AFUB_087520 | AFUB_085550 | AFUB_059760 | AFUB_016830 | AFUB_092780 | AFUB_081050 | AFUB_019530 | AFUB_099510 |
| AFUB_012000 | AFUB_101630 | AFUB_097750 | AFUB_074480 | AFUB_039860 | AFUB_033900 | AFUB_005510 | AFUB_056650 | AFUB_022910 | AFUB_004060 | AFUB_096720 | AFUB_020300 | AFUB_085280 |
| AFUB_047510 | AFUB_044820 | AFUB_074350 | AFUB_003400 | AFUB_030510 | AFUB_033560 | AFUB_093470 | AFUB_081660 | AFUB_041760 | AFUB_004040 | AFUB_011180 | AFUB_022210 | AFUB_101360 |
| AFUB_044600 | AFUB_057610 | AFUB_048750 | AFUB_099040 | AFUB_081680 | AFUB_034690 | AFUB_034740 | AFUB_099190 | AFUB_096830 | AFUB_071200 | AFUB_051380 | AFUB_025470 | AFUB_095010 |
| AFUB_041240 | AFUB_012300 | AFUB_057030 | AFUB_091410 | AFUB_058680 | AFUB_062340 | AFUB_006390 | AFUB_089910 | AFUB_080710 | AFUB_044940 | AFUB_079610 | AFUB_027390 | AFUB_080640 |
| AFUB_096940 | AFUB_035570 | AFUB_094300 | AFUB_059700 | AFUB_044660 | AFUB_087540 | AFUB_041610 | AFUB_051490 | AFUB_063960 | AFUB_035149 | AFUB_097240 | AFUB_029650 | AFUB_090480 |
| AFUB_049180 | AFUB_000930 | AFUB_072940 | AFUB_039020 | AFUB_076320 | AFUB_007800 | AFUB_015900 | AFUB_061550 | AFUB_001360 | AFUB_084570 | AFUB_068290 | AFUB_031120 | AFUB_000190 |
| AFUB_046390 | AFUB_071680 | AFUB_020850 | AFUB_036910 | AFUB_085490 | AFUB_050170 | AFUB_047660 | AFUB_000870 | AFUB_075780 | AFUB_101880 | AFUB_014820 | AFUB_032810 | AFUB_000270 |
| AFUB_096450 | AFUB_062330 | AFUB_064500 | AFUB_073980 | AFUB_074050 | AFUB_001440 | AFUB_017010 | AFUB_096120 | AFUB_094820 | AFUB_003770 | AFUB_063530 | AFUB_033220 | AFUB_000380 |
| AFUB_041670 | AFUB_044830 | AFUB_025780 | AFUB_081210 | AFUB_004030 | AFUB_044240 | AFUB_068210 | AFUB_055880 | AFUB_055560 | AFUB_071000 | AFUB_095120 | AFUB_033230 | AFUB_000860 |
| AFUB_084140 | AFUB_064390 | AFUB_087020 | AFUB_066280 | AFUB_074640 | AFUB_101340 | AFUB_070910 | AFUB_082020 | AFUB_036990 | AFUB_063700 | AFUB_065340 | AFUB_033650 | AFUB_000870 |
| AFUB_024950 | AFUB_099590 | AFUB_065760 | AFUB_082660 | AFUB_049340 | AFUB_057900 | AFUB_055640 | AFUB_081330 | AFUB_017800 | AFUB_062190 | AFUB_022110 | AFUB_033660 | AFUB_001280 |
| AFUB_038280 | AFUB_044590 | AFUB_054290 | AFUB_073390 | AFUB_072390 | AFUB_081270 | AFUB_012550 | AFUB_082930 | AFUB_058760 | AFUB_053180 | AFUB_054890 | AFUB_033670 | AFUB_004510 |
| AFUB_022100 | AFUB_044430 | AFUB_072490 | AFUB_038950 | AFUB_045260 | AFUB_088980 | AFUB_025000 | AFUB_086230 | AFUB_084800 | AFUB_003610 | AFUB_046970 | AFUB_033680 | AFUB_006130 |
| AFUB_100050 | AFUB_071450 | AFUB_043030 | AFUB_025490 | AFUB_013040 | AFUB_084150 | AFUB_096140 | AFUB_083020 | AFUB_032690 | AFUB_063750 | AFUB_100980 | AFUB_033690 | AFUB_007020 |
| AFUB_020900 | AFUB_044960 | AFUB_096560 | AFUB_084670 | AFUB_032110 | AFUB_084110 | AFUB_080250 | AFUB_029550 | AFUB_091070 | AFUB_050510 | AFUB_010610 | AFUB_033710 | AFUB_009190 |
| AFUB_028890 | AFUB_048310 | AFUB_091130 | AFUB_054750 | AFUB_057690 | AFUB_044830 | AFUB_012030 | AFUB_078260 | AFUB_094860 | AFUB_087270 | AFUB_084830 | AFUB_033730 | AFUB_013050 |
| AFUB_078430 | AFUB_033350 | AFUB_099300 | AFUB_005360 | AFUB_004080 | AFUB_041760 | AFUB_029460 | AFUB_100120 | AFUB_046820 | AFUB_077840 | AFUB_087280 | AFUB_033750 | AFUB_013530 |
| AFUB_080950 | AFUB_046830 | AFUB_021580 | AFUB_089810 | AFUB_090060 | AFUB_084920 | AFUB_096940 | AFUB_068710 | AFUB_028160 | AFUB_017010 | AFUB_084160 | AFUB_035460 | AFUB_015060 |
| AFUB_063690 | AFUB_099660 | AFUB_053440 | AFUB_060300 | AFUB_071890 | AFUB_033350 | AFUB_093990 | AFUB_088000 | AFUB_052300 | AFUB_058750 | AFUB_018420 | AFUB_035470 | AFUB_015070 |
| AFUB_001040 | AFUB_062380 | AFUB_092330 | AFUB_065180 | AFUB_016800 | AFUB_034970 | AFUB_101590 | AFUB_090480 | AFUB_033560 | AFUB_030220 | AFUB_063340 | AFUB_036210 | AFUB_015140 |
| AFUB_056240 | AFUB_016830 | AFUB_022210 | AFUB_008310 | AFUB_094610 | AFUB_058270 | AFUB_022590 | AFUB_056180 | AFUB_016570 | AFUB_094830 | AFUB_026810 | AFUB_037680 | AFUB_016350 |
| AFUB_003270 | AFUB_022910 | AFUB_083400 | AFUB_052880 | AFUB_031520 | AFUB_038380 | AFUB_073680 | AFUB_071210 | AFUB_096840 | AFUB_051830 | AFUB_058890 | AFUB_042000 | AFUB_016730 |
| AFUB_044550 | AFUB_041760 | AFUB_088960 | AFUB_004940 | AFUB_055840 | AFUB_090660 | AFUB_093050 | AFUB_062910 | AFUB_034510 | AFUB_017380 | AFUB_060880 | AFUB_045150 | AFUB_016750 |
| AFUB_006820 | AFUB_096830 | AFUB_053130 | AFUB_045350 | AFUB_077770 | AFUB_096090 | AFUB_093290 | AFUB_045860 | AFUB_096850 | AFUB_071230 | AFUB_081700 | AFUB_045610 | AFUB_017060 |
| AFUB_027200 | AFUB_080710 | AFUB_004850 | AFUB_012840 | AFUB_036540 | AFUB_034510 | AFUB_041390 | AFUB_087400 | AFUB_016580 | AFUB_101130 | AFUB_049290 | AFUB_045630 | AFUB_017200 |
| AFUB_090130 | AFUB_094840 | AFUB_075970 | AFUB_018740 | AFUB_049970 | AFUB_025190 | AFUB_102330 | AFUB_065800 | AFUB_086680 | AFUB_094940 | AFUB_099520 | AFUB_049150 | AFUB_017280 |
| AFUB_086960 | AFUB_063960 | AFUB_031710 | AFUB_050020 | AFUB_057860 | AFUB_101320 | AFUB_041040 | AFUB_032120 | AFUB_007800 | AFUB_057620 | AFUB_011030 | AFUB_049200 | AFUB_017290 |
| AFUB_063150 | AFUB_022090 | AFUB_075550 | AFUB_005880 | AFUB_084510 | AFUB_052310 | AFUB_092160 | AFUB_013020 | AFUB_092800 | AFUB_078250 | AFUB_016430 | AFUB_049530 | AFUB_017570 |
| AFUB_092230 | AFUB_001360 | AFUB_100170 | AFUB_003210 | AFUB_095800 | AFUB_037080 | AFUB_025450 | AFUB_059990 | AFUB_090640 | AFUB_068280 | AFUB_087510 | AFUB_049890 | AFUB_018660 |
| AFUB_070860 | AFUB_075780 | AFUB_068500 | AFUB_060790 | AFUB_080900 | AFUB_070710 | AFUB_000960 | AFUB_006130 | AFUB_050660 | AFUB_040020 | AFUB_007410 | AFUB_057130 | AFUB_019100 |
| AFUB_037020 | AFUB_094820 | AFUB_018680 | AFUB_066740 | AFUB_084540 | AFUB_071840 | AFUB_044440 | AFUB_022250 | AFUB_000940 | AFUB_097290 | AFUB_086910 | AFUB_057140 | AFUB_020700 |
| AFUB_060870 | AFUB_081229 | AFUB_073050 | AFUB_034900 | AFUB_050440 | AFUB_050150 | AFUB_069990 | AFUB_086930 | AFUB_045790 | AFUB_081220 | AFUB_003360 | AFUB_057920 | AFUB_021050 |
| AFUB_013430 | AFUB_071250 | AFUB_078130 | AFUB_056120 | AFUB_063880 | AFUB_081690 | AFUB_020030 | AFUB_050350 | AFUB_050160 | AFUB_059520 | AFUB_066570 | AFUB_057930 | AFUB_021270 |
| AFUB_070900 | AFUB_055560 | AFUB_030100 | AFUB_080090 | AFUB_074610 | AFUB_058670 | AFUB_009450 | AFUB_070300 | AFUB_023720 | AFUB_032950 | AFUB_090880 | AFUB_060430 | AFUB_021280 |
| AFUB_054220 | AFUB_036990 | AFUB_054490 | AFUB_083770 | AFUB_075670 | AFUB_013290 | AFUB_020690 | AFUB_015140 | AFUB_062200 | AFUB_094800 | AFUB_006280 | AFUB_063490 | AFUB_021290 |
| AFUB_044610 | AFUB_017800 | AFUB_063440 | AFUB_027320 | AFUB_038140 | AFUB_062330 | AFUB_001820 | AFUB_076340 | AFUB_034660 | AFUB_096520 | AFUB_079380 | AFUB_066400 | AFUB_022250 |
| AFUB_092760 | AFUB_049190 | AFUB_024730 | AFUB_087890 | AFUB_039960 | AFUB_084410 | AFUB_094130 | AFUB_101530 | AFUB_044900 | AFUB_071240 | AFUB_039900 | AFUB_066410 | AFUB_023930 |
| AFUB_011820 | AFUB_058760 | AFUB_040180 | AFUB_068710 | AFUB_087170 | AFUB_009760 | AFUB_065110 | AFUB_045960 | AFUB_017420 | AFUB_091800 | AFUB_062450 | AFUB_066420 | AFUB_029950 |
| AFUB_096380 | AFUB_086740 | AFUB_072000 | AFUB_021140 | AFUB_087500 | AFUB_011190 | AFUB_066710 | AFUB_089560 | AFUB_062040 | AFUB_017940 | AFUB_032760 | AFUB_066440 | AFUB_030940 |
| AFUB_046090 | AFUB_000350 | AFUB_072110 | AFUB_070260 | AFUB_097090 | AFUB_018100 | AFUB_089510 | AFUB_046060 | AFUB_004650 | AFUB_091480 | AFUB_076760 | AFUB_068240 | AFUB_031390 |
| AFUB_071530 | AFUB_033620 | AFUB_082250 | AFUB_021160 | AFUB_042040 | AFUB_017510 | AFUB_064180 | AFUB_020610 | AFUB_055310 | AFUB_044970 | AFUB_100280 | AFUB_070910 | AFUB_031560 |
| AFUB_048020 | AFUB_087030 | AFUB_068630 | AFUB_019270 | AFUB_076670 | AFUB_091080 | AFUB_040960 | AFUB_041930 | AFUB_090270 | AFUB_046740 | AFUB_036060 | AFUB_071140 | AFUB_033120 |
| AFUB_024420 | AFUB_090600 | AFUB_063650 | AFUB_056690 | AFUB_031510 | AFUB_081050 | AFUB_010310 | AFUB_037410 | AFUB_076150 | AFUB_061900 | AFUB_068740 | AFUB_071540 | AFUB_033130 |
| AFUB_064090 | AFUB_084800 | AFUB_036980 | AFUB_065040 | AFUB_040220 | AFUB_034660 | AFUB_026530 | AFUB_062990 | AFUB_051370 | AFUB_055730 | AFUB_025400 | AFUB_071560 | AFUB_033170 |
| AFUB_087330 | AFUB_071220 | AFUB_005890 | AFUB_069710 | AFUB_093370 | AFUB_096720 | AFUB_038590 | AFUB_095860 | AFUB_090650 | AFUB_047120 | AFUB_029990 | AFUB_072010 | AFUB_033180 |
| AFUB_028260 | AFUB_061470 | AFUB_036970 | AFUB_075000 | AFUB_049260 | AFUB_098000 | AFUB_092500 | AFUB_090220 | AFUB_049010 | AFUB_099750 | AFUB_047560 | AFUB_074910 | AFUB_034280 |
| AFUB_080800 | AFUB_032690 | AFUB_072900 | AFUB_029000 | AFUB_038370 | AFUB_001360 | AFUB_073910 | AFUB_038740 | AFUB_044800 | AFUB_086210 | AFUB_046540 | AFUB_075550 | AFUB_034290 |
| AFUB_062170 | AFUB_092790 | AFUB_000360 | AFUB_038770 | AFUB_029960 | AFUB_011180 | AFUB_036100 | AFUB_020600 | AFUB_017830 | AFUB_085960 | AFUB_044500 | AFUB_075820 | AFUB_035440 |

### Table S5

**37 conidiophore (control) vs 50 conidiophore. DESeq2 prefilter low counts > 5. Filtered: padj < 0.05, Log2FC > 2**

negative log2FC value = upreg in 37 conidiophores positive log2FC value = upreg in 50 conidiophores

| phore37v50 | baseMean | log2FoldChang | lfcSE | stat | pvalue | padj | Gene name | Gene ID |
|---|---|---|---|---|---|---|---|---|
| **AFUB_071569** | 287.6391109 | -9.24195114 | 0.840622982 | -10.99416901 | 4.08E-28 | 2.58E-27 | | AFUB_071569 |
| **AFUB_092090** | 2384.679797 | -8.627397935 | 0.249011295 | -34.64661292 | 5.02E-263 | 1.42E-260 | | AFUB_092090 |
| **AFUB_086400** | 2478.633914 | -6.265614752 | 0.143927599 | -43.53310134 | 0 | 0 | | AFUB_086400 |
| **AFUB_017600** | 153.3219827 | -5.99054062 | 0.395382554 | -15.15125177 | 7.43E-52 | 9.59E-51 | | AFUB_017600 |
| **AFUB_057270** | 1647.315864 | -5.972933778 | 0.141221786 | -42.29470497 | 0 | 0 | | AFUB_057270 |
| **AFUB_086410** | 610.7608759 | -5.598858712 | 0.192040597 | -29.15455796 | 7.31E-187 | 1.05E-184 | | AFUB_086410 |
| **AFUB_044790** | 7.813984141 | -5.497757945 | 1.280685733 | -4.292823606 | 1.76E-05 | 3.68E-05 | | AFUB_044790 |
| **AFUB_048750** | 79.17193664 | -5.472731585 | 0.47888924 | -11.4279694 | 3.03E-30 | 2.07E-29 | | AFUB_048750 |
| **AFUB_086320** | 721.055062 | -5.450876613 | 0.169746302 | -32.11190208 | 3.01E-226 | 5.91E-224 | FtmE | AFUB_086320 |
| **AFUB_044840** | 7.356732073 | -5.404634404 | 1.283133 | -4.212060951 | 2.53E-05 | 5.21E-05 | | AFUB_044840 |
| **AFUB_086380** | 57.90513785 | -5.311872621 | 0.520953533 | -10.19644227 | 2.06E-24 | 1.13E-23 | | AFUB_086380 |
| **AFUB_087580** | 28294.13477 | -5.290768882 | 0.079332662 | -66.69092862 | 0 | 0 | | AFUB_087580 |
| **AFUB_089550** | 382.237989 | -5.227346717 | 0.210365909 | -24.84883008 | 2.66E-136 | 1.87E-134 | | AFUB_089550 |
| **AFUB_048760** | 54.09071938 | -5.207569872 | 0.523567531 | -9.946319357 | 2.62E-23 | 1.38E-22 | | AFUB_048760 |
| **AFUB_084610** | 45.95664417 | -4.80750175 | 0.511060162 | -9.406919395 | 5.11E-21 | 2.48E-20 | | AFUB_084610 |
| **AFUB_030300** | 49.83928932 | -4.789263366 | 0.483473371 | -9.905950676 | 3.92E-23 | 2.06E-22 | | AFUB_030300 |
| **AFUB_099930** | 217.6163896 | -4.699127601 | 0.233968097 | -20.08448011 | 1.01E-89 | 3.02E-88 | | AFUB_099930 |
| **AFUB_101340** | 202.2972296 | -4.592834713 | 0.236212054 | -19.44369323 | 3.30E-84 | 8.77E-83 | | AFUB_101340 |
| **AFUB_065420** | 54.68755897 | -4.575984376 | 0.441575114 | -10.36286745 | 3.66E-25 | 2.07E-24 | | AFUB_065420 |
| **AFUB_087570** | 668.1441986 | -4.5625882 | 0.161442197 | -28.26143533 | 1.03E-175 | 1.23E-173 | | AFUB_087570 |
| **AFUB_086360** | 475.9359706 | -4.512029185 | 0.169025831 | -26.69431749 | 5.48E-157 | 5.28E-155 | | AFUB_086360 |
| **AFUB_071390** | 59.33996857 | -4.394998915 | 0.399781886 | -10.99349188 | 4.11E-28 | 2.60E-27 | | AFUB_071390 |
| **AFUB_071550** | 102.652884 | -4.219341595 | 0.302902762 | -13.92969008 | 4.18E-44 | 4.42E-43 | | AFUB_071550 |
| **AFUB_003160** | 713.4600824 | -4.213836019 | 0.136617054 | -30.84414352 | 6.71E-209 | 1.15E-206 | | AFUB_003160 |
| **AFUB_086300** | 377.511121 | -4.140577763 | 0.163494175 | -25.32553697 | 1.67E-141 | 1.25E-139 | FtmP450 | AFUB_086300 |
| **AFUB_095240** | 795.0871169 | -4.138419824 | 0.129131699 | -32.04805524 | 2.34E-225 | 4.41E-223 | | AFUB_095240 |
| **AFUB_002200** | 209.8847757 | -4.102435113 | 0.214022073 | -19.16828041 | 6.81E-82 | 1.72E-80 | | AFUB_002200 |
| **AFUB_097090** | 118.3744055 | -4.070140705 | 0.276304652 | -14.73062679 | 4.10E-49 | 4.92E-48 | | AFUB_097090 |
| **AFUB_086370** | 1002.797889 | -4.024180226 | 0.136679469 | -29.44246323 | 1.57E-190 | 2.37E-188 | | AFUB_086370 |
| **AFUB_091940** | 457.2070825 | -4.021774997 | 0.152392756 | -26.39085406 | 1.74E-153 | 1.59E-151 | | AFUB_091940 |
| **AFUB_071560** | 360.9940925 | -4.013515586 | 0.162509875 | -24.69705668 | 1.15E-134 | 7.81E-133 | | AFUB_071560 |
| **AFUB_091950** | 568.8113359 | -3.98993645 | 0.137329426 | -29.05376185 | 1.38E-185 | 1.93E-183 | | AFUB_091950 |
| **AFUB_084140** | 1129.162341 | -3.987723873 | 0.141008299 | -28.280065 | 6.08E-176 | 7.32E-174 | | AFUB_084140 |
| **AFUB_095000** | 103.1458713 | -3.977515079 | 0.277377757 | -14.33970455 | 1.24E-46 | 1.40E-45 | | AFUB_095000 |
| **AFUB_046940** | 1.371263956 | -3.974903342 | 1.685801632 | -2.357871334 | 0.018380062 | 0.027424943 | | AFUB_046940 |
| **AFUB_096860** | 1.362873851 | -3.969504463 | 1.68732801 | -2.352538711 | 0.018645749 | 0.027786953 | | AFUB_096860 |
| **AFUB_068560** | 274.5638642 | -3.898250967 | 0.190185874 | -20.49705839 | 2.29E-93 | 7.45E-92 | | AFUB_068560 |
| **AFUB_087590** | 9437.585495 | -3.880384072 | 0.083266682 | -46.60188168 | 0 | 0 | | AFUB_087590 |
| **AFUB_068010** | 543.4778251 | -3.874823814 | 0.136281304 | -28.4325413 | 8.01E-178 | 1.00E-175 | | AFUB_068010 |
| **AFUB_069980** | 228.1547897 | -3.826437341 | 0.188099605 | -20.34261231 | 5.40E-92 | 1.70E-90 | | AFUB_069980 |
| **AFUB_100670** | 77.18946481 | -3.78292427 | 0.309698712 | -12.21485308 | 2.59E-34 | 2.02E-33 | | AFUB_100670 |
| **AFUB_011030** | 19.9552412 | -3.710360561 | 0.575875138 | -6.442994868 | 1.17E-10 | 3.49E-10 | | AFUB_011030 |
| **AFUB_089720** | 419.5354858 | -3.698683835 | 0.149670353 | -24.71220091 | 7.91E-135 | 5.40E-133 | | AFUB_089720 |
| **AFUB_081370** | 1021.430289 | -3.696233751 | 0.115010271 | -32.13829263 | 1.29E-226 | 2.58E-224 | | AFUB_081370 |
| **AFUB_096700** | 760.3038285 | -3.695422095 | 0.128274619 | -28.80867717 | 1.67E-182 | 2.20E-180 | | AFUB_096700 |
| **AFUB_020570** | 100.9377891 | -3.687970031 | 0.271578359 | -13.57976401 | 5.28E-42 | 5.22E-41 | | AFUB_020570 |
| **AFUB_089800** | 189.7792777 | -3.641951793 | 0.197332242 | -18.45593885 | 4.67E-76 | 1.01E-74 | | AFUB_089800 |
| **AFUB_086390** | 302.1194816 | -3.638297038 | 0.167411372 | -21.73267562 | 1.01E-104 | 4.13E-103 | | AFUB_086390 |
| **AFUB_086420** | 913.353195 | -3.608477364 | 0.109335521 | -33.00370573 | 7.19E-239 | 1.57E-236 | | AFUB_086420 |
| **AFUB_086330** | 1113.380333 | -3.590188297 | 0.102840805 | -34.91015355 | 5.21E-267 | 1.57E-264 | FtmPT1 | AFUB_086330 |
| **AFUB_086290** | 675.1394459 | -3.556878412 | 0.129847335 | -27.39277182 | 3.34E-165 | 3.43E-163 | FtmPT2 | AFUB_086290 |
| **AFUB_096090** | 196.5001869 | -3.53139085 | 0.191782739 | -18.41349676 | 1.02E-75 | 2.21E-74 | | AFUB_096090 |
| **AFUB_087600** | 3.904012495 | -3.51759894 | 1.250475082 | -2.813010023 | 0.004908012 | 0.007919224 | | AFUB_087600 |
| **AFUB_071890** | 37.03477479 | -3.503844215 | 0.409931538 | -8.547388748 | 1.26E-17 | 5.33E-17 | | AFUB_071890 |
| **AFUB_080710** | 454.0466879 | -3.500706279 | 0.162183399 | -21.58486199 | 2.49E-103 | 9.92E-102 | | AFUB_080710 |
| **AFUB_018320** | 275.2913076 | -3.4906298 | 0.165427126 | -21.10070994 | 7.84E-99 | 2.85E-97 | | AFUB_018320 |
| **AFUB_000870** | 109.8791813 | -3.46032147 | 0.279299394 | -12.38929098 | 2.99E-35 | 2.40E-34 | | AFUB_000870 |
| **AFUB_072490** | 250.9089488 | -3.447714638 | 0.173859249 | -19.83049306 | 1.62E-87 | 4.71E-86 | | AFUB_072490 |
| **AFUB_055040** | 108.4930721 | -3.432113409 | 0.24474355 | -14.02330486 | 1.12E-44 | 1.21E-43 | | AFUB_055040 |
| **AFUB_086340** | 1709.3446 | -3.421411662 | 0.09560294 | -35.78772435 | 1.71E-280 | 6.35E-278 | FtmD | AFUB_086340 |
| **AFUB_000750** | 909.4184694 | -3.365242767 | 0.128513356 | -26.185938 | 3.84E-151 | 3.40E-149 | | AFUB_000750 |
| **AFUB_047370** | 28.74273796 | -3.364500497 | 0.448776401 | -7.497053076 | 6.53E-14 | 2.33E-13 | | AFUB_047370 |
| **AFUB_037050** | 303.3626416 | -3.308903819 | 0.156125655 | -21.19385066 | 1.09E-99 | 4.00E-98 | | AFUB_037050 |
| **AFUB_006430** | 462.6865732 | -3.297101839 | 0.142475576 | -23.14152319 | 1.77E-118 | 9.42E-117 | | AFUB_006430 |
| **AFUB_067280** | 20.1590707 | -3.263830822 | 0.521477228 | -6.258817539 | 3.88E-10 | 1.11E-09 | | AFUB_067280 |
| **AFUB_047010** | 63.22592634 | -3.247126735 | 0.309008174 | -10.50822279 | 7.92E-26 | 4.60E-25 | | AFUB_047010 |
| **AFUB_081350** | 92.89089229 | -3.239354976 | 0.258877709 | -12.51307029 | 6.33E-36 | 5.20E-35 | ustYa | AFUB_081350 |
| **AFUB_062130** | 281.6667995 | -3.238258953 | 0.156886271 | -20.64080521 | 1.18E-94 | 3.95E-93 | | AFUB_062130 |
| **AFUB_003970** | 165.0387713 | -3.222876394 | 0.193148063 | -16.68604047 | 1.66E-62 | 2.73E-61 | | AFUB_003970 |
| **AFUB_055430** | 37.13950049 | -3.206300763 | 0.384046287 | -8.34873522 | 6.90E-17 | 2.81E-16 | | |
| **AFUB_026440** | 905.5154747 | -3.160727825 | 0.106289198 | -29.73705599 | 2.55E-194 | 4.03E-192 | mpdA | AFUB_026440 |
| **AFUB_050560** | 543.8452988 | -3.148611107 | 0.121287355 | -25.95992881 | 1.40E-148 | 1.16E-146 | | AFUB_050560 |
| **AFUB_057060** | 550.7267628 | -3.131209805 | 0.120795681 | -25.92153771 | 3.81E-148 | 3.11E-146 | | AFUB_057060 |
| **AFUB_033710** | 73.33171106 | -3.114792086 | 0.27140781 | -11.47642763 | 1.73E-30 | 1.20E-29 | easN | AFUB_033710 |
| **AFUB_082930** | 266.0612937 | -3.079949012 | 0.15582934 | -19.76488523 | 5.97E-87 | 1.71E-85 | | AFUB_082930 |
| **AFUB_096100** | 3855.26409 | -3.068351765 | 0.089030852 | -34.46391559 | 2.79E-260 | 7.26E-258 | | AFUB_096100 |
| **AFUB_046630** | 81.25896436 | -3.067471272 | 0.267641142 | -11.46113506 | 2.07E-30 | 1.43E-29 | | AFUB_046630 |
| **AFUB_028190** | 465.3691354 | -3.047833284 | 0.131859683 | -23.11421664 | 3.33E-118 | 1.75E-116 | | AFUB_028190 |
| **AFUB_063780** | 78.27450611 | -3.024188786 | 0.266758448 | -11.33680603 | 8.62E-30 | 5.80E-29 | | AFUB_063780 |
| **AFUB_000770** | 4176.809912 | -3.016364306 | 0.084804433 | -35.56847427 | 4.31E-277 | 1.48E-274 | | AFUB_000770 |
| **AFUB_086260** | 40.94463676 | -2.990528921 | 0.351557078 | -8.506524579 | 1.79E-17 | 7.50E-17 | | AFUB_086260 |
| **AFUB_035480** | 69.47800007 | -2.97733905 | 0.292379478 | -10.18313278 | 2.36E-24 | 1.29E-23 | | AFUB_035480 |
| **AFUB_032760** | 146.617347 | -2.967322878 | 0.199188935 | -14.89702666 | 3.45E-50 | 4.25E-49 | | AFUB_032760 |
| **AFUB_052640** | 290.4861508 | -2.957230687 | 0.150826057 | -19.60689523 | 1.35E-85 | 3.72E-84 | | AFUB_052640 |
| **AFUB_065340** | 1254.669419 | -2.95274792 | 0.095198202 | -31.01684558 | 3.20E-211 | 5.70E-209 | | AFUB_065340 |
| **AFUB_077130** | 2034.813248 | -2.950367727 | 0.083628896 | -35.27928573 | 1.22E-272 | 4.06E-270 | | AFUB_077130 |
| **AFUB_001440** | 738.8638181 | -2.949442011 | 0.132988245 | -22.17821586 | 5.57E-109 | 2.50E-107 | | AFUB_001440 |
| **AFUB_078260** | 152.1459766 | -2.948495237 | 0.19648086 | -15.00652653 | 6.65E-51 | 8.40E-50 | | AFUB_078260 |
| **AFUB_071690** | 122.6820262 | -2.928160701 | 0.211522379 | -13.84326666 | 1.40E-43 | 1.45E-42 | | AFUB_071690 |
| **AFUB_095120** | 17.43133677 | -2.903296364 | 0.529589043 | -5.48216849 | 4.20E-08 | 1.07E-07 | | AFUB_095120 |
| **AFUB_036330** | 798.1674607 | -2.894615234 | 0.128168478 | -22.58445515 | 6.16E-113 | 3.00E-111 | sec11 | AFUB_036330 |
| **AFUB_083440** | 2175.037587 | -2.891149761 | 0.114649662 | -25.21725509 | 2.59E-140 | 1.89E-138 | | AFUB_083440 |
| **AFUB_087020** | 19.76209065 | -2.886544786 | 0.516220027 | -5.591694696 | 2.25E-08 | 5.83E-08 | | AFUB_087020 |
| **AFUB_032630** | 185.0281841 | -2.880140927 | 0.184691784 | -15.59431 | 7.96E-55 | 1.10E-53 | | AFUB_032630 |
| **AFUB_055440** | 706.2014305 | -2.875258371 | 0.115026084 | -24.99657709 | 6.66E-138 | 4.75E-136 | | |
| **AFUB_010960** | 71.89247482 | -2.866036956 | 0.266667269 | -10.74761429 | 6.08E-27 | 3.68E-26 | | AFUB_010960 |
| **AFUB_033230** | 1618.488266 | -2.848776335 | 0.087260294 | -32.64688001 | 8.87E-234 | 1.82E-231 | Arb1 | AFUB_033230 |
| **AFUB_046640** | 3.810413463 | -2.838211525 | 1.125059169 | -2.522722006 | 0.011645042 | 0.017859561 | | AFUB_046640 |
| **AFUB_024360** | 159.9009381 | -2.831503429 | 0.188738921 | -15.00222323 | 7.10E-51 | 8.94E-50 | | AFUB_024360 |
| **AFUB_014730** | 321.4414921 | -2.812946931 | 0.15115619 | -18.60953847 | 2.69E-77 | 5.96E-76 | | AFUB_014730 |
| **AFUB_064440** | 996.0879106 | -2.812491374 | 0.098596854 | -28.52516346 | 5.71E-179 | 7.24E-177 | | AFUB_064440 |
| **AFUB_025690** | 1560.538727 | -2.812333064 | 0.093272528 | -30.15178338 | 1.02E-199 | 1.66E-197 | | AFUB_025690 |
| **AFUB_067250** | 914.2777335 | -2.805488594 | 0.102413218 | -27.39381346 | 3.25E-165 | 3.37E-163 | | AFUB_067250 |
| **AFUB_011020** | 98.55720217 | -2.799249277 | 0.229896978 | -12.17610295 | 4.17E-34 | 3.23E-33 | qutD | AFUB_011020 |
| **AFUB_055050** | 911.0272928 | -2.798992518 | 0.143096104 | -19.5602287 | 3.38E-85 | 9.19E-84 | | AFUB_055050 |
| **AFUB_099120** | 1112.807788 | -2.797275977 | 0.098575608 | -28.37695895 | 3.89E-177 | 4.81E-175 | | AFUB_099120 |
| **AFUB_011560** | 154.8793288 | -2.774982225 | 0.192115526 | -14.44434126 | 2.72E-47 | 3.15E-46 | | AFUB_011560 |
| **AFUB_085650** | 218.5295021 | -2.76818565 | 0.178382429 | -15.51826413 | 2.61E-54 | 3.59E-53 | | AFUB_085650 |
| **AFUB_085170** | 12.24492029 | -2.767709857 | 0.616171903 | -4.491781989 | 7.06E-06 | 1.53E-05 | | AFUB_085170 |
| **AFUB_035320** | 46.87322504 | -2.754287818 | 0.326393278 | -8.43855559 | 3.21E-17 | 1.33E-16 | | AFUB_035320 |
| **AFUB_053830** | 435.6725013 | -2.754250715 | 0.127818684 | -21.54810721 | 5.52E-103 | 2.14E-101 | | AFUB_053830 |
| **AFUB_077260** | 323.5190384 | -2.751385894 | 0.140764574 | -19.5460109 | 4.46E-85 | 1.21E-83 | | AFUB_077260 |
| **AFUB_024020** | 351.9561427 | -2.751018414 | 0.14491284 | -18.98395216 | 2.32E-80 | 5.55E-79 | | AFUB_024020 |
| **AFUB_074300** | 250.1748727 | -2.728565671 | 0.156607391 | -17.42296868 | 5.52E-68 | 1.03E-66 | | AFUB_074300 |
| **AFUB_021150** | 2950.977551 | -2.713471108 | 0.096575226 | -28.09696873 | 1.07E-173 | 1.24E-171 | | AFUB_021150 |
| **AFUB_083110** | 1086.74497 | -2.700815855 | 0.101933633 | -26.49582661 | 1.08E-154 | 1.00E-152 | | AFUB_083110 |
| **AFUB_012360** | 157.8856054 | -2.690975055 | 0.184900601 | -14.55363064 | 5.54E-48 | 6.46E-47 | | AFUB_012360 |
| **AFUB_000790** | 592.6226953 | -2.681260234 | 0.118743802 | -22.58021206 | 6.78E-113 | 3.28E-111 | | AFUB_000790 |
| **AFUB_038970** | 260.6047323 | -2.662424183 | 0.156646739 | -16.99635879 | 8.74E-65 | 1.50E-63 | | AFUB_038970 |
| **AFUB_036900** | 1702.920489 | -2.660112264 | 0.112372533 | -23.67226391 | 6.96E-124 | 4.09E-122 | | AFUB_036900 |
| **AFUB_023120** | 291.3180512 | -2.654218544 | 0.153853427 | -17.25160498 | 1.09E-66 | 1.97E-65 | | AFUB_023120 |
| **AFUB_080510** | 8.083057951 | -2.651905766 | 0.736057507 | -3.602851329 | 0.000314746 | 0.000581013 | | AFUB_080510 |
| **AFUB_006040** | 41.09779565 | -2.635698165 | 0.327867249 | -8.038918716 | 9.06E-16 | 3.50E-15 | | AFUB_006040 |
| **AFUB_100260** | 60.12419119 | -2.628379607 | 0.282317083 | -9.310026792 | 1.28E-20 | 6.09E-20 | | AFUB_100260 |
| **AFUB_063260** | 432.4942101 | -2.624919182 | 0.123393616 | -21.27273085 | 2.03E-100 | 7.62E-99 | | AFUB_063260 |
| **AFUB_091160** | 88.15702182 | -2.621582289 | 0.236221067 | -11.09800376 | 1.28E-28 | 8.29E-28 | | AFUB_091160 |
| **AFUB_052590** | 224.8136202 | -2.619365919 | 0.178485266 | -14.67553024 | 9.25E-49 | 1.11E-47 | | AFUB_052590 |
| **AFUB_002520** | 245.8802471 | -2.617856701 | 0.152544906 | -17.16122002 | 5.18E-66 | 9.26E-65 | | AFUB_002520 |
| **AFUB_016760** | 183.5083321 | -2.610690912 | 0.171008936 | -15.26640055 | 1.28E-52 | 1.68E-51 | | AFUB_016760 |
| **AFUB_047060** | 85.17069596 | -2.604729575 | 0.235224586 | -11.07337298 | 1.69E-28 | 1.09E-27 | | AFUB_047060 |
| **AFUB_053710** | 741.4810109 | -2.603520174 | 0.107653096 | -24.18435023 | 3.25E-129 | 2.06E-127 | | AFUB_053710 |
| **AFUB_025970** | 226.2997582 | -2.601525101 | 0.154784238 | -16.80742905 | 2.15E-63 | 3.60E-62 | | AFUB_025970 |
| **AFUB_041640** | 72.73048916 | -2.597488103 | 0.259664837 | -10.00323389 | 1.47E-23 | 7.86E-23 | | AFUB_041640 |
| **AFUB_030090** | 25.71806347 | -2.59740249 | 0.410157423 | -6.332696527 | 2.41E-10 | 7.03E-10 | | AFUB_030090 |
| **AFUB_047280** | 370.0565414 | -2.595598708 | 0.132295653 | -19.61968256 | 1.05E-85 | 2.92E-84 | | AFUB_047280 |
| **AFUB_093120** | 805.8192996 | -2.594431283 | 0.110761353 | -23.42361488 | 2.46E-121 | 1.38E-119 | | AFUB_093120 |
| **AFUB_081360** | 484.6032283 | -2.576646709 | 0.12392033 | -20.79276834 | 5.03E-96 | 1.73E-94 | | AFUB_081360 |
| **AFUB_010080** | 90.27075328 | -2.574751102 | 0.224978382 | -11.44443783 | 2.51E-30 | 1.72E-29 | | AFUB_010080 |
| **AFUB_027100** | 1739.117178 | -2.569592163 | 0.097859648 | -26.25793385 | 5.80E-152 | 5.22E-150 | | AFUB_027100 |
| **AFUB_020040** | 436.9856788 | -2.569376418 | 0.119826383 | -21.44249339 | 5.37E-102 | 2.07E-100 | | AFUB_020040 |
| **AFUB_084600** | 40.51902069 | -2.562372008 | 0.32989627 | -7.767205145 | 8.02E-15 | 2.98E-14 | | AFUB_084600 |
| **AFUB_045270** | 76.7139725 | -2.560659307 | 0.252781103 | -10.12994754 | 4.07E-24 | 2.21E-23 | | AFUB_045270 |

### Table S5

| | | | | | | | | |
|---|---|---|---|---|---|---|---|---|
| **AFUB_086310** | 1106.91944 | -2.559313309 | 0.104145253 | -24.57445965 | 2.37E-133 | 1.57E-131 | | AFUB_086310 |
| **AFUB_036170** | 18.52216144 | -2.553454255 | 0.480368736 | -5.315612913 | 1.06E-07 | 2.62E-07 | | AFUB_036170 |
| **AFUB_050980** | 922.7399698 | -2.551127475 | 0.104237032 | -24.47429131 | 2.78E-132 | 1.82E-130 | | AFUB_050980 |
| **AFUB_047020** | 420.5750815 | -2.549837412 | 0.130937521 | -19.47369549 | 1.84E-84 | 4.94E-83 | | AFUB_047020 |
| **AFUB_006820** | 113.8728782 | -2.548040704 | 0.210917758 | -12.08073101 | 1.34E-33 | 1.02E-32 | | AFUB_006820 |
| **AFUB_015200** | 848.4231126 | -2.547050941 | 0.100874646 | -25.24966436 | 1.14E-140 | 8.47E-139 | Arb1 ortholog | AFUB_015200 |
| **AFUB_099290** | 13.9303856 | -2.529178425 | 0.554155774 | -4.564020701 | 5.02E-06 | 1.10E-05 | | AFUB_099290 |
| **AFUB_029990** | 335.5859153 | -2.527205108 | 0.142285544 | -17.76150289 | 1.40E-70 | 2.78E-69 | | AFUB_029990 |
| **AFUB_035230** | 261.4736934 | -2.522072452 | 0.147327874 | -17.11877318 | 1.08E-65 | 1.90E-64 | | AFUB_035230 |
| **AFUB_014320** | 708.1262145 | -2.52147902 | 0.119823333 | -21.0433057 | 2.63E-98 | 9.47E-97 | | AFUB_014320 |
| **AFUB_060350** | 332.1615653 | -2.517340464 | 0.134267359 | -18.74871508 | 1.98E-78 | 4.51E-77 | | AFUB_060350 |
| **AFUB_056460** | 703.0232685 | -2.516045251 | 0.108292434 | -23.23380452 | 2.07E-119 | 1.13E-117 | | AFUB_056460 |
| **AFUB_068800** | 308.2856079 | -2.515003518 | 0.160410417 | -15.67855487 | 2.12E-55 | 2.99E-54 | alp1 | AFUB_068800 |
| **AFUB_052140** | 378.9778119 | -2.511520649 | 0.132828332 | -18.90801924 | 9.80E-80 | 2.31E-78 | | AFUB_052140 |
| **AFUB_011810** | 174.6553926 | -2.508433301 | 0.17189184 | -14.59309122 | 3.11E-48 | 3.65E-47 | | AFUB_011810 |
| **AFUB_087540** | 122.9924927 | -2.487666423 | 0.207028808 | -12.01603992 | 2.93E-33 | 2.21E-32 | | AFUB_087540 |
| **AFUB_081090** | 10.45273833 | -2.479876423 | 0.627376014 | -3.952775318 | 7.72E-05 | 0.000152007 | | AFUB_081090 |
| **AFUB_074640** | 64.2538889 | -2.470516231 | 0.260760204 | -9.474284016 | 2.69E-21 | 1.31E-20 | | AFUB_074640 |
| **AFUB_058040** | 43.73863733 | -2.46743242 | 0.327461714 | -7.535025669 | 4.88E-14 | 1.75E-13 | | |
| **AFUB_000640** | 278.7130719 | -2.459848051 | 0.159563021 | -15.41615365 | 1.27E-53 | 1.72E-52 | | AFUB_000640 |
| **AFUB_088770** | 110.2073284 | -2.45903554 | 0.204983076 | -11.99628566 | 3.72E-33 | 2.79E-32 | | AFUB_088770 |
| **AFUB_077210** | 481.3052123 | -2.457594812 | 0.120883838 | -20.3302182 | 6.95E-92 | 2.18E-90 | | AFUB_077210 |
| **AFUB_040930** | 244.7903327 | -2.453408407 | 0.160927406 | -15.24543559 | 1.77E-52 | 2.31E-51 | | AFUB_040930 |
| **AFUB_076140** | 183.4931209 | -2.449474766 | 0.16906251 | -14.48857447 | 1.43E-47 | 1.66E-46 | | AFUB_076140 |
| **AFUB_050770** | 115.4883516 | -2.4472355 | 0.201133275 | -12.16723336 | 4.65E-34 | 3.60E-33 | | AFUB_050770 |
| **AFUB_018940** | 354.3724764 | -2.440454223 | 0.130918669 | -18.64099475 | 1.49E-77 | 3.32E-76 | | AFUB_018940 |
| **AFUB_076190** | 1023.430603 | -2.434127067 | 0.100198647 | -24.29301342 | 2.32E-130 | 1.49E-128 | | AFUB_076190 |
| **AFUB_094750** | 642.9446092 | -2.4339343 | 0.110764395 | -21.97397728 | 5.11E-107 | 2.22E-105 | | AFUB_094750 |
| **AFUB_031210** | 674.6578522 | -2.432829904 | 0.119458479 | -20.36548532 | 3.38E-92 | 1.07E-90 | | AFUB_031210 |
| **AFUB_033650** | 292.1534957 | -2.430423995 | 0.138627205 | -17.53208533 | 8.15E-69 | 1.55E-67 | easA | AFUB_033650 |
| **AFUB_082180** | 633.877024 | -2.428281171 | 0.120283132 | -20.18804414 | 1.25E-90 | 3.85E-89 | | AFUB_082180 |
| **AFUB_096720** | 124.1690965 | -2.42632429 | 0.198829304 | -12.20305177 | 2.99E-34 | 2.33E-33 | | AFUB_096720 |
| **AFUB_038690** | 593.000895 | -2.422665961 | 0.116066407 | -20.87310216 | 9.40E-97 | 3.27E-95 | | AFUB_038690 |
| **AFUB_056510** | 731.7409196 | -2.413573824 | 0.104621694 | -23.06953499 | 9.37E-118 | 4.85E-116 | | AFUB_056510 |
| **AFUB_069940** | 446.0859528 | -2.406916166 | 0.126832465 | -18.97712989 | 2.64E-80 | 6.30E-79 | tvp23 | AFUB_069940 |
| **AFUB_001500** | 685.7462631 | -2.406179965 | 0.103646089 | -23.2153475 | 3.19E-119 | 1.73E-117 | | AFUB_001500 |
| **AFUB_078490** | 320.0678524 | -2.39324082 | 0.138002162 | -17.34205309 | 2.27E-67 | 4.17E-66 | | AFUB_078490 |
| **AFUB_014940** | 242.2252638 | -2.391987848 | 0.15429738 | -15.50245281 | 3.34E-54 | 4.57E-53 | | AFUB_014940 |
| **AFUB_059370** | 1099.595129 | -2.387552129 | 0.106049944 | -22.5134691 | 3.06E-112 | 1.47E-110 | | AFUB_059370 |
| **AFUB_014680** | 572.004363 | -2.387324857 | 0.114902678 | -20.77692968 | 7.00E-96 | 2.38E-94 | | AFUB_014680 |
| **AFUB_027960** | 196.830935 | -2.374379589 | 0.158628691 | -14.96815972 | 1.19E-50 | 1.49E-49 | | AFUB_027960 |
| **AFUB_028590** | 167.5698975 | -2.373323374 | 0.175033933 | -13.55921872 | 6.99E-42 | 6.86E-41 | | AFUB_028590 |
| **AFUB_000740** | 10.75973072 | -2.371306285 | 0.60681467 | -3.907793273 | 9.31E-05 | 0.000181796 | | AFUB_000740 |
| **AFUB_071530** | 23.15104334 | -2.37090284 | 0.435748218 | -5.440992629 | 5.30E-08 | 1.34E-07 | | AFUB_071530 |
| **AFUB_056470** | 342.8660671 | -2.364494915 | 0.137495526 | -17.19688625 | 2.80E-66 | 5.04E-65 | | AFUB_056470 |
| **AFUB_010230** | 715.7089819 | -2.36178341 | 0.106620281 | -22.15135227 | 1.01E-108 | 4.47E-107 | | AFUB_010230 |
| **AFUB_099560** | 2012.177431 | -2.361119732 | 0.092380635 | -25.5586004 | 4.41E-144 | 3.40E-142 | | AFUB_099560 |
| **AFUB_091700** | 62.78808323 | -2.351349802 | 0.26361997 | -8.91946769 | 4.69E-19 | 2.09E-18 | | AFUB_091700 |
| **AFUB_024290** | 745.0251546 | -2.350360018 | 0.101483457 | -23.16003109 | 1.15E-118 | 6.17E-117 | | AFUB_024290 |
| **AFUB_066390** | 5.88268424 | -2.350152008 | 0.829838545 | -2.832059345 | 0.004624926 | 0.007486274 | | AFUB_066390 |
| **AFUB_016510** | 1170.38432 | -2.346982154 | 0.104892506 | -22.37511767 | 6.88E-111 | 3.25E-109 | | AFUB_016510 |
| **AFUB_039080** | 48.1459022 | -2.346254129 | 0.292392348 | -8.024334933 | 1.02E-15 | 3.94E-15 | | AFUB_039080 |
| **AFUB_023230** | 9816.089834 | -2.343724429 | 0.071215441 | -32.91034054 | 1.56E-237 | 3.35E-235 | | AFUB_023230 |
| **AFUB_088160** | 91.67743256 | -2.341690778 | 0.224081091 | -10.45019357 | 1.46E-25 | 8.42E-25 | | AFUB_088160 |
| **AFUB_031900** | 5.656486285 | -2.33603287 | 0.871605668 | -2.680148783 | 0.007358944 | 0.011624719 | | AFUB_031900 |
| **AFUB_059570** | 211.5914142 | -2.333446103 | 0.15491585 | -15.06266852 | 2.85E-51 | 3.62E-50 | | AFUB_059570 |
| **AFUB_062000** | 72.1017512 | -2.332922674 | 0.250233401 | -9.322986738 | 1.13E-20 | 5.40E-20 | | AFUB_062000 |
| **AFUB_076910** | 284.5577209 | -2.332480366 | 0.138869459 | -16.79620838 | 2.60E-63 | 4.34E-62 | | AFUB_076910 |
| **AFUB_022610** | 210.8596224 | -2.327695634 | 0.168409947 | -13.8216042 | 1.89E-43 | 1.95E-42 | | AFUB_022610 |
| **AFUB_031780** | 194.2046679 | -2.322146981 | 0.168783643 | -13.75812812 | 4.55E-43 | 4.65E-42 | | AFUB_031780 |
| **AFUB_094740** | 4.762008159 | -2.31897718 | 0.902745937 | -2.568803785 | 0.010205022 | 0.015781671 | | AFUB_094740 |
| **AFUB_087250** | 37.86412446 | -2.311204111 | 0.330446342 | -6.994188811 | 2.67E-12 | 8.74E-12 | | AFUB_087250 |
| **AFUB_008510** | 180.1469417 | -2.311022306 | 0.164808449 | -14.02247471 | 1.14E-44 | 1.22E-43 | | AFUB_008510 |
| **AFUB_032730** | 264.0441776 | -2.309390415 | 0.14435741 | -15.99772687 | 1.33E-57 | 1.95E-56 | | AFUB_032730 |
| **AFUB_071840** | 10.359861 | -2.292562024 | 0.624088679 | -3.673455556 | 0.000239292 | 0.000447385 | | AFUB_071840 |
| **AFUB_076670** | 35.49037469 | -2.290255501 | 0.345146551 | -6.635603044 | 3.23E-11 | 9.97E-11 | | AFUB_076670 |
| **AFUB_082730** | 96.03721395 | -2.288766905 | 0.214691097 | -10.66074438 | 1.55E-26 | 9.22E-26 | | AFUB_082730 |
| **AFUB_004220** | 349.5278512 | -2.282151496 | 0.143214787 | -15.93516668 | 3.61E-57 | 5.29E-56 | | AFUB_004220 |
| **AFUB_022820** | 1383.489166 | -2.278356973 | 0.092095011 | -24.73920083 | 4.05E-135 | 2.79E-133 | | AFUB_022820 |
| **AFUB_064550** | 13.80123991 | -2.277481144 | 0.53317248 | -4.271565445 | 1.94E-05 | 4.04E-05 | | AFUB_064550 |
| **AFUB_066410** | 123.81408 | -2.275623311 | 0.196125753 | -11.60287867 | 3.98E-31 | 2.81E-30 | | AFUB_066410 |
| **AFUB_002700** | 201.8719049 | -2.274244574 | 0.156170786 | -14.56254805 | 4.86E-48 | 5.68E-47 | | AFUB_002700 |
| **AFUB_008810** | 515.2354017 | -2.272301 | 0.136669186 | -16.62628621 | 4.50E-62 | 7.32E-61 | | AFUB_008810 |
| **AFUB_037530** | 471.4457576 | -2.271062699 | 0.121636786 | -18.67085421 | 8.55E-78 | 1.92E-76 | | AFUB_037530 |
| **AFUB_081630** | 6.540165347 | -2.269966753 | 0.777908393 | -2.918038645 | 0.003522407 | 0.005786211 | plyE | AFUB_081630 |
| **AFUB_009900** | 181.1860237 | -2.262511154 | 0.176914413 | -12.78873281 | 1.90E-37 | 1.64E-36 | | AFUB_009900 |
| **AFUB_033180** | 38.49761909 | -2.261489923 | 0.320095811 | -7.065040676 | 1.61E-12 | 5.33E-12 | | AFUB_033180 |
| **AFUB_037770** | 8031.105549 | -2.261249677 | 0.0884451 | -25.56670392 | 3.58E-144 | 2.78E-142 | | AFUB_037770 |
| **AFUB_000820** | 1120.167005 | -2.260860644 | 0.09375853 | -24.11365282 | 1.80E-128 | 1.12E-126 | | AFUB_000820 |
| **AFUB_095770** | 198.706182 | -2.260314912 | 0.157262595 | -14.37287049 | 7.66E-47 | 8.78E-46 | | AFUB_095770 |
| **AFUB_045210** | 75.5623632 | -2.251256914 | 0.238818071 | -9.426660677 | 4.23E-21 | 2.06E-20 | | AFUB_045210 |
| **AFUB_064130** | 486.4084637 | -2.250033436 | 0.118256988 | -19.02664257 | 1.03E-80 | 2.49E-79 | | AFUB_064130 |
| **AFUB_046130** | 271.7865755 | -2.241097337 | 0.173071859 | -12.94894128 | 2.38E-38 | 2.10E-37 | | AFUB_046130 |
| **AFUB_041120** | 101.3818162 | -2.240959691 | 0.210075033 | -10.66742517 | 1.45E-26 | 8.60E-26 | | AFUB_041120 |
| **AFUB_043820** | 459.8236927 | -2.239733643 | 0.120971911 | -18.51449333 | 1.58E-76 | 3.45E-75 | | AFUB_043820 |
| **AFUB_012630** | 541.3602991 | -2.23922196 | 0.118793071 | -18.84976912 | 2.95E-79 | 6.85E-78 | | AFUB_012630 |
| **AFUB_005120** | 484.4975618 | -2.238649926 | 0.116198001 | -19.26582133 | 1.04E-82 | 2.65E-81 | | AFUB_005120 |
| **AFUB_076150** | 55.24050053 | -2.238472065 | 0.271722359 | -8.238085655 | 1.75E-16 | 7.01E-16 | | AFUB_076150 |
| **AFUB_068150** | 14.47279906 | -2.237479515 | 0.517345803 | -4.324920593 | 1.53E-05 | 3.20E-05 | | AFUB_068150 |
| **AFUB_083810** | 778.4425532 | -2.236091869 | 0.098258159 | -22.75731503 | 1.21E-114 | 6.13E-113 | vps10 | AFUB_083810 |
| **AFUB_015960** | 5285.988469 | -2.233833687 | 0.086048837 | -25.9600682 | 1.40E-148 | 1.16E-146 | brlA | AFUB_015960 |
| **AFUB_024550** | 320.3807999 | -2.230605909 | 0.136687071 | -16.31907026 | 7.22E-60 | 1.11E-58 | | AFUB_024550 |
| **AFUB_040860** | 1948.259943 | -2.227115668 | 0.085025788 | -26.19341411 | 3.16E-151 | 2.82E-149 | | AFUB_040860 |
| **AFUB_020170** | 1236.873304 | -2.220761282 | 0.095039841 | -23.36663501 | 9.34E-121 | 5.20E-119 | | AFUB_020170 |
| **AFUB_100390** | 26.86223275 | -2.220523547 | 0.380192642 | -5.840522153 | 5.20E-09 | 1.40E-08 | | AFUB_100390 |
| **AFUB_041370** | 667.3321428 | -2.212716749 | 0.110440264 | -20.03541704 | 2.71E-89 | 8.05E-88 | | AFUB_041370 |
| **AFUB_026700** | 159.1681346 | -2.210978307 | 0.171585549 | -12.88557408 | 5.43E-38 | 4.75E-37 | | AFUB_026700 |
| **AFUB_097210** | 47.94295286 | -2.210451139 | 0.288281998 | -7.6676697 | 1.75E-14 | 6.42E-14 | | AFUB_097210 |
| **AFUB_092070** | 209.9230061 | -2.210272218 | 0.165226805 | -13.37720121 | 8.22E-41 | 7.79E-40 | | AFUB_092070 |
| **AFUB_000890** | 174.3123467 | -2.208135391 | 0.163871461 | -13.47480136 | 2.20E-41 | 2.14E-40 | | AFUB_000890 |
| **AFUB_067230** | 321.7821594 | -2.207991644 | 0.138626421 | -15.92763942 | 4.07E-57 | 5.94E-56 | | AFUB_067230 |
| **AFUB_030510** | 4.476499871 | -2.207798351 | 0.931333525 | -2.370577555 | 0.017760317 | 0.026561915 | | AFUB_030510 |
| **AFUB_064030** | 1024.337843 | -2.203084915 | 0.091437279 | -24.09394669 | 2.89E-128 | 1.80E-126 | | AFUB_064030 |
| **AFUB_068810** | 29.1331945 | -2.197233639 | 0.36481685 | -6.022840334 | 1.71E-09 | 4.75E-09 | | AFUB_068810 |
| **AFUB_069290** | 554.314364 | -2.19560924 | 0.116814696 | -18.79565937 | 8.20E-79 | 1.87E-77 | | AFUB_069290 |
| **AFUB_027630** | 37.17353651 | -2.191663732 | 0.329850357 | -6.644418256 | 3.04E-11 | 9.39E-11 | | AFUB_027630 |
| **AFUB_088170** | 816.5189606 | -2.191318758 | 0.104278173 | -21.01416526 | 4.87E-98 | 1.74E-96 | | AFUB_088170 |
| **AFUB_093750** | 145.0052662 | -2.189455251 | 0.174488811 | -12.54782604 | 4.09E-36 | 3.37E-35 | | AFUB_093750 |
| **AFUB_062220** | 21.94499479 | -2.182481305 | 0.422076932 | -5.170813994 | 2.33E-07 | 5.60E-07 | | AFUB_062220 |
| **AFUB_054350** | 65.70158225 | -2.180725441 | 0.248095109 | -8.7898768 | 1.50E-18 | 6.58E-18 | | AFUB_054350 |
| **AFUB_003230** | 354.066788 | -2.176732389 | 0.138375668 | -15.73060078 | 9.33E-56 | 1.33E-54 | | AFUB_003230 |
| **AFUB_032570** | 94.09529033 | -2.172230711 | 0.22205793 | -9.782270383 | 1.34E-22 | 6.90E-22 | | AFUB_032570 |
| **AFUB_073930** | 598.6195278 | -2.171934852 | 0.104872968 | -20.71014956 | 2.81E-95 | 9.49E-94 | | AFUB_073930 |
| **AFUB_006630** | 156.380451 | -2.170794683 | 0.170291287 | -12.7475382 | 3.22E-37 | 2.76E-36 | | AFUB_006630 |
| **AFUB_056100** | 578.2574426 | -2.169720907 | 0.10768839 | -20.14814136 | 2.79E-90 | 8.55E-89 | | AFUB_056100 |
| **AFUB_090880** | 43.44911328 | -2.167987469 | 0.299130224 | -7.247637625 | 4.24E-13 | 1.45E-12 | | AFUB_090880 |
| **AFUB_090700** | 747.5785201 | -2.166988919 | 0.097681336 | -22.1842678 | 4.87E-109 | 2.20E-107 | | AFUB_090700 |
| **AFUB_040740** | 210.4924736 | -2.162482986 | 0.160845958 | -13.44443473 | 3.32E-41 | 3.20E-40 | | AFUB_040740 |
| **AFUB_005930** | 133.9705443 | -2.159915307 | 0.196020305 | -11.01883452 | 3.10E-28 | 1.97E-27 | | AFUB_005930 |
| **AFUB_098780** | 382.0618586 | -2.155387172 | 0.142527637 | -15.12259113 | 1.15E-51 | 1.48E-50 | | AFUB_098780 |
| **AFUB_015360** | 676.5800681 | -2.155134457 | 0.107471649 | -20.05305096 | 1.90E-89 | 5.66E-88 | | AFUB_015360 |
| **AFUB_036740** | 349.5903995 | -2.153969054 | 0.126185054 | -17.06992226 | 2.49E-65 | 4.35E-64 | | AFUB_036740 |
| **AFUB_065300** | 618.0454894 | -2.153895557 | 0.127991257 | -16.82845846 | 1.51E-63 | 2.53E-62 | | AFUB_065300 |
| **AFUB_087980** | 340.335053 | -2.153583903 | 0.125562776 | -17.15145187 | 6.13E-66 | 1.09E-64 | | AFUB_087980 |
| **AFUB_076810** | 245.2551672 | -2.144937142 | 0.147225105 | -14.56909907 | 4.42E-48 | 5.17E-47 | | AFUB_076810 |
| **AFUB_056680** | 16.2764129 | -2.140283803 | 0.481198088 | -4.44782275 | 8.67E-06 | 1.86E-05 | | AFUB_056680 |
| **AFUB_016030** | 540.2688677 | -2.139726388 | 0.110612692 | -19.34431171 | 2.28E-83 | 5.94E-82 | | AFUB_016030 |
| **AFUB_018330** | 23.17014119 | -2.139631305 | 0.416133905 | -5.141689442 | 2.72E-07 | 6.52E-07 | | AFUB_018330 |
| **AFUB_025530** | 476.130495 | -2.137439499 | 0.112760241 | -18.95561306 | 3.97E-80 | 9.42E-79 | | AFUB_025530 |
| **AFUB_003850** | 274.0011415 | -2.13737718 | 0.143437116 | -14.90114443 | 3.24E-50 | 4.02E-49 | | AFUB_003850 |
| **AFUB_089650** | 734.2083538 | -2.135710261 | 0.106884577 | -19.98146328 | 7.98E-89 | 2.36E-87 | | AFUB_089650 |
| **AFUB_067590** | 260.2615024 | -2.135632102 | 0.13767304 | -15.51234795 | 2.86E-54 | 3.93E-53 | | AFUB_067590 |
| **AFUB_095970** | 465.8476935 | -2.131841263 | 0.117649009 | -18.12035032 | 2.20E-73 | 4.55E-72 | | AFUB_095970 |
| **AFUB_052020** | 1162.300272 | -2.128719925 | 0.091577128 | -23.24510465 | 1.59E-119 | 8.73E-118 | | AFUB_052020 |
| **AFUB_065710** | 409.6046693 | -2.120731878 | 0.127852343 | -16.58735248 | 8.60E-62 | 1.40E-60 | | AFUB_065710 |
| **AFUB_024480** | 1027.62288 | -2.117916975 | 0.10052485 | -21.06859129 | 1.54E-98 | 5.60E-97 | | AFUB_024480 |
| **AFUB_004450** | 84.4342926 | -2.117462085 | 0.230743139 | -9.176706583 | 4.44E-20 | 2.08E-19 | | AFUB_004450 |
| **AFUB_081100** | 20.93979365 | -2.117329616 | 0.426585855 | -4.963431378 | 6.93E-07 | 1.60E-06 | | AFUB_081100 |
| **AFUB_030070** | 41.3101123 | -2.1156902 | 0.314090841 | -6.735918156 | 1.63E-11 | 5.12E-11 | | AFUB_030070 |
| **AFUB_029780** | 333.8643932 | -2.112660054 | 0.132229035 | -15.97727802 | 1.84E-57 | 2.70E-56 | | AFUB_029780 |
| **AFUB_070070** | 340.8540836 | -2.103946547 | 0.131122699 | -16.04563185 | 6.13E-58 | 9.12E-57 | | AFUB_070070 |
| **AFUB_083450** | 114.7631988 | -2.103306999 | 0.19310164 | -10.89222755 | 1.26E-27 | 7.80E-27 | | AFUB_083450 |
| **AFUB_060190** | 504.5379416 | -2.102294754 | 0.116085056 | -18.10995173 | 2.66E-73 | 5.49E-72 | | AFUB_060190 |
| **AFUB_073230** | 168.6218371 | -2.101826539 | 0.168098201 | -12.50356354 | 7.14E-36 | 5.85E-35 | | AFUB_073230 |
| **AFUB_006800** | 163.1212207 | -2.100893966 | 0.177250799 | -11.85266291 | 2.08E-32 | 1.53E-31 | | AFUB_006800 |
| **AFUB_099880** | 1579.342807 | -2.098804621 | 0.114993823 | -18.25145535 | 2.01E-74 | 4.25E-73 | | AFUB_099880 |
| **AFUB_060530** | 130.9489466 | -2.098132256 | 0.18141146 | -11.56559929 | 6.16E-31 | 4.33E-30 | | AFUB_060530 |
| **AFUB_030100** | 11.76827159 | -2.095865758 | 0.559093194 | -3.748687664 | 0.000177762 | 0.000336262 | | AFUB_030100 |
| **AFUB_020680** | 83.41337573 | -2.092790694 | 0.225722448 | -9.271522242 | 1.84E-20 | 8.68E-20 | | AFUB_020680 |
| **AFUB_035340** | 221.0119859 | -2.092665804 | 0.149665892 | -13.98224923 | 2.00E-44 | 2.15E-43 | | AFUB_035340 |
| **AFUB_084630** | 1635.428215 | -2.09228025 | 0.096936588 | -21.58400966 | 2.54E-103 | 1.01E-101 | | AFUB_084630 |
| **AFUB_058810** | 52.20508305 | -2.090984123 | 0.280682329 | -7.449646487 | 9.36E-14 | 3.32E-13 | | AFUB_058810 |
| **AFUB_003410** | 755.2737094 | -2.089926536 | 0.108324624 | -19.29318058 | 6.13E-83 | 1.57E-81 | | AFUB_003410 |
| **AFUB_053420** | 707.2368476 | -2.087768645 | 0.104354007 | -20.00659771 | 4.82E-89 | 1.43E-87 | | AFUB_053420 |
| **AFUB_081980** | 353.8217815 | -2.085611563 | 0.131005346 | -15.92004933 | 4.60E-57 | 6.70E-56 | | AFUB_081980 |

### Table S5

| | | | | | | | | |
|---|---|---|---|---|---|---|---|---|
| **AFUB_036660** | 527.3464077 | -2.079655771 | 0.107807839 | -19.29039471 | 6.47E-83 | 1.65E-81 | | AFUB_036660 |
| **AFUB_063170** | 93.32649206 | -2.077358643 | 0.213264582 | -9.740757791 | 2.02E-22 | 1.03E-21 | | AFUB_063170 |
| **AFUB_076030** | 511.7866902 | -2.072403481 | 0.121928934 | -16.99681452 | 8.67E-65 | 1.49E-63 | | AFUB_076030 |
| **AFUB_021550** | 243.3476924 | -2.069819357 | 0.144001031 | -14.37364258 | 7.57E-47 | 8.69E-46 | | AFUB_021550 |
| **AFUB_060960** | 153.0343179 | -2.065174935 | 0.180207354 | -11.45999258 | 2.10E-30 | 1.44E-29 | | AFUB_060960 |
| **AFUB_018210** | 546.999983 | -2.059316039 | 0.108592443 | -18.96371404 | 3.40E-80 | 8.10E-79 | | AFUB_018210 |
| **AFUB_085610** | 964.459283 | -2.055777128 | 0.096915666 | -21.21202077 | 7.40E-100 | 2.73E-98 | | AFUB_085610 |
| **AFUB_062330** | 134.6489236 | -2.055361632 | 0.182361755 | -11.27079322 | 1.83E-29 | 1.22E-28 | | AFUB_062330 |
| **AFUB_040910** | 591.1077015 | -2.054961922 | 0.106040255 | -19.37907366 | 1.16E-83 | 3.04E-82 | | AFUB_040910 |
| **AFUB_052060** | 3704.397926 | -2.053951744 | 0.079439695 | -25.85548377 | 2.11E-147 | 1.70E-145 | | AFUB_052060 |
| **AFUB_028110** | 372.5817545 | -2.053508819 | 0.124070466 | -16.55114939 | 1.57E-61 | 2.54E-60 | | AFUB_028110 |
| **AFUB_018590** | 112.0782204 | -2.047607728 | 0.200006852 | -10.23768788 | 1.34E-24 | 7.46E-24 | | AFUB_018590 |
| **AFUB_041260** | 348.0313247 | -2.044373446 | 0.123355474 | -16.57302576 | 1.09E-61 | 1.77E-60 | dapB | AFUB_041260 |
| **AFUB_046950** | 361.4884579 | -2.042709149 | 0.126803003 | -16.10931204 | 2.19E-58 | 3.28E-57 | | AFUB_046950 |
| **AFUB_076130** | 741.4440663 | -2.042142316 | 0.119306023 | -17.11684178 | 1.11E-65 | 1.96E-64 | | AFUB_076130 |
| **AFUB_089830** | 2028.252832 | -2.041584405 | 0.084944575 | -24.03431196 | 1.22E-127 | 7.52E-126 | | AFUB_089830 |
| **AFUB_063320** | 145.6930804 | -2.040688989 | 0.17372577 | -11.7466107 | 7.35E-32 | 5.33E-31 | | AFUB_063320 |
| **AFUB_036540** | 67.87432579 | -2.040244271 | 0.239517414 | -8.51814586 | 1.62E-17 | 6.81E-17 | | AFUB_036540 |
| **AFUB_055790** | 289.5401672 | -2.038115389 | 0.141893211 | -14.36372725 | 8.74E-47 | 9.99E-46 | | AFUB_055790 |
| **AFUB_092340** | 70.16680175 | -2.035951749 | 0.246669261 | -8.253771621 | 1.53E-16 | 6.16E-16 | | AFUB_092340 |
| **AFUB_058780** | 299.0816202 | -2.032109425 | 0.132193524 | -15.37223131 | 2.51E-53 | 3.36E-52 | | AFUB_058780 |
| **AFUB_016770** | 481.6790813 | -2.030013406 | 0.115600198 | -17.56063956 | 4.93E-69 | 9.39E-68 | | AFUB_016770 |
| **AFUB_081680** | 9.525069955 | -2.029424122 | 0.628367288 | -3.229678182 | 0.001239296 | 0.002152074 | | AFUB_081680 |
| **AFUB_003070** | 1392.164654 | -2.02927254 | 0.095617366 | -21.22284503 | 5.88E-100 | 2.19E-98 | | AFUB_003070 |
| **AFUB_090820** | 1388.694548 | -2.02771174 | 0.091340692 | -22.19943491 | 3.48E-109 | 1.58E-107 | | AFUB_090820 |
| **AFUB_063310** | 48.13118363 | -2.02664549 | 0.280019878 | -7.237505782 | 4.57E-13 | 1.56E-12 | | AFUB_063310 |
| **AFUB_017920** | 72.67428144 | -2.025883083 | 0.241514405 | -8.388249477 | 4.93E-17 | 2.03E-16 | | AFUB_017920 |
| **AFUB_021140** | 15.19215084 | -2.025567735 | 0.494359701 | -4.097356096 | 4.18E-05 | 8.42E-05 | | AFUB_021140 |
| **AFUB_026640** | 196.2990197 | -2.019844761 | 0.168524846 | -11.98544194 | 4.24E-33 | 3.18E-32 | | AFUB_026640 |
| **AFUB_093530** | 446.5631176 | -2.019635761 | 0.117949898 | -17.12282756 | 1.00E-65 | 1.78E-64 | | AFUB_093530 |
| **AFUB_008250** | 1376.756571 | -2.015019378 | 0.096205785 | -20.94488777 | 2.09E-97 | 7.35E-96 | | AFUB_008250 |
| **AFUB_094990** | 23.24757974 | -2.01296239 | 0.401282812 | -5.016318489 | 5.27E-07 | 1.23E-06 | | AFUB_094990 |
| **AFUB_035960** | 175.6368818 | -2.009531971 | 0.159149806 | -12.62666934 | 1.51E-36 | 1.26E-35 | | AFUB_035960 |
| **AFUB_086910** | 34.9606329 | -2.009105827 | 0.331902288 | -6.053305148 | 1.42E-09 | 3.95E-09 | | AFUB_086910 |
| **AFUB_097080** | 39.0185956 | -2.006603546 | 0.310908372 | -6.454002943 | 1.09E-10 | 3.25E-10 | | AFUB_097080 |
| **AFUB_098340** | 1126.953434 | -2.006156874 | 0.100425175 | -19.9766331 | 8.80E-89 | 2.59E-87 | | AFUB_098340 |
| **AFUB_088550** | 36.39459749 | -2.002871652 | 0.319194623 | -6.27476627 | 3.50E-10 | 1.01E-09 | | AFUB_088550 |
| **AFUB_061230** | 478.8911605 | 2.004480369 | 0.11811077 | 16.97119038 | 1.34E-64 | 2.30E-63 | | AFUB_061230 |
| **AFUB_078890** | 9.524126875 | 2.007838662 | 0.628280933 | 3.195765708 | 0.001394603 | 0.002412638 | | AFUB_078890 |
| **AFUB_013370** | 602.0826303 | 2.010340351 | 0.108890165 | 18.46209292 | 4.17E-76 | 9.07E-75 | | AFUB_013370 |
| **AFUB_091190** | 10.45341379 | 2.010693586 | 0.598578258 | 3.359115636 | 0.000781923 | 0.001388101 | | AFUB_091190 |
| **AFUB_097700** | 418.6689072 | 2.010875368 | 0.122407839 | 16.42766829 | 1.21E-60 | 1.92E-59 | | AFUB_097700 |
| **AFUB_046760** | 6.912596855 | 2.011827597 | 0.727478181 | 2.765481699 | 0.005683879 | 0.009105546 | | AFUB_046760 |
| **AFUB_068210** | 33.27208158 | 2.012647137 | 0.340565841 | 5.909715223 | 3.43E-09 | 9.34E-09 | | AFUB_068210 |
| **AFUB_041420** | 219.8424416 | 2.012957724 | 0.161243885 | 12.48393215 | 9.14E-36 | 7.47E-35 | | AFUB_041420 |
| **AFUB_075990** | 376.6969203 | 2.013392342 | 0.124708924 | 16.14473351 | 1.24E-58 | 1.85E-57 | | AFUB_075990 |
| **AFUB_022920** | 1019.050501 | 2.014396976 | 0.107733095 | 18.69803302 | 5.14E-78 | 1.16E-76 | | AFUB_022920 |
| **AFUB_087350** | 13.18434326 | 2.014998744 | 0.542737658 | 3.712656965 | 0.000205095 | 0.000386146 | | AFUB_087350 |
| **AFUB_085540** | 53.28090786 | 2.015252605 | 0.274723022 | 7.335579621 | 2.21E-13 | 7.67E-13 | | AFUB_085540 |
| **AFUB_015670** | 882.3108895 | 2.017770241 | 0.113414103 | 17.79117574 | 8.27E-71 | 1.64E-69 | | AFUB_015670 |
| **AFUB_082460** | 443.3100585 | 2.018101075 | 0.11566996 | 17.44706295 | 3.62E-68 | 6.77E-67 | | AFUB_082460 |
| **AFUB_039700** | 46.16002723 | 2.018688831 | 0.291049219 | 6.935901891 | 4.04E-12 | 1.31E-11 | | AFUB_039700 |
| **AFUB_067870** | 1523.12267 | 2.019789659 | 0.091613211 | 22.04692579 | 1.02E-107 | 4.50E-106 | | AFUB_067870 |
| **AFUB_092810** | 56.04819815 | 2.021658347 | 0.264333507 | 7.648135009 | 2.04E-14 | 7.44E-14 | | AFUB_092810 |
| **AFUB_060650** | 241.9594366 | 2.023655064 | 0.144905606 | 13.96533312 | 2.54E-44 | 2.72E-43 | | AFUB_060650 |
| **AFUB_059350** | 29.95718835 | 2.024965689 | 0.363592869 | 5.56932179 | 2.56E-08 | 6.60E-08 | | AFUB_059350 |
| **AFUB_028710** | 1017.851602 | 2.027119841 | 0.120080793 | 16.88129954 | 6.18E-64 | 1.04E-62 | | AFUB_028710 |
| **AFUB_013590** | 6.105695614 | 2.02915974 | 0.775557049 | 2.61639004 | 0.008886498 | 0.013885243 | | AFUB_013590 |
| **AFUB_019270** | 107.4882552 | 2.02986817 | 0.203781475 | 9.961004414 | 2.26E-23 | 1.20E-22 | | AFUB_019270 |
| **AFUB_067420** | 333.1066321 | 2.031181534 | 0.129538907 | 15.6800885 | 2.07E-55 | 2.92E-54 | | AFUB_067420 |
| **AFUB_032030** | 810.6145386 | 2.034781179 | 0.097771668 | 20.81156258 | 3.40E-96 | 1.17E-94 | | AFUB_032030 |
| **AFUB_075380** | 63.15688397 | 2.03555276 | 0.256315143 | 7.941601651 | 2.00E-15 | 7.60E-15 | | |
| **AFUB_054070** | 1470.530914 | 2.037053386 | 0.090148058 | 22.59675291 | 4.66E-113 | 2.28E-111 | | AFUB_054070 |
| **AFUB_091800** | 4.362014856 | 2.037283194 | 0.965510206 | 2.110058682 | 0.034853302 | 0.049718196 | | AFUB_091800 |
| **AFUB_087320** | 16.78788898 | 2.038322981 | 0.4726656 | 4.31239968 | 1.61E-05 | 3.39E-05 | | AFUB_087320 |
| **AFUB_019410** | 2953.440265 | 2.038739625 | 0.079566317 | 25.62314951 | 8.42E-145 | 6.60E-143 | | AFUB_019410 |
| **AFUB_038090** | 9.705465063 | 2.040163885 | 0.627718605 | 3.250124924 | 0.001153543 | 0.002011499 | | AFUB_038090 |
| **AFUB_070100** | 58.73418986 | 2.040305046 | 0.278867399 | 7.316398595 | 2.55E-13 | 8.83E-13 | | AFUB_070100 |
| **AFUB_067730** | 4.512334848 | 2.041170268 | 0.926287948 | 2.203602316 | 0.027552324 | 0.039941958 | | AFUB_067730 |
| **AFUB_097620** | 5.271000702 | 2.041707779 | 0.841392124 | 2.426582946 | 0.015241766 | 0.023031152 | | AFUB_097620 |
| **AFUB_055320** | 44.93427543 | 2.043463164 | 0.307197379 | 6.651955068 | 2.89E-11 | 8.95E-11 | | AFUB_055320 |
| **AFUB_003780** | 679.6049045 | 2.043681037 | 0.103166537 | 19.80953413 | 2.46E-87 | 7.11E-86 | | AFUB_003780 |
| **AFUB_066430** | 119.8365794 | 2.044588018 | 0.190674406 | 10.7229285 | 7.94E-27 | 4.78E-26 | | AFUB_066430 |
| **AFUB_100310** | 493.4324276 | 2.048129376 | 0.11271857 | 18.17029235 | 8.87E-74 | 1.86E-72 | | AFUB_100310 |
| **AFUB_044530** | 12.51959215 | 2.049661205 | 0.555560791 | 3.6893554 | 0.000224823 | 0.000421314 | | AFUB_044530 |
| **AFUB_015690** | 146.0681036 | 2.051615715 | 0.175662328 | 11.67931528 | 1.63E-31 | 1.16E-30 | | AFUB_015690 |
| **AFUB_072740** | 256.384104 | 2.053595788 | 0.144564325 | 14.20541199 | 8.48E-46 | 9.43E-45 | | AFUB_072740 |
| **AFUB_018520** | 950.7446825 | 2.054649328 | 0.094150903 | 21.82293816 | 1.41E-105 | 5.89E-104 | | AFUB_018520 |
| **AFUB_013750** | 48.1861972 | 2.055703491 | 0.291546971 | 7.051019887 | 1.78E-12 | 5.89E-12 | | AFUB_013750 |
| **AFUB_055650** | 1609.943371 | 2.055771636 | 0.095025522 | 21.63388933 | 8.62E-104 | 3.46E-102 | | AFUB_055650 |
| **AFUB_041060** | 394.6305311 | 2.057223949 | 0.148306535 | 13.87143159 | 9.44E-44 | 9.86E-43 | | AFUB_041060 |
| **AFUB_010660** | 5.316984803 | 2.057413807 | 0.875056014 | 2.351179553 | 0.018714001 | 0.027875733 | | AFUB_010660 |
| **AFUB_055190** | 22.26226488 | 2.061183323 | 0.419887845 | 4.908890194 | 9.16E-07 | 2.10E-06 | | AFUB_055190 |
| **AFUB_009450** | 116.3689554 | 2.061471634 | 0.20608805 | 10.00286838 | 1.48E-23 | 7.89E-23 | | AFUB_009450 |
| **AFUB_006140** | 24.07279059 | 2.062166289 | 0.39803674 | 5.180844081 | 2.21E-07 | 5.33E-07 | | AFUB_006140 |
| **AFUB_080150** | 19.77961277 | 2.062700769 | 0.449580977 | 4.588051707 | 4.47E-06 | 9.79E-06 | | AFUB_080150 |
| **AFUB_018840** | 4.389424185 | 2.064038294 | 0.945690808 | 2.182572016 | 0.029067342 | 0.041986645 | | AFUB_018840 |
| **AFUB_064080** | 6.309630968 | 2.065021096 | 0.782173216 | 2.640107143 | 0.008287982 | 0.013009121 | | AFUB_064080 |
| **AFUB_051520** | 731.3865407 | 2.065261643 | 0.120356151 | 17.15958528 | 5.33E-66 | 9.51E-65 | | AFUB_051520 |
| **AFUB_085960** | 103.1461659 | 2.065903753 | 0.202121663 | 10.22109022 | 1.60E-24 | 8.84E-24 | | AFUB_085960 |
| **AFUB_091240** | 30359.84174 | 2.071205784 | 0.071605594 | 28.92519503 | 5.76E-184 | 7.81E-182 | | AFUB_091240 |
| **AFUB_091360** | 142.149625 | 2.071442849 | 0.178134008 | 11.62856478 | 2.95E-31 | 2.09E-30 | | AFUB_091360 |
| **AFUB_076040** | 1963.862108 | 2.073567848 | 0.090390648 | 22.94007066 | 1.85E-116 | 9.39E-115 | | AFUB_076040 |
| **AFUB_034180** | 23.72644897 | 2.074051782 | 0.414333033 | 5.005760142 | 5.56E-07 | 1.30E-06 | | AFUB_034180 |
| **AFUB_022390** | 167.1286286 | 2.074204361 | 0.16471068 | 12.59301679 | 2.31E-36 | 1.92E-35 | | AFUB_022390 |
| **AFUB_092970** | 786.7138958 | 2.074848722 | 0.103092719 | 20.12604518 | 4.36E-90 | 1.31E-88 | | AFUB_092970 |
| **AFUB_002370** | 957.4460924 | 2.076664199 | 0.108554362 | 19.13017737 | 1.42E-81 | 3.54E-80 | | AFUB_002370 |
| **AFUB_096580** | 36.0899351 | 2.078302756 | 0.33045939 | 6.289132101 | 3.19E-10 | 9.25E-10 | | AFUB_096580 |
| **AFUB_044540** | 24.38716549 | 2.078706822 | 0.401168218 | 5.181633854 | 2.20E-07 | 5.30E-07 | | AFUB_044540 |
| **AFUB_090090** | 566.9094527 | 2.083668975 | 0.113467459 | 18.36358189 | 2.57E-75 | 5.49E-74 | | AFUB_090090 |
| **AFUB_064150** | 730.5684068 | 2.083761939 | 0.100912326 | 20.64923116 | 9.92E-95 | 3.33E-93 | | AFUB_064150 |
| **AFUB_057640** | 11.7874558 | 2.090041781 | 0.569136084 | 3.672305872 | 0.000240372 | 0.000449316 | | AFUB_057640 |
| **AFUB_062180** | 5.471650855 | 2.090635743 | 0.839405667 | 2.490614284 | 0.012752248 | 0.019477042 | | AFUB_062180 |
| **AFUB_079780** | 53.9235166 | 2.091183317 | 0.274814145 | 7.609445713 | 2.75E-14 | 9.98E-14 | | AFUB_079780 |
| **AFUB_041350** | 168.4498524 | 2.091265785 | 0.173326954 | 12.06543895 | 1.61E-33 | 1.23E-32 | | AFUB_041350 |
| **AFUB_044910** | 21.08468854 | 2.092776208 | 0.431948167 | 4.844970688 | 1.27E-06 | 2.89E-06 | | AFUB_044910 |
| **AFUB_086760** | 41.93244533 | 2.097582051 | 0.305502986 | 6.865995258 | 6.60E-12 | 2.12E-11 | | AFUB_086760 |
| **AFUB_100430** | 102.2158054 | 2.100838633 | 0.206281737 | 10.18431717 | 2.33E-24 | 1.28E-23 | | AFUB_100430 |
| **AFUB_031730** | 409.7077158 | 2.101184016 | 0.122252125 | 17.18730052 | 3.31E-66 | 5.93E-65 | | AFUB_031730 |
| **AFUB_063750** | 3388.093019 | 2.102539458 | 0.097441733 | 21.57740209 | 2.93E-103 | 1.16E-101 | | AFUB_063750 |
| **AFUB_020640** | 117.1993576 | 2.10351405 | 0.197935745 | 10.6272571 | 2.23E-26 | 1.32E-25 | | AFUB_020640 |
| **AFUB_033130** | 23.83610577 | 2.10408051 | 0.406269341 | 5.179028528 | 2.23E-07 | 5.37E-07 | | AFUB_033130 |
| **AFUB_045790** | 5.557103932 | 2.104466615 | 0.841298893 | 2.501449405 | 0.012368611 | 0.018921089 | | AFUB_045790 |
| **AFUB_078230** | 209.2920221 | 2.104710553 | 0.154596927 | 13.61418104 | 3.30E-42 | 3.28E-41 | | AFUB_078230 |
| **AFUB_025700** | 421.8246536 | 2.105026897 | 0.118583016 | 17.75150414 | 1.68E-70 | 3.31E-69 | | AFUB_025700 |
| **AFUB_077450** | 551.6477267 | 2.109817959 | 0.113111712 | 18.65251551 | 1.20E-77 | 2.68E-76 | | AFUB_077450 |
| **AFUB_053550** | 2545.113564 | 2.110192773 | 0.096190002 | 21.93775571 | 1.13E-106 | 4.85E-105 | | AFUB_053550 |
| **AFUB_065110** | 5732.017156 | 2.11040836 | 0.083012806 | 25.42268437 | 1.42E-142 | 1.07E-140 | | AFUB_065110 |
| **AFUB_069970** | 69.41509586 | 2.112314625 | 0.247230762 | 8.543898867 | 1.30E-17 | 5.48E-17 | | AFUB_069970 |
| **AFUB_032070** | 509.3692625 | 2.113644778 | 0.115529738 | 18.29524427 | 9.03E-75 | 1.91E-73 | | AFUB_032070 |
| **AFUB_076750** | 14.56459123 | 2.11367617 | 0.515337418 | 4.101538324 | 4.10E-05 | 8.28E-05 | | AFUB_076750 |
| **AFUB_037180** | 11.13000058 | 2.113787014 | 0.583607412 | 3.621933122 | 0.00029241 | 0.000542171 | | AFUB_037180 |
| **AFUB_015000** | 3123.172707 | 2.116089456 | 0.098082242 | 21.574644 | 3.11E-103 | 1.22E-101 | | AFUB_015000 |
| **AFUB_057440** | 5.509545436 | 2.11853923 | 0.837625401 | 2.529220374 | 0.011431622 | 0.0175546 | | AFUB_057440 |
| **AFUB_067650** | 1422.86136 | 2.118639482 | 0.100657241 | 21.04805843 | 2.38E-98 | 8.60E-97 | | AFUB_067650 |
| **AFUB_030800** | 35.13280814 | 2.119008995 | 0.343707122 | 6.165158826 | 7.04E-10 | 2.00E-09 | | AFUB_030800 |
| **AFUB_059070** | 56.45290084 | 2.120849733 | 0.282979927 | 7.49470027 | 6.65E-14 | 2.37E-13 | | AFUB_059070 |
| **AFUB_028820** | 6.488475976 | 2.122855533 | 0.774682479 | 2.740291139 | 0.006138478 | 0.009790279 | | AFUB_028820 |
| **AFUB_047320** | 14.75407763 | 2.12358136 | 0.517330335 | 4.104884666 | 4.05E-05 | 8.16E-05 | | AFUB_047320 |
| **AFUB_084840** | 23.22703038 | 2.123821701 | 0.410434888 | 5.174564258 | 2.28E-07 | 5.50E-07 | | AFUB_084840 |
| **AFUB_048830** | 5.64999478 | 2.131406835 | 0.829172499 | 2.570522826 | 0.010154513 | 0.015713649 | | AFUB_048830 |
| **AFUB_027240** | 1277.402471 | 2.132430266 | 0.089604148 | 23.79834318 | 3.47E-125 | 2.08E-123 | arg2 | AFUB_027240 |
| **AFUB_025450** | 22.43907643 | 2.136407737 | 0.417638866 | 5.115442816 | 3.13E-07 | 7.45E-07 | | AFUB_025450 |
| **AFUB_065860** | 561.412222 | 2.137259748 | 0.139417599 | 15.32991365 | 4.83E-53 | 6.42E-52 | | AFUB_065860 |
| **AFUB_035080** | 127.7074586 | 2.142238592 | 0.191071537 | 11.21170963 | 3.57E-29 | 2.35E-28 | | AFUB_035080 |
| **AFUB_100720** | 220.8513887 | 2.143763809 | 0.154667535 | 13.86046402 | 1.10E-43 | 1.15E-42 | | AFUB_100720 |
| **AFUB_014690** | 7533.445404 | 2.147019469 | 0.200929761 | 10.68542292 | 1.19E-26 | 7.12E-26 | | AFUB_014690 |
| **AFUB_006520** | 17.74923647 | 2.147540585 | 0.475608917 | 4.515349713 | 6.32E-06 | 1.37E-05 | | AFUB_006520 |
| **AFUB_067340** | 263.6055567 | 2.147709925 | 0.14088919 | 15.24396532 | 1.81E-52 | 2.36E-51 | | AFUB_067340 |
| **AFUB_095250** | 358.6319641 | 2.148168153 | 0.125901072 | 17.06234995 | 2.83E-65 | 4.94E-64 | | AFUB_095250 |
| **AFUB_027560** | 549.4252543 | 2.149026702 | 0.111297315 | 19.30888178 | 4.52E-83 | 1.17E-81 | | AFUB_027560 |
| **AFUB_014270** | 265.707889 | 2.152054947 | 0.143506619 | 14.99620687 | 7.77E-51 | 9.78E-50 | | AFUB_014270 |
| **AFUB_045460** | 12.27212368 | 2.152079776 | 0.559954645 | 3.843310876 | 0.000121386 | 0.000234024 | | AFUB_045460 |
| **AFUB_015250** | 385.9111408 | 2.152511856 | 0.129303829 | 16.64693048 | 3.19E-62 | 5.21E-61 | | AFUB_015250 |
| **AFUB_045240** | 18.99941566 | 2.153585115 | 0.463639816 | 4.64495291 | 3.40E-06 | 7.52E-06 | | AFUB_045240 |
| **AFUB_047640** | 315.2165769 | 2.158331364 | 0.132709424 | 16.26358773 | 1.79E-59 | 2.72E-58 | | AFUB_047640 |
| **AFUB_076700** | 12223.9214 | 2.159367309 | 0.093533816 | 23.08648786 | 6.33E-118 | 3.31E-116 | | AFUB_076700 |
| **AFUB_050510** | 301.4347371 | 2.1602512 | 0.136038054 | 15.87975667 | 8.75E-57 | 1.27E-55 | | AFUB_050510 |
| **AFUB_053300** | 288.3483125 | 2.160669057 | 0.138950847 | 15.54988048 | 1.59E-54 | 2.19E-53 | | AFUB_053300 |
| **AFUB_030420** | 1302.551229 | 2.160921711 | 0.105748135 | 20.4346082 | 8.24E-93 | 2.66E-91 | | AFUB_030420 |
| **AFUB_034420** | 7.740145362 | 2.163662886 | 0.725475679 | 2.982405817 | 0.002859926 | 0.004755681 | | AFUB_034420 |
| **AFUB_094690** | 160.8572583 | 2.163679537 | 0.173497943 | 12.47092328 | 1.08E-35 | 8.76E-35 | | AFUB_094690 |
| **AFUB_023060** | 8.600638434 | 2.164569556 | 0.673909118 | 3.21196063 | 0.001318324 | 0.002286013 | | AFUB_023060 |
| **AFUB_021010** | 4.667217279 | 2.164674409 | 0.92012848 | 2.352578424 | 0.018643758 | 0.027786953 | | AFUB_021010 |
| **AFUB_045330** | 38.21688954 | 2.165625151 | 0.328696534 | 6.588524451 | 4.44E-11 | 1.35E-10 | | AFUB_045330 |

### Table S5

| | | | | | | | | |
|---|---|---|---|---|---|---|---|---|
| **AFUB_035190** | 178.3094238 | 2.166415505 | 0.173910265 | 12.45708819 | 1.28E-35 | 1.04E-34 | | AFUB_035190 |
| **AFUB_019490** | 153.8580057 | 2.16651493 | 0.173880214 | 12.45981292 | 1.24E-35 | 1.01E-34 | | AFUB_019490 |
| **AFUB_076990** | 19.13977919 | 2.166966928 | 0.459327411 | 4.717695651 | 2.39E-06 | 5.33E-06 | | AFUB_076990 |
| **AFUB_016910** | 6.763452224 | 2.167520125 | 0.808389234 | 2.681282772 | 0.007334051 | 0.011591097 | | AFUB_016910 |
| **AFUB_095700** | 30.35986722 | 2.170125836 | 0.3689969 | 5.881149229 | 4.07E-09 | 1.11E-08 | | AFUB_095700 |
| **AFUB_001820** | 678.9209671 | 2.17077944 | 0.110406004 | 19.66178791 | 4.58E-86 | 1.29E-84 | | AFUB_001820 |
| **AFUB_058470** | 42.00349389 | 2.176930017 | 0.310513124 | 7.010750432 | 2.37E-12 | 7.81E-12 | | AFUB_058470 |
| **AFUB_087370** | 8.562868632 | 2.177387154 | 0.682601002 | 3.189838788 | 0.001423522 | 0.002457813 | | AFUB_087370 |
| **AFUB_034110** | 32.43794652 | 2.178872719 | 0.349065212 | 6.242021968 | 4.32E-10 | 1.24E-09 | | AFUB_034110 |
| **AFUB_092160** | 44.32948967 | 2.179048884 | 0.31156617 | 6.993855864 | 2.67E-12 | 8.76E-12 | | AFUB_092160 |
| **AFUB_101670** | 28.65780285 | 2.180977424 | 0.373825114 | 5.834218575 | 5.40E-09 | 1.46E-08 | | AFUB_101670 |
| **AFUB_045680** | 96.32952862 | 2.181559784 | 0.218097907 | 10.00266261 | 1.48E-23 | 7.90E-23 | | AFUB_045680 |
| **AFUB_099370** | 3189.020156 | 2.182046538 | 0.115990047 | 18.81236009 | 5.98E-79 | 1.38E-77 | | AFUB_099370 |
| **AFUB_070750** | 6.773997779 | 2.18451915 | 0.759194406 | 2.877417341 | 0.004009449 | 0.006539447 | | AFUB_070750 |
| **AFUB_047610** | 31.64102104 | 2.191976754 | 0.356231019 | 6.153245041 | 7.59E-10 | 2.15E-09 | | AFUB_047610 |
| **AFUB_100850** | 15.41846449 | 2.192890253 | 0.511224776 | 4.28948352 | 1.79E-05 | 3.74E-05 | | AFUB_100850 |
| **AFUB_079420** | 28.96130996 | 2.193548753 | 0.374290466 | 5.860552032 | 4.61E-09 | 1.25E-08 | | AFUB_079420 |
| **AFUB_051790** | 91.31580319 | 2.194097126 | 0.228595891 | 9.598147713 | 8.14E-22 | 4.06E-21 | | AFUB_051790 |
| **AFUB_060410** | 8.657390315 | 2.194178208 | 0.671715913 | 3.266527061 | 0.001088754 | 0.001902999 | | AFUB_060410 |
| **AFUB_101100** | 11.65109413 | 2.195639422 | 0.586357892 | 3.744538024 | 0.000180726 | 0.0003416 | | AFUB_101100 |
| **AFUB_056020** | 82.03318214 | 2.19660582 | 0.227144474 | 9.670522846 | 4.02E-22 | 2.03E-21 | | AFUB_056020 |
| **AFUB_030750** | 130.2449659 | 2.19715498 | 0.187324389 | 11.7291453 | 9.04E-32 | 6.52E-31 | | AFUB_030750 |
| **AFUB_092260** | 321.7885089 | 2.198012653 | 0.134036728 | 16.39858482 | 1.96E-60 | 3.07E-59 | | AFUB_092260 |
| **AFUB_013600** | 46.34786283 | 2.20226055 | 0.304262604 | 7.238025708 | 4.55E-13 | 1.56E-12 | | AFUB_013600 |
| **AFUB_087070** | 14.47197625 | 2.202352415 | 0.520816085 | 4.228656673 | 2.35E-05 | 4.86E-05 | | AFUB_087070 |
| **AFUB_101180** | 99.41762786 | 2.20304574 | 0.210229043 | 10.47926447 | 1.08E-25 | 6.22E-25 | | AFUB_101180 |
| **AFUB_043770** | 3386.608428 | 2.203338769 | 0.078696914 | 27.99777851 | 1.73E-172 | 1.94E-170 | | AFUB_043770 |
| **AFUB_055490** | 713.3291496 | 2.204581996 | 0.101732206 | 21.67044332 | 3.90E-104 | 1.59E-102 | | AFUB_055490 |
| **AFUB_063740** | 4.844572098 | 2.207255704 | 0.897386798 | 2.459648068 | 0.013907332 | 0.021127392 | | AFUB_063740 |
| **AFUB_059520** | 28.106193 | 2.210241044 | 0.377645173 | 5.852692432 | 4.84E-09 | 1.31E-08 | | AFUB_059520 |
| **AFUB_085270** | 7.712977554 | 2.211073422 | 0.723626486 | 3.055545181 | 0.002246518 | 0.003787166 | | AFUB_085270 |
| **AFUB_066500** | 30.09450087 | 2.213697534 | 0.368198636 | 6.012237197 | 1.83E-09 | 5.06E-09 | | AFUB_066500 |
| **AFUB_021330** | 1494.986849 | 2.214741845 | 0.117152968 | 18.90470113 | 1.04E-79 | 2.45E-78 | | AFUB_021330 |
| **AFUB_034540** | 1521.159388 | 2.217303513 | 0.085185976 | 26.02897361 | 2.33E-149 | 1.97E-147 | plb3 | AFUB_034540 |
| **AFUB_022280** | 72.98216217 | 2.219096599 | 0.246155459 | 9.015020871 | 1.97E-19 | 8.95E-19 | | AFUB_022280 |
| **AFUB_098630** | 381.763915 | 2.219369467 | 0.12752618 | 17.40324583 | 7.80E-68 | 1.44E-66 | | AFUB_098630 |
| **AFUB_088320** | 103.5141178 | 2.220104413 | 0.209163791 | 10.61419095 | 2.56E-26 | 1.51E-25 | | AFUB_088320 |
| **AFUB_101750** | 208.1874786 | 2.220696705 | 0.157796116 | 14.07320255 | 5.55E-45 | 6.01E-44 | | AFUB_101750 |
| **AFUB_013490** | 8.787221798 | 2.222360523 | 0.666187851 | 3.335936731 | 0.000850125 | 0.00150474 | | AFUB_013490 |
| **AFUB_051660** | 7.85032715 | 2.22343131 | 0.729368538 | 3.048433262 | 0.00230038 | 0.003875256 | | AFUB_051660 |
| **AFUB_028430** | 6.973364473 | 2.225714734 | 0.768219084 | 2.897239577 | 0.003764621 | 0.006158895 | | AFUB_028430 |
| **AFUB_084820** | 236.4003634 | 2.22816863 | 0.151189252 | 14.73761261 | 3.70E-49 | 4.45E-48 | | AFUB_084820 |
| **AFUB_063690** | 155.4695088 | 2.228903996 | 0.175485169 | 12.70138101 | 5.81E-37 | 4.93E-36 | | AFUB_063690 |
| **AFUB_059550** | 343.3511315 | 2.229091539 | 0.135158933 | 16.49237299 | 4.16E-61 | 6.65E-60 | | AFUB_059550 |
| **AFUB_050600** | 12.93142918 | 2.22941689 | 0.557653458 | 3.997853609 | 6.39E-05 | 0.000126552 | | AFUB_050600 |
| **AFUB_020230** | 30.51571791 | 2.231393679 | 0.363586295 | 6.137177632 | 8.40E-10 | 2.37E-09 | | AFUB_020230 |
| **AFUB_081240** | 4.991907474 | 2.23413343 | 0.892007763 | 2.504612092 | 0.012258575 | 0.018758716 | | AFUB_081240 |
| **AFUB_050290** | 844.9179231 | 2.234827558 | 0.105341893 | 21.21499328 | 6.94E-100 | 2.57E-98 | | AFUB_050290 |
| **AFUB_037520** | 95.61897314 | 2.236901594 | 0.221965943 | 10.07767931 | 6.93E-24 | 3.73E-23 | | AFUB_037520 |
| **AFUB_040420** | 125.8241499 | 2.237264358 | 0.196092608 | 11.40922334 | 3.76E-30 | 2.56E-29 | | AFUB_040420 |
| **AFUB_089190** | 798.4153642 | 2.237530511 | 0.100117631 | 22.3490157 | 1.23E-110 | 5.80E-109 | | AFUB_089190 |
| **AFUB_021540** | 37.65067927 | 2.240954908 | 0.329404575 | 6.803047312 | 1.02E-11 | 3.25E-11 | | AFUB_021540 |
| **AFUB_095140** | 4.909598952 | 2.242038002 | 0.895722172 | 2.503050691 | 0.012312791 | 0.018838687 | | AFUB_095140 |
| **AFUB_001110** | 70.0341369 | 2.242508641 | 0.246758213 | 9.087878437 | 1.01E-19 | 4.65E-19 | | AFUB_001110 |
| **AFUB_043760** | 6535.176696 | 2.246538386 | 0.076523795 | 29.35738344 | 1.92E-189 | 2.85E-187 | | AFUB_043760 |
| **AFUB_079250** | 106.771827 | 2.251015148 | 0.211535977 | 10.64128752 | 1.91E-26 | 1.13E-25 | | AFUB_079250 |
| **AFUB_016940** | 7.061721823 | 2.252171146 | 0.753944562 | 2.987184019 | 0.002815602 | 0.004687476 | | AFUB_016940 |
| **AFUB_070780** | 85.33495098 | 2.253338168 | 0.229441229 | 9.820981958 | 9.14E-23 | 4.75E-22 | | AFUB_070780 |
| **AFUB_011700** | 5.046246518 | 2.254327138 | 0.899346158 | 2.506628974 | 0.012188858 | 0.018666853 | | AFUB_011700 |
| **AFUB_042780** | 6.033461679 | 2.254845297 | 0.831634137 | 2.711342882 | 0.006701129 | 0.010634393 | | |
| **AFUB_050860** | 5858.343018 | 2.255362533 | 0.092027736 | 24.5074217 | 1.23E-132 | 8.13E-131 | | AFUB_050860 |
| **AFUB_072060** | 519.628464 | 2.255621747 | 0.110493758 | 20.41401957 | 1.26E-92 | 4.02E-91 | | AFUB_072060 |
| **AFUB_086200** | 644.6225328 | 2.256765385 | 0.104293366 | 21.63862838 | 7.78E-104 | 3.14E-102 | | AFUB_086200 |
| **AFUB_092680** | 32.97245979 | 2.259520064 | 0.356952479 | 6.330030457 | 2.45E-10 | 7.15E-10 | | AFUB_092680 |
| **AFUB_033020** | 60.31434075 | 2.261833717 | 0.271248552 | 8.338601993 | 7.52E-17 | 3.06E-16 | | AFUB_033020 |
| **AFUB_047480** | 14.05554892 | 2.262910954 | 0.534720313 | 4.23195248 | 2.32E-05 | 4.79E-05 | | AFUB_047480 |
| **AFUB_028340** | 11.1001847 | 2.263291461 | 0.599564345 | 3.774893355 | 0.000160076 | 0.000304059 | | AFUB_028340 |
| **AFUB_045520** | 3.975092 | 2.264055759 | 0.996198679 | 2.272695003 | 0.023044566 | 0.033870872 | | AFUB_045520 |
| **AFUB_064460** | 39.956543 | 2.26531132 | 0.323245994 | 7.008010497 | 2.42E-12 | 7.95E-12 | | AFUB_064460 |
| **AFUB_018270** | 929.611528 | 2.267242187 | 0.110264094 | 20.56192642 | 6.02E-94 | 1.99E-92 | | AFUB_018270 |
| **AFUB_057550** | 10.14896786 | 2.268496203 | 0.636152623 | 3.565962192 | 0.000362524 | 0.000665074 | | AFUB_057550 |
| **AFUB_045530** | 37.20191089 | 2.270241365 | 0.332672811 | 6.824246804 | 8.84E-12 | 2.82E-11 | | AFUB_045530 |
| **AFUB_099430** | 86.24607485 | 2.278634935 | 0.236446584 | 9.636996642 | 5.58E-22 | 2.80E-21 | | AFUB_099430 |
| **AFUB_053630** | 396.0184693 | 2.279151667 | 0.127692486 | 17.84875322 | 2.96E-71 | 5.97E-70 | | AFUB_053630 |
| **AFUB_038480** | 166.6961489 | 2.279304969 | 0.170089477 | 13.40062306 | 6.00E-41 | 5.72E-40 | | AFUB_038480 |
| **AFUB_090790** | 3776.775419 | 2.281634036 | 0.082004109 | 27.82341125 | 2.26E-170 | 2.45E-168 | | AFUB_090790 |
| **AFUB_098960** | 7.198324207 | 2.283801338 | 0.764546621 | 2.987131557 | 0.002816085 | 0.004687476 | | AFUB_098960 |
| **AFUB_046520** | 111.3322748 | 2.285608926 | 0.202017295 | 11.31392698 | 1.12E-29 | 7.50E-29 | | AFUB_046520 |
| **AFUB_017200** | 7.131176692 | 2.286473422 | 0.770779565 | 2.966442708 | 0.003012665 | 0.00499141 | | AFUB_017200 |
| **AFUB_036100** | 750.3857467 | 2.289231672 | 0.112060886 | 20.42846315 | 9.34E-93 | 3.01E-91 | | AFUB_036100 |
| **AFUB_075370** | 19.41961889 | 2.294495577 | 0.45518727 | 5.040772735 | 4.64E-07 | 1.09E-06 | | |
| **AFUB_012540** | 107.1120884 | 2.294548734 | 0.209999623 | 10.92644218 | 8.62E-28 | 5.39E-27 | | AFUB_012540 |
| **AFUB_084650** | 36.91203825 | 2.295037217 | 0.336534709 | 6.819615198 | 9.13E-12 | 2.91E-11 | | AFUB_084650 |
| **AFUB_090530** | 20.52244883 | 2.298795638 | 0.446943504 | 5.143369621 | 2.70E-07 | 6.46E-07 | | AFUB_090530 |
| **AFUB_017220** | 8.262075743 | 2.303049889 | 0.705531196 | 3.264277896 | 0.001097435 | 0.001917128 | | AFUB_017220 |
| **AFUB_086010** | 748.3994653 | 2.304549957 | 0.11197826 | 20.58033367 | 4.12E-94 | 1.37E-92 | | AFUB_086010 |
| **AFUB_062160** | 6.157476207 | 2.305233627 | 0.813488586 | 2.833762719 | 0.004600348 | 0.007448992 | | AFUB_062160 |
| **AFUB_046150** | 10.26388023 | 2.309606843 | 0.632104383 | 3.653837728 | 0.00025835 | 0.000481986 | | AFUB_046150 |
| **AFUB_086100** | 103.5736491 | 2.30995994 | 0.214474287 | 10.77033508 | 4.75E-27 | 2.90E-26 | | AFUB_086100 |
| **AFUB_017070** | 5.128554666 | 2.310191675 | 0.877745889 | 2.631959551 | 0.008489397 | 0.013301436 | | AFUB_017070 |
| **AFUB_069420** | 4141.213007 | 2.314467816 | 0.085606645 | 27.03607681 | 5.57E-161 | 5.65E-159 | | AFUB_069420 |
| **AFUB_094840** | 20.69470839 | 2.316530489 | 0.443717418 | 5.220733731 | 1.78E-07 | 4.33E-07 | | AFUB_094840 |
| **AFUB_094810** | 53.88138062 | 2.324368156 | 0.287513222 | 8.084386994 | 6.25E-16 | 2.44E-15 | | AFUB_094810 |
| **AFUB_000350** | 24.97402373 | 2.325080181 | 0.40720328 | 5.70987586 | 1.13E-08 | 2.99E-08 | | AFUB_000350 |
| **AFUB_063970** | 1929.059474 | 2.327788582 | 0.088959494 | 26.16683698 | 6.34E-151 | 5.55E-149 | | AFUB_063970 |
| **AFUB_084290** | 16.56347141 | 2.328979155 | 0.501111889 | 4.647623028 | 3.36E-06 | 7.42E-06 | | AFUB_084290 |
| **AFUB_073780** | 13.67153874 | 2.329155818 | 0.551008072 | 4.227081122 | 2.37E-05 | 4.89E-05 | | AFUB_073780 |
| **AFUB_099630** | 22.58296594 | 2.330449415 | 0.447460624 | 5.20816646 | 1.91E-07 | 4.62E-07 | | AFUB_099630 |
| **AFUB_084520** | 166.3536303 | 2.331824447 | 0.177808049 | 13.11427946 | 2.73E-39 | 2.46E-38 | | AFUB_084520 |
| **AFUB_014050** | 22.06783882 | 2.336162339 | 0.433066448 | 5.394466257 | 6.87E-08 | 1.72E-07 | | AFUB_014050 |
| **AFUB_012000** | 3140.16768 | 2.336545704 | 0.09487926 | 24.62651682 | 6.57E-134 | 4.40E-132 | | AFUB_012000 |
| **AFUB_031340** | 10.46108146 | 2.338676948 | 0.645950098 | 3.62052263 | 0.000294009 | 0.000544925 | | AFUB_031340 |
| **AFUB_047110** | 18.72557968 | 2.341924706 | 0.476438674 | 4.915479864 | 8.86E-07 | 2.04E-06 | | AFUB_047110 |
| **AFUB_079770** | 886.303233 | 2.343226333 | 0.104104337 | 22.50844112 | 3.43E-112 | 1.64E-110 | | AFUB_079770 |
| **AFUB_048180** | 7408.119558 | 2.343591978 | 0.106325986 | 22.04157288 | 1.15E-107 | 5.04E-106 | eglC | AFUB_048180 |
| **AFUB_044960** | 145.8042374 | 2.350185165 | 0.181045421 | 12.98119084 | 1.56E-38 | 1.39E-37 | | AFUB_044960 |
| **AFUB_037310** | 3980.838129 | 2.350327061 | 0.085590293 | 27.46020571 | 5.25E-166 | 5.56E-164 | | AFUB_037310 |
| **AFUB_034520** | 11.59027935 | 2.352761474 | 0.595439279 | 3.951303782 | 7.77E-05 | 0.000152852 | | AFUB_034520 |
| **AFUB_048330** | 11.60397144 | 2.354305023 | 0.603200417 | 3.903022871 | 9.50E-05 | 0.000185268 | | AFUB_048330 |
| **AFUB_058760** | 273.9968507 | 2.354473866 | 0.144904473 | 16.24845538 | 2.29E-59 | 3.48E-58 | | AFUB_058760 |
| **AFUB_097780** | 11.54664778 | 2.354645876 | 0.596160578 | 3.949683969 | 7.83E-05 | 0.000153858 | | |
| **AFUB_000670** | 12.80799073 | 2.355868676 | 0.580974552 | 4.055029035 | 5.01E-05 | 0.000100277 | | AFUB_000670 |
| **AFUB_092780** | 1475.695579 | 2.358487082 | 0.093510424 | 25.22164888 | 2.32E-140 | 1.71E-138 | | |
| **AFUB_080970** | 86.75083285 | 2.360254737 | 0.237224649 | 9.949449805 | 2.54E-23 | 1.34E-22 | | AFUB_080970 |
| **AFUB_057970** | 41.28948041 | 2.36272347 | 0.320362662 | 7.375152438 | 1.64E-13 | 5.74E-13 | | AFUB_057970 |
| **AFUB_003610** | 5.370911198 | 2.362856238 | 0.876104832 | 2.697001718 | 0.006996691 | 0.011076082 | | AFUB_003610 |
| **AFUB_008420** | 133.6378464 | 2.365652516 | 0.190304334 | 12.43089146 | 1.78E-35 | 1.43E-34 | | AFUB_008420 |
| **AFUB_046690** | 13.77942444 | 2.366877577 | 0.556115223 | 4.256092043 | 2.08E-05 | 4.32E-05 | | AFUB_046690 |
| **AFUB_059150** | 44.80050698 | 2.367067193 | 0.311062326 | 7.609623531 | 2.75E-14 | 9.97E-14 | | AFUB_059150 |
| **AFUB_024580** | 1052.70786 | 2.370709352 | 0.124875761 | 18.98454377 | 2.29E-80 | 5.50E-79 | | AFUB_024580 |
| **AFUB_033030** | 95.08203662 | 2.372231226 | 0.222806784 | 10.64703322 | 1.80E-26 | 1.07E-25 | | AFUB_033030 |
| **AFUB_045050** | 4.191901912 | 2.372487364 | 1.017452863 | 2.331790936 | 0.019711692 | 0.029262342 | | AFUB_045050 |
| **AFUB_093110** | 802.0074205 | 2.37395751 | 0.111405502 | 21.30915866 | 9.34E-101 | 3.51E-99 | | AFUB_093110 |
| **AFUB_076480** | 11.85077148 | 2.375363755 | 0.602536931 | 3.942270811 | 8.07E-05 | 0.0001585 | | AFUB_076480 |
| **AFUB_070540** | 6.41880517 | 2.375940633 | 0.803967401 | 2.955269867 | 0.003123957 | 0.005166028 | | AFUB_070540 |
| **AFUB_042300** | 12.7975951 | 2.376854217 | 0.595090645 | 3.994104491 | 6.49E-05 | 0.000128544 | | AFUB_042300 |
| **AFUB_027330** | 7.533696365 | 2.378643868 | 0.73499481 | 3.236273012 | 0.001211015 | 0.002105239 | | AFUB_027330 |
| **AFUB_061310** | 556.92479 | 2.379158088 | 0.115995226 | 20.51082759 | 1.72E-93 | 5.63E-92 | | AFUB_061310 |
| **AFUB_059560** | 19.25861775 | 2.379330686 | 0.46569445 | 5.109209884 | 3.24E-07 | 7.69E-07 | | AFUB_059560 |
| **AFUB_046780** | 221.4092947 | 2.381335592 | 0.155826046 | 15.28201258 | 1.01E-52 | 1.33E-51 | | AFUB_046780 |
| **AFUB_074690** | 7.440957421 | 2.382187049 | 0.754747948 | 3.156268333 | 0.001598018 | 0.002743363 | | AFUB_074690 |
| **AFUB_080320** | 3174.447645 | 2.384975835 | 0.096654612 | 24.6752409 | 1.97E-134 | 1.33E-132 | | AFUB_080320 |
| **AFUB_073190** | 9.857928506 | 2.389988792 | 0.65399401 | 3.654450585 | 0.000257733 | 0.000480929 | | AFUB_073190 |
| **AFUB_075710** | 6.438359028 | 2.391508291 | 0.800289448 | 2.988304164 | 0.002805302 | 0.004673564 | | AFUB_075710 |
| **AFUB_036270** | 66.86756226 | 2.393458506 | 0.257186498 | 9.306314785 | 1.32E-20 | 6.30E-20 | | AFUB_036270 |
| **AFUB_096370** | 208.0614606 | 2.397480119 | 0.162954466 | 14.7125769 | 5.35E-49 | 6.42E-48 | | AFUB_096370 |
| **AFUB_033210** | 216.2540943 | 2.39911815 | 0.168442943 | 14.24291279 | 4.96E-46 | 5.55E-45 | | AFUB_033210 |
| **AFUB_101559** | 4.389592962 | 2.399355489 | 0.974826656 | 2.461315017 | 0.013842876 | 0.021046064 | | AFUB_101559 |
| **AFUB_047450** | 8.625266713 | 2.401255665 | 0.696742356 | 3.446404033 | 0.0005681 | 0.001023757 | | AFUB_047450 |
| **AFUB_048010** | 280.3961731 | 2.403388492 | 0.143718043 | 16.72294196 | 8.92E-63 | 1.48E-61 | | AFUB_048010 |
| **AFUB_083500** | 333.9059112 | 2.40471677 | 0.138706958 | 17.3366701 | 2.49E-67 | 4.56E-66 | | AFUB_083500 |
| **AFUB_006510** | 36.01242284 | 2.405899644 | 0.350157216 | 6.870912652 | 6.38E-12 | 2.05E-11 | | AFUB_006510 |
| **AFUB_009180** | 333.779554 | 2.406360162 | 0.134887201 | 17.83979608 | 3.47E-71 | 6.99E-70 | | AFUB_009180 |
| **AFUB_063960** | 200.8333455 | 2.407928032 | 0.165534667 | 14.5463671 | 6.16E-48 | 7.18E-47 | | AFUB_063960 |
| **AFUB_100810** | 640.7175702 | 2.419399535 | 0.113121684 | 21.38758421 | 1.74E-101 | 6.64E-100 | | AFUB_100810 |
| **AFUB_039140** | 213.0521554 | 2.423718736 | 0.164348881 | 14.74740033 | 3.20E-49 | 3.86E-48 | | AFUB_039140 |
| **AFUB_029620** | 639.318465 | 2.427825626 | 0.106969767 | 22.69637195 | 4.87E-114 | 2.42E-112 | | AFUB_029620 |
| **AFUB_018570** | 23.05815417 | 2.434423637 | 0.431999307 | 5.635248937 | 1.75E-08 | 4.56E-08 | | AFUB_018570 |
| **AFUB_088150** | 412.6915422 | 2.43657493 | 0.124582274 | 19.55795842 | 3.53E-85 | 9.58E-84 | | AFUB_088150 |
| **AFUB_080090** | 13.45751718 | 2.441008987 | 0.568573375 | 4.293217186 | 1.76E-05 | 3.68E-05 | | AFUB_080090 |
| **AFUB_015650** | 137.48705 | 2.443452916 | 0.191182764 | 12.78071758 | 2.10E-37 | 1.81E-36 | | AFUB_015650 |
| **AFUB_087560** | 230.4005215 | 2.443709636 | 0.157867706 | 15.47947766 | 4.77E-54 | 6.48E-53 | | AFUB_087560 |
| **AFUB_101950** | 12.27292499 | 2.44882142 | 0.592978992 | 4.129693382 | 3.63E-05 | 7.36E-05 | | AFUB_101950 |
| **AFUB_056290** | 19.86285628 | 2.450503732 | 0.478525968 | 5.120942011 | 3.04E-07 | 7.25E-07 | | AFUB_056290 |
| **AFUB_002540** | 121.6470056 | 2.450644801 | 0.201680016 | 12.15115334 | 5.66E-34 | 4.38E-33 | | AFUB_002540 |

### Table S5

| | | | | | | | | | | | | | | | | | | | | | | | | | | | | | |
|---|---|---|---|---|---|---|---|---|---|---|---|---|---|---|---|---|---|---|---|---|---|---|---|---|---|---|---|---|---|
| **AFUB_079520** | 3.355496195 | 2.452082851 | 1.105942905 | 2.21718756 | 0.026610277 | 0.038686879 | | AFUB_079520 | Ortholog(s) have catalytic activity and role | Cytochrome P450 | Cytochrome P450;Cytochror | protein coding gene | | GO:0003824;G | obsolete oxidation-reduction pro | | catalytic activity;electron t | 1635 | 518 | | 355 | 2 | OG6_100895 | PF00067 | B0Y937 | IPR001128;IPR | 1.14.-.- (Acting | 0 | scf_000006_A_fumigatus_A1163:2,096,528..2,098,162(-) |
| **AFUB_052740** | 2571.620381 | 2.464407739 | 0.083066146 | 29.66801599 | 1.99E-193 | 3.09E-191 | | AFUB_052740 | Protein of unknown function | Mitochondrial substrate/solute | Mitochondrial substrate/solut | protein coding gene | | | | | | 2114 | 315 | | 351 | 0 | OG6_108609 | PF00153 | | IPR018108;IPR023395 | | 0 | scf_000004_A_fumigatus_A1163:1,054,759..1,056,872(-) |
| **AFUB_045590** | 14.5740585 | 2.46445507 | 0.540165779 | 4.562405036 | 5.06E-06 | 1.10E-05 | | AFUB_045590 | Protein of unknown function | N/A | N/A | protein coding gene | | | | | | 333 | 95 | | 2 | 0 | OG6_455348 | N/A | B0Y045 | N/A | | 0 | scf_000003_A_fumigatus_A1163:3,409,403..3,409,735(-) |
| **AFUB_009580** | 11.27364857 | 2.46529579 | 0.618096413 | 3.988529523 | 6.65E-05 | 0.000131521 | | AFUB_009580 | Protein of unknown function | Phytanoyl-CoA dioxygenase | Phytanoyl-CoA dioxygenase | protein coding gene | | GO:0051213 | | | dioxygenase activity | 1104 | 337 | | 151 | 0 | OG6_119779 | PF05721 | B0XQD6 | IPR008775 | | 0 | scf_000001_A_fumigatus_A1163:2,691,463..2,692,566(-) |
| **AFUB_091840** | 4.533081378 | 2.465465225 | 0.963648607 | 2.558469142 | 0.010513415 | 0.01622214 | | AFUB_091840 | Has domain(s) with predicted protein bindi | Nucleoside phosphorylase dom | Nucleoside phosphorylase d | protein coding gene | | GO:0003824;G | nucleoside metabolic process | | catalytic activity;protein bi | 4204 | 1257 | | 1183 | 2 | OG6_111122 | PF01048;PF057 | B0YB77 | IPR000845;IPR002110;IPR0071 | | 0 | scf_000008_A_fumigatus_A1163:1,543,103..1,547,306(+) |
| **AFUB_092720** | 21.54322565 | 2.465481315 | 0.454416102 | 5.425602885 | 5.78E-08 | 1.45E-07 | | AFUB_092720 | Has domain(s) with predicted DNA binding | Transcription factor, MADS-box | Transcription factor, MADS-b | protein coding gene | GO:0005634 | GO:0003677;GO:0046983 | | nucleus | DNA binding;protein dime | 464 | 103 | | 117 | 1 | OG6_166547 | PF00319 | B0YBG4 | IPR002100;IPR036879 | | 0 | scf_000008_A_fumigatus_A1163:1,797,469..1,797,932(+) |
| **AFUB_049910** | 3.366088541 | 2.467197964 | 1.131146766 | 2.18114752 | 0.029172507 | 0.042106992 | | AFUB_049910 | Has domain(s) with predicted metallopepti | Lysine-specific metallo-endope | Lysine-specific metallo-endo | protein coding gene | | GO:0046872;GO:0004222;GO:0008237 | | | metal ion binding;metalloe | 1345 | 331 | | 10 | 0 | OG6r21_10601 | PF14521 | B0Y1N3 | IPR029463 | | 0 | scf_000004_A_fumigatus_A1163:286,929..288,273(+) |
| **AFUB_049790** | 48.54942997 | 2.468177957 | 0.30569617 | 8.073957737 | 6.81E-16 | 2.65E-15 | | AFUB_049790 | Ortholog(s) have intracellular localization | N/A | Quinone oxidoreductase/zeta | protein coding gene | GO:0005622 | GO:0016491;G | obsolete oxidati | intracellular ana | oxidoreductase activity;zi | 1374 | 330 | | 1199 | 4 | OG6_100491 | PF13602 | B0Y1M1 | IPR002364;IPR | 1.1.1.1 (Alcohol | 0 | scf_000004_A_fumigatus_A1163:250,729..252,102(+) |
| **AFUB_047630** | 8.956025403 | 2.468616816 | 0.688388958 | 3.586078463 | 0.000335688 | 0.000617661 | | AFUB_047630 | Has domain(s) with predicted transmembr | N/A | N/A | protein coding gene | GO:0016021;G | GO:0004888 | cell surface rece | integral compon | transmembrane signaling | 1260 | 355 | | 119 | 0 | OG6_119585 | PF05462 | B0XX39 | N/A | | 7 | scf_000003_A_fumigatus_A1163:3,941,997..3,943,256(-) |
| **AFUB_035560** | 5.639055871 | 2.46896988 | 0.862722728 | 2.861834746 | 0.004211965 | 0.006854668 | | AFUB_035560 | Protein of unknown function | N/A | N/A | protein coding gene | | | | | | 866 | 265 | | 296 | 3 | OG6_152993 | N/A | B0Y0C9 | N/A | | 0 | scf_000003_A_fumigatus_A1163:461,528..462,393(-) |
| **AFUB_047390** | 5.639055871 | 2.46896988 | 0.862722728 | 2.861834746 | 0.004211965 | 0.006854668 | | AFUB_047390 | Has domain(s) with predicted oxidoreduct | TauD/TfdA-like domain | TauD/TfdA-like domain | protein coding gene | | GO:0051213;G | obsolete oxidation-reduction pro | | dioxygenase activity;oxid | 1281 | 388 | | 534 | 2 | OG6_105576 | PF02668 | B0XX16 | IPR003819 | 1.14.11.17 (Tau | 0 | scf_000003_A_fumigatus_A1163:3,879,655..3,880,935(-) |
| **AFUB_057570** | 20.29541288 | 2.47011957 | 0.457748296 | 5.396239789 | 6.81E-08 | 1.70E-07 | | AFUB_057570 | Protein of unknown function | N/A | N/A | protein coding gene | | | | | | 1089 | 301 | MRPLSILLVVS | 36 | 0 | OG6_455196 | N/A | B0Y4E8 | N/A | | 0 | scf_000004_A_fumigatus_A1163:2,469,417..2,470,505(-) |
| **AFUB_029980** | 243.6452486 | 2.470877974 | 0.156578398 | 15.78045247 | 4.24E-56 | 6.05E-55 | | AFUB_029980 | Ortholog(s) have glucan endo-1,3-beta-glu | Glycoside hydrolase family 16 | Glycoside hydrolase family 1 | protein coding gene | GO:0031225;G | GO:0052861;G | carbohydrate m | anchored comp | glucan endo-1,3-beta-glu | 2519 | 652 | MAPSSLLLSVG | 898 | 3 | OG6_105686 | PF00722 | B0XTU6 | IPR000757;IPR | 3.2.1.- (Glycosid | 0 | scf_000002_A_fumigatus_A1163:3,733,363..3,735,881(+) |
| **AFUB_097810** | 520.5896311 | 2.472527642 | 0.121248488 | 20.39223493 | 1.96E-92 | 6.25E-91 | | AFUB_097810 | Has domain(s) with predicted sequence-s | Zn(2)-C6 fungal-type DNA-binc | Zn(2)-C6 fungal-type DNA-b | protein coding gene | GO:0042025;G | GO:0000981;G | regulation of tra | host cell nucleus | DNA-binding transcription | 2188 | 485 | | 694 | 5 | OG6_101136 | PF00172 | | IPR001138;IPR036864 | | 0 | scf_000009_A_fumigatus_A1163:1,470,617..1,472,804(+) |
| **AFUB_080630** | 32.97697706 | 2.476222271 | 0.368901146 | 6.712427694 | 1.91E-11 | 5.98E-11 | | AFUB_080630 | Ortholog(s) have extracellular region local | N/A | N/A | protein coding gene | GO:0005576 | | | extracellular region | | 836 | 272 | MRFLPWAVAIL | 72 | 0 | OG6_165847 | PF17660 | B0Y9D3 | N/A | | 0 | scf_000007_A_fumigatus_A1163:264,107..264,942(-) |
| **AFUB_002320** | 183.4942063 | 2.476870749 | 0.174754841 | 14.17340281 | 1.34E-45 | 1.48E-44 | | AFUB_002320 | Protein of unknown function | Protein of unknown function DU | Protein of unknown function | protein coding gene | | | | | | 1356 | 433 | | 91 | 0 | OG6_186579 | PF11905 | B0XRF8 | IPR021833 | | 0 | scf_000001_A_fumigatus_A1163:645,667..647,022(+) |
| **AFUB_092500** | 7.89776131 | 2.477024417 | 0.745901832 | 3.320845066 | 0.000897453 | 0.001584147 | | AFUB_092500 | Protein of unknown function | Glyoxalase/fosfomycin resistan | Glyoxalase/fosfomycin resist | protein coding gene | | | | | | 1086 | 361 | | 67 | 0 | OG6_453853 | PF00903 | B0YBE3 | IPR004360;IPR | 1.13.11.- (With i | 0 | scf_000008_A_fumigatus_A1163:1,691,333..1,692,418(-) |
| **AFUB_026530** | 6457.429431 | 2.477392555 | 0.084683783 | 29.25462788 | 3.92E-188 | 5.72E-186 | | AFUB_026530 | Has domain(s) with predicted nucleic acid | DEAD/DEAH box helicase dom | ATP-dependent RNA helicas | protein coding gene | | GO:0005524;GO:0004386;GO:0003676 | | | ATP binding;helicase acti | 3198 | 549 | | 322 | 0 | OG6_100364 | PF00270;PF00271 | | IPR000629;IPR | 3.4.24.- (Metallo | 0 | scf_000002_A_fumigatus_A1163:2,729,493..2,732,690(+) |
| **AFUB_050110** | 177.0458066 | 2.478468505 | 0.175213867 | 14.14539015 | 1.99E-45 | 2.19E-44 | | AFUB_050110 | Protein of unknown function | N/A | N/A | protein coding gene | | | | | | 1014 | 337 | | 70 | 0 | OG6_137031 | N/A | B0Y1W9 | N/A | | 0 | scf_000004_A_fumigatus_A1163:335,299..336,312(-) |
| **AFUB_071790** | 4.536902824 | 2.47885818 | 0.970460961 | 2.554310043 | 0.010639847 | 0.01639622 | | AFUB_071790 | Ortholog(s) have transcription coactivator | N/A | Winged helix DNA-binding d | protein coding gene | | GO:0003713 | aflatoxin biosynthetic process;m | | transcription coactivator a | 1366 | 437 | | 631 | 1 | OG6_112909 | N/A | B0Y590 | IPR036390 | | 0 | scf_000005_A_fumigatus_A1163:2,535,673..2,537,038(-) |
| **AFUB_098970** | 28.4279523 | 2.485420794 | 0.397014585 | 6.260275788 | 3.84E-10 | 1.10E-09 | | AFUB_098970 | Ortholog(s) have holo-[acyl-carrier-protein | 4'-phosphopantetheinyl transfe | Holo-[acyl carrier protein] sy | protein coding gene | GO:0005739 | GO:0008897;G | fatty acid biosyn | mitochondrion | holo-[acyl-carrier-protein] | 1268 | 153 | | 93 | 0 | OG6_454838 | PF01648 | B0YEF2 | IPR002582;IPR | 2.7.8.7 (Holo-[ac | 0 | scf_000010_A_fumigatus_A1163:237,055..238,322(+) |
| **AFUB_064750** | 677.6843537 | 2.486861051 | 0.131382164 | 18.92845245 | 6.65E-80 | 1.57E-78 | | AFUB_064750 | Has domain(s) with predicted nucleic acid | DEAD/DEAH box helicase dom | ATP-dependent RNA helicas | protein coding gene | | GO:0005524;GO:0003724;GO:0004386;GO:000 | | | ATP binding;RNA helicas | 4209 | 674 | | 323 | 0 | OG6_100461 | PF00270;PF002 | B0Y5V9 | IPR000629;IPR | 3.6.1.- (In phosp | 0 | scf_000005_A_fumigatus_A1163:539,042..543,250(+) |
| **AFUB_101590** | 6.807669522 | 2.48715655 | 0.793601992 | 3.134009961 | 0.00172435 | 0.002949208 | | AFUB_101590 | Protein of unknown function | Metallo-beta-lactamase | Metallo-beta-lactamase;Ribo | protein coding gene | | GO:0016787 | | | hydrolase activity | 846 | 281 | | 173 | 2 | OG6_119662 | PF12706 | B0YF52 | IPR001279;IPR036866 | | 0 | scf_000011_A_fumigatus_A1163:318,824..319,669(+) |
| **AFUB_051330** | 469.7401854 | 2.487600493 | 0.122159893 | 20.36347958 | 3.53E-92 | 1.11E-90 | | AFUB_051330 | Has domain(s) with predicted amino acid t | Amino acid/polyamine transpor | Amino acid/polyamine transp | protein coding gene | GO:0016021;G | GO:0015171;G | amino acid trans | integral compon | amino acid transmembrar | 2981 | 554 | | 163 | 0 | OG6_137061 | PF13520 | B0Y2G1 | IPR002293 | 2.1.1.314 (Dipht | 11 | scf_000004_A_fumigatus_A1163:673,257..676,237(+) |
| **AFUB_020760** | 152.6044932 | 2.488014724 | 0.18735929 | 13.27937744 | 3.05E-40 | 2.84E-39 | | AFUB_020760 | Protein of unknown function | N/A | N/A | protein coding gene | | | | | | 1453 | 371 | MQALRILSMVV | 269 | 0 | OG6_119610 | N/A | | N/A | | 1 | scf_000002_A_fumigatus_A1163:970,297..971,749(-) |
| **AFUB_039990** | 53.92598576 | 2.489099413 | 0.290240062 | 8.576002212 | 9.82E-18 | 4.17E-17 | | AFUB_039990 | Has domain(s) with predicted hydrolase a | Glycoside hydrolase, family 43 | Glycoside hydrolase, family | protein coding gene | | GO:0046558;G | arabinan catabolic process;carb | | arabinan endo-1,5-alpha- | 1605 | 415 | MFRIQLLLQAC | 433 | 4 | OG6_110274 | PF04616 | B0XY82 | IPR006710;IPR | 3.2.1.8 (Endo-1, | 0 | scf_000003_A_fumigatus_A1163:1,724,169..1,725,773(-) |
| **AFUB_018260** | 72.61409225 | 2.49263883 | 0.265154369 | 9.400708125 | 5.42E-21 | 2.62E-20 | | AFUB_018260 | Has domain(s) with predicted flavin adenir | Glucose-methanol-choline oxid | Glucose-methanol-choline o | protein coding gene | | GO:0030246;G | carbohydrate catabolic process; | | carbohydrate binding;cata | 2841 | 820 | MRSFLFGGLV | 205 | 0 | OG6_186654 | PF00732;PF051 | B0XT73 | IPR000172;IPR | 1.1.99.18 (Cello | 0 | scf_000002_A_fumigatus_A1163:267,194..270,034(+) |
| **AFUB_078210** | 6.840111822 | 2.50089796 | 0.793350145 | 3.152325584 | 0.001619756 | 0.002777712 | xlnB | AFUB_078210 | Ortholog(s) have endo-1,4-beta-xylanase | Glycosyl hydrolases family 11 ( | Glycoside hydrolase family 1 | protein coding gene | GO:0005576 | GO:0031176;G | carbohydrate m | extracellular reg | endo-1,4-beta-xylanase a | 712 | 221 | MVSFSSLVLAA | 336 | 2 | OG6_115937 | PF00457 | B0Y8Q8 | IPR001137;IPR | 3.2.1.4 (Cellulas | 0 | scf_000006_A_fumigatus_A1163:1,757,336..1,758,047(+) |
| **AFUB_046980** | 32.2808429 | 2.502018491 | 0.371374589 | 6.73718279 | 1.61E-11 | 5.08E-11 | | AFUB_046980 | Protein of unknown function | Serine hydrolase FSH | Serine hydrolase FSH;Alpha | protein coding gene | | | | | | 776 | 225 | | 640 | 1 | OG6_102370 | PF03959 | B0XWQ8 | IPR005645;IPR029058 | | 0 | scf_000003_A_fumigatus_A1163:3,755,108..3,755,883(+) |
| **AFUB_000920** | 39.28199382 | 2.505164953 | 0.33992355 | 7.369789336 | 1.71E-13 | 5.97E-13 | | AFUB_000920 | Protein of unknown function | N/A | SKP1/BTB/POZ domain sup | protein coding gene | | | | | | 1186 | 360 | | 653 | 7 | OG6_121403 | N/A | B0XM49 | IPR011333 | | 0 | scf_000001_A_fumigatus_A1163:237,652..238,837(-) |
| **AFUB_021820** | 316.8312299 | 2.505783078 | 0.141562858 | 17.70085112 | 4.13E-70 | 8.11E-69 | | AFUB_021820 | Has domain(s) with predicted protein bindi | NUC153 | WD40 repeat;NUC153;WD4 | protein coding gene | GO:0005730;G | GO:0005515 | | nucleolus;nucle | protein binding | 2234 | 721 | | 307 | 0 | OG6_102419 | PF08159 | B0XV64 | IPR001680;IPR012580;IPR0363 | | 0 | scf_000002_A_fumigatus_A1163:1,303,136..1,305,369(-) |
| **AFUB_048890** | 4.655385101 | 2.506454799 | 0.979010616 | 2.560191643 | 0.010461446 | 0.016147123 | | AFUB_048890 | Has domain(s) with predicted sequence-s | Zn(2)-C6 fungal-type DNA-binc | Zn(2)-C6 fungal-type DNA-b | protein coding gene | GO:0042025;G | GO:0003677;G | regulation of tra | host cell nucleus | DNA binding;DNA-binding | 1862 | 526 | | 122 | 0 | OG6_454177 | PF00172 | B0Y170 | IPR001138;IPR036864 | | 0 | scf_000004_A_fumigatus_A1163:50,292..52,153(+) |
| **AFUB_090250** | 675.5527218 | 2.507123758 | 0.108460162 | 23.11561871 | 3.23E-118 | 1.71E-116 | | AFUB_090250 | Has domain(s) with predicted protein bindi | Zinc finger, NF-X1-type;R3H d | Zinc finger, NF-X1-type;R3H | protein coding gene | GO:0005634 | GO:0003700;G | regulation of tra | nucleus | DNA-binding transcription | 3438 | 1145 | | 260 | 0 | OG6_103027 | PF01422;PF014 | B0YCH9 | IPR000967;IPR | 2.3.2.- (Aminoac | 0 | scf_000008_A_fumigatus_A1163:1,078,477..1,081,914(+) |
| **AFUB_068100** | 38.344401 | 2.509227187 | 0.341320104 | 7.351536444 | 1.96E-13 | 6.82E-13 | | AFUB_068100 | Has domain(s) with predicted role in trans | Major facilitator superfamily | Major facilitator superfamily; | protein coding gene | GO:0016021;G | GO:0022857 | transmembrane | integral compon | transmembrane transport | 1819 | 574 | | 1751 | 6 | OG6_104355 | PF07690 | B0Y6T8 | IPR011701;IPR | 2.7.-.- (Transfer | 12 | scf_000005_A_fumigatus_A1163:1,440,619..1,442,437(-) |
| **AFUB_016310** | 10.5005434 | 2.515701744 | 0.647194668 | 3.887086636 | 0.000101455 | 0.000197299 | xlnD | AFUB_016310 | Ortholog(s) have xylosyltransferase, alpha | Glycoside hydrolase, family 3, | Glycoside hydrolase, family | protein coding gene | GO:0005576;G | GO:0046556;G | carbohydrate m | extracellular reg | alpha-L-arabinofuranosid | 2379 | 792 | MSVAKSIAAVL | 2041 | 9 | OG6_100201 | PF00933;PF01 | B0XP71 | IPR001764;IPR | 3.2.1.21 (Beta-g | 0 | scf_000001_A_fumigatus_A1163:4,632,488..4,634,866(-) |
| **AFUB_034030** | 5.806701965 | 2.517849302 | 0.859356844 | 2.929922905 | 0.003390461 | 0.005586608 | | AFUB_034030 | Has domain(s) with predicted molybdenun | Molybdopterin oxidoreductase; | Molybdopterin oxidoreductas | protein coding gene | | GO:0046872;G | nitrate assimilation;obsolete oxid | | metal ion binding;molybde | 2792 | 909 | | 63 | 0 | OG6_106344 | PF00384;PF01 | B0XZI9 | IPR006656;IPR | 1.17.99.7 (Form | 0 | scf_000003_A_fumigatus_A1163:70,530..73,321(-) |
| **AFUB_082860** | 87.02972979 | 2.520031896 | 0.233299613 | 10.8016977 | 3.38E-27 | 2.07E-26 | | AFUB_082860 | Has domain(s) with predicted catalytic acti | RNA 3'-terminal phosphate cyc | RNA 3'-terminal phosphate c | protein coding gene | GO:0005730 | GO:0003824 | RNA processing | nucleolus | catalytic activity | 1798 | 403 | | 293 | 0 | OG6_102522 | PF01137;PF051 | B0YA00 | IPR000228;IPR | 6.5.1.4 (RNA 3'- | 0 | scf_000007_A_fumigatus_A1163:872,192..873,989(+) |
| **AFUB_047890** | 17.26257583 | 2.52107314 | 0.51293246 | 4.915019693 | 8.88E-07 | 2.04E-06 | | AFUB_047890 | Protein of unknown function | N/A | N/A | protein coding gene | | | | | | 512 | 153 | | 52 | 1 | OG6_186711 | N/A | B0XX64 | N/A | | 0 | scf_000003_A_fumigatus_A1163:4,010,219..4,010,730(+) |
| **AFUB_097970** | 240.1565354 | 2.521699663 | 0.154239848 | 16.34921 | 4.41E-60 | 6.84E-59 | | AFUB_097970 | Has domain(s) with predicted hydrolase a | GNAT domain;Glycoside hydro | GNAT domain;Glycoside hy | protein coding gene | | GO:0008080;G | carbohydrate metabolic process; | | N-acetyltransferase activi | 2229 | 742 | | 305 | 2 | OG6_106267 | PF00583;PF00 | B0YE62 | IPR000182;IPR001764;IPR0161 | | 0 | scf_000009_A_fumigatus_A1163:1,533,016..1,535,244(+) |
| **AFUB_088380** | 95.17359272 | 2.523287857 | 0.239870936 | 10.51935636 | 7.04E-26 | 4.10E-25 | | AFUB_088380 | Has domain(s) with predicted sequence-s | Zn(2)-C6 fungal-type DNA-binc | Zn(2)-C6 fungal-type DNA-b | protein coding gene | GO:0042025;G | GO:0003677;G | regulation of tra | host cell nucleus | DNA binding;DNA-binding | 2595 | 864 | | 387 | 5 | OG6_133375 | PF00172;PF04 | B0YBZ7 | IPR001138;IPR | 1.1.1.85 (3-isop | 0 | scf_000008_A_fumigatus_A1163:480,667..483,261(-) |
| **AFUB_092270** | 46.67848932 | 2.524489589 | 0.31377465 | 8.045549857 | 8.59E-16 | 3.32E-15 | | AFUB_092270 | Protein of unknown function | N/A | FAD/NAD(P)-binding domair | protein coding gene | | GO:0004497 | | | monooxygenase activity | 2405 | 599 | | 661 | 4 | OG6_104745 | PF13738 | B0YBC0 | IPR036188 | 1.14.13.8 (Flavi | 0 | scf_000008_A_fumigatus_A1163:1,632,639..1,635,043(+) |
| **AFUB_073750** | 99.05409029 | 2.526552565 | 0.225354233 | 11.21147151 | 3.58E-29 | 2.35E-28 | | AFUB_073750 | Protein of unknown function | N/A | N/A | protein coding gene | | | | | | 387 | 101 | | 1 | 0 | OG6_454200 | N/A | B0Y7G7 | N/A | | 0 | scf_000006_A_fumigatus_A1163:437,406..437,792(+) |
| **AFUB_000280** | 82.12041591 | 2.530408862 | 0.240715514 | 10.51203064 | 7.60E-26 | 4.43E-25 | bglH | AFUB_000280 | Has domain(s) with predicted hydrolase a | Glycoside hydrolase, family 3, | Glycoside hydrolase, family | protein coding gene | GO:0005576 | GO:0008422;G | carbohydrate m | extracellular reg | beta-glucosidase activity; | 2825 | 829 | | 743 | 4 | OG6_110843 | PF00933;PF01 | B0XM94 | IPR001764;IPR | 3.2.1.21 (Beta-g | 0 | scf_000001_A_fumigatus_A1163:82,059..84,883(-) |
| **AFUB_006050** | 19.9860328 | 2.535295276 | 0.467555104 | 5.422452359 | 5.88E-08 | 1.48E-07 | | AFUB_006050 | Protein of unknown function | N/A | N/A | protein coding gene | | | | | | 2965 | 344 | | 186 | 1 | OG6_152840 | N/A | B0XP61 | N/A | | 0 | scf_000001_A_fumigatus_A1163:1,712,959..1,715,923(+) |
| **AFUB_039440** | 21.300451 | 2.535799689 | 0.462134979 | 5.487140777 | 4.08E-08 | 1.04E-07 | | AFUB_039440 | Protein of unknown function | N/A | N/A | protein coding gene | | | | | | 2658 | 586 | | 13 | 0 | OG6_454461 | N/A | | N/A | | 0 | scf_000003_A_fumigatus_A1163:1,578,223..1,580,880(+) |
| **AFUB_088740** | 522.2050068 | 2.537538125 | 0.123839248 | 20.4905808 | 2.61E-93 | 8.48E-92 | | AFUB_088740 | Has domain(s) with predicted nucleic acid | Zinc finger, CCHC-type | Zinc finger, CCHC-type;Zinc | protein coding gene | | GO:0003676;GO:0008270 | | | nucleic acid binding;zinc i | 1741 | 243 | | 430 | 1 | OG6_101988 | PF00098 | | IPR001878;IPR036875 | | 0 | scf_000008_A_fumigatus_A1163:585,541..587,281(+) |
| **AFUB_073670** | 22.71255274 | 2.539207579 | 0.462759626 | 5.487098351 | 4.09E-08 | 1.04E-07 | | AFUB_073670 | Has domain(s) with predicted role in trans | Mitochondrial substrate/solute | Mitochondrial carrier protein; | protein coding gene | GO:0016021;G | GO:0005347;G | transmembrane | integral compon | ATP transmembrane tran | 1869 | 316 | | 237 | 0 | OG6_101430 | PF00153 | B0Y7F9 | IPR002067;IPR | 2.7.-.- (Transfer | 0 | scf_000006_A_fumigatus_A1163:407,681..409,549(-) |
| **AFUB_073090** | 52.0243639 | 2.544248015 | 0.303229067 | 8.39051493 | 4.84E-17 | 1.99E-16 | | AFUB_073090 | Has domain(s) with predicted methyltransf | Methyltransferase type 12 | Methyltransferase type 12;M | protein coding gene | | GO:0008168 | methylation | | methyltransferase activity | 1101 | 366 | | 295 | 0 | OG6_103169 | PF08242 | B0Y7A1 | IPR013217;IPR | 2.1.1.268 (tRNA | 0 | scf_000006_A_fumigatus_A1163:261,980..263,080(+) |
| **AFUB_080930** | 43.7107695 | 2.544712886 | 0.321902797 | 7.905221415 | 2.67E-15 | 1.01E-14 | | AFUB_080930 | Has domain(s) with predicted nucleotide b | N/A | Short-chain dehydrogenase/ | protein coding gene | | | | | | 1260 | 365 | | 76 | 0 | OG6_455153 | PF13561 | B0Y9G3 | IPR002347;IPR036291 | | 0 | scf_000007_A_fumigatus_A1163:345,292..346,551(-) |
| **AFUB_040390** | 267.3226323 | 2.546342943 | 0.156053498 | 16.3171548 | 7.46E-60 | 1.15E-58 | | AFUB_040390 | Has domain(s) with predicted protein kina | Protein kinase domain | Protein kinase domain;Serin | protein coding gene | | GO:0005524;G | protein phosphorylation | | ATP binding;protein kinas | 2368 | 413 | | 116 | 0 | OG6_186745 | PF00069 | B0XYC2 | IPR000719;IPR008271;IPR0110 | | 0 | scf_000003_A_fumigatus_A1163:1,820,411..1,822,778(-) |
| **AFUB_009560** | 38.93824588 | 2.546469774 | 0.341709895 | 7.452139401 | 9.18E-14 | 3.26E-13 | | AFUB_009560 | Protein of unknown function | N/A | S-adenosyl-L-methionine-de | protein coding gene | | | | | | 1040 | 227 | | 192 | 0 | OG6_106858 | N/A | | IPR029063 | | 0 | scf_000001_A_fumigatus_A1163:2,686,790..2,687,829(-) |
| **AFUB_091820** | 35.25122829 | 2.546984218 | 0.364057195 | 6.996110106 | 2.63E-12 | 8.63E-12 | | AFUB_091820 | Has domain(s) with predicted phosphoric | N/A | HD/PDEase domain;Cyanan | protein coding gene | | GO:0046872;GO:0008081 | | | metal ion binding;phosph | 780 | 259 | | 300 | 1 | OG6_129919 | N/A | B0YB75 | IPR003607;IPR | 4.2.1.69 (Cyana | 0 | scf_000008_A_fumigatus_A1163:1,536,686..1,537,465(+) |
| **AFUB_037040** | 173.9129422 | 2.548081428 | 0.173794529 | 14.66145941 | 1.14E-48 | 1.36E-47 | | AFUB_037040 | Protein of unknown function | N/A | N/A | protein coding gene | | | | | | 1023 | 340 | | 157 | 0 | OG6_166366 | N/A | B0XX71 | N/A | | 0 | scf_000003_A_fumigatus_A1163:875,845..876,867(-) |
| **AFUB_021260** | 174.6267512 | 2.550070794 | 0.174564535 | 14.60818371 | 2.49E-48 | 2.93E-47 | | AFUB_021260 | Protein of unknown function | N/A | N/A | protein coding gene | | | | | | 3371 | 705 | | 251 | 0 | OG6_105273 | N/A | B0XV10 | N/A | | 0 | scf_000002_A_fumigatus_A1163:1,150,804..1,154,174(+) |
| **AFUB_075550** | 48.76991292 | 2.550427157 | 0.305009731 | 8.361789469 | 6.18E-17 | 2.52E-16 | | AFUB_075550 | Has domain(s) with predicted unfolded pro | DnaJ domain | DnaJ domain;DnaJ domain, | protein coding gene | | | | | | 794 | 241 | | 22 | 0 | OG6_454596 | PF00226 | B0Y7Z4 | IPR001623;IPR018253;IPR0368 | | 0 | scf_000006_A_fumigatus_A1163:1,014,561..1,015,354(-) |
| **AFUB_089320** | 100.4511369 | 2.552282411 | 0.225105348 | 11.33816871 | 8.49E-30 | 5.71E-29 | | AFUB_089320 | Has domain(s) with predicted nucleic acid | RNA recognition motif domain; | RNA recognition motif doma | protein coding gene | | GO:0003723;GO:0003676;GO:0000166 | | | RNA binding;nucleic acid | 3674 | 672 | | 177 | 0 | OG6_143880 | PF00076;PF04146 | | IPR000504;IPR007275;IPR0359 | | 0 | scf_000008_A_fumigatus_A1163:817,508..821,181(-) |
| **AFUB_051580** | 5.828703821 | 2.554293999 | 0.869062509 | 2.939137256 | 0.003291273 | 0.005434322 | | AFUB_051580 | Has domain(s) with predicted protein bindi | WD40 repeat | WD40 repeat;WD40-repeat- | protein coding gene | | GO:0005515 | | | protein binding | 1278 | 407 | | 273 | 0 | OG6_103232 | PF00400 | B0Y2Q4 | IPR001680;IPR036322 | | 0 | scf_000004_A_fumigatus_A1163:763,540..764,817(-) |
| **AFUB_099590** | 356.3792215 | 2.555541557 | 0.134181547 | 19.04540238 | 7.17E-81 | 1.76E-79 | | AFUB_099590 | Ortholog(s) have nucleus localization | Myc-type, basic helix-loop-heli | Myc-type, basic helix-loop-h | protein coding gene | GO:0005634 | GO:0046983 | cellular respons | nucleus | protein dimerization activi | 2470 | 299 | | 58 | 1 | OG6_186636 | PF00010 | B0YEL4 | IPR011598;IPR036638 | | 0 | scf_000010_A_fumigatus_A1163:386,321..388,790(-) |
| **AFUB_075500** | 690.1302079 | 2.560077474 | 0.109381987 | 23.40492752 | 3.81E-121 | 2.13E-119 | | AFUB_075500 | Ortholog(s) have oxidoreductase activity | Cytochrome P450 | Cytochrome P450;Cytochror | protein coding gene | GO:0016021 | GO:0009055;G | obsolete oxidati | integral compon | electron transfer activity;h | 1829 | 522 | | 545 | 0 | OG6_106077 | PF00067 | B0Y7Y9 | IPR001128;IPR | 1.14.-.- (Acting | 1 | scf_000006_A_fumigatus_A1163:993,095..994,923(+) |
| **AFUB_064180** | 41.94051019 | 2.564744359 | 0.334436171 | 7.668860562 | 1.74E-14 | 6.36E-14 | | AFUB_064180 | Has domain(s) with predicted nucleic acid | N/A | Zinc finger C2H2-type;Zinc fi | protein coding gene | | | | | | 3648 | 710 | | 189 | 0 | OG6_143760 | N/A | | IPR013087;IPR036236 | | 0 | scf_000005_A_fumigatus_A1163:389,238..392,885(+) |
| **AFUB_090430** | 166.3714154 | 2.565407949 | 0.181870579 | 14.10566473 | 3.50E-45 | 3.81E-44 | | AFUB_090430 | Has domain(s) with predicted transferase | UDP-glucuronosyl/UDP-glucos | UDP-glucuronosyl/UDP-gluc | protein coding gene | | GO:0008194;G | carbohydrate metabolic process; | | UDP-glycosyltransferase | 2553 | 787 | | 1149 | 2 | OG6_102789 | PF00201;PF03 | B0YCJ7 | IPR002213;IPR | 2.4.1.141 (N-ace | 0 | scf_000008_A_fumigatus_A1163:1,144,925..1,147,477(+) |
| **AFUB_017020** | 4.821597444 | 2.565940055 | 0.945108279 | 2.714969397 | 0.00662819 | 0.010523848 | | AFUB_017020 | Has domain(s) with predicted sequence-s | Zinc finger C2H2-type;Zn(2)-C | Zn(2)-C6 fungal-type DNA-b | protein coding gene | GO:0042025;G | GO:0003677;G | regulation of tra | host cell nucleus | DNA binding;DNA-binding | 2719 | 866 | | 417 | 1 | OG6_111246 | PF00096;PF001 | B0XPK8 | IPR001138;IPR007219;IPR0130 | | 0 | scf_000001_A_fumigatus_A1163:4,850,511..4,853,229(+) |
| **AFUB_092410** | 109.3563307 | 2.568047348 | 0.216925643 | 11.8383761 | 2.47E-32 | 1.82E-31 | | AFUB_092410 | Has domain(s) with predicted role in trans | Major facilitator superfamily | Major facilitator superfamily; | protein coding gene | GO:0016021;G | GO:0022857 | transmembrane | integral compon | transmembrane transport | 1230 | 409 | | 700 | 4 | OG6_108411 | PF07690 | B0YBD4 | IPR011701;IPR036259 | | 11 | scf_000008_A_fumigatus_A1163:1,667,522..1,668,751(-) |
| **AFUB_048320** | 24.07935087 | 2.568642308 | 0.438619576 | 5.856196237 | 4.74E-09 | 1.28E-08 | | AFUB_048320 | Protein of unknown function | Amino acid transporter, transm | Amino acid transporter, trans | protein coding gene | GO:0016021 | | | integral component of membrane | | 1649 | 453 | | 410 | 0 | OG6_122907 | PF01490 | B0XXH2 | IPR013057 | | 11 | scf_000003_A_fumigatus_A1163:4,113,290..4,114,938(-) |
| **AFUB_049800** | 76.63741367 | 2.568955253 | 0.25326342 | 10.14341216 | 3.54E-24 | 1.94E-23 | | AFUB_049800 | Protein of unknown function | N/A | N/A | protein coding gene | GO:0016021 | | | integral component of membrane | | 1794 | 597 | MAYINFYLLVLL | 9 | 0 | OG6_186848 | N/A | B0Y1M2 | N/A | | 9 | scf_000004_A_fumigatus_A1163:252,362..254,155(-) |
| **AFUB_036570** | 3.499064581 | 2.569898079 | 1.127574267 | 2.279138637 | 0.022658825 | 0.033359884 | | AFUB_036570 | Protein of unknown function | N/A | N/A | protein coding gene | | | | | | 1301 | 225 | | 17 | 0 | OG6_453942 | N/A | B0XWV7 | N/A | | 0 | scf_000003_A_fumigatus_A1163:725,547..726,847(+) |
| **AFUB_058280** | 1543.931759 | 2.572658791 | 0.096735873 | 26.59467185 | 7.82E-156 | 7.46E-154 | | AFUB_058280 | Protein of unknown function | Pentatricopeptide repeat;Penta | Pentatricopeptide repeat | protein coding gene | | GO:0005515 | | | protein binding | 3819 | 1272 | | 462 | 0 | OG6_100693 | PF13041;PF13 | B0Y0R6 | IPR002885 | | 0 | scf_000004_A_fumigatus_A1163:2,622,789..2,626,607(-) |
| **AFUB_068950** | 494.0441453 | 2.574346215 | 0.132560484 | 19.42016308 | 5.21E-84 | 1.38E-82 | nar1 | AFUB_068950 | Protein of unknown function | Iron hydrogenase, large subun | Iron hydrogenase, large sub | protein coding gene | | GO:0051539;GO:0046872 | | | 4 iron, 4 sulfur cluster bin | 2333 | 597 | | 305 | 0 | OG6_101268 | PF02906 | B0Y4F9 | IPR004108;IPR | 1.12.7.2 (Ferred | 0 | scf_000005_A_fumigatus_A1163:1,710,286..1,712,618(+) |
| **AFUB_058510** | 572.9345132 | 2.574393031 | 0.113258359 | 22.73026957 | 2.25E-114 | 1.13E-112 | | AFUB_058510 | Ortholog(s) have extracellular region local | Polysaccharide lyase | Polysaccharide lyase | protein coding gene | GO:0005576 | | | extracellular region | | 1429 | 292 | | 53 | 0 | OG6_153201 | PF14099 | B0Y0T9 | IPR025975 | | 0 | scf_000004_A_fumigatus_A1163:2,697,090..2,698,518(+) |
| **AFUB_076620** | 65.16070429 | 2.575585521 | 0.270411566 | 9.524686971 | 1.66E-21 | 8.16E-21 | | AFUB_076620 | Protein of unknown function | Extracellular membrane proteir | Extracellular membrane prot | protein coding gene | GO:0031225;GO:0005618;GO:0005886 | | | anchored component of membrane;cell wa | | 795 | 202 | MHFSRTSLILF | 41 | 0 | OG6_454053 | PF05730 | B0Y8A1 | IPR008427 | | 0 | scf_000006_A_fumigatus_A1163:1,305,295..1,306,089(+) |
| **AFUB_090110** | 696.6036695 | 2.582791799 | 0.118253129 | 21.84121311 | 9.42E-106 | 3.96E-104 | | AFUB_090110 | Has domain(s) with predicted ATP binding | Cation-transporting P-type ATF | P-type ATPase;Cation-trans | protein coding gene | GO:0016021 | GO:0005524;G | cation transport | integral compon | ATP binding;ATPase-cou | 3318 | 1105 | | 418 | 1 | OG6_100215 | PF00122;PF00 | B0YCG5 | IPR001757;IPR | 3.2.1.52 (Beta-N | 10 | scf_000008_A_fumigatus_A1163:1,032,657..1,035,974(+) |
| **AFUB_084920** | 6.079335326 | 2.583714899 | 0.849401272 | 3.041807194 | 0.002351625 | 0.003951911 | | AFUB_084920 | Protein of unknown function | N/A | N/A | protein coding gene | | | | | | 593 | 123 | | 0 | 0 | OG6r21_23999 | N/A | B0YAJ9 | N/A | | 0 | scf_000007_A_fumigatus_A1163:1,511,538..1,512,130(+) |
| **AFUB_045440** | 3.644012261 | 2.584566041 | 1.105046759 | 2.338874821 | 0.019341913 | 0.028753266 | | AFUB_045440 | Has domain(s) with predicted hydrolase a | Alpha/beta hydrolase fold-3 | Alpha/beta hydrolase fold-3; | protein coding gene | | GO:0016787 | metabolic process | | hydrolase activity | 684 | 227 | | 2599 | 8 | OG6_100160 | PF07859 | B0Y031 | IPR013094;IPR | 3.1.1.- (Carboxy | 0 | scf_000003_A_fumigatus_A1163:3,368,773..3,369,456(+) |
| **AFUB_037220** | 68.39852256 | 2.592723365 | 0.273134831 | 9.492466992 | 2.26E-21 | 1.11E-20 | | AFUB_037220 | Has domain(s) with predicted protein bindi | Fork head domain;Forkhead-as | Forkhead-associated (FHA) | protein coding gene | GO:0005634 | GO:0003700;G | regulation of tra | nucleus | DNA-binding transcription | 3070 | 722 | | 277 | 0 | OG6_104629 | PF00250;PF00 | B0XX89 | IPR000253;IPR001766;IPR0089 | | 0 | scf_000003_A_fumigatus_A1163:923,499..926,568(+) |
| **AFUB_065500** | 40.19218884 | 2.59331329 | 0.338179506 | 7.668451953 | 1.74E-14 | 6.38E-14 | | AFUB_065500 | Protein of unknown function | Protein of unknown function DU | Protein of unknown function | protein coding gene | | | | | | 654 | 217 | | 17 | 0 | OG6_453765 | PF04402 | B0Y633 | IPR007497 | | 0 | scf_000005_A_fumigatus_A1163:721,785..722,438(-) |
| **AFUB_072000** | 340.2750083 | 2.599533733 | 0.138042752 | 18.83136711 | 4.18E-79 | 9.66E-78 | | AFUB_072000 | Has domain(s) with predicted ATP binding | Protein kinase domain | Protein kinase domain;Serin | protein coding gene | | GO:0005524;G | protein phosphorylation | | ATP binding;protein kinas | 1624 | 360 | | 79 | 0 | OG6_102479 | PF00069 | | IPR000719;IPR | 2.7.11.1 (Non-sp | 0 | scf_000005_A_fumigatus_A1163:2,585,549..2,587,172(+) |
| **AFUB_003700** | 29.22585452 | 2.603889419 | 0.398902945 | 6.527626466 | 6.68E-11 | 2.02E-10 | | AFUB_003700 | Has domain(s) with predicted transmembr | Major facilitator, sugar transpor | Major facilitator, sugar trans | protein coding gene | GO:0016021 | GO:0022857 | transmembrane | integral compon | transmembrane transport | 1170 | 343 | | 26 | 0 | OG6_454902 | PF00083 | B0XMY1 | IPR005828;IPR036259 | | 7 | scf_000001_A_fumigatus_A1163:1,026,261..1,027,430(-) |
| **AFUB_075430** | 2131.878729 | 2.605691734 | 0.089516414 | 29.1085356 | 2.80E-186 | 3.97E-184 | | AFUB_075430 | Has domain(s) with predicted chromatin bi | N/A | Homeobox-like domain supe | protein coding gene | | GO:0003677;GO:0003682 | | | DNA binding;chromatin bi | 2483 | 630 | | 634 | 3 | OG6_100129 | PF13921 | B0Y7Y3 | IPR009057 | | 0 | scf_000006_A_fumigatus_A1163:949,135..951,617(-) |
| **AFUB_084150** | 230.2834018 | 2.607137768 | 0.157369483 | 16.5669844 | 1.21E-61 | 1.96E-60 | | AFUB_084150 | Has domain(s) with predicted iron ion bind | Fatty acid hydroxylase | Fatty acid hydroxylase | protein coding gene | | GO:0005506;G | fatty acid biosynthetic process;lip | | iron ion binding;oxidoredu | 1001 | 302 | | 388 | 1 | OG6_102540 | PF04116 | B0YAC2 | IPR006694 | 1.1.3.- (With oxy | 0 | scf_000007_A_fumigatus_A1163:1,282,062..1,283,062(+) |
| **AFUB_023450** | 7.390573439 | 2.611559982 | 0.772121701 | 3.382316514 | 0.000718773 | 0.001281187 | | AFUB_023450 | Protein of unknown function | N/A | N/A | protein coding gene | | | | | | 478 | 79 | | 0 | 0 | OG6r21_23986 | N/A | B0XW37 | N/A | | 0 | scf_000002_A_fumigatus_A1163:1,857,591..1,858,068(+) |
| **AFUB_058990** | 618.8281431 | 2.61236157 | 0.10957339 | 23.84120421 | 1.25E-125 | 7.52E-124 | | AFUB_058990 | Protein of unknown function | N/A | N/A | protein coding gene | | | | | | 3871 | 764 | | 97 | 0 | OG6_454118 | N/A | B0Y0Y8 | N/A | | 0 | scf_000004_A_fumigatus_A1163:2,836,051..2,839,921(-) |
| **AFUB_071800** | 13.4275703 | 2.613446696 | 0.585226272 | 4.46570296 | 7.98E-06 | 1.72E-05 | | AFUB_071800 | Protein of unknown function | Beta-ketoacyl synthase, N-term | Phosphopantetheine binding | protein coding gene | | GO:0003824;G | arugosin biosynthetic process;as | | catalytic activity;phosphop | 5532 | 1741 | | 3303 | 12 | OG6_100178 | PF00109;PF00 | B0Y591 | IPR009081;IPR | 1.1.1.100 (3-oxo | 0 | scf_000005_A_fumigatus_A1163:2,537,344..2,542,875(-) |
| **AFUB_090410** | 310.2593636 | 2.61625523 | 0.139137697 | 18.80335295 | 7.09E-79 | 1.63E-77 | | AFUB_090410 | Has domain(s) with predicted rRNA (aden | Ribosomal RNA adenine methy | Ribosomal RNA adenine me | protein coding gene | | GO:0052909;G | rRNA modification;rRNA process | | 18S rRNA (adenine(1779 | 1670 | 392 | | 338 | 0 | OG6_100569 | PF00398 | B0YCJ5 | IPR001737;IPR | 2.1.1.- (Methyltr | 0 | scf_000008_A_fumigatus_A1163:1,138,525..1,140,194(+) |
| **AFUB_077620** | 7.298626516 | 2.618355386 | 0.780553129 | 3.354487078 | 0.000795123 | 0.001410753 | | AFUB_077620 | Ortholog(s) have cellulase activity and role | Glycoside hydrolase, family 5;C | Cellulose-binding domain, fu | protein coding gene | GO:0005576 | GO:0003824;G | carbohydrate m | extracellular reg | catalytic activity;cellulase | 1397 | 397 | MKASTIICALLP | 351 | 2 | OG6_110872 | PF00150;PF00 | B0Y8K1 | IPR000254;IPR | 3.2.1.4 (Cellulas | 0 | scf_000006_A_fumigatus_A1163:1,563,551..1,564,947(-) |
| **AFUB_048370** | 13.62323267 | 2.619406365 | 0.589140912 | 4.446145753 | 8.74E-06 | 1.87E-05 | | AFUB_048370 | Has domain(s) with predicted metal ion bir | Class II aldolase/adducin N-ter | Class II aldolase/adducin N- | protein coding gene | | GO:0046872 | | | metal ion binding | 1021 | 294 | | 698 | 2 | OG6_100477 | PF00596 | B0XXH7 | IPR001303;IPR | 4.1.2.- (Aldehyd | 0 | scf_000003_A_fumigatus_A1163:4,128,026..4,129,046(-) |
| **AFUB_091860** | 3.723907573 | 2.620689171 | 1.115463643 | 2.349416933 | 0.018802839 | 0.027990755 | | AFUB_091860 | Has domain(s) with predicted transferase | Aminoglycoside phosphotransf | Aminoglycoside phosphotrar | protein coding gene | | GO:0016772 | | | transferase activity, transf | 957 | 318 | | 173 | 0 | OG6_454583 | PF01636 | B0YB79 | IPR002575;IPR011009 | | 0 | scf_000008_A_fumigatus_A1163:1,548,056..1,549,012(-) |
| **AFUB_051480** | 4.987809788 | 2.621404245 | 0.960024149 | 2.730560733 | 0.006322668 | 0.01006197 | | AFUB_051480 | Protein of unknown function | N/A | N/A | protein coding gene | | | | | | 445 | 94 | | 1 | 0 | OG6_454507 | N/A | B0Y2P4 | N/A | | 0 | scf_000004_A_fumigatus_A1163:722,869..723,313(-) |
| **AFUB_072550** | 37.05360767 | 2.624041277 | 0.353031659 | 7.432878063 | 1.06E-13 | 3.76E-13 | | AFUB_072550 | Protein of unknown function | Prenylated rab acceptor PRA1 | Prenylated rab acceptor PR | protein coding gene | GO:0016021 | | | integral component of membrane | | 525 | 174 | | 325 | 0 | OG6_102188 | PF03208 | B0Y748 | IPR004895 | | 3 | scf_000006_A_fumigatus_A1163:100,427..100,951(+) |
| **AFUB_017710** | 8.674821533 | 2.627514538 | 0.71867352 | 3.65606143 | 0.00025612 | 0.000478104 | | AFUB_017710 | Protein of unknown function | N/A | Glycosyl hydrolase, five-blad | protein coding gene | GO:0016021 | | | integral component of membrane | | 1583 | 482 | | 137 | 0 | OG6_137726 | N/A | B0XSV2 | IPR023296 | | 1 | scf_000002_A_fumigatus_A1163:133,322..134,904(+) |
| **AFUB_003650** | 7.397389148 | 2.631746276 | 0.786550408 | 3.345934666 | 0.000820057 | 0.001453923 | | AFUB_003650 | Has domain(s) with predicted ATP binding | ABC transporter-like | ABC transporter-like;P-loop | protein coding gene | GO:0016021 | GO:0005524;GO:0016887 | | integral compon | ATP binding;ATP hydroly | 437 | 139 | | 1 | 0 | OG6_455216 | PF00005 | B0XMX6 | IPR003439;IPR027417;IPR0394 | | 0 | scf_000001_A_fumigatus_A1163:1,016,582..1,017,018(-) |
| **AFUB_091480** | 139.3579107 | 2.633930283 | 0.193891211 | 13.584578 | 4.94E-42 | 4.90E-41 | | AFUB_091480 | Has domain(s) with predicted catalytic, mo | Molybdenum cofactor sulfurase | Molybdenum cofactor sulfura | protein coding gene | GO:0016021 | GO:0003824;GO:0030151;GO:0 | | integral compon | catalytic activity;molybder | 1365 | 454 | | 574 | 2 | OG6_101731 | PF03473;PF03 | B0YB41 | IPR005302;IPR005303;IPR0110 | | 2 | scf_000008_A_fumigatus_A1163:1,446,900..1,448,264(+) |
| **AFUB_048290** | 43.83416925 | 2.635535847 | 0.327359923 | 8.050881202 | 8.22E-16 | 3.18E-15 | | AFUB_048290 | Has domain(s) with predicted nucleotide b | Short-chain dehydrogenase/re | Short-chain dehydrogenase/ | protein coding gene | | GO:0016491 | metabolic process | | oxidoreductase activity | 852 | 267 | | 345 | 2 | OG6_119558 | PF00106 | B0XXG9 | IPR002347;IPR | 1.1.1.1 (Alcohol | 0 | scf_000003_A_fumigatus_A1163:4,099,922..4,100,773(+) |
| **AFUB_098600** | 120.0834349 | 2.635878025 | 0.210038463 | 12.54950157 | 4.00E-36 | 3.31E-35 | | AFUB_098600 | Has domain(s) with predicted protein bindi | IBR domain | IBR domain;E3 ubiquitin liga | protein coding gene | | GO:0046872;G | protein ubiquitination | | metal ion binding;ubiquitir | 1111 | 333 | | 115 | 0 | OG6_111184 | PF01485 | B0YEB6 | IPR002867;IPR | 2.3.2.31 (RBR-t | 0 | scf_000010_A_fumigatus_A1163:123,032..124,142(+) |
| **AFUB_022380** | 32.39680431 | 2.637541742 | 0.377091812 | 6.994428574 | 2.66E-12 | 8.73E-12 | | AFUB_022380 | Has domain(s) with predicted role in trans | Major facilitator superfamily | Major facilitator superfamily; | protein coding gene | GO:0016021;G | GO:0022857 | transmembrane | integral compon | transmembrane transport | 1510 | 426 | | 112 | 0 | OG6_103185 | PF07690 | B0XVI6 | IPR011701;IPR036259 | | 11 | scf_000002_A_fumigatus_A1163:1,476,046..1,477,555(+) |
| **AFUB_072080** | 315.7910785 | 2.637714431 | 0.152426436 | 17.30483574 | 4.33E-67 | 7.88E-66 | | AFUB_072080 | Has domain(s) with predicted transferase | Transferase | Transferase | protein coding gene | | GO:0016747 | | | acyltransferase activity, tr | 1710 | 478 | | 355 | 3 | OG6_107457 | PF02458 | B0Y5B9 | IPR003480 | 2.3.1.- (Transfer | 0 | scf_000005_A_fumigatus_A1163:2,601,826..2,603,535(+) |
| **AFUB_000960** | 81.93165939 | 2.646518391 | 0.245947868 | 10.76048519 | 5.29E-27 | 3.22E-26 | | AFUB_000960 | Has domain(s) with predicted sequence-s | Zn(2)-C6 fungal-type DNA-binc | Zn(2)-C6 fungal-type DNA-b | protein coding gene | GO:0042025;G | GO:0003677;G | regulation of tra | host cell nucleus | DNA binding;DNA-binding | 2353 | 712 | | 58 | 0 | OG6_454098 | PF00172;PF11 | B0XM53 | IPR001138;IPR021858;IPR0368 | | 0 | scf_000001_A_fumigatus_A1163:247,785..250,137(-) |
| **AFUB_094710** | 2256.447657 | 2.648835436 | 0.098058583 | 27.01278517 | 1.05E-160 | 1.05E-158 | | AFUB_094710 | Has domain(s) with predicted ATP binding | ABC transporter-like;Chromo d | Chromo/chromo shadow dor | protein coding gene | | GO:0005524;GO:0016887;GO:0017111;GO:000 | | | ATP binding;ATP hydroly | 3912 | 1117 | | 638 | 1 | OG6_105271 | PF00005;PF00385 | | IPR000953;IPR | 3.4.25.1 (Protea | 0 | scf_000009_A_fumigatus_A1163:520,767..524,678(+) |
| **AFUB_071640** | 678.1778057 | 2.649096292 | 0.108498585 | 24.41595247 | 1.16E-131 | 7.54E-130 | | AFUB_071640 | Has domain(s) with predicted chromatin bi | N/A | SANT/Myb domain;Homeob | protein coding gene | | GO:0003677;GO:0003682 | | | DNA binding;chromatin bi | 1545 | 514 | | 0 | 0 | OG6_507312 | N/A | B0Y575 | IPR001005;IPR009057 | | 0 | scf_000005_A_fumigatus_A1163:2,471,007..2,472,551(-) |
| **AFUB_089510** | 1240.338006 | 2.651644026 | 0.096569891 | 27.45828949 | 5.53E-166 | 5.79E-164 | | AFUB_089510 | Has domain(s) with predicted protein bindi | Proteasome component (PCI) | Proteasome component (PC | protein coding gene | GO:0005737;G | GO:0003743 | formation of cyt | cytoplasm;euka | translation initiation factor | 1806 | 468 | | 302 | 0 | OG6_103094 | PF01399;PF18 | B0YCA6 | IPR000717;IPR027528;IPR0407 | | 0 | scf_000008_A_fumigatus_A1163:887,147..888,952(+) |
| **AFUB_051450** | 572.6054383 | 2.654588964 | 0.118843165 | 22.33690905 | 1.62E-110 | 7.57E-109 | | AFUB_051450 | Has domain(s) with predicted protein bindi | L-asparaginase, N-terminal;An | Ankyrin repeat;Asparaginase | protein coding gene | | GO:0004067;G | cellular amino acid metabolic pro | | asparaginase activity;prot | 2390 | 560 | | 442 | 1 | OG6_100658 | PF00710;PF12 | B0Y2P1 | IPR002110;IPR | 3.1.1.47 (1-alkyl | 0 | scf_000004_A_fumigatus_A1163:710,581..712,970(-) |
| **AFUB_041390** | 5535.054867 | 2.655379183 | 0.075424636 | 35.20572743 | 1.63E-271 | 5.25E-269 | | AFUB_041390 | Has domain(s) with predicted nucleic acid | RNA recognition motif domain | RNA recognition motif doma | protein coding gene | | GO:0003723;GO:0003676;GO:0000166 | | | RNA binding;nucleic acid | 1747 | 546 | | 426 | 1 | OG6_102437 | PF00076 | B0XYM1 | IPR000504;IPR034276;IPR0359 | | 0 | scf_000003_A_fumigatus_A1163:2,101,758..2,103,504(-) |
| **AFUB_077920** | 5.153103348 | 2.656692504 | 0.942808409 | 2.817849816 | 0.004834642 | 0.007808684 | | AFUB_077920 | Ortholog(s) have hydrolase activity, hydrol | Zn(2)-C6 fungal-type DNA-binc | Zn(2)-C6 fungal-type DNA-b | protein coding gene | GO:0042025;G | GO:0003677;G | cellulose catabo | host cell nucleus | DNA binding;DNA-binding | 2455 | 676 | | 107 | 0 | OG6_454839 | PF00172 | B0Y8N0 | IPR001138;IPR036864 | | 0 | scf_000006_A_fumigatus_A1163:1,669,796..1,672,250(+) |
| **AFUB_030250** | 437.6424781 | 2.658461647 | 0.142505366 | 18.65516871 | 1.15E-77 | 2.56E-76 | | AFUB_030250 | Protein of unknown function | N/A | N/A | protein coding gene | | | | | | 955 | 281 | | 56 | 0 | OG6_186697 | N/A | B0XU42 | N/A | | 0 | scf_000002_A_fumigatus_A1163:3,803,679..3,804,633(-) |
| **AFUB_035100** | 54.30932224 | 2.66463369 | 0.297316234 | 8.962287928 | 3.18E-19 | 1.43E-18 | | AFUB_035100 | Protein of unknown function | N/A | N/A | protein coding gene | | | | | | 1519 | 204 | | 42 | 0 | OG6_453897 | N/A | B0Y082 | N/A | | 0 | scf_000003_A_fumigatus_A1163:306,715..308,233(+) |
| **AFUB_088750** | 34.45771883 | 2.667574315 | 0.374698555 | 7.119254348 | 1.09E-12 | 3.64E-12 | | AFUB_088750 | Ortholog(s) have role in mitotic recombina | DNA recombination and repair | AAA+ ATPase domain;DNA | protein coding gene | GO:0005634 | GO:0005524;G | DNA metabolic | nucleus | ATP binding;ATP-depend | 1518 | 338 | | 460 | 1 | OG6_100483 | PF08423 | B0YC34 | IPR003593;IPR | 2.7.11.1 (Non-sp | 0 | scf_000008_A_fumigatus_A1163:587,562..589,079(+) |
| **AFUB_075920** | 8.910336842 | 2.667992093 | 0.722760866 | 3.691389809 | 0.000223032 | 0.000418283 | | AFUB_075920 | Protein of unknown function | N/A | N/A | protein coding gene | | | | | | 1459 | 426 | MLLSATFLLLGI | 57 | 1 | OG6_186649 | N/A | B0Y831 | N/A | | 0 | scf_000006_A_fumigatus_A1163:1,121,039..1,122,497(-) |
| **AFUB_048980** | 19.07684103 | 2.66923508 | 0.492653336 | 5.41807978 | 6.02E-08 | 1.51E-07 | | AFUB_048980 | Protein of unknown function | Protein of unknown function DU | Protein of unknown function | protein coding gene | | | | | | 1371 | 456 | | 45 | 0 | OG6_153141 | PF11175 | B0Y179 | IPR021345 | | 0 | scf_000004_A_fumigatus_A1163:73,972..75,342(-) |
| **AFUB_016090** | 790.3129247 | 2.671553367 | 0.113633456 | 23.5102711 | 3.20E-122 | 1.84E-120 | | AFUB_016090 | Has domain(s) with predicted integral to m | ELO family | ELO family | protein coding gene | GO:0016021 | GO:0102336;G | fatty acid biosyn | integral compon | 3-oxo-arachidoyl-CoA syr | 2563 | 548 | | 249 | 0 | OG6_111476 | PF01151 | B0XNY3 | IPR002076 | 2.3.1.199 (Very- | 7 | scf_000001_A_fumigatus_A1163:4,561,769..4,564,331(-) |
| **AFUB_086080** | 63.90875413 | 2.672564706 | 0.274751569 | 9.72720453 | 2.31E-22 | 1.18E-21 | | AFUB_086080 | Has domain(s) with predicted nucleotide b | Polyketide synthase, ketoreduc | Polyketide synthase, ketorec | protein coding gene | | | | | | 3155 | 1017 | | 2 | 0 | OG6_454626 | PF08659;PF14 | B0YAW7 | IPR013968;IPR | 1.1.1.- (With NA | 0 | scf_000007_A_fumigatus_A1163:1,840,463..1,843,617(-) |
| **AFUB_082120** | 90.00575125 | 2.673837053 | 0.241712956 | 11.06203448 | 1.92E-28 | 1.23E-27 | | AFUB_082120 | Protein of unknown function | N/A | N/A | protein coding gene | | | | | | 954 | 317 | | 52 | 0 | OG6_453807 | N/A | B0Y9S7 | N/A | | 0 | scf_000007_A_fumigatus_A1163:670,654..671,607(-) |
| **AFUB_074700** | 6.343713382 | 2.674453407 | 0.893357658 | 2.99370961 | 0.002756081 | 0.004597126 | | AFUB_074700 | Protein of unknown function | Xylose isomerase-like, TIM bar | Xylose isomerase-like, TIM b | protein coding gene | | | | | | 1142 | 353 | | 497 | 3 | OG6_112484 | PF01261 | B0Y7R2 | IPR013022;IPR | 4.2.1.- (Hydro-ly | 0 | scf_000006_A_fumigatus_A1163:729,730..730,871(+) |
| **AFUB_084710** | 12.93644293 | 2.67450439 | 0.607592331 | 4.401807356 | 1.07E-05 | 2.28E-05 | | AFUB_084710 | Protein of unknown function | N/A | N/A | protein coding gene | GO:0016021 | | | integral component of membrane | | 1256 | 278 | MLVSSTSASIA | 78 | 0 | OG6_454824 | N/A | B0YAH7 | N/A | | 1 | scf_000007_A_fumigatus_A1163:1,461,115..1,462,370(-) |
| **AFUB_014020** | 346.3243342 | 2.676167391 | 0.134144215 | 19.9499277 | 1.50E-88 | 4.41E-87 | | AFUB_014020 | Has domain(s) with predicted catalytic, py | Aminotransferase, class I/class | Aminotransferase, class I/cla | protein coding gene | | GO:0003824;G | biosynthetic process | | catalytic activity;pyridoxal | 1617 | 446 | | 732 | 2 | OG6_100686 | PF00155 | B0XMS7 | IPR004839;IPR | 2.6.1.- (Transam | 0 | scf_000001_A_fumigatus_A1163:3,938,911..3,940,527(-) |
| **AFUB_097190** | 7.709947761 | 2.680029468 | 0.771994821 | 3.471564051 | 0.000517436 | 0.000935989 | | AFUB_097190 | Protein of unknown function | N/A | N/A | protein coding gene | | | | | | 921 | 278 | | 55 | 0 | OG6_453997 | N/A | B0YDY1 | N/A | | 0 | scf_000009_A_fumigatus_A1163:1,245,934..1,246,854(+) |
| **AFUB_016820** | 101.0215641 | 2.682998569 | 0.227419165 | 11.79759219 | 4.02E-32 | 2.93E-31 | | AFUB_016820 | Has domain(s) with predicted sequence-s | Zn(2)-C6 fungal-type DNA-binc | Zn(2)-C6 fungal-type DNA-b | protein coding gene | GO:0042025;G | GO:0003677;G | regulation of tra | host cell nucleus | DNA binding;DNA-binding | 2053 | 562 | | 40 | 0 | OG6_454382 | PF00172;PF11 | B0XPC2 | IPR001138;IPR021858;IPR0368 | | 0 | scf_000001_A_fumigatus_A1163:4,791,215..4,793,267(+) |
| **AFUB_067620** | 134.6800945 | 2.687883508 | 0.196919792 | 13.64963614 | 2.03E-42 | 2.03E-41 | | AFUB_067620 | Protein of unknown function | DPH-type metal-binding domai | DPH-type metal-binding dor | protein coding gene | | GO:0046872 | peptidyl-diphthamide biosyntheti | | metal ion binding | 321 | 106 | | 245 | 0 | OG6_103160 | PF05207 | B0Y6N9 | IPR007872;IPR036671 | | 0 | scf_000005_A_fumigatus_A1163:1,314,016..1,314,336(-) |
| **AFUB_000460** | 29.62386465 | 2.688773544 | 0.400253403 | 6.717678161 | 1.85E-11 | 5.78E-11 | | AFUB_000460 | Protein of unknown function | N/A | Nuclear mRNA export protei | protein coding gene | | | | | | 2356 | 631 | | 249 | 0 | OG6_121412 | N/A | B0XMA7 | IPR040114 | | 0 | scf_000001_A_fumigatus_A1163:131,773..134,128(-) |
| **AFUB_087760** | 6.446242631 | 2.690681472 | 0.85464152 | 3.148315883 | 0.001642141 | 0.002813098 | | AFUB_087760 | Protein of unknown function | Carboxylesterase, type B | Carboxylesterase, type B;Ca | protein coding gene | | GO:0016787 | | | hydrolase activity | 1965 | 587 | | 1660 | 5 | OG6_100080 | PF00135 | B0YBT5 | IPR002018;IPR | 3.1.1.1 (Carboxy | 1 | scf_000008_A_fumigatus_A1163:313,022..314,986(-) |
| **AFUB_046040** | 5.110123158 | 2.697360615 | 0.951426428 | 2.835070098 | 0.004581563 | 0.007422317 | | AFUB_046040 | Has domain(s) with predicted sulfuric este | Sulfatase, N-terminal | Sulfatase, N-terminal;Alkalin | protein coding gene | | GO:0003824;G | metabolic process | | catalytic activity;sulfuric e | 1861 | 599 | | 296 | 1 | OG6_100690 | PF00884 | B0Y0F8 | IPR000917;IPR | 3.1.6.1 (Arylsulf | 0 | scf_000003_A_fumigatus_A1163:3,545,579..3,547,439(-) |
| **AFUB_002910** | 103.7662048 | 2.698700072 | 0.223620185 | 12.06823109 | 1.55E-33 | 1.19E-32 | | AFUB_002910 | Has domain(s) with predicted substrate-sp | Major facilitator, sugar transpor | Sugar/inositol transporter;Ma | protein coding gene | GO:0016021;G | GO:0022857 | carbohydrate tra | integral compon | transmembrane transport | 1640 | 503 | | 141 | 0 | OG6_129090 | PF00083 | B0XMJ1 | IPR003663;IPR005828;IPR0058 | | 12 | scf_000001_A_fumigatus_A1163:809,903..811,542(+) |
| **AFUB_090390** | 12.96902711 | 2.704972972 | 0.604595316 | 4.474022373 | 7.68E-06 | 1.65E-05 | | AFUB_090390 | Has domain(s) with predicted catalytic acti | N/A | Six-hairpin glycosidase supe | protein coding gene | | GO:0003824 | | | catalytic activity | 3184 | 707 | | 18 | 0 | OG6_455294 | N/A | B0YCJ3 | IPR008928 | | 0 | scf_000008_A_fumigatus_A1163:1,132,810..1,135,993(+) |
| **AFUB_071040** | 150.9430419 | 2.70585977 | 0.194007014 | 13.94722652 | 3.27E-44 | 3.47E-43 | | AFUB_071040 | Protein of unknown function | N/A | N/A | protein coding gene | | | | | | 972 | 233 | | 2 | 0 | OG6r21_13789 | N/A | B0Y514 | N/A | | 0 | scf_000005_A_fumigatus_A1163:2,260,633..2,261,604(+) |
| **AFUB_060200** | 344.3995575 | 2.70737323 | 0.142726425 | 18.9689697 | 3.08E-80 | 7.34E-79 | | AFUB_060200 | Ortholog(s) have role in protein ubiquitinat | LDB19, N-terminal | Immunoglobulin E-set;LDB1 | protein coding gene | | | protein ubiquitination | | | 2352 | 525 | | 257 | 0 | OG6_133449 | PF13002 | B0Y1G0 | IPR014756;IPR024391 | | 0 | scf_000004_A_fumigatus_A1163:3,151,222..3,153,573(-) |
| **AFUB_034300** | 394.2354711 | 2.707438157 | 0.129643277 | 20.88375288 | 7.52E-97 | 2.63E-95 | | AFUB_034300 | Ortholog(s) have enzyme inhibitor activity | Proteinase inhibitor I78 | Proteinase inhibitor I78 | protein coding gene | GO:0005576 | GO:0004857;GO:0004867 | | extracellular reg | enzyme inhibitor activity;s | 1038 | 87 | MKFSLACLLAL | 10 | 0 | OG6_453930 | PF11720 | B0XZT4 | IPR021719 | | 0 | scf_000003_A_fumigatus_A1163:116,174..117,211(+) |
| **AFUB_019750** | 15.51983234 | 2.707735992 | 0.545850258 | 4.960583883 | 7.03E-07 | 1.63E-06 | | AFUB_019750 | Protein of unknown function | N/A | N/A | protein coding gene | GO:0016021 | | | integral component of membrane | | 3297 | 1098 | | 7 | 0 | OG6_454580 | N/A | B0XU07 | N/A | | 1 | scf_000002_A_fumigatus_A1163:670,915..674,211(+) |
| **AFUB_085860** | 86.12935866 | 2.709484637 | 0.241159677 | 11.2352308 | 2.74E-29 | 1.81E-28 | | AFUB_085860 | Has domain(s) with predicted role in defen | Antimicrobial protein MiAMP1 | Gamma-crystallin-like;Antimi | protein coding gene | | | defense response;negative regulation of growth | | | 306 | 101 | MKFLAIATLLAS | 9 | 0 | OG6_454834 | PF09117 | B0YAU4 | IPR011024;IPR015201 | | 0 | scf_000007_A_fumigatus_A1163:1,781,493..1,781,798(-) |
| **AFUB_030650** | 266.6517111 | 2.710342548 | 0.154335119 | 17.56141164 | 4.87E-69 | 9.28E-68 | | AFUB_030650 | Has domain(s) with predicted calcium ion | Mechanosensitive ion channel | Mechanosensitive ion chann | protein coding gene | GO:0005789;G | GO:0005509 | transmembrane | endoplasmic ret | calcium ion binding | 3915 | 949 | | 221 | 0 | OG6_112962 | PF00924 | B0XU81 | IPR006685;IPR010920;IPR0119 | | 5 | scf_000002_A_fumigatus_A1163:3,925,189..3,929,103(+) |
| **AFUB_091080** | 93.3721145 | 2.714830206 | 0.236320493 | 11.48791698 | 1.52E-30 | 1.05E-29 | | AFUB_091080 | Ortholog(s) have glutathione transferase, | Glutathione S-transferase, C-te | Glutathione S-transferase, N | protein coding gene | GO:0005622 | GO:0004602;GO:0004364;GO:0 | | intracellular ana | glutathione peroxidase ac | 1907 | 230 | | 1178 | 3 | OG6_100130 | PF00043;PF02 | B0YCR0 | IPR004045;IPR | 1.11.1.- (Peroxid | 0 | scf_000008_A_fumigatus_A1163:1,329,313..1,331,219(+) |
| **AFUB_001390** | 5.215110425 | 2.717160553 | 0.950048669 | 2.860022481 | 0.00423611 | 0.006891635 | | AFUB_001390 | unspecified product | N/A | N/A | ncRNA gene | | | | | | 111 | | | 0 | 0 | N/A (orthology r | N/A | | N/A | | | scf_000001_A_fumigatus_A1163:368,081..368,191(+) |
| **AFUB_029830** | 3926.131091 | 2.719388653 | 0.078581855 | 34.60580899 | 2.07E-262 | 5.53E-260 | | AFUB_029830 | Ortholog(s) have dihydroxy-acid dehydrat | Dihydroxy-acid/6-phosphogluc | Dihydroxy-acid/6-phosphogl | protein coding gene | GO:0005622 | GO:0051539;G | branched-chain | intracellular ana | 4 iron, 4 sulfur cluster bin | 2245 | 642 | MMLTILGCGLA | 553 | 1 | OG6_102200 | PF00920 | B0XTT1 | IPR000581;IPR | 4.2.1.9 (Dihydro | 0 | scf_000002_A_fumigatus_A1163:3,694,210..3,696,454(+) |
| **AFUB_014560** | 1527.304575 | 2.719938756 | 0.096336518 | 28.2337249 | 2.26E-175 | 2.65E-173 | | AFUB_014560 | Has domain(s) with predicted 2-isopropyln | Pyruvate carboxyltransferase;2 | Pyruvate carboxyltransferase | protein coding gene | | GO:0003852;G | carboxylic acid metabolic proces | | 2-isopropylmalate synthas | 2369 | 643 | | 315 | 0 | OG6_110817 | PF00682;PF08 | B0XN47 | IPR000891;IPR | 2.3.1.182 ((R)-c | 0 | scf_000001_A_fumigatus_A1163:4,096,465..4,098,833(+) |
| **AFUB_075490** | 5.304911546 | 2.725715548 | 0.927408876 | 2.939065626 | 0.003292034 | 0.005434647 | | AFUB_075490 | Protein of unknown function | N/A | LamG-like jellyroll fold;Conca | protein coding gene | GO:0016021 | GO:0016787 | | integral compon | hydrolase activity | 5153 | 1681 | | 2 | 0 | OG6r21_12618 | PF13385 | B0Y7Y8 | IPR006558;IPR013320;IPR0368 | | 2 | scf_000006_A_fumigatus_A1163:979,612..984,764(+) |
| **AFUB_097540** | 15.7351169 | 2.727687341 | 0.552510178 | 4.936899718 | 7.94E-07 | 1.83E-06 | | AFUB_097540 | Protein of unknown function | N/A | Macro domain-like | protein coding gene | | | | | | 533 | 156 | | 135 | 1 | OG6_132750 | N/A | B0YE19 | IPR043472 | | 0 | scf_000009_A_fumigatus_A1163:1,361,160..1,361,692(+) |
| **AFUB_078070** | 1990.367 | 2.731412746 | 0.101762303 | 26.84110595 | 1.07E-158 | 1.05E-156 | | AFUB_078070 | Protein of unknown function | AMP-dependent synthetase/lig | AMP-dependent synthetase/ | protein coding gene | GO:0031410 | GO:0003824;G | brevianamide F | cytoplasmic ves | catalytic activity;ligase act | 11990 | 3955 | | 2509 | 12 | OG6_100454 | PF00501;PF00 | B0Y8P5 | IPR000873;IPR | 2.1.-.- (Transfer | 0 | scf_000006_A_fumigatus_A1163:1,711,510..1,723,499(-) |
| **AFUB_017850** | 459.9339317 | 2.733582408 | 0.12940357 | 21.12447446 | 4.74E-99 | 1.73E-97 | | AFUB_017850 | Has domain(s) with predicted oxidoreduct | FAD-binding domain | FAD-binding domain;FAD/N | protein coding gene | | GO:0071949;G | metabolic process | | FAD binding;oxidoreducta | 1381 | 442 | | 165 | 2 | OG6_152350 | PF01494 | B0XT33 | IPR002938;IPR | 1.14.13.1 (Salic | 0 | scf_000002_A_fumigatus_A1163:163,179..164,559(-) |
| **AFUB_017790** | 17.33553029 | 2.734049012 | 0.536326151 | 5.097735781 | 3.44E-07 | 8.16E-07 | | AFUB_017790 | Ortholog(s) have alpha-amylase activity, r | Glycosyl hydrolase, family 13, | Carbohydrate binding modul | protein coding gene | GO:0030428;G | GO:0004556;G | carbohydrate ca | cell septum;cell | alpha-amylase activity;alp | 2370 | 630 | MKWISQLFPLS | 389 | 2 | OG6_104504 | PF00128;PF00 | B0XT27 | IPR002044;IPR | 3.2.1.1 (Alpha-a | 0 | scf_000002_A_fumigatus_A1163:149,091..151,460(+) |
| **AFUB_006670** | 124.6285138 | 2.734277026 | 0.207845806 | 13.15531487 | 1.59E-39 | 1.44E-38 | | AFUB_006670 | Has domain(s) with predicted protein bindi | WD40 repeat;Sof1-like protein | WD40 repeat;Sof1-like prote | protein coding gene | | GO:0005515 | | | protein binding | 1459 | 466 | | 304 | 0 | OG6_102358 | PF00400;PF04158 | | IPR001680;IPR | 2.3.1.48 (Histon | 1 | scf_000001_A_fumigatus_A1163:1,882,384..1,883,842(-) |
| **AFUB_093040** | 41.05782629 | 2.735974836 | 0.341636662 | 8.008434525 | 1.16E-15 | 4.47E-15 | | AFUB_093040 | Protein of unknown function | N/A | N/A | protein coding gene | | | | | | 656 | 131 | | 0 | 0 | OG6r21_23991 | N/A | B0YCS6 | N/A | | 0 | scf_000009_A_fumigatus_A1163:52,189..52,844(-) |
| **AFUB_081520** | 3.967208029 | 2.736589842 | 1.072253991 | 2.552184337 | 0.010704986 | 0.016488691 | | AFUB_081520 | unspecified product | N/A | N/A | ncRNA gene | | | | | | 85 | | | 0 | 0 | N/A (orthology r | N/A | | N/A | | | scf_000007_A_fumigatus_A1163:508,465..508,549(-) |
| **AFUB_057840** | 484.8691412 | 2.741213652 | 0.127058846 | 21.57436296 | 3.13E-103 | 1.23E-101 | | AFUB_057840 | Protein of unknown function | LPS-induced tumour necrosis f | LPS-induced tumour necrosi | protein coding gene | GO:0016020 | GO:0046872 | | membrane | metal ion binding | 895 | 210 | | 225 | 0 | OG6_100529 | PF10601 | B0Y0M5 | IPR006629;IPR | 3.1.13.4 (Poly(A | 0 | scf_000004_A_fumigatus_A1163:2,524,186..2,525,080(-) |
| **AFUB_017380** | 126.749156 | 2.747152983 | 0.202877127 | 13.54096949 | 8.96E-42 | 8.77E-41 | | AFUB_017380 | Has domain(s) with predicted iron ion bind | Fatty acid hydroxylase | Fatty acid hydroxylase | protein coding gene | GO:0016021 | GO:0005506;G | fatty acid biosyn | integral compon | iron ion binding;oxidoredu | 1134 | 335 | | 422 | 2 | OG6_102250 | PF04116 | B0XSS0 | IPR006694 | 1.14.19.20 (Delt | 3 | scf_000002_A_fumigatus_A1163:52,861..53,994(+) |
| **AFUB_077060** | 18.57834707 | 2.750553191 | 0.504867696 | 5.448067307 | 5.09E-08 | 1.29E-07 | | AFUB_077060 | Has domain(s) with predicted nucleic acid | RNA recognition motif domain | RNA recognition motif doma | protein coding gene | | GO:0003723;GO:0003676;GO:0000166 | | | RNA binding;nucleic acid | 2120 | 624 | | 210 | 0 | OG6_105551 | PF00076 | B0Y8E5 | IPR000504;IPR035979 | | 0 | scf_000006_A_fumigatus_A1163:1,407,769..1,409,888(+) |
| **AFUB_077330** | 3001.114676 | 2.752090557 | 0.082816576 | 33.23115589 | 3.82E-242 | 8.77E-240 | | AFUB_077330 | Ortholog(s) have carbamoyl-phosphate sy | Glutamine amidotransferase;A | Aspartate carbamoyltransfer | protein coding gene | | GO:0005524;G | de novo' pyrimidine nucleobase | | ATP binding;amino acid b | 7640 | 2256 | | 665 | 1 | OG6_100435 | PF00117;PF00185;PF00988;PF | | IPR002082;IPR | 2.1.3.2 (Asparta | 0 | scf_000006_A_fumigatus_A1163:1,488,432..1,496,071(+) |
| **AFUB_044820** | 181.9054287 | 2.75308786 | 0.179007955 | 15.37969558 | 2.24E-53 | 3.01E-52 | | AFUB_044820 | Has domain(s) with predicted ATP binding | ABC transporter-like;ABC trans | ABC transporter-like;AAA+ A | protein coding gene | GO:0016021 | GO:0005524;G | transmembrane | integral compon | ATP binding;ATP hydroly | 4060 | 1315 | | 2370 | 9 | OG6_100059 | PF00005;PF00 | B0XZQ0 | IPR003439;IPR | 2.7.2.8 (Acetylgl | 11 | scf_000003_A_fumigatus_A1163:3,194,528..3,198,587(+) |
| **AFUB_056410** | 4.030578908 | 2.753938507 | 1.068386124 | 2.577662181 | 0.009947119 | 0.015417475 | | AFUB_056410 | Protein of unknown function | Nitrogen regulatory protein are | Nitrogen regulatory protein a | protein coding gene | | | | | | 1687 | 476 | | 41 | 0 | OG6_453817 | PF08550 | B0Y439 | IPR013860 | | 0 | scf_000004_A_fumigatus_A1163:2,177,153..2,178,839(-) |
| **AFUB_013940** | 529.3228687 | 2.755450508 | 0.134913427 | 20.42384191 | 1.03E-92 | 3.30E-91 | | AFUB_013940 | Has domain(s) with predicted NAD binding | D-isomer specific 2-hydroxyaci | D-isomer specific 2-hydroxy | protein coding gene | | GO:0051287;G | metabolic process;obsolete oxid | | NAD binding;obsolete cof | 1335 | 335 | | 680 | 1 | OG6_100397 | PF00389;PF02 | B0XMR9 | IPR006139;IPR | 1.1.1.215 (Gluc | 0 | scf_000001_A_fumigatus_A1163:3,919,155..3,920,489(+) |
| **AFUB_040960** | 33.78320716 | 2.761002534 | 0.380976351 | 7.247175658 | 4.26E-13 | 1.46E-12 | | AFUB_040960 | Protein of unknown function | N/A | N/A | protein coding gene | | | | | | 1137 | 355 | | 48 | 0 | OG6_455275 | N/A | B0XYH7 | N/A | | 0 | scf_000003_A_fumigatus_A1163:1,961,327..1,962,463(+) |
| **AFUB_011180** | 9.453867246 | 2.763330869 | 0.726121616 | 3.805603372 | 0.000141459 | 0.000270349 | | AFUB_011180 | Protein of unknown function | Protein of unknown function DU | Protein of unknown function | protein coding gene | GO:0016021 | | | integral component of membrane | | 795 | 264 | | 309 | 0 | OG6_129920 | PF17648 | B0XQU4 | IPR040841 | | 1 | scf_000001_A_fumigatus_A1163:3,156,409..3,157,203(-) |
| **AFUB_018490** | 9.58087641 | 2.769509179 | 0.714933904 | 3.873797512 | 0.000107152 | 0.000207793 | | AFUB_018490 | Protein of unknown function | Cyanovirin-N | Cyanovirin-N;Cyanovirin-N s | protein coding gene | | | | | | 421 | 111 | | 20 | 0 | OG6_454760 | PF08881 | B0XTG4 | IPR011058;IPR036673 | | 0 | scf_000002_A_fumigatus_A1163:328,632..329,052(-) |
| **AFUB_092120** | 503.9417439 | 2.769682828 | 0.124481935 | 22.2496768 | 1.14E-109 | 5.21E-108 | | AFUB_092120 | Has domain(s) with predicted lyase, pyrido | Aromatic amino acid beta-elimi | Aromatic amino acid beta-eli | protein coding gene | | GO:0003824;G | cellular amino acid metabolic pro | | catalytic activity;lyase acti | 1141 | 360 | | 67 | 0 | OG6_120858 | PF01212 | B0YBA5 | IPR001597;IPR | 4.1.2.5 (L-threon | 0 | scf_000008_A_fumigatus_A1163:1,599,566..1,600,706(+) |
| **AFUB_081040** | 64.77591655 | 2.771836916 | 0.279421776 | 9.919903002 | 3.41E-23 | 1.80E-22 | | AFUB_081040 | Has domain(s) with predicted role in trans | Major facilitator superfamily | Major facilitator superfamily; | protein coding gene | GO:0016021;G | GO:0022857 | transmembrane | integral compon | transmembrane transport | 1729 | 513 | | 624 | 2 | OG6_112461 | PF07690 | B0Y9H3 | IPR011701;IPR036259 | | 14 | scf_000007_A_fumigatus_A1163:378,958..380,686(+) |
| **AFUB_071300** | 6.73098206 | 2.773463376 | 0.83793199 | 3.309890789 | 0.000933324 | 0.00164235 | | AFUB_071300 | Has domain(s) with predicted transferase | UDP-glucuronosyl/UDP-glucos | UDP-glucuronosyl/UDP-gluc | protein coding gene | | GO:0008194;G | metabolic process | | UDP-glycosyltransferase | 2663 | 855 | | 537 | 2 | OG6_100399 | PF00201;PF03 | B0Y540 | IPR002213;IPR | 2.3.1.20 (Diacyl | 0 | scf_000005_A_fumigatus_A1163:2,332,101..2,334,763(+) |
| **AFUB_046300** | 6.699411752 | 2.779812886 | 0.844265472 | 3.292581513 | 0.000992721 | 0.001742416 | | AFUB_046300 | Has domain(s) with predicted nucleotide b | Short-chain dehydrogenase/re | Short-chain dehydrogenase/ | protein coding gene | | GO:0016491 | metabolic process | | oxidoreductase activity | 1083 | 330 | | 2995 | 7 | OG6_100091 | PF00106 | B0Y0I3 | IPR002347;IPR | 1.1.1.100 (3-oxo | 0 | scf_000003_A_fumigatus_A1163:3,602,023..3,603,105(+) |
| **AFUB_001130** | 17.85057464 | 2.792437613 | 0.518334769 | 5.387324521 | 7.15E-08 | 1.78E-07 | | AFUB_001130 | Ortholog(s) have endo-1,4-beta-xylanase | Glycoside hydrolase family 10 | Cellulose-binding domain, fu | protein coding gene | GO:0005576 | GO:0003824;G | carbohydrate m | extracellular reg | catalytic activity;cellulose | 1415 | 397 | MVHLSSLAAAL | 515 | 2 | OG6_102896 | PF00331;PF00 | B0XM69 | IPR000254;IPR | 3.2.1.155 (Xylog | 0 | scf_000001_A_fumigatus_A1163:304,672..306,086(-) |
| **AFUB_051860** | 177.56429 | 2.792780637 | 0.184054811 | 15.1736356 | 5.29E-52 | 6.83E-51 | | AFUB_051860 | Protein of unknown function | Questin oxidase-like | Questin oxidase-like | protein coding gene | | GO:0016491 | | | oxidoreductase activity | 2399 | 507 | | 535 | 3 | OG6_109991 | PF14027 | | IPR025337 | 1.14.13.43 (Que | 0 | scf_000004_A_fumigatus_A1163:829,940..832,338(+) |

### Table S5

| | | | | | | | | | | | | | | | | | | | | | | | | | | | | | |
|---|---|---|---|---|---|---|---|---|---|---|---|---|---|---|---|---|---|---|---|---|---|---|---|---|---|---|---|---|---|
| **AFUB_014140** | 1258.065455 | 2.794761477 | 0.092387992 | 30.25026761 | 5.18E-201 | 8.60E-199 | | AFUB_014140 | Protein of unknown function | Lipin, N-terminal;Lipin/Ned1/Sr | Lipin, N-terminal;Lipin/Ned1/ | protein coding gene | | | | | | 2991 | 765 | | 316 | 0 | OG6_100990 | PF04571;PF082 | B0XMT9 | IPR007651;IPR( | 3.1.3.4 (Phosph | 0 | scf_000001_A_fumigatus_A1163:3,968,330..3,971,320(-) |
| **AFUB_013280** | 29.09399468 | 2.802598982 | 0.410797858 | 6.822331054 | 8.96E-12 | 2.86E-11 | | AFUB_013280 | Has domain(s) with predicted role in transı | Major facilitator superfamily | Major facilitator superfamily;l | protein coding gene | GO:0016021;G( | GO:0022857 | transmembrane | integral compon | transmembrane transporte | 3365 | 607 | | 517 | 2 | OG6_121378 | PF07690 | B0XRN3 | IPR011701;IPR036259 | | 12 | scf_000001_A_fumigatus_A1163:3,751,896..3,755,260(-) |
| **AFUB_075410** | 8.416029375 | 2.803724384 | 0.761323411 | 3.682698238 | 0.000230778 | 0.000432221 | | AFUB_075410 | Protein of unknown function | N/A | N/A | protein coding gene | | | | | | 1010 | 181 | | 0 | 0 | OG6r21_23991! | N/A | B0Y7Y1 | N/A | | 0 | scf_000006_A_fumigatus_A1163:944,321..945,330(-) |
| **AFUB_033160** | 26.19719358 | 2.805335441 | 0.429070957 | 6.538162035 | 6.23E-11 | 1.88E-10 | | AFUB_033160 | Protein of unknown function | Amino acid transporter, transm | Amino acid transporter, trans | protein coding gene | GO:0016021 | | | integral component of membrane | | 1751 | 474 | | 614 | 5 | OG6_124686 | PF01490 | B0XVP4 | IPR013057 | | 10 | scf_000002_A_fumigatus_A1163:4,621,602..4,623,352(+) |
| **AFUB_024247** | 8.290766178 | 2.806455024 | 0.753878797 | 3.722687301 | 0.000197114 | 0.000371482 | | AFUB_024247 | Has domain(s) with predicted NAD binding | Malic enzyme, N-terminal doma | Malic oxidoreductase;Malic e | protein coding gene | | GO:0051287;G( | malate metabolic process;obsole | | NAD binding;malate dehy | 2355 | 661 | | 560 | 1 | OG6_100205 | PF00390;PF03949 | | IPR001891;IPR( | 1.1.1.38 (Malate | 0 | scf_000002_A_fumigatus_A1163:2,117,595..2,119,949(-) |
| **AFUB_061610** | 59.75517061 | 2.807693767 | 0.291264675 | 9.639664575 | 5.44E-22 | 2.73E-21 | | AFUB_061610 | Protein of unknown function | N/A | N/A | protein coding gene | | | | | | 315 | 85 | | 395 | 1 | OG6_101458 | N/A | B0Y282 | N/A | | 0 | scf_000004_A_fumigatus_A1163:3,548,907..3,549,221(-) |
| **AFUB_076300** | 23.34969695 | 2.80810133 | 0.456971346 | 6.145027156 | 7.99E-10 | 2.26E-09 | | AFUB_076300 | Ortholog(s) have role in pathogenesis, cel | Signal transduction response r | PAS domain;Signal transduc | protein coding gene | GO:0016020 | GO:0005524;G( | cellular respons | membrane | ATP binding;phosphorela | 3101 | 708 | | 148 | 0 | OG6_108005 | PF00072;PF005 | B0Y869 | IPR000014;IPR( | 2.7.-.- (Transfer | 0 | scf_000006_A_fumigatus_A1163:1,218,755..1,221,855(+) |
| **AFUB_097120** | 96.62005428 | 2.808653754 | 0.232149056 | 12.0984931 | 1.08E-33 | 8.25E-33 | | AFUB_097120 | Protein of unknown function | Fungal transcription factor | Fungal transcription factor | protein coding gene | | | | | | 1140 | 379 | MHHFVTQTIRV | 59 | 0 | OG6_454867 | PF11951 | B0YDX4 | IPR021858 | | 0 | scf_000009_A_fumigatus_A1163:1,236,188..1,237,327(+) |
| **AFUB_083490** | 60.946118 | 2.809025957 | 0.289816765 | 9.692420523 | 3.25E-22 | 1.65E-21 | | AFUB_083490 | Has domain(s) with predicted glucosamine | Glucosamine/galactosamine-6- | Glucosamine-6-phosphate is | protein coding gene | | GO:0004342 | N-acetylglucosamine metabolic p | | glucosamine-6-phosphate | 1938 | 358 | | 298 | 0 | OG6_102139 | PF01182 | | IPR004547;IPR( | 3.5.99.6 (Glucos | 0 | scf_000007_A_fumigatus_A1163:1,049,023..1,050,960(-) |
| **AFUB_101870** | 344.6514011 | 2.809951662 | 0.139488446 | 20.14469119 | 3.00E-90 | 9.14E-89 | | AFUB_101870 | Has domain(s) with predicted sequence-s| | Zn(2)-C6 fungal-type DNA-binc | Zn(2)-C6 fungal-type DNA-b | protein coding gene | GO:0042025;G( | GO:0003677;G( | regulation of tra | host cell nucleus | DNA binding;DNA-binding | 3284 | 819 | | 22 | 0 | OG6_186794 | PF00172 | B0YF81 | IPR001138;IPR036864 | | 0 | scf_000011_A_fumigatus_A1163:396,961..400,244(+) |
| **AFUB_023200** | 25.1183714 | 2.810751756 | 0.442576458 | 6.350884029 | 2.14E-10 | 6.28E-10 | | AFUB_023200 | Protein of unknown function | N/A | N/A | protein coding gene | | | | | | 3155 | 659 | | 48 | 0 | OG6_454291 | N/A | | N/A | | 0 | scf_000002_A_fumigatus_A1163:1,722,508..1,725,662(-) |
| **AFUB_076220** | 26.31583778 | 2.811271982 | 0.428487247 | 6.560923344 | 5.35E-11 | 1.62E-10 | | AFUB_076220 | Protein of unknown function | Meiotically up-regulated gene 1 | Meiotically up-regulated gen | protein coding gene | | | | | | 987 | 328 | | 62 | 0 | OG6_166549 | PF15474 | B0Y861 | IPR029167 | | 0 | scf_000006_A_fumigatus_A1163:1,201,055..1,202,041(-) |
| **AFUB_097470** | 75.04845544 | 2.815838535 | 0.26283617 | 10.713284 | 8.82E-27 | 5.29E-26 | | AFUB_097470 | Protein of unknown function | N/A | N/A | protein coding gene | | | | | | 792 | 263 | | 179 | 1 | OG6_453771 | N/A | B0YE12 | N/A | | 0 | scf_000009_A_fumigatus_A1163:1,335,101..1,335,892(+) |
| **AFUB_037000** | 20.97390402 | 2.816737701 | 0.480430052 | 5.862950681 | 4.55E-09 | 1.23E-08 | | AFUB_037000 | Has domain(s) with predicted sequence-s| | Zn(2)-C6 fungal-type DNA-binc | Zn(2)-C6 fungal-type DNA-b | protein coding gene | GO:0042025;G( | GO:0003677;G( | regulation of tra | host cell nucleus | DNA binding;DNA-binding | 2380 | 692 | | 523 | 4 | OG6_143648 | PF00172;PF04( | B0XX67 | IPR001138;IPR007219;IPR0368 | | 0 | scf_000003_A_fumigatus_A1163:862,261..864,640(-) |
| **AFUB_003629** | 7.07836102 | 2.819345907 | 0.829576992 | 3.398534356 | 0.00067748 | 0.001210596 | | AFUB_003629 | Protein of unknown function | N/A | N/A | protein coding gene | | | | | | 927 | 167 | | 1 | 0 | OG6_453828 | N/A | B0XMX3 | N/A | | 0 | scf_000001_A_fumigatus_A1163:1,004,048..1,004,974(+) |
| **AFUB_015560** | 261.3423985 | 2.821257788 | 0.155079048 | 18.19238522 | 5.93E-74 | 1.24E-72 | | AFUB_015560 | Has domain(s) with predicted sequence-s| | Zn(2)-C6 fungal-type DNA-binc | Zn(2)-C6 fungal-type DNA-b | protein coding gene | GO:0042025;G( | GO:0003677;G( | regulation of tra | host cell nucleus | DNA binding;DNA-binding | 3286 | 753 | | 409 | 0 | OG6_152901 | PF00172;PF04082 | | IPR001138;IPR007219;IPR0368 | | 0 | scf_000001_A_fumigatus_A1163:4,422,216..4,425,501(+) |
| **AFUB_064050** | 9.689043022 | 2.822438003 | 0.702090352 | 4.020049549 | 5.82E-05 | 0.000115604 | | AFUB_064050 | Protein of unknown function | N/A | N/A | protein coding gene | | | | | | 3125 | 853 | | 32 | 0 | OG6_455361 | N/A | B0Y5N9 | N/A | | 0 | scf_000005_A_fumigatus_A1163:352,269..355,393(-) |
| **AFUB_026050** | 5.483255098 | 2.823120203 | 0.950182426 | 2.971134939 | 0.002967014 | 0.004920851 | | AFUB_026050 | unspecified product | N/A | N/A | ncRNA gene | | | | | | 72 | | | 0 | 0 | N/A (orthology r | N/A | | N/A | | | scf_000002_A_fumigatus_A1163:2,594,798..2,594,869(+) |
| **AFUB_043340** | 110.8011464 | 2.824967136 | 0.223658651 | 12.63070812 | 1.43E-36 | 1.20E-35 | | AFUB_043340 | Has domain(s) with predicted role in transı | Major facilitator superfamily | Major facilitator superfamily;l | protein coding gene | GO:0016021;G( | GO:0022857 | transmembrane | integral compon | transmembrane transporte | 1700 | 484 | | 2612 | 9 | OG6_100246 | PF07690 | B0XZ68 | IPR011701;IPR( | 2.8.2.11 (Galact | 12 | scf_000003_A_fumigatus_A1163:2,666,923..2,668,622(-) |
| **AFUB_004020** | 256.9234556 | 2.829390834 | 0.161919308 | 17.4740793 | 2.26E-68 | 4.22E-67 | | AFUB_004020 | Has domain(s) with predicted oxygen bind | N/A | N/A | protein coding gene | | GO:0020037;G( | oxygen transport | | heme binding;iron ion bin | 978 | 282 | | 259 | 0 | OG6_116977 | PF11563 | B0XN13 | N/A | | 0 | scf_000001_A_fumigatus_A1163:1,115,922..1,116,899(+) |
| **AFUB_074880** | 1306.843988 | 2.832212151 | 0.09136312 | 30.99951233 | 5.47E-211 | 9.59E-209 | | AFUB_074880 | Has domain(s) with predicted oxidoreducta | Oxidoreductase FAD/NAD(P)-t | Oxidoreductase FAD/NAD(P | protein coding gene | GO:0005739 | GO:0003824;G( | metabolic proce | mitochondrion | catalytic activity;oxidoredu | 3141 | 1046 | | 284 | 0 | OG6_111529 | PF00175;PF006 | B0Y7T0 | IPR001433;IPR( | 1.6.2.2 (Cytochr | 0 | scf_000006_A_fumigatus_A1163:782,132..785,272(-) |
| **AFUB_034330** | 5.62583475 | 2.833288281 | 0.931128459 | 3.04285435 | 0.002343457 | 0.003940247 | | AFUB_034330 | Protein of unknown function | N/A | N/A | protein coding gene | | | | | | 948 | 315 | MTPLVKVGSLL | 33 | 1 | OG6_186772 | N/A | B0XZT7 | N/A | | 1 | scf_000003_A_fumigatus_A1163:121,626..122,573(-) |
| **AFUB_099610** | 101.7963065 | 2.838823242 | 0.232244164 | 12.22344275 | 2.33E-34 | 1.82E-33 | | AFUB_099610 | Protein of unknown function | N/A | N/A | protein coding gene | | | | | | 1191 | 376 | MKSIFALTGLLS | 45 | 1 | OG6_153083 | N/A | B0YEL6 | N/A | | 0 | scf_000010_A_fumigatus_A1163:392,881..394,071(-) |
| **AFUB_022400** | 78.88145338 | 2.838880265 | 0.257956168 | 11.00528158 | 3.60E-28 | 2.29E-27 | | AFUB_022400 | Protein of unknown function | N/A | N/A | protein coding gene | | | | | | 792 | 122 | | 1 | 0 | OG6_455060 | N/A | B0XVI8 | N/A | | 0 | scf_000002_A_fumigatus_A1163:1,482,170..1,482,961(+) |
| **AFUB_038600** | 8.230549503 | 2.839480877 | 0.786506162 | 3.610246192 | 0.000305907 | 0.00056578 | | AFUB_038600 | Has domain(s) with predicted sequence-s| | Zn(2)-C6 fungal-type DNA-binc | Zn(2)-C6 fungal-type DNA-b | protein coding gene | GO:0042025;G( | GO:0003677;G( | regulation of tra | host cell nucleus | DNA binding;DNA-binding | 363 | 120 | | 101 | 2 | OG6_166352 | PF00172 | B0XXU5 | IPR001138;IPR036864 | | 0 | scf_000003_A_fumigatus_A1163:1,339,000..1,339,362(+) |
| **AFUB_077580** | 207.1261686 | 2.840242736 | 0.177417024 | 16.00885123 | 1.11E-57 | 1.64E-56 | | AFUB_077580 | Protein of unknown function | N/A | N/A | protein coding gene | GO:0016021 | | | integral component of membrane | | 1427 | 387 | | 700 | 5 | OG6_129931 | N/A | B0Y8J7 | N/A | | 6 | scf_000006_A_fumigatus_A1163:1,557,728..1,559,154(+) |
| **AFUB_070830** | 5.679381773 | 2.848681797 | 0.943477435 | 3.019342796 | 0.002533237 | 0.004246003 | pgaA | AFUB_070830 | Ortholog(s) have polygalacturonase activi | Glycoside hydrolase, family 28 | Glycoside hydrolase, family : | protein coding gene | | GO:0004650 | carbohydrate metabolic process; | | polygalacturonase activity | 1238 | 368 | MRSVKLFGLAA | 988 | 5 | OG6_102442 | PF00295 | B0Y4Z4 | IPR000743;IPR( | 3.2.1.15 (Endo-| | 0 | scf_000005_A_fumigatus_A1163:2,194,877..2,196,114(+) |
| **AFUB_030950** | 12.69899503 | 2.849119508 | 0.623489858 | 4.569632482 | 4.89E-06 | 1.07E-05 | | AFUB_030950 | Protein of unknown function | N/A | N/A | protein coding gene | | | | | | 1677 | 297 | | 278 | 0 | OG6_152984 | N/A | | N/A | | 0 | scf_000002_A_fumigatus_A1163:4,023,438..4,025,114(+) |
| **AFUB_047660** | 85.44373373 | 2.849900306 | 0.250896589 | 11.35886429 | 6.70E-30 | 4.52E-29 | | AFUB_047660 | Has domain(s) with predicted N-acetyltran | GNAT domain | GNAT domain;Acyl-CoA N-a | protein coding gene | | GO:0008080 | | | N-acetyltransferase activi | 468 | 155 | | 14 | 0 | OG6_129405 | PF00583 | B0XX42 | IPR000182;IPR( | 2.3.1.- (Transfer | 0 | scf_000003_A_fumigatus_A1163:3,947,092..3,947,559(-) |
| **AFUB_015380** | 55.85932254 | 2.851730678 | 0.307210286 | 9.282666644 | 1.65E-20 | 7.83E-20 | | AFUB_015380 | Has domain(s) with predicted zinc ion binc | Transcription factor domain, fu | Transcription factor domain, | protein coding gene | GO:0005634 | GO:0003677;G( | transcription, DN | nucleus | DNA binding;zinc ion bind | 2299 | 727 | | 387 | 5 | OG6_133375 | PF04082 | B0XNJ6 | IPR007219 | 1.1.1.85 (3-isop | 0 | scf_000001_A_fumigatus_A1163:4,353,186..4,355,484(-) |
| **AFUB_080070** | 8.524170846 | 2.854021761 | 0.755661853 | 3.776850389 | 0.000158824 | 0.000301885 | | AFUB_080070 | Has domain(s) with predicted sequence-s| | Heat shock factor (HSF)-type, I | Heat shock factor (HSF)-type | protein coding gene | GO:0042025;G( | GO:0003700;G( | regulation of tra | host cell nucleus | DNA-binding transcription | 852 | 225 | | 0 | 0 | OG6_119133 | PF00447 | B0Y979 | IPR000232;IPR036390 | | 0 | scf_000007_A_fumigatus_A1163:33,810..34,661(-) |
| **AFUB_029640** | 354.1766752 | 2.857647831 | 0.144161611 | 19.82252984 | 1.90E-87 | 5.51E-86 | | AFUB_029640 | Protein of unknown function | N/A | N/A | protein coding gene | | | | | | 1956 | 651 | | 78 | 0 | OG6_454145 | N/A | B0XTR2 | N/A | | 0 | scf_000002_A_fumigatus_A1163:3,639,147..3,641,102(+) |
| **AFUB_088790** | 54.30714578 | 2.85914602 | 0.306582848 | 9.32585121 | 1.10E-20 | 5.26E-20 | | AFUB_088790 | Protein of unknown function | N/A | N/A | protein coding gene | | | | | | 2047 | 514 | | 88 | 0 | OG6_454888 | N/A | B0YC38 | N/A | | 0 | scf_000008_A_fumigatus_A1163:600,465..602,511(+) |
| **AFUB_043980** | 71.46001204 | 2.859777997 | 0.272107248 | 10.50974578 | 7.79E-26 | 4.53E-25 | | AFUB_043980 | Has domain(s) with predicted monooxyge | Cytochrome P450 | Cytochrome P450;Cytochror | protein coding gene | | GO:0009055;G( | obsolete oxidation-reduction pro | | electron transfer activity;h | 1793 | 561 | | 3790 | 17 | OG6_100014 | PF00067 | B0XZD0 | IPR001128;IPR( | 1.14.-.- (Acting ( | 0 | scf_000003_A_fumigatus_A1163:2,897,528..2,899,320(-) |
| **AFUB_016300** | 7.179616459 | 2.869847689 | 0.81801751 | 3.50829616 | 0.000450987 | 0.000821184 | | AFUB_016300 | Has domain(s) with predicted role in transı | Major facilitator superfamily | Tetracycline resistance prote | protein coding gene | GO:0016021;G( | GO:0022857 | gliotoxin biosynt | integral compon | transmembrane transporte | 1823 | 561 | | 316 | 0 | OG6_152905 | PF07690 | B0XP70 | IPR001411;IPR011701;IPR0362 | | 12 | scf_000001_A_fumigatus_A1163:4,629,820..4,631,642(+) |
| **AFUB_023950** | 46.06942715 | 2.870061279 | 0.332619522 | 8.628661558 | 6.21E-18 | 2.65E-17 | | AFUB_023950 | Has domain(s) with predicted DNA binding | N/A | Homeobox-like domain supe | protein coding gene | | GO:0003677 | | | DNA binding | 339 | 112 | | 86 | 0 | OG6_166237 | N/A | B0XW87 | IPR009057 | | 0 | scf_000002_A_fumigatus_A1163:2,015,464..2,015,802(+) |
| **AFUB_069450** | 450.2480363 | 2.872732458 | 0.133213894 | 21.56481107 | 3.84E-103 | 1.50E-101 | | AFUB_069450 | Protein of unknown function | N/A | N/A | protein coding gene | | | | | | 769 | 99 | | 15 | 0 | OG6_454783 | N/A | B0Y4K9 | N/A | | 0 | scf_000005_A_fumigatus_A1163:1,844,298..1,845,066(+) |
| **AFUB_022210** | 77.54298021 | 2.876825861 | 0.262338696 | 10.96607518 | 5.56E-28 | 3.51E-27 | | AFUB_022210 | Has domain(s) with predicted nucleic acid | DNA2/NAM7 helicase, AAA do | Zinc finger, CCCH-type;Zinc | protein coding gene | GO:0005634 | GO:0003700;G( | regulation of tra | nucleus | DNA-binding transcription | 5798 | 1914 | | 549 | 1 | OG6_101077 | PF13086;PF13( | B0XVG9 | IPR000571;IPR( | 3.6.4.13 (RNA h | 0 | scf_000002_A_fumigatus_A1163:1,423,664..1,429,461(+) |
| **AFUB_049700** | 5.873588734 | 2.8811412 | 0.929890584 | 3.098365818 | 0.001945911 | 0.003307019 | | AFUB_049700 | Has domain(s) with predicted role in respc | RTA-like protein | RTA-like protein | protein coding gene | GO:0016021 | | response to stre | integral component of membrane | | 1051 | 328 | | 857 | 1 | OG6_112506 | PF04479 | B0Y1L3 | IPR007568 | | 7 | scf_000004_A_fumigatus_A1163:234,976..236,026(+) |
| **AFUB_080830** | 11.67074073 | 2.88422505 | 0.65031229 | 4.435138459 | 9.20E-06 | 1.97E-05 | | AFUB_080830 | Protein of unknown function | N/A | N/A | protein coding gene | | | | | | 1906 | 519 | | 83 | 0 | OG6_454805 | N/A | B0Y9F3 | N/A | | 0 | scf_000007_A_fumigatus_A1163:318,578..320,483(+) |
| **AFUB_008720** | 45.01097401 | 2.884321954 | 0.337006692 | 8.558648903 | 1.14E-17 | 4.84E-17 | | AFUB_008720 | Protein of unknown function | N/A | N/A | protein coding gene | | | | | | 1204 | 381 | | 167 | 0 | OG6_186600 | N/A | B0XQ52 | N/A | | 0 | scf_000001_A_fumigatus_A1163:2,484,535..2,485,738(-) |
| **AFUB_101640** | 2.911866266 | 2.886072475 | 1.29466942 | 2.22919645 | 0.025800837 | 0.037617924 | | AFUB_101640 | Ortholog(s) have intracellular localization | Dienelactone hydrolase | Dienelactone hydrolase;Alph | protein coding gene | GO:0005622 | GO:0016787 | | intracellular ana | hydrolase activity | 832 | 233 | | 961 | 3 | OG6_102257 | PF01738 | B0YF58 | IPR002925;IPR( | 3.1.1.- (Carboxy | 0 | scf_000011_A_fumigatus_A1163:333,087..333,918(+) |
| **AFUB_044740** | 13.28034042 | 2.892971601 | 0.624205301 | 4.634647601 | 3.58E-06 | 7.88E-06 | | AFUB_044740 | Protein of unknown function | N/A | N/A | protein coding gene | | | | | | 420 | 131 | | 0 | 0 | OG6r21_23991( | N/A | B0XZP2 | N/A | | 0 | scf_000003_A_fumigatus_A1163:3,177,517..3,177,936(+) |
| **AFUB_020630** | 87.65746273 | 2.900323763 | 0.247729657 | 11.70761629 | 1.17E-31 | 8.37E-31 | | AFUB_020630 | Ortholog(s) have role in secondary metab( | Major facilitator superfamily | Major facilitator superfamily;l | protein coding gene | GO:0016021;G( | GO:0022857 | cellular respons | integral compon | transmembrane transporte | 1637 | 520 | | 2612 | 9 | OG6_100246 | PF07690 | B0XUM8 | IPR011701;IPR( | 2.8.2.11 (Galact | 12 | scf_000002_A_fumigatus_A1163:930,791..932,427(+) |
| **AFUB_038260** | 8.651881668 | 2.901908424 | 0.763643812 | 3.80008111 | 0.000144649 | 0.000275952 | | AFUB_038260 | Protein of unknown function | N/A | N/A | protein coding gene | | | | | | 789 | 262 | | 1 | 0 | OG6_454017 | N/A | B0XXQ8 | N/A | | 0 | scf_000003_A_fumigatus_A1163:1,239,947..1,240,735(-) |
| **AFUB_041310** | 11.71616307 | 2.901975025 | 0.6494279 | 4.468509935 | 7.88E-06 | 1.70E-05 | | AFUB_041310 | Protein of unknown function | N/A | N/A | protein coding gene | GO:0016021 | | | integral component of membrane | | 1283 | 380 | | 39 | 0 | OG6_454471 | N/A | B0XYL3 | N/A | | 1 | scf_000003_A_fumigatus_A1163:2,082,150..2,083,432(+) |
| **AFUB_004800** | 10.27179665 | 2.904839679 | 0.698253098 | 4.160152937 | 3.18E-05 | 6.48E-05 | | AFUB_004800 | Protein of unknown function | N/A | N/A | protein coding gene | | | | | | 546 | 127 | | 1 | 0 | OG6_454159 | N/A | B0XNM4 | N/A | | 0 | scf_000001_A_fumigatus_A1163:1,344,491..1,345,036(+) |
| **AFUB_050990** | 113.0166831 | 2.905654724 | 0.220215099 | 13.1946208 | 9.42E-40 | 8.67E-39 | | AFUB_050990 | Protein of unknown function | Arylsulfotransferase-like | Arylsulfotransferase-like | protein coding gene | | | | | | 1935 | 624 | MMLRDLWFAV | 664 | 5 | OG6_103894 | PF14269 | | IPR039535 | 2.8.2.22 (Aryl-su | 1 | scf_000004_A_fumigatus_A1163:568,874..570,808(+) |
| **AFUB_049850** | 5.872841521 | 2.907017512 | 0.915616437 | 3.174929364 | 0.00149873 | 0.002579807 | plb2 | AFUB_049850 | Has domain(s) with predicted phospholipa | Lysophospholipase, catalytic d | Lysophospholipase, catalytic | protein coding gene | | GO:0004622;G( | metabolic process;phospholipid ( | | lysophospholipase activity | 1965 | 588 | | 693 | 2 | OG6_108039 | PF01735 | B0Y1M7 | IPR002642;IPR( | 3.1.1.5 (Lysoph( | 0 | scf_000004_A_fumigatus_A1163:266,662..268,626(-) |
| **AFUB_003590** | 6.007893413 | 2.908058585 | 0.917920694 | 3.168093501 | 0.001534422 | 0.002637946 | | AFUB_003590 | unspecified product | N/A | N/A | ncRNA gene | | | | | | 97 | | | 0 | 0 | N/A (orthology r | N/A | | N/A | | | scf_000001_A_fumigatus_A1163:989,812..989,908(+) |
| **AFUB_028690** | 1324.514616 | 2.910238169 | 0.109804445 | 26.50382845 | 8.76E-155 | 8.19E-153 | | AFUB_028690 | Ortholog(s) have role in cellular response | Calcipressin | Calcipressin;RNA-binding dc | protein coding gene | GO:0005737 | GO:0003676 | calcium ion hom | cytoplasm | nucleic acid binding | 867 | 288 | | 232 | 0 | OG6_103325 | PF04847 | B0XTA0 | IPR006931;IPR( | 2.7.11.- (Protein | 0 | scf_000002_A_fumigatus_A1163:3,327,994..3,328,860(+) |
| **AFUB_030260** | 11.79022176 | 2.912721016 | 0.646001616 | 4.508844785 | 6.52E-06 | 1.41E-05 | | AFUB_030260 | unspecified product | N/A | N/A | ncRNA gene | | | | | | 113 | | | 0 | 0 | N/A (orthology r | N/A | | N/A | | | scf_000002_A_fumigatus_A1163:3,804,914..3,805,026(+) |
| **AFUB_091500** | 9391.901807 | 2.915066171 | 0.074591451 | 39.08043241 | 0 | 0 | | AFUB_091500 | Ortholog(s) have role in unsaturated fatty | Cytochrome b5-like heme/sterc | Cytochrome b5-like heme/st | protein coding gene | GO:0016021 | GO:0020037;G( | asexual sporula | integral compon | heme binding;metal ion bi | 1433 | 456 | | 585 | 0 | OG6_100468 | PF00173;PF004 | B0YB43 | IPR001199;IPR( | 1.13.11.47 (3-hy | 3 | scf_000008_A_fumigatus_A1163:1,453,091..1,454,523(+) |
| **AFUB_094850** | 88.30080094 | 2.917493607 | 0.249483284 | 11.69414462 | 1.37E-31 | 9.79E-31 | | AFUB_094850 | Has domain(s) with predicted oxidoreducta | FAD dependent oxidoreductas | FAD dependent oxidoreduct | protein coding gene | | GO:0016491 | obsolete oxidation-reduction pro | | oxidoreductase activity | 1562 | 497 | | 9 | 0 | OG6_454857 | PF01266 | B0YDA4 | IPR006076;IPR( | 1.5.3.- (With oxy | 1 | scf_000009_A_fumigatus_A1163:564,907..566,468(+) |
| **AFUB_083460** | 104.9099621 | 2.918717646 | 0.230362659 | 12.6700988 | 8.66E-37 | 7.30E-36 | nagA | AFUB_083460 | Has domain(s) with predicted N-acetylgluc | Amidohydrolase-related | N-acetylglucosamine-6-phos | protein coding gene | | GO:0047419;G( | N-acetylglucosamine metabolic p | | N-acetylgalactosamine-6- | 2135 | 430 | | 283 | 0 | OG6_102909 | PF01979 | B0YA54 | IPR003764;IPR( | 3.5.1.25 (N-acet | 0 | scf_000007_A_fumigatus_A1163:1,041,226..1,043,360(+) |
| **AFUB_047880** | 159.7216778 | 2.920067397 | 0.191189455 | 15.27316135 | 1.15E-52 | 1.52E-51 | | AFUB_047880 | Has domain(s) with predicted catalytic, mc | Molybdenum cofactor sulfurase | Molybdenum cofactor sulfura | protein coding gene | | GO:0003824;GO:0030151;GO:0030170 | | | catalytic activity;molybder | 848 | 200 | | 69 | 0 | OG6_122945 | PF03473 | B0XX63 | IPR005302;IPR011037 | | 0 | scf_000003_A_fumigatus_A1163:4,007,916..4,008,763(-) |
| **AFUB_092380** | 14.89317975 | 2.931381635 | 0.583038142 | 5.027769927 | 4.96E-07 | 1.16E-06 | | AFUB_092380 | Has domain(s) with predicted carbon-nitro | Amidase signature domain | Amidase;Amidase signature | protein coding gene | | GO:0004040;GO:0016884;GO:0016787;GO:001( | | | amidase activity;carbon-n | 2130 | 553 | | 231 | 0 | OG6_166595 | PF01425 | B0YBD1 | IPR000120;IPR( | 6.3.5.7 (Glutami | 0 | scf_000008_A_fumigatus_A1163:1,657,453..1,659,582(-) |
| **AFUB_046220** | 3.048538973 | 2.932244487 | 1.285184341 | 2.281575019 | 0.022514443 | 0.033162515 | | AFUB_046220 | Ortholog(s) have role in secondary metab( | Major facilitator superfamily | Major facilitator superfamily;l | protein coding gene | GO:0016021;G( | GO:0022857 | cellular respons | integral compon | transmembrane transporte | 1516 | 449 | | 118 | 0 | OG6_112023 | PF07690 | B0Y0H6 | IPR011701;IPR036259 | | 10 | scf_000003_A_fumigatus_A1163:3,586,129..3,587,644(+) |
| **AFUB_015920** | 247.3743776 | 2.939238889 | 0.167056923 | 17.59423573 | 2.73E-69 | 5.23E-68 | | AFUB_015920 | Protein of unknown function | MJ1316 RNA cyclic group end | MJ1316 RNA cyclic group er | protein coding gene | | | | | | 636 | 153 | | 55 | 0 | OG6_110219 | PF04457 | B0XNW7 | IPR040459 | | 0 | scf_000001_A_fumigatus_A1163:4,500,533..4,501,168(-) |
| **AFUB_098300** | 22.37472694 | 2.941293545 | 0.48521887 | 6.061787222 | 1.35E-09 | 3.75E-09 | | AFUB_098300 | Protein of unknown function | N/A | N/A | protein coding gene | | GO:0005515 | | | protein binding | 837 | 278 | | 61 | 0 | OG6_455407 | N/A | B0YE86 | N/A | | 0 | scf_000010_A_fumigatus_A1163:49,347..50,183(+) |
| **AFUB_090690** | 19.4984047 | 2.941643056 | 0.517240535 | 5.687185858 | 1.29E-08 | 3.41E-08 | | AFUB_090690 | Has domain(s) with predicted protein bindi | BTB/POZ domain | BTB/POZ domain;SKP1/BTE | protein coding gene | | GO:0005515 | | | protein binding | 801 | 266 | | 653 | 7 | OG6_121403 | PF00651 | B0YCM2 | IPR000210;IPR011333 | | 0 | scf_000008_A_fumigatus_A1163:1,208,529..1,209,329(+) |
| **AFUB_025000** | 51.39788944 | 2.943729687 | 0.327299582 | 8.993991582 | 2.38E-19 | 1.08E-18 | | AFUB_025000 | Has domain(s) with predicted protein bindi | C-5 cytosine methyltransferase | SNF2-related, N-terminal do | protein coding gene | GO:0005634 | GO:0005524;G( | DNA methylatio | nucleus | ATP binding;DNA binding | 6568 | 2115 | | 178 | 0 | OG6_129939 | PF00145;PF001 | B0XRV8 | IPR000330;IPR( | 2.1.1.37 (DNA (( | 0 | scf_000002_A_fumigatus_A1163:2,319,506..2,326,073(+) |
| **AFUB_012600** | 88.76127139 | 2.947194928 | 0.250878514 | 11.74749832 | 7.27E-32 | 5.28E-31 | | AFUB_012600 | Protein of unknown function | N/A | N/A | protein coding gene | | | | | | 640 | 143 | | 1 | 0 | OG6_454307 | N/A | B0XR98 | N/A | | 0 | scf_000001_A_fumigatus_A1163:3,551,174..3,551,813(+) |
| **AFUB_084560** | 415.7817791 | 2.949331454 | 0.141560913 | 20.83436304 | 2.11E-96 | 7.32E-95 | | AFUB_084560 | Protein of unknown function | Mannosyltransferase 1, CMT1 | Mannosyltransferase 1, CMT | protein coding gene | | | | | | 1822 | 585 | | 202 | 1 | OG6_152934 | PF11735 | B0YAG2 | IPR021047 | | 0 | scf_000007_A_fumigatus_A1163:1,409,263..1,411,084(+) |
| **AFUB_033480** | 51.31474013 | 2.950057825 | 0.323726279 | 9.112815422 | 8.03E-20 | 3.72E-19 | | AFUB_033480 | Has domain(s) with predicted role in respc | RTA-like protein | RTA-like protein | protein coding gene | GO:0016021 | | response to stre | integral component of membrane | | 1230 | 360 | | 252 | 1 | OG6_166199 | PF04479 | B0XVY9 | IPR007568 | | 6 | scf_000002_A_fumigatus_A1163:4,692,992..4,694,221(+) |
| **AFUB_092280** | 74.15178552 | 2.954832541 | 0.272392187 | 10.84771401 | 2.04E-27 | 1.26E-26 | | AFUB_092280 | Has domain(s) with predicted transition m( | Rieske [2Fe-2S] iron-sulphur d | Protein of unknown function | protein coding gene | | GO:0051537;G( | obsolete oxidation-reduction pro | | 2 iron, 2 sulfur cluster bin | 1664 | 451 | | 130 | 0 | OG6_109515 | PF00355;PF043 | B0YBC1 | IPR007402;IPR009078;IPR0179 | | 0 | scf_000008_A_fumigatus_A1163:1,635,214..1,636,877(+) |
| **AFUB_026340** | 633.9437796 | 2.958065288 | 0.132718589 | 22.28825143 | 4.80E-110 | 2.24E-108 | | AFUB_026340 | Has domain(s) with predicted nucleic acid | N/A | Zinc finger C2H2-type;Zinc fi | protein coding gene | | GO:0003676 | | | nucleic acid binding | 3721 | 529 | | 82 | 0 | OG6_454401 | N/A | | IPR013087;IPR036236 | | 0 | scf_000002_A_fumigatus_A1163:2,672,124..2,675,844(-) |
| **AFUB_000140** | 16.49751384 | 2.958081145 | 0.555945394 | 5.320812398 | 1.03E-07 | 2.55E-07 | | AFUB_000140 | Protein of unknown function | Haloacid dehalogenase-like hy | HAD-like superfamily;Haloac | protein coding gene | | | | | | 714 | 237 | | 244 | 1 | OG6_129404 | PF13419 | B0XMG2 | IPR036412;IPR( | 3.8.1.2 ((S)-2-ha | 0 | scf_000001_A_fumigatus_A1163:33,460..34,173(-) |
| **AFUB_011940** | 27.4314487 | 2.959393416 | 0.432752589 | 6.838534288 | 8.00E-12 | 2.56E-11 | | AFUB_011940 | Protein of unknown function | N/A | N/A | protein coding gene | | | | | | 438 | 145 | | 20 | 0 | OG6_455241 | N/A | B0XR19 | N/A | | 0 | scf_000001_A_fumigatus_A1163:3,352,621..3,353,058(-) |
| **AFUB_047540** | 21.21840679 | 2.96292874 | 0.492421657 | 6.017056108 | 1.78E-09 | 4.91E-09 | | AFUB_047540 | Ortholog(s) have orotidine-5-prime -phosp | Orotidine 5'-phosphate decarb( | Orotidine 5'-phosphate deca | protein coding gene | | GO:0003824;G( | de novo' UMP biosynthetic proce | | catalytic activity;orotidine- | 1116 | 277 | | 572 | 2 | OG6_100519 | PF00215 | B0XX31 | IPR001754;IPR( | 4.1.1.23 (Orotidi | 0 | scf_000003_A_fumigatus_A1163:3,919,781..3,920,896(+) |
| **AFUB_084640** | 74.77607427 | 2.968367138 | 0.273591745 | 10.84962246 | 2.00E-27 | 1.24E-26 | pgaB | AFUB_084640 | Ortholog(s) have polygalacturonase activi | Glycoside hydrolase, family 28 | Glycoside hydrolase, family : | protein coding gene | | GO:0004650 | carbohydrate metabolic process; | | polygalacturonase activity | 1149 | 364 | MHFFQSSLVAA | 988 | 5 | OG6_102442 | PF00295 | B0YAH0 | IPR000743;IPR( | 3.2.1.15 (Endo-| | 0 | scf_000007_A_fumigatus_A1163:1,439,070..1,440,218(-) |
| **AFUB_030400** | 507.2924254 | 2.972354637 | 0.12404668 | 23.96158159 | 7.00E-127 | 4.27E-125 | | AFUB_030400 | Has domain(s) with predicted ATP binding | ATPase, AAA-type, core;BCS1 | AAA+ ATPase domain;ATPa | protein coding gene | | GO:0005524;G( | mitochondrial respiratory chain c | | ATP binding;nucleoside-tr | 2299 | 579 | | 795 | 2 | OG6_101424 | PF00004;PF08740 | | IPR003593;IPR( | 2.7.11.1 (Non-sp | 0 | scf_000002_A_fumigatus_A1163:3,861,889..3,864,187(-) |
| **AFUB_074450** | 4.706408833 | 2.972735156 | 1.054869395 | 2.818107312 | 0.004830766 | 0.007805536 | | AFUB_074450 | Has domain(s) with predicted magnesium | Inorganic pyrophosphatase | Inorganic pyrophosphatase;I | protein coding gene | GO:0005737 | GO:0004427;G( | phosphate-conta | cytoplasm | inorganic diphosphatase a | 1211 | 321 | | 458 | 1 | OG6_100346 | PF00719 | B0Y7N7 | IPR008162;IPR( | 3.6.1.1 (Inorgan | 0 | scf_000006_A_fumigatus_A1163:652,665..653,875(+) |
| **AFUB_049360** | 202.6273325 | 2.979505622 | 0.175999095 | 16.92909625 | 2.75E-64 | 4.69E-63 | | AFUB_049360 | Ortholog(s) have intracellular localization | Beta-lactamase-related | Beta-lactamase-related;Beta | protein coding gene | GO:0005622 | | | intracellular anatomical structure | | 1227 | 408 | | 674 | 3 | OG6_104484 | PF00144 | B0Y1H9 | IPR001466;IPR( | 2.3.1.238 (Mona | 0 | scf_000004_A_fumigatus_A1163:167,851..169,077(+) |
| **AFUB_080810** | 19.88890793 | 2.981367209 | 0.506592491 | 5.885138963 | 3.98E-09 | 1.08E-08 | | AFUB_080810 | Has domain(s) with predicted lysozyme, c | Glycoside hydrolase, family 25 | Glycoside hydrolase, family : | protein coding gene | | GO:0003824;G( | carbohydrate metabolic process; | | catalytic activity;lysozyme | 1478 | 309 | MKFSIVAIATIA( | 77 | 1 | OG6_119717 | PF01183 | B0Y9F1 | IPR002053;IPR( | 3.2.1.17 (Lysozy | 0 | scf_000007_A_fumigatus_A1163:312,758..314,235(+) |
| **AFUB_000150** | 11.05102819 | 2.983596263 | 0.712962375 | 4.184787819 | 2.85E-05 | 5.84E-05 | | AFUB_000150 | Has domain(s) with predicted 4-aminobuty | Aminotransferase class-III | 4-aminobutyrate aminotransf | protein coding gene | | GO:0003867;G( | gamma-aminobutyric acid metab | | 4-aminobutyrate transami | 1578 | 467 | | 372 | 1 | OG6_103520 | PF00202 | B0XMG3 | IPR004631;IPR( | 2.6.1.19 (4-amir | 0 | scf_000001_A_fumigatus_A1163:34,544..36,121(+) |
| **AFUB_071570** | 10.88732028 | 2.987544468 | 0.683804423 | 4.369004306 | 1.25E-05 | 2.64E-05 | | AFUB_071570 | Has domain(s) with predicted sequence-s| | Zn(2)-C6 fungal-type DNA-binc | Zn(2)-C6 fungal-type DNA-b | protein coding gene | GO:0042025;G( | GO:0003677;G( | regulation of tra | host cell nucleus | DNA binding;DNA-binding | 1185 | 373 | | 178 | 1 | OG6_153139 | PF00172;PF11¢ | B0Y568 | IPR001138;IPR021858;IPR0368 | | 0 | scf_000005_A_fumigatus_A1163:2,422,001..2,423,185(-) |
| **AFUB_028460** | 74.5760328 | 2.998246174 | 0.274673913 | 10.91565683 | 9.70E-28 | 6.06E-27 | | AFUB_028460 | Has domain(s) with predicted transferase | UDP-glucuronosyl/UDP-glucos | UDP-glucuronosyl/UDP-gluc | protein coding gene | GO:0016021 | GO:0008194;G( | metabolic proce | integral compon | UDP-glycosyltransferase | 1628 | 523 | | 326 | 0 | OG6_100052 | PF00201 | B0XT08 | IPR002213 | 1.2.1.84 (Alcohc | 1 | scf_000002_A_fumigatus_A1163:3,269,109..3,270,736(+) |
| **AFUB_060420** | 7.838816467 | 3.003939053 | 0.804658138 | 3.733186693 | 0.000189072 | 0.000357025 | | AFUB_060420 | Protein of unknown function | N/A | N/A | protein coding gene | | | | | | 1601 | 488 | | 128 | 3 | OG6_166300 | N/A | B0Y1Q3 | N/A | | 0 | scf_000004_A_fumigatus_A1163:3,253,856..3,255,456(-) |
| **AFUB_097590** | 6.262507915 | 3.005203155 | 0.918215153 | 3.272874712 | 0.001064597 | 0.001862126 | | AFUB_097590 | Has domain(s) with predicted zinc ion binc | N/A | Zinc finger C2H2-type | protein coding gene | | GO:0008270 | | | zinc ion binding | 1374 | 402 | | 1 | 0 | OG6r21_16210 | N/A | B0YE24 | IPR013087 | | 0 | scf_000009_A_fumigatus_A1163:1,384,674..1,386,047(+) |
| **AFUB_096320** | 303.9016582 | 3.009864193 | 0.32408914 | 9.287149186 | 1.58E-20 | 7.52E-20 | | AFUB_096320 | Has domain(s) with predicted protein kina | Cyclin PHO80-like | Cyclin PHO80-like;Cyclin-lik | protein coding gene | | GO:0019901 | regulation of cyclin-dependent p | | protein kinase binding | 2172 | 702 | | 194 | 0 | OG6_121405 | PF08613 | B0YDP6 | IPR013922;IPR036915 | | 0 | scf_000009_A_fumigatus_A1163:985,684..987,855(+) |
| **AFUB_000390** | 162.0606109 | 3.011367781 | 0.194213595 | 15.50544274 | 3.19E-54 | 4.37E-53 | | AFUB_000390 | Has domain(s) with predicted iron ion bind | Cytochrome P450 | Cytochrome P450;Cytochror | protein coding gene | | GO:0009055;G( | obsolete oxidation-reduction pro | | electron transfer activity;h | 1485 | 461 | | 38 | 0 | OG6_455278 | PF00067 | | IPR001128;IPR( | 1.14.-.- (Acting ( | 1 | scf_000001_A_fumigatus_A1163:111,906..113,390(-) |
| **AFUB_101260** | 6.237476262 | 3.017476833 | 0.915152389 | 3.29723975 | 0.000976401 | 0.001714708 | | AFUB_101260 | Protein of unknown function | N/A | N/A | protein coding gene | | | | | | 761 | 245 | | 57 | 0 | OG6_454215 | N/A | B0YF18 | N/A | | 0 | scf_000011_A_fumigatus_A1163:218,035..218,795(+) |
| **AFUB_053180** | 119.8109001 | 3.024447059 | 0.218632984 | 13.83344361 | 1.60E-43 | 1.66E-42 | | AFUB_053180 | Has domain(s) with predicted role in respc | RTA-like protein | RTA-like protein | protein coding gene | GO:0016021 | | response to stre | integral component of membrane | | 1256 | 279 | | 13 | 0 | OG6_453711 | PF04479 | B0Y373 | IPR007568 | | 5 | scf_000004_A_fumigatus_A1163:1,221,879..1,223,134(-) |
| **AFUB_057820** | 60.08642848 | 3.024809379 | 0.302987556 | 9.983279232 | 1.80E-23 | 9.59E-23 | | AFUB_057820 | Has domain(s) with predicted transferase | Alcohol dehydrogenase, C-term | GroES-like superfamily;Alcol | protein coding gene | | GO:0016747;G( | obsolete oxidation-reduction pro | | acyltransferase activity, tr | 1239 | 350 | | 1153 | 3 | OG6_110113 | PF00107;PF082 | B0Y0M3 | IPR011032;IPR( | 1.1.1.- (With NA | 0 | scf_000004_A_fumigatus_A1163:2,519,752..2,520,990(+) |
| **AFUB_092550** | 52.25860786 | 3.024959643 | 0.329905667 | 9.169165462 | 4.77E-20 | 2.22E-19 | | AFUB_092550 | Has domain(s) with predicted catalytic acti | Fumarylacetoacetase-like, C-te | Fumarylacetoacetase-like, C | protein coding gene | | GO:0003824;G( | metabolic process | | catalytic activity;hydrolase | 1154 | 249 | | 1260 | 4 | OG6_100403 | PF01557 | B0YBE7 | IPR011234;IPR( | 4.1.1.68 (5-oxop | 0 | scf_000008_A_fumigatus_A1163:1,701,416..1,702,569(-) |
| **AFUB_084860** | 374.4110484 | 3.027338737 | 0.136611587 | 22.16019012 | 8.32E-109 | 3.69E-107 | | AFUB_084860 | Has domain(s) with predicted transferase | Glycosyl transferase, family 8 | Glycosyl transferase, family | protein coding gene | | GO:0016757 | | | glycosyltransferase activit | 1448 | 324 | | 387 | 1 | OG6_107247 | PF01501 | B0YAJ3 | IPR002495;IPR( | 2.4.1.- (Hexosyl | 0 | scf_000007_A_fumigatus_A1163:1,500,632..1,502,079(+) |
| **AFUB_082150** | 16.00089903 | 3.033925118 | 0.572783217 | 5.296812178 | 1.18E-07 | 2.90E-07 | | AFUB_082150 | Protein of unknown function | N/A | N/A | protein coding gene | | | | | | 4046 | 1219 | | 80 | 0 | OG6_454786 | N/A | B0Y9T0 | N/A | | 0 | scf_000007_A_fumigatus_A1163:677,887..681,932(+) |
| **AFUB_032700** | 3.168661732 | 3.034462194 | 1.270986126 | 2.387486482 | 0.016964027 | 0.02542625 | | AFUB_032700 | Protein of unknown function | N/A | N/A | protein coding gene | | | | | | 3011 | 951 | | 34 | 0 | OG6_453843 | N/A | B0XVD1 | N/A | | 0 | scf_000002_A_fumigatus_A1163:4,500,794..4,503,804(+) |
| **AFUB_033540** | 11.19025014 | 3.038090981 | 0.681699277 | 4.456643986 | 8.33E-06 | 1.79E-05 | | AFUB_033540 | Has domain(s) with predicted sequence-s| | Zn(2)-C6 fungal-type DNA-binc | Zn(2)-C6 fungal-type DNA-b | protein coding gene | GO:0042025;G( | GO:0003677;G( | regulation of tra | host cell nucleus | DNA binding;DNA-binding | 2268 | 667 | | 44 | 0 | OG6_166287 | PF00172;PF04082 | | IPR001138;IPR007219;IPR0368 | | 0 | scf_000002_A_fumigatus_A1163:4,705,072..4,707,339(+) |
| **AFUB_070960** | 22.32741917 | 3.04298614 | 0.489179858 | 6.220587561 | 4.95E-10 | 1.41E-09 | | AFUB_070960 | Has domain(s) with predicted zinc ion binc | Protein of unknown function DU | Protein of unknown function | protein coding gene | | | | | | 2803 | 773 | | 388 | 7 | OG6_153136 | PF11917 | B0Y506 | IPR021842 | | 0 | scf_000005_A_fumigatus_A1163:2,226,898..2,229,700(+) |
| **AFUB_001000** | 44.89869734 | 3.043593973 | 0.351995295 | 8.646689373 | 5.30E-18 | 2.27E-17 | | AFUB_001000 | Protein of unknown function | Sulfotransferase, S. mansonii-t | P-loop containing nucleoside | protein coding gene | GO:0016021 | | | integral component of membrane | | 955 | 302 | | 713 | 1 | OG6_103690 | PF17784 | B0XM57 | IPR027417;IPR040632 | | 1 | scf_000001_A_fumigatus_A1163:257,484..258,438(-) |
| **AFUB_091340** | 16.14886064 | 3.046480145 | 0.584015951 | 5.216433108 | 1.82E-07 | 4.43E-07 | | AFUB_091340 | Protein of unknown function | N/A | N/A | protein coding gene | | | | | | 799 | 222 | | 98 | 0 | OG6_453961 | N/A | B0YB27 | N/A | | 0 | scf_000008_A_fumigatus_A1163:1,415,320..1,416,118(+) |
| **AFUB_102340** | 43.45189237 | 3.055831526 | 0.352718938 | 8.66364462 | 4.57E-18 | 1.96E-17 | | AFUB_102340 | Protein of unknown function | N/A | N/A | protein coding gene | | | | | | 6194 | 2037 | | 439 | 2 | OG6_142064 | N/A | B0YFA9 | N/A | | 0 | scf_000014_A_fumigatus_A1163:3,657..9,850(-) |
| **AFUB_091470** | 337.1072039 | 3.062978451 | 0.148817818 | 20.58206805 | 3.97E-94 | 1.32E-92 | | AFUB_091470 | Protein of unknown function | DNA recombination and repair | DNA recombination and repa | protein coding gene | | | | | | 910 | 266 | | 177 | 0 | OG6_152708 | PF08423 | | IPR013632;IPR027417 | | 0 | scf_000008_A_fumigatus_A1163:1,445,141..1,446,050(-) |
| **AFUB_079210** | 32.56508904 | 3.069487897 | 0.412095144 | 7.448493246 | 9.44E-14 | 3.35E-13 | | AFUB_079210 | Has domain(s) with predicted protein bindi | Ankyrin repeat-containing doma | Ankyrin repeat;Ankyrin repea | protein coding gene | GO:0016021 | GO:0005515 | | integral compon | protein binding | 1057 | 306 | | 30 | 1 | OG6_186958 | PF12796 | B0Y907 | IPR002110;IPR020683;IPR0367 | | 1 | scf_000006_A_fumigatus_A1163:2,031,486..2,032,542(+) |
| **AFUB_046370** | 14.67288874 | 3.069717164 | 0.605949183 | 5.065964684 | 4.06E-07 | 9.58E-07 | | AFUB_046370 | Has domain(s) with predicted protein bindi | Ankyrin repeat-containing doma | Ankyrin repeat;Ankyrin repea | protein coding gene | | GO:0005515 | | | protein binding | 1630 | 284 | | 1 | 0 | OG6_455183 | PF12796 | B0Y0J0 | IPR002110;IPR020683;IPR0367 | | 0 | scf_000003_A_fumigatus_A1163:3,618,878..3,620,507(-) |
| **AFUB_047080** | 66.37911204 | 3.072204508 | 0.289582124 | 10.60909584 | 2.70E-26 | 1.59E-25 | | AFUB_047080 | Protein of unknown function | N/A | N/A | protein coding gene | | | | | | 1227 | 254 | | 52 | 0 | OG6_455206 | N/A | B0XWR8 | N/A | | 0 | scf_000003_A_fumigatus_A1163:3,796,122..3,797,348(+) |
| **AFUB_052910** | 84.84419607 | 3.073310597 | 0.260714103 | 11.78804891 | 4.50E-32 | 3.28E-31 | | AFUB_052910 | Protein of unknown function | N/A | N/A | protein coding gene | | | | | | 791 | 184 | | 115 | 2 | OG6_113596 | N/A | B0Y346 | N/A | | 0 | scf_000004_A_fumigatus_A1163:1,104,785..1,105,575(-) |
| **AFUB_087500** | 4.951107879 | 3.074007742 | 1.032578481 | 2.977020921 | 0.002910641 | 0.004832347 | | AFUB_087500 | Protein of unknown function | N/A | N/A | protein coding gene | GO:0016021 | | | integral component of membrane | | 1888 | 374 | | 94 | 0 | OG6_454501 | N/A | B0YBQ8 | N/A | | 6 | scf_000008_A_fumigatus_A1163:248,750..250,637(+) |
| **AFUB_081780** | 9.761089524 | 3.076081123 | 0.733838692 | 4.191767424 | 2.77E-05 | 5.67E-05 | | AFUB_081780 | Protein of unknown function | Fungal N-terminal domain of S | Fungal N-terminal domain of | protein coding gene | | | | | | 977 | 303 | | 80 | 0 | OG6_454269 | PF17111 | B0Y9P3 | IPR031348 | | 0 | scf_000007_A_fumigatus_A1163:560,754..561,730(+) |
| **AFUB_063280** | 2981.89421 | 3.078895767 | 0.083688349 | 36.79001671 | 2.67E-296 | 1.22E-293 | | AFUB_063280 | Has domain(s) with predicted branched-ch | Aminotransferase class IV | Aminotransferase class IV;B | protein coding gene | | GO:0004084;G( | branched-chain amino acid meta | | branched-chain-amino-ac | 2236 | 414 | | 978 | 3 | OG6_100313 | PF01063 | | IPR001544;IPR( | 2.6.1.42 (Branch | 0 | scf_000005_A_fumigatus_A1163:127,349..129,584(+) |
| **AFUB_032950** | 16.2126288 | 3.08382187 | 0.586227591 | 5.260451602 | 1.44E-07 | 3.51E-07 | | AFUB_032950 | Has domain(s) with predicted protein bindi | Glutathione S-transferase, N-te | Glutathione S-transferase, N | protein coding gene | | GO:0005515;GO:0016740 | | | protein binding;transferas | 871 | 256 | | 353 | 2 | OG6_109856 | PF02798;PF144 | B0XVM3 | IPR004045;IPR( | 2.5.1.18 (Glutatl | 0 | scf_000002_A_fumigatus_A1163:4,577,031..4,577,901(-) |
| **AFUB_038590** | 271.6253149 | 3.08567378 | 0.159663692 | 19.32608316 | 3.24E-83 | 8.42E-82 | | AFUB_038590 | Has domain(s) with predicted integral to m | AdipoR/Haemolysin-III-related | AdipoR/Haemolysin-III-relate | protein coding gene | GO:0016021 | | | integral component of membrane | | 966 | 321 | | 752 | 2 | OG6_100486 | PF03006 | B0XXU4 | IPR004254 | | 6 | scf_000003_A_fumigatus_A1163:1,336,985..1,337,950(+) |
| **AFUB_066630** | 24.66119893 | 3.086714319 | 0.467918201 | 6.596696419 | 4.20E-11 | 1.29E-10 | | AFUB_066630 | Protein of unknown function | N/A | Uncharacterized membrane | protein coding gene | | | | | | 1806 | 308 | | 97 | 0 | OG6_455162 | N/A | B0Y6E3 | IPR038781 | | 0 | scf_000005_A_fumigatus_A1163:1,042,334..1,044,139(-) |
| **AFUB_038990** | 14.68369329 | 3.087014993 | 0.61457543 | 5.023004246 | 5.09E-07 | 1.19E-06 | | AFUB_038990 | Protein of unknown function | N/A | N/A | protein coding gene | | | | | | 362 | 104 | MKTSTIVLSAVI | 2 | 0 | OG6_455203 | N/A | B0XXY4 | N/A | | 0 | scf_000003_A_fumigatus_A1163:1,441,283..1,441,644(+) |
| **AFUB_017960** | 8.112181175 | 3.087524316 | 0.813774267 | 3.794079563 | 0.000148192 | 0.000282321 | | AFUB_017960 | Has domain(s) with predicted sequence-s| | Zn(2)-C6 fungal-type DNA-binc | Zn(2)-C6 fungal-type DNA-b | protein coding gene | GO:0042025;G( | GO:0003677;G( | regulation of tra | host cell nucleus | DNA binding;DNA-binding | 1896 | 614 | | 938 | 1 | OG6_143641 | PF00172 | B0XT44 | IPR001138;IPR036864 | | 0 | scf_000002_A_fumigatus_A1163:191,085..192,980(+) |
| **AFUB_061950** | 88.81995422 | 3.100575848 | 0.257605158 | 12.03615592 | 2.29E-33 | 1.74E-32 | | AFUB_061950 | Protein of unknown function | DJ-1/PfpI | DJ-1/PfpI;Class I glutamine a | protein coding gene | | | | | | 681 | 226 | | 523 | 1 | OG6_106790 | PF01965 | B0Y2I6 | IPR002818;IPR029062 | | 0 | scf_000004_A_fumigatus_A1163:3,620,377..3,621,057(+) |
| **AFUB_013910** | 8.374081918 | 3.104763842 | 0.809893745 | 3.83354466 | 0.00012631 | 0.000243084 | | AFUB_013910 | Has domain(s) with predicted sequence-s| | Zn(2)-C6 fungal-type DNA-binc | Zn(2)-C6 fungal-type DNA-b | protein coding gene | GO:0042025;G( | GO:0003677;G( | regulation of tra | host cell nucleus | DNA binding;DNA-binding | 2401 | 738 | | 514 | 1 | OG6_129925 | PF00172;PF04( | B0XMR6 | IPR001138;IPR007219;IPR0368 | | 0 | scf_000001_A_fumigatus_A1163:3,912,118..3,914,518(+) |
| **AFUB_094660** | 440.4299568 | 3.106476364 | 0.130817258 | 23.7466861 | 1.19E-124 | 7.03E-123 | | AFUB_094660 | Ortholog(s) have role in biotin biosynthetic | Aminotransferase, class I/class | Aminotransferase, class I/cla | protein coding gene | GO:0005777 | GO:0003824;G( | biosynthetic pro | peroxisome | catalytic activity;pyridoxal | 1394 | 414 | | 595 | 1 | OG6_100359 | PF00155 | B0YD85 | IPR004839;IPR( | 2.1.2.1 (Glycine | 0 | scf_000009_A_fumigatus_A1163:508,023..509,416(+) |
| **AFUB_034230** | 9.955191002 | 3.10812374 | 0.736580701 | 4.219664916 | 2.45E-05 | 5.05E-05 | | AFUB_034230 | Protein of unknown function | Ion channel regulatory protein, | Ion channel regulatory protei | protein coding gene | GO:0016021 | | | integral component of membrane | | 1546 | 465 | | 411 | 0 | OG6_122838 | PF05978 | B0XZS7 | IPR010291;IPR036259 | | 11 | scf_000003_A_fumigatus_A1163:105,586..107,131(+) |
| **AFUB_017320** | 100.1906113 | 3.113692557 | 0.240932568 | 12.92350215 | 3.32E-38 | 2.91E-37 | | AFUB_017320 | Has domain(s) with predicted catalytic acti | N/A | Cyclic phosphodiesterase | protein coding gene | GO:0005622 | GO:0003824 | RNA metabolic | intracellular ana | catalytic activity | 1062 | 303 | | 174 | 1 | OG6_152908 | N/A | B0XSR4 | IPR009097 | | 0 | scf_000002_A_fumigatus_A1163:41,261..42,322(+) |
| **AFUB_094620** | 299.6756435 | 3.113999258 | 0.174817556 | 17.81285201 | 5.62E-71 | 1.12E-69 | | AFUB_094620 | unspecified product | N/A | N/A | ncRNA gene | | | | | | 136 | | | 0 | 0 | N/A (orthology r | N/A | | N/A | | | scf_000009_A_fumigatus_A1163:495,878..496,013(+) |
| **AFUB_054570** | 376.2655527 | 3.118156458 | 0.143220406 | 21.77173306 | 4.30E-105 | 1.79E-103 | | AFUB_054570 | Protein of unknown function | HPP | HPP | protein coding gene | GO:0016021 | | | integral component of membrane | | 1660 | 294 | | 269 | 0 | OG6_106377 | PF04982 | B0Y3K8 | IPR007065 | | 5 | scf_000004_A_fumigatus_A1163:1,636,342..1,638,001(+) |
| **AFUB_049390** | 60.60886381 | 3.121654645 | 0.308925343 | 10.10488364 | 5.26E-24 | 2.85E-23 | | AFUB_049390 | Has domain(s) with predicted sequence-s| | Zn(2)-C6 fungal-type DNA-binc | Zn(2)-C6 fungal-type DNA-b | protein coding gene | GO:0042025;G( | GO:0003677;G( | regulation of tra | host cell nucleus | DNA binding;DNA-binding | 1384 | 443 | | 225 | 1 | OG6_166473 | PF00172 | B0Y1I2 | IPR001138;IPR036864 | | 0 | scf_000004_A_fumigatus_A1163:172,506..173,889(+) |
| **AFUB_097290** | 33.81271557 | 3.123760527 | 0.414439189 | 7.537319371 | 4.80E-14 | 1.72E-13 | | AFUB_097290 | Has domain(s) with predicted nucleotide b | NmrA-like domain | NmrA-like domain;NAD(P)-b | protein coding gene | | | | | | 1003 | 297 | | 1226 | 3 | OG6_105106 | PF05368 | B0YDZ1 | IPR008030;IPR( | 1.23.1.1 ((+)-pin | 0 | scf_000009_A_fumigatus_A1163:1,268,474..1,269,476(-) |
| **AFUB_066200** | 400.8219241 | 3.124396151 | 0.137593 | 22.70752258 | 3.78E-114 | 1.88E-112 | | AFUB_066200 | Has domain(s) with predicted heat shock | | DPH-type metal-binding domai | DnaJ domain;DPH-type met | protein coding gene | | GO:0046872 | peptidyl-diphthamide biosyntheti | | metal ion binding | 1312 | 199 | | 276 | 1 | OG6_104505 | PF05207 | B0Y6A2 | IPR001623;IPR007872;IPR0366 | | 0 | scf_000005_A_fumigatus_A1163:928,084..929,395(-) |
| **AFUB_049320** | 31.83686961 | 3.124593043 | 0.414618354 | 7.536070257 | 4.84E-14 | 1.74E-13 | | AFUB_049320 | Protein of unknown function | N/A | N/A | protein coding gene | | | | | | 424 | 116 | | 4 | 0 | OG6_454400 | N/A | B0Y1B3 | N/A | | 0 | scf_000004_A_fumigatus_A1163:162,538..162,961(+) |
| **AFUB_102330** | 16.84121462 | 3.126811873 | 0.571500273 | 5.471234263 | 4.47E-08 | 1.14E-07 | | AFUB_102330 | Protein of unknown function | Protein of unknown function DU | Protein of unknown function | protein coding gene | | | | | | 1896 | 631 | | 439 | 2 | OG6_142064 | PF12340;PF12¢ | B0YFA8 | IPR022099;IPR022105 | | 0 | scf_000014_A_fumigatus_A1163:1,682..3,577(-) |
| **AFUB_047700** | 171.4397261 | 3.128204668 | 0.203452357 | 15.3756128 | 2.39E-53 | 3.19E-52 | | AFUB_047700 | Has domain(s) with predicted protein bindi | Ubiquitin-associated domain;Y | Ysc84 actin-binding domain; | protein coding gene | | GO:0005515 | | | protein binding | 1601 | 455 | | 364 | 1 | OG6_123003 | PF00627;PF04¢ | B0XX46 | IPR007461;IPR009060;IPR0159 | | 0 | scf_000003_A_fumigatus_A1163:3,951,782..3,953,382(-) |
| **AFUB_018470** | 975.2938037 | 3.130309708 | 0.108117482 | 28.95284528 | 2.58E-184 | 3.56E-182 | | AFUB_018470 | Has domain(s) with predicted methyltrans | Methyltransferase type 11 | Methyltransferase type 11;S | protein coding gene | | GO:0008168 | metabolic process;methylation | | methyltransferase activity | 630 | 209 | | 338 | 0 | OG6_101540 | PF08241 | B0XT94 | IPR013216;IPR( | 2.1.1.- (Methyltr | 0 | scf_000002_A_fumigatus_A1163:326,113..326,742(-) |
| **AFUB_019580** | 3.401951269 | 3.131911532 | 1.302135528 | 2.405211643 | 0.016163102 | 0.024305189 | | AFUB_019580 | Has domain(s) with predicted nucleotide b | Short-chain dehydrogenase/red | Short-chain dehydrogenase/ | protein coding gene | | | | | | 560 | 140 | | 23 | 0 | OG6_115932 | PF00106 | B0XTZ0 | IPR002347;IPR036291 | | 0 | scf_000002_A_fumigatus_A1163:625,434..625,993(+) |
| **AFUB_047940** | 15.36689908 | 3.141052195 | 0.593970294 | 5.288231126 | 1.24E-07 | 3.03E-07 | | AFUB_047940 | Protein of unknown function | N/A | N/A | protein coding gene | | | | | | 1240 | 290 | MFFSRLLFIAAF | 188 | 2 | OG6_153002 | N/A | B0XXD5 | N/A | | 0 | scf_000003_A_fumigatus_A1163:4,015,967..4,017,206(-) |
| **AFUB_000950** | 5.062955665 | 3.154424237 | 1.031064941 | 3.059384634 | 0.002217922 | 0.003742889 | | AFUB_000950 | Has domain(s) with predicted role in transı | Major facilitator superfamily | Major facilitator superfamily;l | protein coding gene | GO:0016021;G( | GO:0022857 | transmembrane | integral compon | transmembrane transporte | 1883 | 558 | | 379 | 1 | OG6_121386 | PF07690 | B0XM52 | IPR011701;IPR036259 | | 9 | scf_000001_A_fumigatus_A1163:245,078..246,960(+) |
| **AFUB_085010** | 13.75134216 | 3.154881135 | 0.631252048 | 4.997815285 | 5.80E-07 | 1.35E-06 | | AFUB_085010 | Has domain(s) with predicted protein bindi | N/A | N/A | protein coding gene | | GO:0005515 | | | protein binding | 1678 | 524 | | 85 | 0 | OG6_454618 | N/A | B0YAK8 | N/A | | 0 | scf_000007_A_fumigatus_A1163:1,542,977..1,544,654(+) |
| **AFUB_071000** | 8.661956255 | 3.158820401 | 0.793754286 | 3.979594766 | 6.90E-05 | 0.000136395 | | AFUB_071000 | Protein of unknown function | N/A | Protein of unknown function | protein coding gene | | | | | | 1258 | 307 | | 267 | 9 | OG6_137775 | N/A | B0Y510 | IPR021264 | | 0 | scf_000005_A_fumigatus_A1163:2,247,023..2,248,280(+) |
| **AFUB_082079** | 5.210967931 | 3.161173648 | 1.035633711 | 3.052405125 | 0.002270155 | 0.003826345 | | AFUB_082079 | Protein of unknown function | N/A | N/A | protein coding gene | GO:0016021 | | | integral component of membrane | | 202 | 61 | | 1 | 0 | OG6_454373 | N/A | B0Y9S2 | N/A | | 1 | scf_000007_A_fumigatus_A1163:660,311..660,512(+) |
| **AFUB_079370** | 6.808505985 | 3.162227155 | 0.893442551 | 3.539073086 | 0.000401535 | 0.000733634 | | AFUB_079370 | Protein of unknown function | N/A | N/A | protein coding gene | | | | | | 384 | 127 | | 19 | 0 | OG6r21_10220 | N/A | B0Y923 | N/A | | 0 | scf_000006_A_fumigatus_A1163:2,073,973..2,074,356(-) |
| **AFUB_029430** | 17.29438252 | 3.162649173 | 0.566819395 | 5.579641771 | 2.41E-08 | 6.23E-08 | | AFUB_029430 | Protein of unknown function | Protein of unknown function DU | Protein of unknown function | protein coding gene | | | | | | 810 | 251 | MYFLTSISIFLA | 335 | 0 | OG6_123068 | PF11937 | B0XTP2 | IPR021851 | | 0 | scf_000002_A_fumigatus_A1163:3,568,007..3,568,816(+) |
| **AFUB_029440** | 168.6530609 | 3.181098214 | 0.193675349 | 16.42489985 | 1.27E-60 | 2.00E-59 | | AFUB_029440 | Has domain(s) with predicted oxidoreducta | Malate/L-lactate dehydrogenas | Malate/L-lactate dehydrogen | protein coding gene | | GO:0016491 | metabolic process;obsolete oxid | | oxidoreductase activity | 1555 | 356 | | 219 | 0 | OG6_104354 | PF02615 | B0XTP3 | IPR003767;IPR( | 1.1.1.237 (Hydro | 0 | scf_000002_A_fumigatus_A1163:3,569,103..3,570,657(+) |
| **AFUB_055730** | 162.9395871 | 3.187242579 | 0.19872185 | 16.03871229 | 6.86E-58 | 1.02E-56 | | AFUB_055730 | Protein of unknown function | N/A | Uncharacterized protein AN1 | protein coding gene | | | | | | 1508 | 262 | | 66 | 0 | OG6_454452 | N/A | B0Y3X2 | IPR038883 | | 0 | scf_000004_A_fumigatus_A1163:1,975,236..1,976,743(-) |
| **AFUB_097600** | 5.305989087 | 3.190523723 | 1.0206588 | 3.125945441 | 0.001772344 | 0.003024865 | | AFUB_097600 | Protein of unknown function | N/A | PLAC8 motif-containing prot | protein coding gene | | | | | | 498 | 122 | | 3 | 1 | OG6_103536 | N/A | B0YE25 | IPR006461 | | 0 | scf_000009_A_fumigatus_A1163:1,390,824..1,391,321(-) |
| **AFUB_045640** | 5.290221513 | 3.190679404 | 1.020226036 | 3.127424015 | 0.001763454 | 0.003011278 | | AFUB_045640 | Has domain(s) with predicted protein bindi | Alcohol dehydrogenase, C-term | GroES-like superfamily;Alcol | protein coding gene | | GO:0016747;G( | metabolic process;obsolete oxid | | acyltransferase activity, tr | 8264 | 2463 | | 3303 | 12 | OG6_100178 | PF00107;PF001 | B0Y051 | IPR011032;IPR( | 1.1.1.100 (3-oxc | 0 | scf_000003_A_fumigatus_A1163:3,427,808..3,436,071(+) |
| **AFUB_096660** | 3.55594043 | 3.192478899 | 1.246316808 | 2.561530808 | 0.0104212 | 0.016095317 | | AFUB_096660 | Protein of unknown function | N/A | Protein kinase-like domain s | protein coding gene | | | | | | 696 | 231 | | 160 | 4 | OG6_152900 | N/A | B0YDS9 | IPR011009 | | 0 | scf_000009_A_fumigatus_A1163:1,105,926..1,106,621(-) |
| **AFUB_086940** | 3.50395392 | 3.192996826 | 1.249542567 | 2.555332576 | 0.010608638 | 0.016358591 | | AFUB_086940 | unspecified product | N/A | N/A | ncRNA gene | | | | | | 74 | | | 0 | 0 | N/A (orthology r | N/A | | N/A | | | scf_000008_A_fumigatus_A1163:105,276..105,349(+) |
| **AFUB_101400** | 66.90864898 | 3.199778175 | 0.296777477 | 10.78174197 | 4.20E-27 | 2.57E-26 | | AFUB_101400 | Protein of unknown function | N/A | P-loop containing nucleoside | protein coding gene | GO:0016021 | | | integral component of membrane | | 2159 | 671 | | 57 | 0 | OG6_453740 | N/A | B0YF32 | IPR027417 | | 1 | scf_000011_A_fumigatus_A1163:270,715..272,873(+) |
| **AFUB_084880** | 3243.232796 | 3.201670362 | 0.273003065 | 11.7275986 | 9.20E-32 | 6.63E-31 | | AFUB_084880 | Ortholog(s) have role in spore germination | N/A | N/A | protein coding gene | GO:0009277 | | spore germinati | fungal-type cell wall | | 504 | 167 | MSFSKFFSLAA | 188 | 2 | OG6_153002 | N/A | B0YAJ5 | N/A | | 0 | scf_000007_A_fumigatus_A1163:1,505,036..1,505,539(-) |
| **AFUB_032940** | 8.861217465 | 3.202574198 | 0.796944911 | 4.018564085 | 5.86E-05 | 0.000116311 | | AFUB_032940 | Protein of unknown function | N/A | N/A | protein coding gene | | | | | | 515 | 151 | MKLTSIAVALFL | 2 | 0 | OG6_454071 | N/A | B0XVM2 | N/A | | 0 | scf_000002_A_fumigatus_A1163:4,575,056..4,575,570(-) |
| **AFUB_101860** | 6.9452235 | 3.203798944 | 0.893623779 | 3.585176466 | 0.00033685 | 0.000619326 | | AFUB_101860 | Has domain(s) with predicted phosphatase | HAD-superfamily hydrolase, su | 2-phosphoglycolate phospha | protein coding gene | | GO:0016787;G( | metabolic process | | hydrolase activity;phospha | 1101 | 331 | | 384 | 1 | OG6_100677 | PF13242;PF13¢ | B0YF80 | IPR006349;IPR( | 3.1.3.18 (Phosp | 0 | scf_000011_A_fumigatus_A1163:394,971..396,071(+) |
| **AFUB_092510** | 7.012335854 | 3.205427629 | 0.88772784 | 3.610822466 | 0.000305228 | 0.000564633 | | AFUB_092510 | Protein of unknown function | FAD-binding domain | FAD-binding domain;FAD/N | protein coding gene | | GO:0071949;GO:0004497 | | | FAD binding;monooxygen | 1509 | 502 | | 139 | 0 | OG6_455137 | PF01494 | B0YBE4 | IPR002938;IPR( | 1.14.13.50 (Pen | 0 | scf_000008_A_fumigatus_A1163:1,693,359..1,694,867(+) |
| **AFUB_047120** | 42.19513284 | 3.206054629 | 0.372109285 | 8.615895268 | 6.94E-18 | 2.96E-17 | | AFUB_047120 | Has domain(s) with predicted N-acetyltran | GNAT domain | GNAT domain;Acyl-CoA N-a | protein coding gene | | GO:0008080 | | | N-acetyltransferase activi | 1306 | 249 | | 86 | 0 | OG6_121098 | PF00583 | B0XWS2 | IPR000182;IPR016181 | | 0 | scf_000003_A_fumigatus_A1163:3,804,964..3,806,269(-) |
| **AFUB_074910** | 97.50104503 | 3.2079937 | 0.252566787 | 12.70156593 | 5.80E-37 | 4.92E-36 | | AFUB_074910 | Protein of unknown function | N/A | N/A | protein coding gene | | | | | | 336 | 111 | | 180 | 0 | OG6_166545 | N/A | B0Y7T3 | N/A | | 0 | scf_000006_A_fumigatus_A1163:790,923..791,258(+) |
| **AFUB_048200** | 15.93869531 | 3.218296778 | 0.592638129 | 5.430458518 | 5.62E-08 | 1.42E-07 | | AFUB_048200 | Ortholog(s) have intracellular localization | N/A | Tetratricopeptide-like helical | protein coding gene | GO:0005622 | GO:0005515 | | intracellular ana | protein binding | 1694 | 552 | | 371 | 3 | OG6_110856 | N/A | B0XXG0 | IPR011990 | | 0 | scf_000003_A_fumigatus_A1163:4,075,619..4,077,312(+) |
| **AFUB_001020** | 3.495678947 | 3.219368488 | 1.265022669 | 2.544909721 | 0.010930599 | 0.016817381 | | AFUB_001020 | Has domain(s) with predicted chitinase, ca | Glycoside hydrolase family 18, | Glycoside hydrolase family 1 | protein coding gene | | GO:0003824;G( | carbohydrate metabolic process; | | catalytic activity;chitin bin | 4461 | 1378 | MSPFFLYASTIL | 485 | 3 | OG6_126992 | PF00704;PF014 | B0XM62 | IPR001223;IPR( | 3.2.1.14 (Chitina | 0 | scf_000001_A_fumigatus_A1163:262,383..266,843(-) |
| **AFUB_073610** | 5.426859059 | 3.222750789 | 1.016189796 | 3.171406368 | 0.001517028 | 0.002610371 | | AFUB_073610 | Protein of unknown function | N/A | N/A | protein coding gene | | | | | | 1089 | 341 | | 5 | 0 | OG6_454194 | N/A | B0Y7F3 | N/A | | 0 | scf_000006_A_fumigatus_A1163:396,182..397,270(+) |
| **AFUB_031130** | 62.46900976 | 3.225246855 | 0.305831105 | 10.54584312 | 5.31E-26 | 3.11E-25 | | AFUB_031130 | Protein of unknown function | N/A | N/A | protein coding gene | | | | | | 1431 | 476 | | 273 | 1 | OG6_186702 | N/A | B0XUJ7 | N/A | | 0 | scf_000002_A_fumigatus_A1163:4,058,826..4,060,256(-) |
| **AFUB_070130** | 128.8568527 | 3.225296467 | 0.225399547 | 14.30924114 | 1.92E-46 | 2.17E-45 | | AFUB_070130 | Protein of unknown function | N/A | N/A | protein coding gene | GO:0016021 | | | integral component of membrane | | 436 | 117 | | 2 | 0 | OG6_453713 | N/A | B0Y4S5 | N/A | | 2 | scf_000005_A_fumigatus_A1163:2,027,878..2,028,313(+) |
| **AFUB_094360** | 282.2424399 | 3.227758032 | 0.165914821 | 19.45430801 | 2.68E-84 | 7.17E-83 | | AFUB_094360 | Has domain(s) with predicted and role in c | Aromatic-ring-hydroxylating dic | Aromatic-ring-hydroxylating | protein coding gene | | GO:0051537;G( | aromatic compound catabolic pr | | 2 iron, 2 sulfur cluster bin | 1471 | 360 | | 104 | 0 | OG6_186893 | PF00848 | B0YD57 | IPR015879 | 1.14.15.7 (Choli | 0 | scf_000009_A_fumigatus_A1163:434,663..436,133(+) |
| **AFUB_046740** | 5.418584085 | 3.23976079 | 1.012070421 | 3.201121901 | 0.001368936 | 0.002370789 | | AFUB_046740 | Protein of unknown function | Carbohydrate-binding WSC;Dc | Carbohydrate-binding WSC; | protein coding gene | | | | | | 1466 | 467 | MRAGAQHLPF | 554 | 1 | OG6_105672 | PF01822;PF09¢ | B0XWN6 | IPR002889;IPR018535 | | 0 | scf_000003_A_fumigatus_A1163:3,702,009..3,703,474(+) |
| **AFUB_001370** | 538.2668607 | 3.246273189 | 0.148100973 | 21.91932381 | 1.70E-106 | 7.25E-105 | | AFUB_001370 | Protein of unknown function | Arrestin-like, N-terminal | Arrestin-like, N-terminal | protein coding gene | | | | | | 2253 | 511 | | 381 | 0 | OG6_153277 | PF00339 | B0XME0 | IPR011021 | | 0 | scf_000001_A_fumigatus_A1163:364,824..367,076(+) |
| **AFUB_062190** | 34.18855316 | 3.251809452 | 0.419228929 | 7.75664375 | 8.72E-15 | 3.23E-14 | | AFUB_062190 | Protein of unknown function | Beta-lactamase-related | Beta-lactamase-related;Beta | protein coding gene | | | | | | 1317 | 438 | MRLSCPLLLAI\ | 674 | 3 | OG6_104484 | PF00144 | B0Y2K9 | IPR001466;IPR( | 2.3.1.238 (Mona | 0 | scf_000004_A_fumigatus_A1163:3,674,600..3,675,916(-) |
| **AFUB_044370** | 11.07052803 | 3.260765851 | 0.720388248 | 4.526400675 | 6.00E-06 | 1.30E-05 | | AFUB_044370 | Protein of unknown function | Protein of unknown function DU | Protein of unknown function | protein coding gene | | | | | | 1683 | 523 | | 267 | 9 | OG6_137775 | PF11001 | B0XZG9 | IPR021264 | | 0 | scf_000003_A_fumigatus_A1163:3,009,467..3,011,149(+) |
| **AFUB_042900** | 21.98479396 | 3.267090891 | 0.509724377 | 6.409524517 | 1.46E-10 | 4.31E-10 | | AFUB_042900 | Ortholog(s) have role in ascospore formati | Mating-type protein MAT alpha | Mating-type protein MAT alp | protein coding gene | GO:0005634 | GO:0003677;G( | ascospore forma | nucleus | DNA binding;DNA binding | 1158 | 368 | | 35 | 0 | OG6r21_10058 | PF04769 | B0XZ24 | IPR006856 | | 0 | scf_000003_A_fumigatus_A1163:2,522,745..2,523,902(+) |
| **AFUB_047190** | 57.19196258 | 3.272308142 | 0.323647389 | 10.11072003 | 4.95E-24 | 2.69E-23 | | AFUB_047190 | Protein of unknown function | N/A | N/A | protein coding gene | GO:0016021 | | | integral component of membrane | | 526 | 58 | | 118 | 0 | OG6_133400 | N/A | B0XWS9 | N/A | | 1 | scf_000003_A_fumigatus_A1163:3,821,877..3,822,402(+) |
| **AFUB_044360** | 38.46898786 | 3.272527011 | 0.395291724 | 8.278764303 | 1.24E-16 | 5.01E-16 | bafB | AFUB_044360 | Protein of unknown function | N/A | N/A | protein coding gene | | | | | | 635 | 193 | MVWYRAILVCM | 13 | 2 | OG6_453939 | N/A | B0XZG8 | N/A | | 0 | scf_000003_A_fumigatus_A1163:3,007,141..3,007,775(+) |

### Table S5

| | | | | | | | | |
|---|---|---|---|---|---|---|---|---|
| **AFUB_045580** | 9.254765608 | 3.278664596 | 0.783734277 | 4.183388035 | 2.87E-05 | 5.87E-05 | | AFUB_045580 |
| **AFUB_075740** | 7.479837574 | 3.289119313 | 0.876494546 | 3.752583888 | 0.000175021 | 0.000331272 | | AFUB_075740 |
| **AFUB_049350** | 13.12943863 | 3.293627837 | 0.662334407 | 4.972756663 | 6.60E-07 | 1.53E-06 | | AFUB_049350 |
| **AFUB_000130** | 11.19590598 | 3.294936852 | 0.717341618 | 4.593260403 | 4.36E-06 | 9.55E-06 | | AFUB_000130 |
| **AFUB_081400** | 239.6892778 | 3.29785777 | 0.173415044 | 19.01713765 | 1.23E-80 | 2.98E-79 | | AFUB_081400 |
| **AFUB_046960** | 618.8371568 | 3.303276962 | 0.126576456 | 26.09708848 | 3.93E-150 | 3.42E-148 | | AFUB_046960 |
| **AFUB_085300** | 5.586951878 | 3.306272584 | 1.024006864 | 3.228760176 | 0.001243281 | 0.002158605 | | AFUB_085300 |
| **AFUB_074520** | 137.2489497 | 3.308784707 | 0.223203339 | 14.82408246 | 1.02E-49 | 1.25E-48 | | AFUB_074520 |
| **AFUB_071380** | 185.6628557 | 3.314065895 | 0.191682259 | 17.28937204 | 5.66E-67 | 1.03E-65 | | AFUB_071380 |
| **AFUB_033010** | 66.44501857 | 3.31415828 | 0.314503019 | 10.53776299 | 5.79E-26 | 3.39E-25 | | AFUB_033010 |
| **AFUB_035010** | 15.16725614 | 3.318867092 | 0.619308785 | 5.358985972 | 8.37E-08 | 2.08E-07 | | AFUB_035010 |
| **AFUB_058140** | 9.623329262 | 3.318947118 | 0.799911334 | 4.149143756 | 3.34E-05 | 6.79E-05 | | AFUB_058140 |
| **AFUB_097940** | 53.58098954 | 3.328210014 | 0.337730215 | 9.854640958 | 6.55E-23 | 3.41E-22 | | AFUB_097940 |
| **AFUB_100610** | 2.034031225 | 3.329618213 | 1.529013495 | 2.177625131 | 0.029433957 | 0.042433514 | | AFUB_100610 |
| **AFUB_097800** | 32.73970763 | 3.330121641 | 0.434108943 | 7.671165712 | 1.70E-14 | 6.25E-14 | | AFUB_097800 |
| **AFUB_065960** | 9.679894078 | 3.333432757 | 0.785960676 | 4.241220789 | 2.22E-05 | 4.60E-05 | | AFUB_065960 |
| **AFUB_046790** | 9.619482301 | 3.340217748 | 0.794997432 | 4.201545331 | 2.65E-05 | 5.45E-05 | | AFUB_046790 |
| **AFUB_052780** | 1004.761113 | 3.340447104 | 0.105840615 | 31.56110819 | 1.26E-218 | 2.30E-216 | | AFUB_052780 |
| **AFUB_014400** | 3.858870289 | 3.34141657 | 1.236832081 | 2.70159274 | 0.006900823 | 0.010929706 | | AFUB_014400 |
| **AFUB_018870** | 3.922918058 | 3.343727559 | 1.247322542 | 2.680724068 | 0.007346306 | 0.011606658 | | AFUB_018870 |
| **AFUB_097930** | 125.8223825 | 3.348823049 | 0.229471642 | 14.59362485 | 3.08E-48 | 3.62E-47 | | AFUB_097930 |
| **AFUB_009650** | 2.054412266 | 3.349027092 | 1.509020201 | 2.219338807 | 0.026463683 | 0.038485369 | | AFUB_009650 |
| **AFUB_079750** | 2.070215 | 3.349030333 | 1.507364155 | 2.221779205 | 0.026298231 | 0.038273637 | | AFUB_079750 |
| **AFUB_048070** | 11.6165261 | 3.350670841 | 0.710352764 | 4.716911101 | 2.39E-06 | 5.35E-06 | lacE | AFUB_048070 |
| **AFUB_044940** | 7.834753943 | 3.360319546 | 0.871624938 | 3.855235662 | 0.000115618 | 0.000223311 | | AFUB_044940 |
| **AFUB_081770** | 33.2866948 | 3.361936102 | 0.428421718 | 7.847258819 | 4.25E-15 | 1.59E-14 | | AFUB_081770 |
| **AFUB_073910** | 335.6573195 | 3.366183368 | 0.153323509 | 21.95477653 | 7.80E-107 | 3.37E-105 | | AFUB_073910 |
| **AFUB_033420** | 5.862649825 | 3.367643839 | 1.001355193 | 3.36308621 | 0.000770763 | 0.001369298 | | AFUB_033420 |
| **AFUB_052160** | 2.13362104 | 3.376198745 | 1.534376535 | 2.200371726 | 0.027780532 | 0.040230419 | | AFUB_052160 |
| **AFUB_066310** | 9.966966373 | 3.389291536 | 0.782690168 | 4.330310604 | 1.49E-05 | 3.13E-05 | | AFUB_066310 |
| **AFUB_046770** | 4.015132015 | 3.389839649 | 1.221354162 | 2.775476396 | 0.00551209 | 0.008850941 | | AFUB_046770 |
| **AFUB_015780** | 125.2616287 | 3.391528066 | 0.230230312 | 14.7310232 | 4.07E-49 | 4.90E-48 | | AFUB_015780 |
| **AFUB_045560** | 6.085347015 | 3.411140923 | 0.997537938 | 3.419560091 | 0.000627225 | 0.001124244 | | AFUB_045560 |
| **AFUB_057750** | 10.00618922 | 3.41883068 | 0.779214362 | 4.387535504 | 1.15E-05 | 2.43E-05 | | AFUB_057750 |
| **AFUB_014540** | 34.41184529 | 3.421738902 | 0.425100429 | 8.049248284 | 8.33E-16 | 3.22E-15 | | AFUB_014540 |
| **AFUB_047690** | 487.9992458 | 3.422805012 | 0.135822042 | 25.20065939 | 3.94E-140 | 2.85E-138 | | AFUB_047690 |
| **AFUB_054970** | 32.66332236 | 3.428983645 | 0.439569137 | 7.800783435 | 6.15E-15 | 2.30E-14 | | |
| **AFUB_055480** | 87.76330607 | 3.431273682 | 0.274267923 | 12.51066346 | 6.53E-36 | 5.36E-35 | | AFUB_055480 |
| **AFUB_080700** | 32.51545037 | 3.43321527 | 0.437851015 | 7.841058155 | 4.47E-15 | 1.67E-14 | | AFUB_080700 |
| **AFUB_079490** | 10.11806252 | 3.434348148 | 0.778571383 | 4.411089622 | 1.03E-05 | 2.19E-05 | | AFUB_079490 |
| **AFUB_053600** | 4.037133871 | 3.440449137 | 1.242788232 | 2.768330958 | 0.005634421 | 0.009035327 | | AFUB_053600 |
| **AFUB_021650** | 14.40650479 | 3.441256485 | 0.660031525 | 5.213775937 | 1.85E-07 | 4.48E-07 | | AFUB_021650 |
| **AFUB_102200** | 4.135966826 | 3.441904496 | 1.214345674 | 2.834369627 | 0.004591619 | 0.007436108 | | AFUB_102200 |
| **AFUB_023050** | 2.198647894 | 3.444115479 | 1.503553242 | 2.290650828 | 0.021983617 | 0.032410363 | | AFUB_023050 |
| **AFUB_058090** | 1317.055613 | 3.446360155 | 0.099317872 | 34.70030193 | 7.80E-264 | 2.28E-261 | | AFUB_058090 |
| **AFUB_092790** | 141.2684856 | 3.455314093 | 0.223983126 | 15.42667141 | 1.08E-53 | 1.46E-52 | | |
| **AFUB_023070** | 4.136723686 | 3.45648556 | 1.218722639 | 2.836154387 | 0.004566037 | 0.007400897 | | AFUB_023070 |
| **AFUB_090750** | 2.225834997 | 3.457411865 | 1.489312769 | 2.321481382 | 0.020260878 | 0.02999905 | | AFUB_090750 |
| **AFUB_015910** | 43.56963239 | 3.459692491 | 0.386677094 | 8.947239297 | 3.64E-19 | 1.64E-18 | | AFUB_015910 |
| **AFUB_003180** | 4.159412452 | 3.46212545 | 1.215875267 | 2.847434719 | 0.004407313 | 0.007154459 | | AFUB_003180 |
| **AFUB_092170** | 10.34151657 | 3.465230187 | 0.772932281 | 4.483226113 | 7.35E-06 | 1.59E-05 | | AFUB_092170 |
| **AFUB_017520** | 2.237861094 | 3.469046462 | 1.483646461 | 2.338189423 | 0.019377424 | 0.028797171 | | AFUB_017520 |
| **AFUB_068230** | 10.50159009 | 3.46956114 | 0.771875452 | 4.494975361 | 6.96E-06 | 1.51E-05 | | AFUB_068230 |
| **AFUB_075310** | 729.8975591 | 3.480815417 | 0.121725207 | 28.59568292 | 7.60E-180 | 9.77E-178 | | AFUB_075310 |
| **AFUB_011790** | 35.78340601 | 3.486298289 | 0.423651639 | 8.229162763 | 1.89E-16 | 7.54E-16 | | AFUB_011790 |
| **AFUB_087640** | 6.345963927 | 3.491638923 | 0.989320483 | 3.529330469 | 0.000416613 | 0.00076003 | | AFUB_087640 |
| **AFUB_071370** | 38.30986512 | 3.500001746 | 0.411010924 | 8.515592989 | 1.66E-17 | 6.96E-17 | | AFUB_071370 |
| **AFUB_020800** | 61.62103232 | 3.500927954 | 0.327503322 | 10.68974793 | 1.14E-26 | 6.80E-26 | | AFUB_020800 |
| **AFUB_074850** | 304.196973 | 3.511290132 | 0.161346171 | 21.76246333 | 5.26E-105 | 2.18E-103 | | AFUB_074850 |
| **AFUB_049660** | 8.639597816 | 3.524484689 | 0.864630126 | 4.076291796 | 4.58E-05 | 9.19E-05 | | AFUB_049660 |
| **AFUB_011880** | 8.544621469 | 3.526714405 | 0.863956223 | 4.082052205 | 4.46E-05 | 8.98E-05 | | AFUB_011880 |
| **AFUB_016520** | 4.343618141 | 3.527430392 | 1.212943158 | 2.908158036 | 0.003635646 | 0.005960034 | | AFUB_016520 |
| **AFUB_082470** | 1179.017527 | 3.527859102 | 0.102429385 | 34.44186555 | 5.96E-260 | 1.51E-257 | | AFUB_082470 |
| **AFUB_032000** | 131.6230769 | 3.54086003 | 0.235494237 | 15.035867 | 4.27E-51 | 5.40E-50 | | AFUB_032000 |
| **AFUB_062410** | 45.806985 | 3.542466603 | 0.390299096 | 9.076286969 | 1.12E-19 | 5.17E-19 | | AFUB_062410 |
| **AFUB_086860** | 2.330075052 | 3.545071126 | 1.502200529 | 2.359918704 | 0.01827894 | 0.027286734 | | AFUB_086860 |
| **AFUB_073890** | 423.3853083 | 3.545288038 | 0.150213381 | 23.60167928 | 3.70E-123 | 2.15E-121 | creD | AFUB_073890 |
| **AFUB_070180** | 4.388228794 | 3.551227109 | 1.224433712 | 2.900301645 | 0.003728037 | 0.006103188 | | AFUB_070180 |
| **AFUB_035580** | 6.665408572 | 3.552061334 | 0.990444625 | 3.586330062 | 0.000335364 | 0.000617183 | | AFUB_035580 |
| **AFUB_086820** | 2.381384672 | 3.569074662 | 1.471538055 | 2.425404257 | 0.015291349 | 0.023095209 | | AFUB_086820 |
| **AFUB_067680** | 4.46298404 | 3.583463834 | 1.206311964 | 2.970594622 | 0.002972238 | 0.00492782 | | AFUB_067680 |
| **AFUB_046510** | 9.132690965 | 3.588620174 | 0.857864012 | 4.183204011 | 2.87E-05 | 5.87E-05 | cbhC | AFUB_046510 |
| **AFUB_044380** | 11.41668645 | 3.595789543 | 0.765357797 | 4.698181109 | 2.62E-06 | 5.84E-06 | | AFUB_044380 |
| **AFUB_061850** | 8.986496598 | 3.599987781 | 0.869300499 | 4.141246652 | 3.45E-05 | 7.01E-05 | | AFUB_061850 |
| **AFUB_070790** | 2.449253886 | 3.600529218 | 1.50234115 | 2.396612259 | 0.016547424 | 0.024832733 | | AFUB_070790 |
| **AFUB_099440** | 61.0383718 | 3.60121082 | 0.339082526 | 10.62045533 | 2.39E-26 | 1.41E-25 | | AFUB_099440 |
| **AFUB_066720** | 2.44551241 | 3.607787836 | 1.550398552 | 2.327006711 | 0.019964908 | 0.029606318 | | AFUB_066720 |
| **AFUB_060540** | 2.484039072 | 3.611090049 | 1.510941488 | 2.389960219 | 0.016850198 | 0.025271363 | | AFUB_060540 |
| **AFUB_014780** | 130.175836 | 3.61208564 | 0.237779058 | 15.19093261 | 4.06E-52 | 5.28E-51 | | AFUB_014780 |
| **AFUB_017010** | 18.33746639 | 3.614900819 | 0.603036388 | 5.994498657 | 2.04E-09 | 5.63E-09 | | AFUB_017010 |
| **AFUB_078970** | 126.1232054 | 3.618491477 | 0.243549211 | 14.85733194 | 6.24E-50 | 7.66E-49 | | AFUB_078970 |
| **AFUB_084670** | 22.95647991 | 3.618793206 | 0.545425569 | 6.634806676 | 3.25E-11 | 1.00E-10 | | AFUB_084670 |
| **AFUB_097660** | 4.604119922 | 3.620736402 | 1.203931311 | 3.007427723 | 0.002634688 | 0.004406848 | | AFUB_097660 |
| **AFUB_091070** | 154.3872352 | 3.624294978 | 0.222042349 | 16.32253936 | 6.82E-60 | 1.05E-58 | | AFUB_091070 |
| **AFUB_045040** | 9.301220681 | 3.633436691 | 0.851560225 | 4.266799439 | 1.98E-05 | 4.13E-05 | | AFUB_045040 |
| **AFUB_045290** | 11.86004014 | 3.665947356 | 0.759082393 | 4.829445905 | 1.37E-06 | 3.11E-06 | | AFUB_045290 |
| **AFUB_025280** | 2.553564263 | 3.669706176 | 1.450116588 | 2.53062837 | 0.011385841 | 0.017487084 | | AFUB_025280 |
| **AFUB_099480** | 4.809580627 | 3.683728751 | 1.210591168 | 3.042917253 | 0.002342967 | 0.003940111 | | AFUB_099480 |
| **AFUB_100830** | 2.572476392 | 3.689236771 | 1.464121372 | 2.519761572 | 0.011743435 | 0.018007597 | | AFUB_100830 |
| **AFUB_099510** | 2.600340385 | 3.694323552 | 1.462868767 | 2.525396424 | 0.011556784 | 0.017735495 | | AFUB_099510 |
| **AFUB_045760** | 7.343520709 | 3.707234411 | 0.980295033 | 3.781753744 | 0.000155727 | 0.000296208 | | AFUB_045760 |
| **AFUB_042510** | 464.5896198 | 3.709412552 | 0.149391236 | 24.83018851 | 4.23E-136 | 2.96E-134 | | AFUB_042510 |
| **AFUB_088290** | 371.4018702 | 3.711377428 | 0.164184314 | 22.60494523 | 3.87E-113 | 1.90E-111 | | AFUB_088290 |
| **AFUB_084250** | 1.247546186 | 3.71260335 | 1.703001651 | 2.180035085 | 0.029254861 | 0.042200575 | | AFUB_084250 |
| **AFUB_098940** | 1.247546186 | 3.71260335 | 1.703001651 | 2.180035085 | 0.029254861 | 0.042200575 | | AFUB_098940 |
| **AFUB_079880** | 1.252044522 | 3.716398346 | 1.716883675 | 2.164618606 | 0.030416905 | 0.043726286 | | AFUB_079880 |
| **AFUB_061440** | 7.417322011 | 3.718307592 | 0.984504979 | 3.776829646 | 0.000158837 | 0.000301885 | | AFUB_061440 |
| **AFUB_060550** | 1.259607444 | 3.722532185 | 1.693089549 | 2.198662314 | 0.027901943 | 0.040394098 | | AFUB_060550 |
| **AFUB_020600** | 1.264105781 | 3.726372745 | 1.726746339 | 2.158031357 | 0.030925394 | 0.044410894 | | AFUB_020600 |
| **AFUB_024210** | 426.6560295 | 3.731432863 | 0.146748635 | 25.42737703 | 1.26E-142 | 9.53E-141 | | AFUB_024210 |
| **AFUB_055810** | 1.271668703 | 3.732346217 | 1.698309908 | 2.197682648 | 0.027971729 | 0.040482965 | | AFUB_055810 |
| **AFUB_044760** | 17.12297536 | 3.732412544 | 0.648569668 | 5.754836727 | 8.67E-09 | 2.31E-08 | | AFUB_044760 |
| **AFUB_044230** | 1.279231625 | 3.738200831 | 1.689367452 | 2.212781373 | 0.026912722 | 0.039091195 | | AFUB_044230 |
| **AFUB_051200** | 1.279231625 | 3.738200831 | 1.689367452 | 2.212781373 | 0.026912722 | 0.039091195 | | AFUB_051200 |
| **AFUB_046880** | 15.06894781 | 3.745403281 | 0.691056878 | 5.419819122 | 5.97E-08 | 1.50E-07 | | AFUB_046880 |
| **AFUB_084980** | 12.49444074 | 3.747001753 | 0.75549023 | 4.959695842 | 7.06E-07 | 1.63E-06 | | AFUB_084980 |
| **AFUB_016950** | 1.291292884 | 3.748256218 | 1.70914354 | 2.193061103 | 0.028302977 | 0.040937779 | | AFUB_016950 |
| **AFUB_088370** | 22.4405021 | 3.753498237 | 0.573297016 | 6.547213981 | 5.86E-11 | 1.78E-10 | | AFUB_088370 |
| **AFUB_035620** | 2.709149099 | 3.761831106 | 1.44365092 | 2.605776128 | 0.009166635 | 0.014274353 | | AFUB_035620 |
| **AFUB_034740** | 68666.27945 | 3.769204672 | 0.081689505 | 46.14062326 | 0 | 0 | | AFUB_034740 |
| **AFUB_046250** | 5.07166645 | 3.77561867 | 1.182953077 | 3.191689293 | 0.001414434 | 0.002443874 | | AFUB_046250 |
| **AFUB_081300** | 51.13989634 | 3.789324146 | 0.383495584 | 9.881011157 | 5.03E-23 | 2.64E-22 | | AFUB_081300 |
| **AFUB_042120** | 38.73303898 | 3.806102826 | 0.438223987 | 8.685290934 | 3.78E-18 | 1.63E-17 | | AFUB_042120 |
| **AFUB_043990** | 26.05595344 | 3.814746639 | 0.532094916 | 7.169297284 | 7.54E-13 | 2.56E-12 | | AFUB_043990 |
| **AFUB_084940** | 13.11211983 | 3.819128248 | 0.750183224 | 5.090927288 | 3.56E-07 | 8.44E-07 | | AFUB_084940 |
| **AFUB_035640** | 81.01773506 | 3.826335922 | 0.313460725 | 12.20674749 | 2.86E-34 | 2.23E-33 | | AFUB_035640 |
| **AFUB_021230** | 7.992885231 | 3.833459169 | 0.960145516 | 3.992581443 | 6.54E-05 | 0.000129347 | | AFUB_021230 |
| **AFUB_021570** | 2.842045168 | 3.840302794 | 1.428533657 | 2.688283034 | 0.007182049 | 0.011358317 | | AFUB_021570 |
| **AFUB_058320** | 15.86735814 | 3.842548069 | 0.682869701 | 5.627058956 | 1.83E-08 | 4.78E-08 | | AFUB_058320 |
| **AFUB_046850** | 456.2820484 | 3.847469942 | 0.149472017 | 25.74040289 | 4.13E-146 | 3.29E-144 | | AFUB_046850 |
| **AFUB_035550** | 2.833770194 | 3.847567409 | 1.424708733 | 2.700599302 | 0.006921467 | 0.010960601 | | AFUB_035550 |
| **AFUB_084020** | 2500.191223 | 3.848212242 | 0.091697275 | 41.9664841 | 0 | 0 | | AFUB_084020 |
| **AFUB_090310** | 106.6216672 | 3.850560229 | 0.275963329 | 13.95315909 | 3.01E-44 | 3.20E-43 | | AFUB_090310 |
| **AFUB_097580** | 21.70904886 | 3.858419734 | 0.598862285 | 6.442916565 | 1.17E-10 | 3.49E-10 | | AFUB_097580 |
| **AFUB_004210** | 53.67208162 | 3.861542609 | 0.380814712 | 10.14021382 | 3.66E-24 | 2.00E-23 | ordA | AFUB_004210 |
| **AFUB_013900** | 320.5489719 | 3.86663907 | 0.173815558 | 22.24564431 | 1.24E-109 | 5.68E-108 | easD | AFUB_013900 |
| **AFUB_090100** | 376.2652707 | 3.866857462 | 0.16002662 | 24.16383882 | 5.34E-129 | 3.36E-127 | | AFUB_090100 |
| **AFUB_084730** | 8.053423397 | 3.876902117 | 0.963448307 | 4.023985605 | 5.72E-05 | 0.000113805 | | AFUB_084730 |
| **AFUB_013130** | 2.943195496 | 3.882996224 | 1.455779352 | 2.667297223 | 0.007646402 | 0.012049178 | | AFUB_013130 |
| **AFUB_080640** | 1.407629358 | 3.885637183 | 1.649063809 | 2.356268545 | 0.018459567 | 0.027526523 | | AFUB_080640 |
| **AFUB_096520** | 1366.846626 | 3.887845476 | 0.107649017 | 36.11594031 | 1.27E-285 | 4.91E-283 | | AFUB_096520 |
| **AFUB_058740** | 5.391120742 | 3.888831625 | 1.181880738 | 3.290375671 | 0.001000537 | 0.001755175 | | AFUB_058740 |
| **AFUB_071440** | 76.86011314 | 3.888991775 | 0.323227801 | 12.03173665 | 2.42E-33 | 1.83E-32 | | AFUB_071440 |
| **AFUB_003740** | 8.276832638 | 3.89486478 | 0.956701355 | 4.071139607 | 4.68E-05 | 9.38E-05 | | AFUB_003740 |
| **AFUB_039690** | 16.42859916 | 3.896653577 | 0.689858106 | 5.6484856 | 1.62E-08 | 4.24E-08 | | AFUB_039690 |
| **AFUB_088390** | 8.253466982 | 3.899389556 | 0.955941932 | 4.079107134 | 4.52E-05 | 9.09E-05 | | AFUB_088390 |
| **AFUB_055180** | 8.301747177 | 3.900326641 | 0.96000249 | 4.062829715 | 4.85E-05 | 9.71E-05 | | AFUB_055180 |
| **AFUB_101900** | 1.439314797 | 3.908368798 | 1.644475228 | 2.376666265 | 0.017469886 | 0.026147845 | | AFUB_101900 |
| **AFUB_067040** | 96.78393722 | 3.910656588 | 0.291242508 | 13.42749247 | 4.17E-41 | 3.99E-40 | | AFUB_067040 |
| **AFUB_101930** | 1.443813133 | 3.912260804 | 1.683995424 | 2.323201565 | 0.020168327 | 0.029875787 | | AFUB_101930 |
| **AFUB_046010** | 1.446877719 | 3.913602638 | 1.643401539 | 2.381403781 | 0.017246796 | 0.025821958 | | AFUB_046010 |
| **AFUB_082420** | 1.446877719 | 3.913602638 | 1.643401539 | 2.381403781 | 0.017246796 | 0.025821958 | | AFUB_082420 |
| **AFUB_044410** | 25.23404186 | 3.916611429 | 0.565334176 | 6.927958008 | 4.27E-12 | 1.39E-11 | | AFUB_044410 |
| **AFUB_071900** | 2.993096506 | 3.924873435 | 1.425819211 | 2.752714654 | 0.005910339 | 0.009446346 | | AFUB_071900 |
| **AFUB_091430** | 81.48729479 | 3.934282536 | 0.316307287 | 12.43816597 | 1.62E-35 | 1.31E-34 | | AFUB_091430 |
| **AFUB_077880** | 25.52554217 | 3.941890418 | 0.557787597 | 7.067009805 | 1.58E-12 | 5.26E-12 | | AFUB_077880 |
| **AFUB_092390** | 3.025538805 | 3.94202064 | 1.412060976 | 2.791678763 | 0.005243538 | 0.008433773 | | AFUB_092390 |
| **AFUB_094800** | 74.73829716 | 3.959270065 | 0.337535354 | 11.72994181 | 8.95E-32 | 6.48E-31 | | AFUB_094800 |
| **AFUB_069990** | 432.1936065 | 3.962304053 | 0.155260277 | 25.52039787 | 1.17E-143 | 8.95E-142 | | AFUB_069990 |
| **AFUB_044970** | 336.9504975 | 3.962917251 | 0.17527545 | 22.60965378 | 3.48E-113 | 1.72E-111 | | AFUB_044970 |
| **AFUB_085580** | 57.7800615 | 3.975933583 | 0.375442047 | 10.59000613 | 3.32E-26 | 1.95E-25 | | AFUB_085580 |
| **AFUB_001380** | 8.64548591 | 3.979861918 | 0.969359264 | 4.105662436 | 4.03E-05 | 8.14E-05 | | AFUB_001380 |
| **AFUB_081220** | 3.125315685 | 3.999877297 | 1.419359089 | 2.81808693 | 0.004831073 | 0.007805536 | | AFUB_081220 |
| **AFUB_085090** | 20.85133747 | 4.003250045 | 0.647060049 | 6.18682926 | 6.14E-10 | 1.74E-09 | | AFUB_085090 |
| **AFUB_084850** | 38.3061742 | 4.004376566 | 0.464934617 | 8.612773529 | 7.13E-18 | 3.04E-17 | | AFUB_084850 |

### Table S5

| | | | | | | | | | | | | | | | | | | | | | | | | | | | | | |
|---|---|---|---|---|---|---|---|---|---|---|---|---|---|---|---|---|---|---|---|---|---|---|---|---|---|---|---|---|---|
| AFUB_085360 | 73.88392088 | 4.00880867 | 0.337580819 | 11.87510796 | 1.59E-32 | 1.17E-31 | | AFUB_085360 | Protein of unknown function | Phosphatidylglycerol lysyltrans | Phosphatidylglycerol lysyltra | protein coding gene | | | | | | 2157 | 509 | | 88 | 0 | OG6_117021 | PF09924 | | IPR024320 | | 0 | scf_000007_A_fumigatus_A1163:1,659,241..1,661,397(+) |
| AFUB_044650 | 1.523965832 | 4.009773906 | 1.66303576 | 2.411117068 | 0.015903743 | 0.023933854 | | AFUB_044650 | Has domain(s) with predicted oleate hydra | Oleate hydratase | Oleate hydratase;FAD/NAD( | protein coding gene | | GO:0071949;G( | fatty acid metabolic process | | FAD binding;oleate hydra | 1665 | 538 | | 126 | 1 | OG6_137768 | PF06100 | B0XZN3 | IPR010354;IPR( | 5.2.-.- (Cis-trans | 0 | scf_000003_A_fumigatus_A1163:3,153,324..3,154,988(-) |
| AFUB_018180 | 1140.067188 | 4.020667285 | 0.112385245 | 35.77575748 | 2.63E-280 | 9.39E-278 | | AFUB_018180 | Protein of unknown function | Pirin, N-terminal domain | Pirin, N-terminal domain;Rm | protein coding gene | | | | | | 1078 | 287 | | 159 | 0 | OG6_107618 | PF02678 | B0XT65 | IPR003829;IPR( | 1.13.11.24 (Que | 0 | scf_000002_A_fumigatus_A1163:250,236..251,313(+) |
| AFUB_052580 | 116.3271338 | 4.024207272 | 0.2741785 | 14.67732619 | 9.01E-49 | 1.08E-47 | | AFUB_052580 | Has domain(s) with predicted catalytic, AT | Spo11/DNA topoisomerase VI, | Spo11/DNA topoisomerase | protein coding gene | GO:0005694 | GO:0005524;G( | DNA catabolic p | chromosome | ATP binding;DNA binding | 1724 | 383 | | 304 | 0 | OG6_101269 | PF04406 | | IPR002815;IPR( | 5.6.2.2 (DNA to | 0 | scf_000004_A_fumigatus_A1163:1,012,077..1,013,800(-) |
| AFUB_033150 | 39.05715703 | 4.034810115 | 0.46498814 | 8.677232311 | 4.06E-18 | 1.75E-17 | | AFUB_033150 | Protein of unknown function | N/A | Transient receptor potential | protein coding gene | GO:0016021 | | | integral component of membrane | | 2507 | 753 | MLPESYSLRCL | 53 | 0 | OG6_454301 | N/A | B0XVP3 | IPR040241 | | 10 | scf_000002_A_fumigatus_A1163:4,617,222..4,619,728(-) |
| AFUB_101520 | 1.560149608 | 4.034978452 | 1.634420775 | 2.468751323 | 0.01355854 | 0.020643087 | | AFUB_101520 | Protein of unknown function | N/A | N/A | protein coding gene | | | | | | 702 | 174 | | 460 | 3 | OG6_100774 | N/A | B0YF44 | N/A | | 0 | scf_000011_A_fumigatus_A1163:299,873..300,574(+) |
| AFUB_033040 | 1.570777116 | 4.041579453 | 1.63308017 | 2.474819991 | 0.013330334 | 0.020318112 | | AFUB_033040 | Protein of unknown function | N/A | N/A | protein coding gene | | | | | | 770 | 199 | | 7 | 0 | OG6_454386 | N/A | B0XVN2 | N/A | | 0 | scf_000002_A_fumigatus_A1163:4,599,033..4,599,802(-) |
| AFUB_097870 | 1.575275452 | 4.044876392 | 1.601537479 | 2.525620815 | 0.011549406 | 0.017726996 | | AFUB_097870 | Has domain(s) with predicted protein bindi | Ankyrin repeat;Ankyrin repeat- | Ankyrin repeat;Ankyrin repea | protein coding gene | | GO:0005515 | | | protein binding | 1089 | 272 | | 938 | 1 | OG6_104829 | PF00023;PF127 | B0YE52 | IPR002110;IPR020683;IPR0367 | | 0 | scf_000009_A_fumigatus_A1163:1,487,339..1,488,427(+) |
| AFUB_032910 | 15.33543314 | 4.055683284 | 0.741931203 | 5.466387272 | 4.59E-08 | 1.17E-07 | | AFUB_032910 | Has domain(s) with predicted hydrolase a | Calcineurin-like phosphoestera | Calcineurin-like phosphoeste | protein coding gene | | GO:0016787 | | | hydrolase activity | 999 | 332 | | 806 | 3 | OG6_101637 | PF00149 | B0XVL9 | IPR004843 | | 0 | scf_000002_A_fumigatus_A1163:4,570,747..4,571,745(-) |
| AFUB_045010 | 6.147068206 | 4.073353349 | 1.161254275 | 3.507718711 | 0.000451967 | 0.000822813 | | AFUB_045010 | Ortholog(s) have role in secondary metab | Major facilitator superfamily | Major facilitator superfamily; | protein coding gene | GO:0016021;G( | GO:0022857 | cellular respons | integral compon | transmembrane transport | 1361 | 420 | | 73 | 0 | OG6_113922 | PF07690 | B0XZR8 | IPR011701;IPR036259 | | 12 | scf_000003_A_fumigatus_A1163:3,245,065..3,246,425(+) |
| AFUB_102220 | 1.622086735 | 4.075121134 | 1.62160946 | 2.513010212 | 0.011970587 | 0.018344239 | | AFUB_102220 | Protein of unknown function | NYN domain, limkain-b1-type;( | NYN domain, limkain-b1-typ | protein coding gene | | | | | | 630 | 178 | | 1 | 0 | OG6_106891 | PF01936;PF128 | B0YFA6 | IPR021139;IPR025605 | | 0 | scf_000012_A_fumigatus_A1163:23,215..23,844(+) |
| AFUB_099940 | 3.317796348 | 4.077014721 | 1.394493498 | 2.923652729 | 0.003459505 | 0.005690643 | | AFUB_099940 | Has domain(s) with predicted catalytic acti | Glycosyl transferase, family 20 | Glycosyl transferase, family | protein coding gene | | GO:0003824 | trehalose biosynthetic process | | catalytic activity | 1665 | 475 | | 840 | 4 | OG6_100616 | PF00982 | B0YEP8 | IPR001830 | 2.4.1.15 (Alpha, | 0 | scf_000010_A_fumigatus_A1163:478,249..479,913(-) |
| AFUB_080010 | 3.31326285 | 4.079321559 | 1.395025831 | 2.924190698 | 0.003453531 | 0.005682757 | | AFUB_080010 | Protein of unknown function | Protein of unknown function DU | Protein of unknown function | protein coding gene | | | | | | 2565 | 759 | | 276 | 3 | OG6_121380 | PF11917 | B0YB02 | IPR021842 | | 0 | scf_000007_A_fumigatus_A1163:5,126..7,690(+) |
| AFUB_014470 | 1720.044222 | 4.082630355 | 0.10608364 | 38.48501403 | 0 | 0 | | AFUB_014470 | Protein of unknown function | N/A | N/A | protein coding gene | | | | | | 1571 | 124 | | 54 | 0 | OG6_454275 | N/A | B0XN38 | N/A | | 0 | scf_000001_A_fumigatus_A1163:4,069,780..4,071,350(+) |
| AFUB_032890 | 3.31401971 | 4.084695663 | 1.393449209 | 2.931355973 | 0.003374858 | 0.005563752 | | AFUB_032890 | Has domain(s) with predicted zinc ion binc | N/A | N/A | protein coding gene | | | | | | 1021 | 177 | | 4 | 0 | OG6_455090 | N/A | B0XVL7 | N/A | | 0 | scf_000002_A_fumigatus_A1163:4,567,287..4,568,307(+) |
| AFUB_000100 | 3.341918865 | 4.084702183 | 1.392210748 | 2.933968287 | 0.003346584 | 0.005519973 | | AFUB_000100 | Has domain(s) with predicted magnesium | Isopropylmalate dehydrogenas | Tartrate dehydrogenase;Isop | protein coding gene | | GO:0051287;G( | obsolete oxidation-reduction pro( | | NAD binding;magnesium | 1278 | 360 | | 679 | 4 | OG6_103350 | PF00180 | B0XMF8 | IPR011829;IPR( | 1.1.1.83 (D-mal | 0 | scf_000001_A_fumigatus_A1163:24,630..25,907(+) |
| AFUB_071860 | 21.98612617 | 4.103306146 | 0.622554964 | 6.591074497 | 4.37E-11 | 1.33E-10 | | AFUB_071860 | Has domain(s) with predicted protein bindi | Zinc finger, RING-type | Zinc finger, RING-type | protein coding gene | | GO:0008270 | | | zinc ion binding | 902 | 134 | | 12 | 0 | OG6_454271 | PF13639 | B0Y597 | IPR001841 | | 0 | scf_000005_A_fumigatus_A1163:2,554,816..2,555,717(+) |
| AFUB_048850 | 3.399322495 | 4.106646771 | 1.407255965 | 2.918194608 | 0.003520645 | 0.005784303 | | AFUB_048850 | Has domain(s) with predicted catalytic, py | Aminotransferase, class I/class | Aminotransferases, class-I, p | protein coding gene | | GO:0003824;G( | biosynthetic process | | catalytic activity;pyridoxal | 1378 | 443 | | 455 | 2 | OG6_103084 | PF00155 | B0Y166 | IPR004838;IPR( | 2.6.1.- (Transam | 0 | scf_000004_A_fumigatus_A1163:39,723..41,100(-) |
| AFUB_071780 | 6.275465939 | 4.106925401 | 1.162270923 | 3.533535358 | 0.000410041 | 0.000748751 | | AFUB_071780 | Ortholog(s) have sequence-specific DNA I | Zn(2)-C6 fungal-type DNA-binc | Zn(2)-C6 fungal-type DNA-b | protein coding gene | GO:0042025;G( | GO:0003677;G( | aflatoxin biosynt | host cell nucleus | DNA binding;DNA-binding | 1191 | 396 | | 123 | 0 | OG6_455265 | PF00172;PF084 | B0Y589 | IPR001138;IPR013700;IPR0368 | | 0 | scf_000005_A_fumigatus_A1163:2,533,111..2,534,301(-) |
| AFUB_033430 | 35.13570439 | 4.125098316 | 0.499669976 | 8.255645749 | 1.51E-16 | 6.07E-16 | | AFUB_033430 | Has domain(s) with predicted role in trans | Major facilitator superfamily | Major facilitator superfamily; | protein coding gene | GO:0016021;G( | GO:0022857 | transmembrane | integral compon | transmembrane transport | 1545 | 514 | | 120 | 0 | OG6_454019 | PF07690 | B0XVR9 | IPR011701;IPR036259 | | 12 | scf_000002_A_fumigatus_A1163:4,679,156..4,680,700(-) |
| AFUB_029480 | 124.6339334 | 4.129808388 | 0.271930737 | 15.18698636 | 4.31E-52 | 5.60E-51 | | AFUB_029480 | Has domain(s) with predicted protein bindi | CBM21 (carbohydrate binding | CBM21 (carbohydrate bindin | protein coding gene | | GO:0005515 | | | protein binding | 3770 | 741 | | 287 | 0 | OG6_115051 | PF03370 | | IPR005036 | | 0 | scf_000002_A_fumigatus_A1163:3,588,200..3,591,969(+) |
| AFUB_064910 | 158.2916155 | 4.139121493 | 0.248263304 | 16.67230487 | 2.08E-62 | 3.42E-61 | | AFUB_064910 | Protein of unknown function | Pyridoxal-phosphate depender | Serine/threonine dehydratas | protein coding gene | | GO:0030170 | cellular amino acid metabolic prc | | pyridoxal phosphate bindi | 1453 | 460 | | 341 | 1 | OG6_105174 | PF00291 | B0Y5X4 | IPR000634;IPR( | 4.3.1.17 (L-serin | 0 | scf_000005_A_fumigatus_A1163:591,166..592,618(-) |
| AFUB_102560 | 51.59267351 | 4.142608012 | 0.417426248 | 9.92416752 | 3.27E-23 | 1.72E-22 | | AFUB_102560 | unspecified product | N/A | N/A | ncRNA gene | | | | | | 74 | | | 0 | 0 | N/A (orthology r | N/A | | N/A | | | scf_000015_A_fumigatus_A1163:10,949..11,022(-) |
| AFUB_086210 | 19.52153423 | 4.149829855 | 0.669546997 | 6.197966492 | 5.72E-10 | 1.63E-09 | | AFUB_086210 | Protein of unknown function | N/A | Papain-like cysteine peptida | protein coding gene | | | | | | 893 | 264 | MRTVFTPGIKA | 24 | 0 | OG6_152566 | N/A | B0YAY0 | IPR038765 | | 0 | scf_000007_A_fumigatus_A1163:1,869,710..1,870,602(-) |
| AFUB_017730 | 3.480908944 | 4.152900579 | 1.385584409 | 2.997219477 | 0.002724545 | 0.00454846 | | AFUB_017730 | Protein of unknown function | N/A | N/A | protein coding gene | | | | | | 501 | 166 | | 6 | 0 | OG6_454367 | N/A | B0XSV3 | N/A | | 0 | scf_000002_A_fumigatus_A1163:135,958..136,458(-) |
| AFUB_036230 | 127.4888627 | 4.154826769 | 0.274307533 | 15.14660107 | 7.98E-52 | 1.03E-50 | | AFUB_036230 | Ortholog(s) have role in gliotoxin biosynth | Cytochrome P450 | Cytochrome P450;Cytochror | protein coding gene | | GO:0009055;G( | gliotoxin biosynthetic process;ob | | electron transfer activity;h | 2193 | 539 | | 145 | 1 | OG6_166373 | PF00067 | | IPR001128;IPR( | 1.8.1.- (With NA | 0 | scf_000003_A_fumigatus_A1163:616,436..618,628(+) |
| AFUB_033070 | 16.30278305 | 4.1567602 | 0.731220581 | 5.684687094 | 1.31E-08 | 3.46E-08 | | AFUB_033070 | Has domain(s) with predicted 3-beta-hydr | 3-beta hydroxysteroid dehydro | 3-beta hydroxysteroid dehyd | protein coding gene | | GO:0003854;G( | obsolete oxidation-reduction pro( | | 3-beta-hydroxy-delta5-ste | 1134 | 377 | | 380 | 1 | OG6_102847 | PF01073 | B0XVN5 | IPR002225;IPR( | 1.1.1.145 (3-bet | 0 | scf_000002_A_fumigatus_A1163:4,605,186..4,606,319(+) |
| AFUB_071980 | 176.7286405 | 4.16101541 | 0.236166871 | 17.61896324 | 1.76E-69 | 3.41E-68 | | AFUB_071980 | Has domain(s) with predicted N-acetyltran | GNAT domain | GNAT domain;Acyl-CoA N-a | protein coding gene | | GO:0008080 | | | N-acetyltransferase activit | 594 | 197 | | 80 | 0 | OG6_116608 | PF13302 | B0Y5A9 | IPR000182;IPR( | 2.3.1.- (Transfer | 0 | scf_000005_A_fumigatus_A1163:2,579,198..2,579,791(+) |
| AFUB_049400 | 1.708171521 | 4.167537999 | 1.603450571 | 2.599105999 | 0.009346691 | 0.014538291 | | AFUB_049400 | Protein of unknown function | Tannase/feruloyl esterase | Tannase/feruloyl esterase;Al | protein coding gene | | GO:0030600;GO:0046872 | | | feruloyl esterase activity;m | 1566 | 505 | | 854 | 2 | OG6_115949 | PF07519 | B0Y1I3 | IPR011118;IPR( | 3.1.1.- (Carboxy | 0 | scf_000004_A_fumigatus_A1163:174,074..175,639(-) |
| AFUB_016280 | 3.551877906 | 4.170541174 | 1.388336031 | 3.003985404 | 0.002664681 | 0.004452379 | | AFUB_016280 | Ortholog(s) have intracellular localization | Beta-lactamase-related | Beta-lactamase-related;Beta | protein coding gene | GO:0005622 | | | intracellular anatomical structure | | 1296 | 374 | | 674 | 3 | OG6_104484 | PF00144 | B0XP02 | IPR001466;IPR( | 2.3.1.238 (Mona | 0 | scf_000001_A_fumigatus_A1163:4,620,799..4,622,094(+) |
| AFUB_084780 | 6.553259033 | 4.170906916 | 1.153559413 | 3.615684525 | 0.000299555 | 0.000554672 | | AFUB_084780 | Protein of unknown function | N/A | N/A | protein coding gene | | | | | | 1423 | 246 | | 5 | 0 | OG6_454813 | N/A | B0YAI4 | N/A | | 0 | scf_000007_A_fumigatus_A1163:1,479,621..1,481,043(+) |
| AFUB_079450 | 1.723297366 | 4.176770553 | 1.572366583 | 2.656359272 | 0.007898941 | 0.012426808 | | AFUB_079450 | Has domain(s) with predicted sequence-s | N/A | Zn(2)-C6 fungal-type DNA-b | protein coding gene | GO:0042025;G( | GO:0003677;G( | regulation of tra | host cell nucleus | DNA binding;DNA-binding | 1551 | 499 | | 70 | 0 | OG6_186573 | N/A | B0Y931 | IPR001138;IPR036864 | | 0 | scf_000006_A_fumigatus_A1163:2,084,851..2,086,401(-) |
| AFUB_084660 | 1.723297366 | 4.176770553 | 1.572366583 | 2.656359272 | 0.007898941 | 0.012426808 | | AFUB_084660 | Has domain(s) with predicted catalytic, ca | N/A | N/A | protein coding gene | | GO:0003824 | carbohydrate metabolic process | | catalytic activity | 1503 | 278 | MRFTQLLIAAG | 1 | 0 | OG6_453719 | N/A | B0YAH2 | N/A | | 0 | scf_000007_A_fumigatus_A1163:1,446,816..1,448,318(+) |
| AFUB_078410 | 9.871812241 | 4.177990108 | 0.940666363 | 4.44152175 | 8.93E-06 | 1.91E-05 | gel7 | AFUB_078410 | Ortholog(s) have 1,3-beta-glucanosyltrans | Glucanosyltransferase;X8 dom | Glucanosyltransferase;X8 do | protein coding gene | GO:0031225;G( | GO:0042124;G( | carbohydrate m | anchored comp( | 1,3-beta-glucanosyltransf | 1679 | 541 | MRFSSILAGAIL | 652 | 2 | OG6_107446 | PF03198;PF079 | B0Y8S8 | IPR004886;IPR( | 2.4.1.- (Hexosyl | 0 | scf_000006_A_fumigatus_A1163:1,825,481..1,827,159(+) |
| AFUB_012720 | 722.7720822 | 4.182940316 | 0.135683099 | 30.82874982 | 1.08E-208 | 1.82E-206 | | AFUB_012720 | Has domain(s) with predicted RNA-directe | N/A | N/A | protein coding gene | | GO:0003723;G( | RNA-dependent DNA biosynthet | | RNA binding;RNA-directe | 2688 | 895 | | 188 | 0 | OG6_127168 | N/A | B0XRB0 | N/A | | 0 | scf_000001_A_fumigatus_A1163:3,580,636..3,583,323(+) |
| AFUB_078000 | 1.73985696 | 4.18674743 | 1.584584781 | 2.642173193 | 0.008237592 | 0.012934241 | | | | | | | | | | | | | | | | | | | | | | | |
| AFUB_040600 | 10.01525585 | 4.18792434 | 0.937890459 | 4.46525956 | 8.00E-06 | 1.72E-05 | | AFUB_040600 | Protein of unknown function | N/A | Class I glutamine amidotrans | protein coding gene | | | | | | 663 | 178 | | 5 | 0 | OG6_455120 | N/A | B0XYE2 | IPR029062 | | 0 | scf_000003_A_fumigatus_A1163:1,872,320..1,872,982(-) |
| AFUB_032010 | 5229.562624 | 4.192875331 | 0.083154062 | 50.42297699 | 0 | 0 | | AFUB_032010 | Protein of unknown function | N/A | N/A | protein coding gene | | | | | | 498 | 165 | | 53 | 0 | OG6_454135 | N/A | B0XUZ9 | N/A | | 0 | scf_000002_A_fumigatus_A1163:4,312,730..4,313,227(+) |
| AFUB_084500 | 1.754982805 | 4.195316398 | 1.565528599 | 2.679808214 | 0.007366435 | 0.011634432 | | AFUB_084500 | Protein of unknown function | N/A | N/A | protein coding gene | | | | | | 762 | 189 | | 1 | 0 | OG6_453804 | N/A | B0YAF6 | N/A | | 0 | scf_000007_A_fumigatus_A1163:1,397,604..1,398,365(+) |
| AFUB_024220 | 284.7584935 | 4.198117166 | 0.191980153 | 21.86745398 | 5.30E-106 | 2.25E-104 | | AFUB_024220 | Has domain(s) with predicted sequence-s | Zn(2)-C6 fungal-type DNA-binc | Zn(2)-C6 fungal-type DNA-b | protein coding gene | GO:0042025;G( | GO:0003677;G( | regulation of tra | host cell nucleus | DNA binding;DNA-binding | 2351 | 726 | | 348 | 0 | OG6_152943 | PF00172;PF04( | B0XWB4 | IPR001138;IPR007219;IPR0368 | | 0 | scf_000002_A_fumigatus_A1163:2,110,815..2,113,165(+) |
| AFUB_034380 | 3.589717658 | 4.201746656 | 1.397012897 | 3.007664901 | 0.002632633 | 0.004404175 | | AFUB_034380 | Has domain(s) with predicted transferase | N/A | Protein kinase-like domain s | protein coding gene | | GO:0016772 | | | transferase activity, transf | 669 | 222 | | 192 | 0 | OG6_453808 | N/A | B0XZU2 | IPR011009 | 2.7.11.1 (Non-s | 0 | scf_000003_A_fumigatus_A1163:131,517..132,185(-) |
| AFUB_071480 | 37.51042664 | 4.21788003 | 0.500971918 | 8.419394133 | 3.78E-17 | 1.56E-16 | | AFUB_071480 | Protein of unknown function | N/A | N/A | protein coding gene | | | | | | 779 | 248 | | 24 | 1 | OG6_454648 | N/A | B0Y558 | N/A | | 0 | scf_000005_A_fumigatus_A1163:2,391,964..2,392,742(+) |
| AFUB_045500 | 3.631121468 | 4.222738016 | 1.402934729 | 3.009931916 | 0.002613063 | 0.004373102 | | AFUB_045500 | Has domain(s) with predicted role in trans | Major facilitator superfamily | Major facilitator superfamily; | protein coding gene | GO:0016021;G( | GO:0022857 | transmembrane | integral compon | transmembrane transport | 1738 | 518 | | 95 | 0 | OG6_166307 | PF07690 | B0Y036 | IPR011701;IPR036259 | | 10 | scf_000003_A_fumigatus_A1163:3,380,476..3,382,213(-) |
| AFUB_080620 | 6.890697023 | 4.230250316 | 1.145035913 | 3.694425886 | 0.000220384 | 0.000413639 | | AFUB_080620 | unspecified product | N/A | N/A | ncRNA gene | | | | | | 113 | | | 0 | 0 | N/A (orthology r | N/A | | N/A | | | scf_000007_A_fumigatus_A1163:263,644..263,756(-) |
| AFUB_039880 | 133.3449781 | 4.238325002 | 0.276722177 | 15.31617398 | 5.96E-53 | 7.92E-52 | | AFUB_039880 | Has domain(s) with predicted catalytic acti | CoA-transferase family III | CoA-transferase family III;C | protein coding gene | | GO:0008410;G( | metabolic process | | CoA-transferase activity;c | 1485 | 494 | | 564 | 1 | OG6_121197 | PF02515 | B0XY73 | IPR003673;IPR( | 5.1.99.4 (Alpha- | 0 | scf_000003_A_fumigatus_A1163:1,703,980..1,705,464(-) |
| AFUB_101480 | 639.2702416 | 4.240125902 | 0.143278011 | 29.59369591 | 1.80E-192 | 2.75E-190 | | AFUB_101480 | Protein of unknown function | Protein of unknown function DU | Protein of unknown function | protein coding gene | GO:0016021 | | | integral component of membrane | | 819 | 272 | | 46 | 0 | OG6_186788 | PF15496 | B0YF40 | IPR028018 | | 1 | scf_000011_A_fumigatus_A1163:287,668..288,486(+) |
| AFUB_039370 | 206.1981119 | 4.240317262 | 0.222134733 | 19.08894299 | 3.12E-81 | 7.73E-80 | | AFUB_039370 | Has domain(s) with predicted metal ion bi | Class II aldolase/adducin N-ter | Class II aldolase/adducin N- | protein coding gene | | GO:0046872 | | | metal ion binding | 1229 | 280 | | 698 | 2 | OG6_100477 | PF00596 | | IPR001303;IPR( | 4.1.2.- (Aldehyd | 0 | scf_000003_A_fumigatus_A1163:1,560,977..1,562,205(-) |
| AFUB_089530 | 106.7384791 | 4.241771757 | 0.300817488 | 14.10081505 | 3.75E-45 | 4.08E-44 | | AFUB_089530 | Protein of unknown function | N/A | N/A | protein coding gene | | | | | | 1242 | 413 | | 63 | 0 | OG6_454937 | N/A | B0YCA8 | N/A | | 0 | scf_000008_A_fumigatus_A1163:891,324..892,565(+) |
| AFUB_042710 | 3.727086922 | 4.242254872 | 1.390912409 | 3.049979887 | 0.002288567 | 0.003856704 | | AFUB_042710 | Protein of unknown function | N/A | N/A | protein coding gene | | | | | | 276 | 91 | | 1 | 0 | OG6_454391 | N/A | B0XZ04 | N/A | | 0 | scf_000003_A_fumigatus_A1163:2,476,848..2,477,123(-) |
| AFUB_044890 | 819.7570294 | 4.244254736 | 0.127677922 | 33.24188462 | 2.67E-242 | 6.28E-240 | | AFUB_044890 | Protein of unknown function | N/A | N/A | protein coding gene | | | | | | 1148 | 186 | MRFFLLLATTL/ | 80 | 0 | OG6_454451 | N/A | B0XZQ6 | N/A | | 0 | scf_000003_A_fumigatus_A1163:3,213,116..3,214,263(+) |
| AFUB_102100 | 7.035689512 | 4.25622354 | 1.146789347 | 3.711425774 | 0.000206095 | 0.000387953 | | AFUB_102100 | Protein of unknown function | Protein of unknown function DU | Protein of unknown function | protein coding gene | | | | | | 1179 | 392 | | 3 | 0 | OG6_454584 | PF14033 | B0YFA3 | IPR025340 | | 0 | scf_000011_A_fumigatus_A1163:469,345..470,523(+) |
| AFUB_045780 | 13.87308302 | 4.258695415 | 0.81627722 | 5.217217033 | 1.82E-07 | 4.41E-07 | | AFUB_045780 | Has domain(s) with predicted sequence-s | Zn(2)-C6 fungal-type DNA-binc | Zn(2)-C6 fungal-type DNA-b | protein coding gene | GO:0042025;G( | GO:0003677;G( | regulation of tra | host cell nucleus | DNA binding;DNA-binding | 2144 | 577 | | 798 | 3 | OG6_152930 | PF00172;PF04( | B0Y065 | IPR001138;IPR007219;IPR0368 | | 0 | scf_000003_A_fumigatus_A1163:3,468,030..3,470,173(+) |
| AFUB_049140 | 27.91891542 | 4.264504655 | 0.573982538 | 7.429676642 | 1.09E-13 | 3.84E-13 | | AFUB_049140 | Has domain(s) with predicted cation bindi | Glycoside hydrolase, family 42 | Glycoside hydrolase, family | protein coding gene | GO:0009341 | GO:0004565;G( | carbohydrate m | beta-galactosida | beta-galactosidase activit | 1772 | 565 | | 100 | 0 | OG6_186354 | PF02449;PF181 | B0Y195 | IPR001944;IPR( | 3.2.1.23 (Beta-g | 0 | scf_000004_A_fumigatus_A1163:118,273..120,044(+) |
| AFUB_013930 | 17.5116155 | 4.266885128 | 0.726066539 | 5.876713634 | 4.18E-09 | 1.14E-08 | | AFUB_013930 | Has domain(s) with predicted zinc ion binc | Alcohol dehydrogenase, N-tern | Alcohol dehydrogenase, zinc | protein coding gene | | GO:0016491;G( | obsolete oxidation-reduction pro( | | oxidoreductase activity;zi | 893 | 242 | | 252 | 0 | OG6_126503 | PF08240 | B0XMR8 | IPR002328;IPR( | 1.1.1.- (With NA | 1 | scf_000001_A_fumigatus_A1163:3,917,675..3,918,567(-) |
| AFUB_039820 | 101.4576219 | 4.268277665 | 0.311350995 | 13.70889361 | 8.98E-43 | 9.12E-42 | | AFUB_039820 | Has domain(s) with predicted RNA polyme | Mediator of RNA polymerase II | Mediator of RNA polymerase | protein coding gene | GO:0016592 | GO:0003712 | regulation of tra | mediator comple | transcription coregulator a | 1611 | 167 | | 71 | 0 | OG6_454188 | PF07544 | B0XY67 | IPR011425;IPR037212 | | 0 | scf_000003_A_fumigatus_A1163:1,691,816..1,693,426(-) |
| AFUB_080580 | 3.766360424 | 4.272611711 | 1.37833362 | 3.099838565 | 0.001936261 | 0.003291781 | | AFUB_080580 | Protein of unknown function | N/A | N/A | protein coding gene | | | | | | 521 | 100 | | 0 | 0 | OG6r21_23987: | N/A | B0Y9C9 | N/A | | 0 | scf_000007_A_fumigatus_A1163:255,296..255,816(-) |
| AFUB_096640 | 3.803462984 | 4.290511834 | 1.400734165 | 3.063045038 | 0.002190971 | 0.003700648 | | AFUB_096640 | Has domain(s) with predicted oxidoreduct | Ferric reductase transmembrar | FAD-binding 8;Ferric reducta | protein coding gene | GO:0016021;G( | GO:0016491 | ion transport;ob | integral compon | oxidoreductase activity | 1746 | 529 | | 163 | 5 | OG6_137686 | PF01794;PF08( | B0YDS7 | IPR013112;IPR013121;IPR0131 | | 6 | scf_000009_A_fumigatus_A1163:1,099,583..1,101,328(+) |
| AFUB_041530 | 17.82891435 | 4.294023829 | 0.722871461 | 5.940231505 | 2.85E-09 | 7.79E-09 | | AFUB_041530 | Protein of unknown function | N/A | N/A | protein coding gene | | | | | | 1095 | 254 | | 0 | 0 | OG6r21_23993( | N/A | B0XYN5 | N/A | | 0 | scf_000003_A_fumigatus_A1163:2,143,531..2,144,625(-) |
| AFUB_003730 | 1.871319279 | 4.29754884 | 1.549007906 | 2.774387932 | 0.005530569 | 0.008879133 | | AFUB_003730 | Protein of unknown function | N/A | N/A | protein coding gene | | | | | | 282 | 73 | | 1 | 0 | OG6_186583 | N/A | B0XMY4 | N/A | | 0 | scf_000001_A_fumigatus_A1163:1,030,969..1,031,250(-) |
| AFUB_097180 | 39.17346762 | 4.297697718 | 0.493252078 | 8.712984509 | 2.96E-18 | 1.29E-17 | | AFUB_097180 | Protein of unknown function | N/A | N/A | protein coding gene | | | | | | 597 | 198 | | 1 | 0 | OG6_454040 | N/A | B0YDY0 | N/A | | 0 | scf_000009_A_fumigatus_A1163:1,245,247..1,245,843(+) |
| AFUB_092480 | 3.84108051 | 4.3011182 | 1.372807958 | 3.13308076 | 0.001729818 | 0.002957511 | | AFUB_092480 | Has domain(s) with predicted nucleotide b | Short-chain dehydrogenase/re | Short-chain dehydrogenase/ | protein coding gene | GO:0016021 | GO:0016491 | metabolic proce | integral compon | oxidoreductase activity | 1189 | 276 | | 1441 | 3 | OG6_100207 | PF00106 | B0YBE1 | IPR002347;IPR( | 1.1.1.- (With NA | 1 | scf_000008_A_fumigatus_A1163:1,687,382..1,688,570(+) |
| AFUB_070530 | 1.878882201 | 4.301481127 | 1.554748625 | 2.766673055 | 0.005663152 | 0.009076868 | | AFUB_070530 | Protein of unknown function | N/A | N/A | protein coding gene | | | | | | 803 | 101 | | 1 | 0 | OG6_455091 | N/A | B0Y4W4 | N/A | | 0 | scf_000005_A_fumigatus_A1163:2,123,317..2,124,119(-) |
| AFUB_101840 | 117.9516434 | 4.302467965 | 0.289758117 | 14.84848125 | 7.12E-50 | 8.74E-49 | | AFUB_101840 | Has domain(s) with predicted role in trans | Major facilitator superfamily | Major facilitator superfamily; | protein coding gene | GO:0016021;G( | GO:0022857 | transmembrane | integral compon | transmembrane transport | 2008 | 533 | | 499 | 1 | OG6_123029 | PF07690 | B0YF78 | IPR011701;IPR036259 | | 14 | scf_000011_A_fumigatus_A1163:389,553..391,560(+) |
| AFUB_086720 | 1.922628899 | 4.325138333 | 1.538775419 | 2.810766458 | 0.004942365 | 0.007972764 | | AFUB_086720 | Has domain(s) with predicted nucleotide b | NAD-dependent epimerase/de | NAD-dependent epimerase/ | protein coding gene | | GO:0003824 | | | catalytic activity | 948 | 315 | | 165 | 1 | OG6_111933 | PF01370 | B0YBI5 | IPR001509;IPR( | 1.1.1.- (With NA | 0 | scf_000008_A_fumigatus_A1163:45,584..46,531(-) |
| AFUB_000120 | 3.881984847 | 4.330494178 | 1.374956733 | 3.149549418 | 0.001635224 | 0.002802245 | | AFUB_000120 | Has domain(s) with predicted succinate-se | Aldehyde dehydrogenase dom | Succinate semialdehyde deh | protein coding gene | | GO:0016491;G( | gamma-aminobutyric acid catab | | oxidoreductase activity;ox | 1749 | 529 | | 627 | 2 | OG6_102015 | PF00171 | B0XMG0 | IPR010102;IPR( | 1.2.1.16 (Succin | 0 | scf_000001_A_fumigatus_A1163:30,233..31,981(-) |
| AFUB_101580 | 18.4436711 | 4.339152119 | 0.727182145 | 5.967077371 | 2.42E-09 | 6.64E-09 | | AFUB_101580 | Has domain(s) with predicted role in trans | Major facilitator superfamily | Major facilitator superfamily; | protein coding gene | GO:0016021;G( | GO:0022857 | transmembrane | integral compon | transmembrane transport | 1911 | 573 | | 32 | 0 | OG6_186790 | PF07690 | B0YF51 | IPR011701;IPR036259 | | 12 | scf_000011_A_fumigatus_A1163:316,670..318,580(-) |
| AFUB_030280 | 11.08906193 | 4.340490363 | 0.92903288 | 4.672052473 | 2.98E-06 | 6.61E-06 | | AFUB_030280 | Has domain(s) with predicted protein bindi | N/A | F-box-like domain superfami | protein coding gene | | GO:0005515 | | | protein binding | 1038 | 345 | | 213 | 2 | OG6_152977 | N/A | B0XU44 | IPR036047 | | 0 | scf_000002_A_fumigatus_A1163:3,810,224..3,811,261(-) |
| AFUB_090400 | 25.78454754 | 4.342585315 | 0.613495901 | 7.078425968 | 1.46E-12 | 4.85E-12 | | AFUB_090400 | Protein of unknown function | NADP-dependent oxidoreducta | NADP-dependent oxidoredu | protein coding gene | | | | | | 1272 | 355 | | 75 | 0 | OG6_163837 | PF00248 | B0YCJ4 | IPR023210;IPR036812 | | 0 | scf_000008_A_fumigatus_A1163:1,136,133..1,137,404(-) |
| AFUB_092300 | 3.928796131 | 4.34317038 | 1.367366851 | 3.176302231 | 0.001491655 | 0.002568546 | | AFUB_092300 | Protein of unknown function | Tannase/feruloyl esterase | Tannase/feruloyl esterase;Al | protein coding gene | | GO:0030600;GO:0046872 | | | feruloyl esterase activity;m | 1722 | 573 | MRPSSCIPFSV | 192 | 0 | OG6_186954 | PF07519 | B0YBC3 | IPR011118;IPR( | 3.1.1.- (Carboxy | 0 | scf_000008_A_fumigatus_A1163:1,639,521..1,641,242(+) |
| AFUB_016440 | 104.2181733 | 4.365314304 | 0.312858429 | 13.95300206 | 3.02E-44 | 3.21E-43 | | AFUB_016440 | Protein of unknown function | N/A | N/A | protein coding gene | | | | | | 854 | 198 | MPSVSQLVSA/ | 201 | 1 | OG6_186635 | N/A | B0XP84 | N/A | | 0 | scf_000001_A_fumigatus_A1163:4,662,826..4,663,679(+) |
| AFUB_003600 | 124.0838499 | 4.371154157 | 0.292784958 | 14.92957218 | 2.12E-50 | 2.64E-49 | | AFUB_003600 | Has domain(s) with predicted carbon-sulfu | Glutathione-dependent formalc | Glutathione-dependent form | protein coding gene | | GO:0016846 | metabolic process | | carbon-sulfur lyase activit | 779 | 147 | | 167 | 0 | OG6_119679 | PF04828 | | IPR006913;IPR( | 4.4.1.22 (S-(hyd | 0 | scf_000001_A_fumigatus_A1163:990,489..991,267(-) |
| AFUB_050050 | 440.6205818 | 4.385767516 | 0.171153093 | 25.62482181 | 8.07E-145 | 6.37E-143 | | AFUB_050050 | Ortholog(s) have L-proline transmembran | Amino acid permease/ SLC12A | Amino acid permease, conse | protein coding gene | GO:0000324;G( | GO:0015193;G( | amino acid trans | fungal-type vac | L-proline transmembrane | 1747 | 566 | | 1824 | 4 | OG6_103579 | PF00324 | B0Y1P7 | IPR004840;IPR004841 | | 12 | scf_000004_A_fumigatus_A1163:322,062..323,808(-) |
| AFUB_045620 | 155.9732346 | 4.391158529 | 0.260728869 | 16.84185775 | 1.20E-63 | 2.02E-62 | | AFUB_045620 | Protein of unknown function | N/A | N/A | protein coding gene | | | | | | 1538 | 453 | | 128 | 3 | OG6_166300 | N/A | B0Y049 | N/A | | 0 | scf_000003_A_fumigatus_A1163:3,420,419..3,421,956(+) |
| AFUB_085979 | 2.035900787 | 4.41748957 | 1.530154629 | 2.886956315 | 0.003889882 | 0.006355189 | | AFUB_085979 | Has domain(s) with predicted hydrolase a | Ham1-like protein | Ham1-like protein;Inosine tri | protein coding gene | | GO:0016787;G( | nucleoside triphosphate catabol | | hydrolase activity;nucleos | 540 | 152 | | 1 | 0 | OG6_454997 | PF01725 | B0YAV6 | IPR002637;IPR( | 3.6.1.66 (XTP/d | 0 | scf_000007_A_fumigatus_A1163:1,807,615..1,808,154(-) |
| AFUB_081190 | 4.183213921 | 4.41975509 | 1.362742664 | 3.243279313 | 0.001181623 | 0.002057857 | | AFUB_081190 | Has domain(s) with predicted amino acid t | Amino acid/polyamine transpor | Amino acid/polyamine trans | protein coding gene | GO:0016021;G( | GO:0015171;G( | amino acid trans | integral compon | amino acid transmembrar | 1824 | 553 | | 287 | 2 | OG6_137852 | PF13520 | B0Y9I7 | IPR002293 | | 12 | scf_000007_A_fumigatus_A1163:430,712..432,535(+) |
| AFUB_097730 | 127.9880537 | 4.421463318 | 0.287530733 | 15.37735907 | 2.32E-53 | 3.11E-52 | | AFUB_097730 | Has domain(s) with predicted GTP binding | Dynamin superfamily;Dynamin | Dynamin central domain;Dyr | protein coding gene | | GO:0005525;GO:0003924 | | | GTP binding;GTPase acti | 2315 | 737 | | 910 | 5 | OG6_109505 | PF00350;PF01( | B0YE38 | IPR000375;IPR001401;IPR0228 | | 0 | scf_000009_A_fumigatus_A1163:1,441,365..1,443,679(+) |
| AFUB_046340 | 499.3448614 | 4.421966286 | 0.165138573 | 26.77730719 | 5.94E-158 | 5.78E-156 | | AFUB_046340 | Has domain(s) with predicted role in trans | Major facilitator superfamily | Major facilitator superfamily; | protein coding gene | GO:0016021;G( | GO:0022857 | transmembrane | integral compon | transmembrane transport | 1924 | 538 | | 357 | 0 | OG6_114261 | PF07690 | B0Y0I7 | IPR011701;IPR036259 | | 10 | scf_000003_A_fumigatus_A1163:3,613,218..3,615,141(+) |
| AFUB_051830 | 190.7889674 | 4.432760267 | 0.240171752 | 18.45662627 | 4.61E-76 | 1.00E-74 | | AFUB_051830 | Protein of unknown function | N/A | N/A | protein coding gene | | | | | | 183 | 60 | | 51 | 0 | OG6_453979 | N/A | B0Y2S7 | N/A | | 0 | scf_000004_A_fumigatus_A1163:824,946..825,128(+) |
| AFUB_073680 | 15278.27885 | 4.439676513 | 0.107426272 | 41.32766047 | 0 | 0 | | AFUB_073680 | Ortholog(s) have role in carbon utilization | Phosphoenolpyruvate carboxyl | Phosphoenolpyruvate carbo: | protein coding gene | | GO:0005524;G( | carbon utilization;gluconeogenes | | ATP binding;kinase activit | 4067 | 601 | | 379 | 0 | OG6_102030 | PF01293 | B0Y7G0 | IPR001272;IPR( | 4.1.1.32 (Phosp | 0 | scf_000006_A_fumigatus_A1163:411,173..415,239(-) |
| AFUB_047760 | 11.87387214 | 4.449659841 | 0.929431962 | 4.787504651 | 1.69E-06 | 3.81E-06 | | AFUB_047760 | Protein of unknown function | Protein of unknown function DU | Protein of unknown function | protein coding gene | GO:0016021 | | | integral component of membrane | | 720 | 239 | | 98 | 0 | OG6_166294 | PF06912 | B0XX51 | IPR010699 | | 5 | scf_000003_A_fumigatus_A1163:3,966,159..3,966,878(+) |
| AFUB_065550 | 43.3672902 | 4.452261075 | 0.494215017 | 9.008753118 | 2.08E-19 | 9.47E-19 | | AFUB_065550 | Protein of unknown function | N/A | N/A | protein coding gene | | GO:0008236 | | | serine-type peptidase acti | 642 | 213 | | 167 | 0 | OG6_166427 | N/A | B0Y638 | N/A | | 0 | scf_000005_A_fumigatus_A1163:732,392..733,033(-) |
| AFUB_088530 | 47.56733131 | 4.45294341 | 0.465359067 | 9.568833452 | 1.08E-21 | 5.36E-21 | | AFUB_088530 | Has domain(s) with predicted sequence-s | Zn(2)-C6 fungal-type DNA-binc | Zn(2)-C6 fungal-type DNA-b | protein coding gene | GO:0042025;G( | GO:0003700;G( | regulation of tra | host cell nucleus | DNA-binding transcription | 1303 | 405 | | 602 | 4 | OG6_143612 | PF00172;PF11951 | | IPR001138;IPR021858;IPR0368 | | 0 | scf_000008_A_fumigatus_A1163:530,514..531,816(+) |
| AFUB_057650 | 11.88347538 | 4.452980961 | 0.925012413 | 4.813968868 | 1.48E-06 | 3.35E-06 | | AFUB_057650 | Has domain(s) with predicted heme bindin | Cytochrome b5-like heme/ster | Cytochrome b5-like heme/st | protein coding gene | GO:0016021;G( | GO:0020037;G( | obsolete oxidati | integral compon | heme binding;metal ion bi | 1463 | 471 | | 1271 | 4 | OG6_100179 | PF00173;PF001 | B0Y0K6 | IPR001199;IPR( | 1.6.2.2 (Cytochr | 1 | scf_000004_A_fumigatus_A1163:2,488,541..2,490,003(+) |
| AFUB_046170 | 195.3090548 | 4.464371358 | 0.239179348 | 18.66537137 | 9.47E-78 | 2.12E-76 | | AFUB_046170 | Protein of unknown function | N/A | N/A | protein coding gene | | | | | | 855 | 192 | MATLKSRNIILL | 52 | 0 | OG6_453901 | N/A | B0Y0H1 | N/A | | 0 | scf_000003_A_fumigatus_A1163:3,573,454..3,574,308(-) |
| AFUB_024590 | 550.6462638 | 4.46825174 | 0.15458038 | 28.90568475 | 1.01E-183 | 1.36E-181 | | AFUB_024590 | Has domain(s) with predicted acetyl-CoA | Biotin carboxylase-like, N-term | Biotin/lipoyl attachment;Bioti | protein coding gene | | GO:0005524;G( | fatty acid biosynthetic process;m | | ATP binding;acetyl-CoA c | 7154 | 2292 | | 396 | 0 | OG6_101052 | PF00289;PF003 | B0XRR7 | IPR000089;IPR( | 6.3.4.14 (Biotin | 0 | scf_000002_A_fumigatus_A1163:2,212,111..2,219,264(-) |
| AFUB_028410 | 186.4043506 | 4.488374745 | 0.246196831 | 18.23083885 | 2.94E-74 | 6.18E-73 | | AFUB_028410 | Has domain(s) with predicted protein bindi | VIT domain;von Willebrand fac | von Willebrand factor, type A | protein coding gene | GO:0005694 | | | chromosome | | 2961 | 946 | | 227 | 1 | OG6_100585 | PF08487;PF137 | B0XT03 | IPR002035;IPR( | 2.4.2.- (Pentosy | 0 | scf_000002_A_fumigatus_A1163:3,255,010..3,257,970(-) |
| AFUB_013540 | 52.98618207 | 4.49687979 | 0.44783636 | 10.04134589 | 1.00E-23 | 5.38E-23 | | AFUB_013540 | Has domain(s) with predicted protein bindi | F-box domain | F-box domain;F-box-like don | protein coding gene | | GO:0005515 | | | protein binding | 1640 | 172 | | 115 | 0 | OG6_186618 | PF12937 | | IPR001810;IPR036047 | | 1 | scf_000001_A_fumigatus_A1163:3,821,269..3,822,908(-) |
| AFUB_099650 | 3303.856404 | 4.5199752 | 0.094545532 | 47.80739069 | 0 | 0 | | AFUB_099650 | Ortholog(s) have nitric oxide dioxygenase, | Globin;Oxidoreductase FAD/N | Globin;Oxidoreductase FAD/ | protein coding gene | GO:0005829;G( | GO:0071949;G( | cellular respons | cytosol;intracell | FAD binding;heme binding | 1764 | 413 | | 353 | 1 | OG6_109281 | PF00042;PF001 | B0YEM0 | IPR000971;IPR( | 1.14.12.17 (Nitri | 0 | scf_000010_A_fumigatus_A1163:399,319..401,082(-) |
| AFUB_028310 | 2.194550209 | 4.524963216 | 1.512312115 | 2.992082897 | 0.00277081 | 0.004620894 | | AFUB_028310 | Has domain(s) with predicted role in isopr | Polyprenyl synthetase | Polyprenyl synthetase;Isopre | protein coding gene | | | isoprenoid biosynthetic process | | | 306 | 65 | | 1 | 0 | OG6_453829 | PF00348 | B0XSZ3 | IPR000092;IPR008949 | | 0 | scf_000002_A_fumigatus_A1163:3,236,132..3,236,437(+) |
| AFUB_044630 | 2.203546881 | 4.529572992 | 1.497867173 | 3.024015128 | 0.00249444 | 0.004183885 | | AFUB_044630 | Has domain(s) with predicted protein bindi | Ankyrin repeat-containing dom | Ankyrin repeat;Ankyrin repea | protein coding gene | | GO:0005515 | | | protein binding | 1869 | 622 | | 7 | 0 | OG6_455295 | PF12796;PF136 | B0XZN1 | IPR002110;IPR020683;IPR0367 | | 0 | scf_000003_A_fumigatus_A1163:3,148,841..3,150,709(+) |
| AFUB_055080 | 2.206611467 | 4.530748109 | 1.497713898 | 3.025109212 | 0.002485434 | 0.004170958 | | AFUB_055080 | Has domain(s) with predicted zinc ion binc | Transcription factor domain, fu | Transcription factor domain, | protein coding gene | GO:0005634 | GO:0003677;G( | transcription, DN | nucleus | DNA binding;zinc ion bind | 2284 | 662 | | 83 | 0 | OG6_454782 | PF04082 | B0Y3Q9 | IPR007219 | | 0 | scf_000004_A_fumigatus_A1163:1,781,832..1,784,115(-) |
| AFUB_079760 | 162.9735959 | 4.53306582 | 0.26628332 | 17.02346892 | 5.50E-65 | 9.52E-64 | | AFUB_079760 | Ortholog(s) have role in secondary metab | Major facilitator superfamily | Major facilitator superfamily; | protein coding gene | GO:0016021;G( | GO:0022857 | cellular respons | integral compon | transmembrane transport | 1762 | 440 | | 2612 | 9 | OG6_100246 | PF07690 | B0Y961 | IPR011701;IPR( | 2.8.2.11 (Galact | 12 | scf_000006_A_fumigatus_A1163:2,161,045..2,162,806(+) |
| AFUB_039890 | 2.257921087 | 4.554432396 | 1.500370174 | 3.035539145 | 0.002401061 | 0.004032878 | | AFUB_039890 | unspecified product | N/A | N/A | ncRNA gene | | | | | | 72 | | | 0 | 0 | N/A (orthology r | N/A | | N/A | | | scf_000003_A_fumigatus_A1163:1,705,867..1,705,938(-) |
| AFUB_094370 | 17.14143931 | 4.575836981 | 0.798723811 | 5.728935231 | 1.01E-08 | 2.68E-08 | | AFUB_094370 | Has domain(s) with predicted and role in c | Rieske [2Fe-2S] iron-sulphur d | Aromatic-ring-hydroxylating | protein coding gene | | GO:0051537;G( | aromatic compound catabolic pr | | 2 iron, 2 sulfur cluster bin | 1759 | 498 | | 556 | 2 | OG6_109308 | PF00355 | B0YD58 | IPR001663;IPR( | 1.14.15.7 (Choli | 1 | scf_000009_A_fumigatus_A1163:436,366..438,124(+) |
| AFUB_029460 | 355.9192207 | 4.575947676 | 0.188329871 | 24.29751397 | 2.08E-130 | 1.35E-128 | | AFUB_029460 | Protein of unknown function | Methyltransferase type 11 | Methyltransferase type 11;S | protein coding gene | | GO:0008168 | | | methyltransferase activity | 6345 | 1542 | | 190 | 0 | OG6_166264 | PF08241 | | IPR013216;IPR029063 | | 0 | scf_000002_A_fumigatus_A1163:3,573,991..3,580,335(+) |
| AFUB_054380 | 293.6915034 | 4.607958512 | 0.206946359 | 22.2664392 | 7.82E-110 | 3.62E-108 | | AFUB_054380 | Protein of unknown function | Alpha crystallin/Hsp20 domain | Alpha crystallin/Hsp20 doma | protein coding gene | | | | | | 1083 | 327 | | 10 | 0 | OG6_455078 | PF00011 | B0Y3J1 | IPR002068;IPR008978 | | 0 | scf_000004_A_fumigatus_A1163:1,582,809..1,583,891(-) |
| AFUB_045090 | 4.724713654 | 4.618617203 | 1.333000784 | 3.464827072 | 0.000530573 | 0.000958852 | | AFUB_045090 | Has domain(s) with predicted oxidoreduct | NADP-dependent oxidoreducta | Aldo/keto reductase, conser | protein coding gene | | GO:0016491 | D-xylose catabolic process;obso | | oxidoreductase activity | 1301 | 274 | | 2440 | 9 | OG6_100072 | PF00248 | B0XZS6 | IPR018170;IPR( | 1.1.1.1 (Alcohol | 0 | scf_000003_A_fumigatus_A1163:3,262,177..3,263,477(-) |
| AFUB_028360 | 4.769172403 | 4.622334731 | 1.331386783 | 3.471819602 | 0.000516944 | 0.000935274 | | AFUB_028360 | Protein of unknown function | N/A | N/A | protein coding gene | | | | | | 884 | 221 | | 3 | 0 | OG6_454467 | N/A | B0XSZ8 | N/A | | 0 | scf_000002_A_fumigatus_A1163:3,243,379..3,244,262(+) |
| AFUB_102060 | 66.48804435 | 4.623835129 | 0.41491896 | 11.14394756 | 7.66E-29 | 4.99E-28 | | AFUB_102060 | Protein of unknown function | Amino acid transporter, transm | Amino acid transporter, trans | protein coding gene | GO:0016021 | | | integral component of membrane | | 2309 | 315 | | 3 | 0 | OG6_454261 | PF01490 | B0YFA0 | IPR013057 | | 5 | scf_000011_A_fumigatus_A1163:456,247..458,555(+) |
| AFUB_075690 | 4.788084532 | 4.63243529 | 1.332557407 | 3.47634951 | 0.000508289 | 0.000920135 | | AFUB_075690 | Ortholog(s) have carbon-sulfur lyase activ | Aminotransferase, class I/class | Aminotransferase, class I/cla | protein coding gene | | GO:0016846;G( | biosynthetic process;gliotoxin bi( | | carbon-sulfur lyase activit | 1428 | 435 | | 103 | 0 | OG6_455331 | PF00155 | B0Y808 | IPR004839;IPR( | 2.6.1.- (Transan | 0 | scf_000006_A_fumigatus_A1163:1,050,401..1,051,828(-) |
| AFUB_100500 | 8.94411135 | 4.634193213 | 1.118749831 | 4.142296236 | 3.44E-05 | 6.98E-05 | | AFUB_100500 | Has domain(s) with predicted substrate-sp | Major facilitator, sugar transpo | Sugar/inositol transporter;Ma | protein coding gene | GO:0016021;G( | GO:0022857 | fumiquinazoline | integral compon | transmembrane transport | 1687 | 520 | | 70 | 0 | OG6_454264 | PF00083 | B0YEV4 | IPR003663;IPR005828;IPR0362 | | 10 | scf_000010_A_fumigatus_A1163:640,232..641,918(-) |
| AFUB_036260 | 923.4221232 | 4.638004819 | 0.131846124 | 35.1774075 | 4.43E-271 | 1.38E-268 | | AFUB_036260 | Ortholog(s) have peptidase, dimethylallylt | Aromatic prenyltransferase, DM | Aromatic prenyltransferase D | protein coding gene | | GO:0004161;G( | alkaloid biosynthetic process;alk | | dimethylallyltranstransfera | 1472 | 472 | | 186 | 1 | OG6_143714 | PF11991 | B0XWK7 | IPR012148;IPR( | 2.5.1.- (Transfer | 0 | scf_000003_A_fumigatus_A1163:623,590..625,061(-) |
| AFUB_043420 | 2.425567181 | 4.656823191 | 1.486220118 | 3.133333437 | 0.00172833 | 0.00295549 | | AFUB_043420 | Protein of unknown function | N/A | Homologous-pairing protein | protein coding gene | | | reciprocal meiotic recombination | | | 1116 | 186 | | 193 | 0 | OG6_103529 | N/A | B0XZ76 | IPR040461 | | 0 | scf_000003_A_fumigatus_A1163:2,693,978..2,695,093(+) |
| AFUB_050200 | 91.67774591 | 4.664807967 | 0.3671366 | 12.70591918 | 5.48E-37 | 4.66E-36 | | AFUB_050200 | Has domain(s) with predicted zinc ion binc | Transcription factor domain, fu | Transcription factor domain, | protein coding gene | GO:0005634 | GO:0003677;G( | transcription, DN | nucleus | DNA binding;zinc ion bind | 2317 | 638 | | 411 | 4 | OG6_143688 | PF04082 | B0Y1X8 | IPR007219 | | 0 | scf_000004_A_fumigatus_A1163:355,833..358,149(+) |
| AFUB_086740 | 18.27552654 | 4.665099309 | 0.789353457 | 5.910025815 | 3.42E-09 | 9.32E-09 | | AFUB_086740 | Has domain(s) with predicted transferase | Transferase | Transferase | protein coding gene | | GO:0016747 | | | acyltransferase activity, tr | 1545 | 496 | | 317 | 1 | OG6_153072 | PF02458 | B0YBI7 | IPR003480 | | 0 | scf_000008_A_fumigatus_A1163:50,310..51,854(+) |
| AFUB_077900 | 4.914129731 | 4.668523952 | 1.335495729 | 3.495723612 | 0.000472778 | 0.000859564 | | AFUB_077900 | Has domain(s) with predicted hydrolase a | Glycoside hydrolase, family 3, | Glycoside hydrolase, family | protein coding gene | | GO:0004553 | carbohydrate metabolic process | | hydrolase activity, hydroly | 2834 | 856 | | 743 | 4 | OG6_110843 | PF00933;PF01915;PF07691;PF | | IPR001764;IPR( | 3.2.1.21 (Beta-g | 0 | scf_000006_A_fumigatus_A1163:1,664,209..1,667,042(-) |
| AFUB_101550 | 4.976066859 | 4.681467645 | 1.32778899 | 3.525761759 | 0.000422267 | 0.000770053 | | AFUB_101550 | Has domain(s) with predicted nucleotide b | N/A | Short-chain dehydrogenase/ | protein coding gene | | GO:0016491 | metabolic process | | oxidoreductase activity | 762 | 253 | | 243 | 1 | OG6_133082 | PF13561 | B0YF47 | IPR002347;IPR( | 1.1.1.100 (3-oxo | 0 | scf_000011_A_fumigatus_A1163:304,362..305,123(-) |
| AFUB_092140 | 305.8108264 | 4.713395919 | 0.209172397 | 22.53354641 | 1.95E-112 | 9.38E-111 | | AFUB_092140 | Has domain(s) with predicted cation trans | Cation efflux protein | Cation efflux protein;Cation | protein coding gene | GO:0016021 | GO:0008324 | cation transport; | integral compon | cation transmembrane tra | 2450 | 401 | | 471 | 1 | OG6_102585 | PF01545 | B0YBA7 | IPR002524;IPR( | 1.3.1.74 (2-alke | 6 | scf_000008_A_fumigatus_A1163:1,603,594..1,606,043(+) |
| AFUB_044700 | 2.514716553 | 4.720579696 | 1.459873332 | 3.233554303 | 0.001222601 | 0.00212423 | | AFUB_044700 | Has domain(s) with predicted oxidoreduct | N/A | Nitroreductase-like;Nitroredu | protein coding gene | | GO:0016491;G( | cellular response to oxidative str | | oxidoreductase activity;ox | 687 | 196 | | 19 | 0 | OG6_454482 | N/A | B0XZN8 | IPR000415;IPR033877 | | 0 | scf_000003_A_fumigatus_A1163:3,167,104..3,167,790(+) |
| AFUB_041860 | 2.550900328 | 4.735289345 | 1.459097668 | 3.24535461 | 0.001173044 | 0.002043655 | | AFUB_041860 | Has domain(s) with predicted nucleic acid | DEAD/DEAH box helicase dom | Helicase, C-terminal;DEAD/I | protein coding gene | | GO:0005524;GO:0003724;GO:0004386;GO:000 | | | ATP binding;RNA helicas | 2183 | 587 | | 187 | 0 | OG6_137770 | PF00270;PF002 | B0XYR9 | IPR001650;IPR( | 3.6.4.13 (RNA h | 0 | scf_000003_A_fumigatus_A1163:2,237,931..2,240,113(-) |
| AFUB_063700 | 22460.71938 | 4.74639152 | 0.084204456 | 56.36746256 | 0 | 0 | | AFUB_063700 | Ortholog(s) have glutamate dehydrogenas | Glutamate/phenylalanine/leuci | Glutamate/phenylalanine/leu | protein coding gene | GO:0005829;G( | GO:0004354;G( | cellular amino a | cytosol;extracell | glutamate dehydrogenase | 2083 | 458 | | 282 | 0 | OG6_100455 | PF00208;PF028 | B0Y5K4 | IPR006095;IPR( | 1.4.1.- (With NA | 0 | scf_000005_A_fumigatus_A1163:259,033..261,115(+) |
| AFUB_027040 | 5.196643387 | 4.757415677 | 1.320979924 | 3.601429204 | 0.000316473 | 0.000583866 | | AFUB_027040 | Protein of unknown function | N/A | N/A | protein coding gene | GO:0016021 | | | integral component of membrane | | 732 | 126 | | 1 | 0 | OG6_453981 | N/A | B0XSF8 | N/A | | 1 | scf_000002_A_fumigatus_A1163:2,881,888..2,882,619(-) |
| AFUB_086700 | 5.226021101 | 4.766616634 | 1.329218952 | 3.58602819 | 0.000335753 | 0.000617662 | | AFUB_086700 | Protein of unknown function | Beta-ketoacyl synthase, N-tern | Phosphopantetheine binding | protein coding gene | | GO:0003824;G( | arugosin biosynthetic process;as | | catalytic activity;phospho | 5573 | 1794 | | 3303 | 12 | OG6_100178 | PF00109;PF005 | B0YBI3 | IPR009081;IPR( | 1.3.1.10 (Enoyl- | 0 | scf_000008_A_fumigatus_A1163:37,572..43,144(-) |
| AFUB_045750 | 5.267612349 | 4.769031416 | 1.31850646 | 3.616995107 | 0.000298043 | 0.000551978 | | AFUB_045750 | Protein of unknown function | Alpha/beta hydrolase fold-1 | Alpha/beta hydrolase fold-1; | protein coding gene | | GO:0016787 | | | hydrolase activity | 889 | 268 | | 12 | 0 | OG6_216557 | PF00561 | B0Y062 | IPR000073;IPR029058 | | 0 | scf_000003_A_fumigatus_A1163:3,463,275..3,464,163(-) |
| AFUB_034530 | 2.655175544 | 4.801703787 | 1.448332852 | 3.315331679 | 0.000915344 | 0.001613364 | | AFUB_034530 | Ortholog(s) have role in biotin biosynthetic | Aminotransferase, class I/class | Aminotransferase, class I/cla | protein coding gene | GO:0005777 | GO:0003824;G( | biosynthetic pro | peroxisome | catalytic activity;pyridoxal | 1536 | 470 | | 122 | 0 | OG6_454640 | PF00155 | B0XZV7 | IPR004839;IPR( | 2.3.1.- (Transfer | 0 | scf_000003_A_fumigatus_A1163:166,463..167,998(+) |
| AFUB_061560 | 15.08091754 | 4.818578045 | 0.906260545 | 5.316989768 | 1.05E-07 | 2.60E-07 | | AFUB_061560 | Protein of unknown function | N/A | SKP1/BTB/POZ domain sup | protein coding gene | | | | | | 1016 | 321 | | 653 | 7 | OG6_121403 | N/A | B0Y278 | IPR011333 | | 0 | scf_000004_A_fumigatus_A1163:3,539,484..3,540,499(+) |
| AFUB_071770 | 15.29158863 | 4.830455483 | 0.904417875 | 5.340955346 | 9.25E-08 | 2.29E-07 | | AFUB_071770 | Ortholog(s) have role in xanthone-containi | NAD(P)-binding domain | NAD(P)-binding domain;NAD | protein coding gene | | | monodictyphenone biosynthetic process;secondary metab | | | 792 | 263 | | 86 | 0 | OG6_454081 | PF13460 | B0Y588 | IPR016040;IPR( | 1.1.1.- (With NA | 0 | scf_000005_A_fumigatus_A1163:2,531,133..2,531,924(+) |
| AFUB_069380 | 15.32046687 | 4.83366827 | 0.909680793 | 5.313587256 | 1.07E-07 | 2.65E-07 | | AFUB_069380 | Protein of unknown function | N/A | N/A | protein coding gene | | | | | | 309 | 102 | | 5 | 0 | OG6_453792 | N/A | B0Y4K2 | N/A | | 0 | scf_000005_A_fumigatus_A1163:1,821,834..1,822,142(+) |
| AFUB_044770 | 46.11204229 | 4.843521348 | 0.525802647 | 9.211671667 | 3.21E-20 | 1.50E-19 | | AFUB_044770 | Protein of unknown function | N/A | N/A | protein coding gene | GO:0016021 | | | integral component of membrane | | 643 | 177 | | 92 | 0 | OG6_153022 | N/A | B0XZP5 | N/A | | 2 | scf_000003_A_fumigatus_A1163:3,182,698..3,183,340(+) |
| AFUB_050250 | 10.27630857 | 4.846390514 | 1.107630823 | 4.375456527 | 1.21E-05 | 2.56E-05 | | AFUB_050250 | Protein of unknown function | N/A | N/A | protein coding gene | | | | | | 234 | 77 | | 1 | 0 | OG6_454096 | N/A | B0Y1Y3 | N/A | | 0 | scf_000004_A_fumigatus_A1163:367,126..367,359(+) |
| AFUB_049900 | 61.86048964 | 4.851887691 | 0.459000277 | 10.57055503 | 4.08E-26 | 2.39E-25 | | AFUB_049900 | Has domain(s) with predicted chitinase, ca | Chitin-binding, type 1;Glycosid | Chitin-binding, type 1;Glycos | protein coding gene | | GO:0003824;G( | carbohydrate metabolic process; | | catalytic activity;chitin bin | 4177 | 1142 | MRHLRPSFAFS | 5 | 0 | OG6_455213 | PF00187;PF00704 | | IPR001002;IPR( | 3.2.1.14 (Chitina | 0 | scf_000004_A_fumigatus_A1163:280,710..284,886(-) |
| AFUB_009790 | 311.0140341 | 4.859115306 | 0.221337364 | 21.95343438 | 8.03E-107 | 3.45E-105 | | AFUB_009790 | Has domain(s) with predicted role in trans | Major facilitator superfamily | Major facilitator superfamily; | protein coding gene | GO:0016021;G( | GO:0022857 | transmembrane | integral compon | transmembrane transport | 2480 | 559 | | 314 | 0 | OG6_143632 | PF07690 | B0XQF7 | IPR011701;IPR036259 | | 12 | scf_000001_A_fumigatus_A1163:2,746,030..2,748,509(+) |
| AFUB_049200 | 1388.207944 | 4.865887355 | 0.122637131 | 39.67711338 | 0 | 0 | | AFUB_049200 | Has domain(s) with predicted ATP binding | Cation-transporting P-type ATF | P-type ATPase;Cation-trans | protein coding gene | GO:0016021 | GO:0019829;G( | cation transport | integral compon | ATPase-coupled cation tr | 3671 | 1067 | | 418 | 1 | OG6_100215 | PF00122;PF00689;PF00690;PF | | IPR001757;IPR( | 3.2.1.52 (Beta-N | 8 | scf_000004_A_fumigatus_A1163:131,326..134,996(-) |
| AFUB_090510 | 26.31465091 | 4.876317215 | 0.69964367 | 6.969715333 | 3.18E-12 | 1.04E-11 | | AFUB_090510 | Has domain(s) with predicted triglyceride l | Fungal lipase-like domain | Fungal lipase-like domain;Al | protein coding gene | | GO:0004806 | lipid metabolic process | | triglyceride lipase activity | 947 | 281 | MFHPDISAGLL | 562 | 0 | OG6_100474 | PF01764 | B0YCK4 | IPR002921;IPR( | 3.1.1.- (Carboxy | 0 | scf_000008_A_fumigatus_A1163:1,165,195..1,166,141(-) |
| AFUB_099870 | 21.14295883 | 4.885135779 | 0.783991712 | 6.231106407 | 4.63E-10 | 1.32E-09 | | AFUB_099870 | unspecified product | N/A | N/A | ncRNA gene | | | | | | 88 | | | 0 | 0 | N/A (orthology r | N/A | | N/A | | | scf_000010_A_fumigatus_A1163:456,773..456,860(+) |
| AFUB_034430 | 10.64396309 | 4.890571202 | 1.101189812 | 4.441170041 | 8.95E-06 | 1.91E-05 | | AFUB_034430 | Has domain(s) with predicted shikimate 3- | Shikimate dehydrogenase subs | Shikimate dehydrogenase su | protein coding gene | | GO:0004764 | obsolete oxidation-reduction pro( | | shikimate 3-dehydrogena | 1032 | 319 | | 300 | 3 | OG6_103532 | PF08501 | B0XZU7 | IPR013708;IPR( | 1.1.1.24 (Quinat | 0 | scf_000003_A_fumigatus_A1163:140,427..141,458(-) |
| AFUB_086780 | 16.04535513 | 4.90645319 | 0.909510531 | 5.394608446 | 6.87E-08 | 1.72E-07 | | AFUB_086780 | Has domain(s) with predicted substrate-sp | Major facilitator, sugar transpo | Sugar/inositol transporter;Ma | protein coding gene | GO:0016021;G( | GO:0022857;G( | transmembrane | integral compon | transmembrane transport | 1434 | 424 | | 8 | 0 | OG6_454483 | PF00083 | B0YBJ1 | IPR003663;IPR005828;IPR0362 | | 10 | scf_000008_A_fumigatus_A1163:62,083..63,516(-) |
| AFUB_094970 | 247.117529 | 4.910134108 | 0.241592483 | 20.32403509 | 7.88E-92 | 2.47E-90 | | AFUB_094970 | Has domain(s) with predicted role in trans | Major facilitator superfamily | Major facilitator superfamily; | protein coding gene | GO:0016021;G( | GO:0022857 | transmembrane | integral compon | transmembrane transport | 2131 | 555 | | 3130 | 10 | OG6_100558 | PF07690 | B0YDB4 | IPR011701;IPR036259 | | 13 | scf_000009_A_fumigatus_A1163:595,112..597,242(+) |
| AFUB_091280 | 5.911731322 | 4.94563056 | 1.307369648 | 3.782886169 | 0.00015502 | 0.000295038 | | AFUB_091280 | Protein of unknown function | N/A | N/A | protein coding gene | | | | | | 297 | 98 | | 1 | 0 | OG6_454853 | N/A | B0YB21 | N/A | | 0 | scf_000008_A_fumigatus_A1163:1,401,524..1,401,820(+) |
| AFUB_046280 | 5.933663228 | 4.946710972 | 1.31291175 | 3.767740651 | 0.000164732 | 0.000312533 | | AFUB_046280 | Protein of unknown function | N/A | N/A | protein coding gene | | | | | | 727 | 195 | | 0 | 0 | OG6r21_23988: | N/A | B0Y0I1 | N/A | | 0 | scf_000003_A_fumigatus_A1163:3,596,617..3,597,343(+) |
| AFUB_086710 | 16.61293612 | 4.952981196 | 0.903222248 | 5.483679358 | 4.17E-08 | 1.06E-07 | | AFUB_086710 | Ortholog(s) have peptidase, dimethylallylt | Aromatic prenyltransferase, DM | Aromatic prenyltransferase D | protein coding gene | | GO:0004161;G( | alkaloid biosynthetic process;alk | | dimethylallyltranstransfera | 1362 | 453 | | 186 | 1 | OG6_143714 | PF11991 | B0YBI4 | IPR012148;IPR( | 2.5.1.- (Transfer | 0 | scf_000008_A_fumigatus_A1163:43,626..44,987(+) |
| AFUB_036880 | 5.978191927 | 4.969088347 | 1.311296062 | 3.789448081 | 0.000150982 | 0.000287466 | | AFUB_036880 | Protein of unknown function | N/A | N/A | protein coding gene | | | | | | 397 | 72 | | 0 | 0 | OG6r21_23996( | N/A | B0XWY8 | N/A | | 0 | scf_000003_A_fumigatus_A1163:830,213..830,609(-) |
| AFUB_057730 | 6.021216926 | 4.97175706 | 1.30363503 | 3.813764547 | 0.000136866 | 0.000262195 | | AFUB_057730 | Has domain(s) with predicted sequence-s | N/A | Basic-leucine zipper domain | protein coding gene | | GO:0003700;G( | regulation of transcription, DNA-( | | DNA-binding transcription | 810 | 257 | | 11 | 0 | OG6_453928 | N/A | B0Y0L4 | IPR004827 | | 0 | scf_000004_A_fumigatus_A1163:2,504,767..2,505,576(+) |
| AFUB_057810 | 17.02315095 | 4.990860766 | 0.89963013 | 5.547680764 | 2.89E-08 | 7.46E-08 | | AFUB_057810 | Protein of unknown function | N/A | N/A | protein coding gene | GO:0016021 | | | integral component of membrane | | 855 | 284 | | 84 | 1 | OG6_186871 | N/A | B0Y0M2 | N/A | | 2 | scf_000004_A_fumigatus_A1163:2,518,627..2,519,481(-) |
| AFUB_005950 | 28.63684199 | 5.000631732 | 0.697511036 | 7.169251053 | 7.54E-13 | 2.56E-12 | | AFUB_005950 | Protein of unknown function | N/A | N/A | protein coding gene | | | | | | 204 | 67 | | 1 | 0 | OG6_455049 | N/A | B0XP52 | N/A | | 0 | scf_000001_A_fumigatus_A1163:1,683,596..1,683,799(+) |
| AFUB_075470 | 3.079591963 | 5.001902564 | 1.483138048 | 3.372513146 | 0.000744855 | 0.001326451 | | AFUB_075470 | Protein of unknown function | N/A | N/A | protein coding gene | | | | | | 1482 | 493 | | 1 | 0 | OG6r21_16209( | N/A | B0Y7Y6 | N/A | | 0 | scf_000006_A_fumigatus_A1163:977,210..978,691(-) |
| AFUB_079810 | 59.25123136 | 5.054872325 | 0.493723522 | 10.23826514 | 1.34E-24 | 7.42E-24 | | AFUB_079810 | Has domain(s) with predicted zinc ion binc | Transcription factor domain, fu | Transcription factor domain, | protein coding gene | GO:0005634 | GO:0003677;G( | transcription, DN | nucleus | DNA binding;zinc ion bind | 2210 | 631 | | 1491 | 3 | OG6_107650 | PF04082 | | IPR007219 | | 0 | scf_000006_A_fumigatus_A1163:2,171,308..2,173,517(+) |
| AFUB_092590 | 178.8169354 | 5.116117113 | 0.295298946 | 17.3252129 | 3.04E-67 | 5.55E-66 | | AFUB_092590 | Protein of unknown function | N/A | N/A | protein coding gene | | | | | | 1072 | 298 | | 7 | 0 | OG6r21_11050: | N/A | B0YBF1 | N/A | | 0 | scf_000008_A_fumigatus_A1163:1,748,087..1,749,158(+) |
| AFUB_075790 | 199.8900177 | 5.181558158 | 0.285846469 | 18.12706723 | 1.95E-73 | 4.04E-72 | | AFUB_075790 | Ortholog(s) have flavin adenine dinucleoti | FAD/NAD(P)-binding domain | FAD/NAD(P)-binding domair | protein coding gene | GO:0005576 | GO:0050660;G( | cellular respons | extracellular reg | flavin adenine dinucleotid | 1137 | 334 | | 388 | 1 | OG6_116751 | PF07992 | B0Y818 | IPR023753;IPR( | 1.8.1.9 (Thiored | 0 | scf_000006_A_fumigatus_A1163:1,072,429..1,073,565(+) |
| AFUB_092320 | 32.36678721 | 5.192070475 | 0.693967667 | 7.481718125 | 7.34E-14 | 2.61E-13 | | AFUB_092320 | Has domain(s) with predicted cellulose bir | Cellulose-binding domain, fung | Cellulose-binding domain, fu | protein coding gene | GO:0005576 | GO:0008810;G( | carbohydrate m | extracellular reg | cellulase activity;cellulose | 782 | 237 | | 30 | 0 | OG6_143890 | PF00734;PF033 | B0YBC5 | IPR000254;IPR( | 3.2.1.4 (Cellulas | 0 | scf_000008_A_fumigatus_A1163:1,643,177..1,643,958(+) |
| AFUB_076180 | 343.7133305 | 5.219356875 | 0.226500253 | 23.04349248 | 1.71E-117 | 8.81E-116 | | AFUB_076180 | Ortholog(s) have intracellular localization | Alcohol dehydrogenase, C-tern | GroES-like superfamily;Alcol | protein coding gene | GO:0005622 | GO:0016491;G( | obsolete oxidati | intracellular ana | oxidoreductase activity;zi | 1503 | 347 | | 1021 | 1 | OG6_115538 | PF00107;PF082 | B0Y857 | IPR011032;IPR( | 1.1.1.1 (Alcohol | 0 | scf_000006_A_fumigatus_A1163:1,186,470..1,187,972(-) |
| AFUB_034130 | 7.111032474 | 5.223597221 | 1.284026969 | 4.068136689 | 4.74E-05 | 9.50E-05 | | AFUB_034130 | Protein of unknown function | Leucine-rich repeat | Leucine-rich repeat | protein coding gene | | GO:0005515 | | | protein binding | 2411 | 531 | | 41 | 0 | OG6_132702 | PF13516 | B0XZJ9 | IPR001611 | | 0 | scf_000003_A_fumigatus_A1163:86,055..88,465(-) |
| AFUB_086870 | 7.221926687 | 5.235656528 | 1.285702573 | 4.072214396 | 4.66E-05 | 9.34E-05 | | AFUB_086870 | unspecified product | N/A | N/A | ncRNA gene | | | | | | 111 | | | 0 | 0 | N/A (orthology r | N/A | | N/A | | | scf_000008_A_fumigatus_A1163:88,832..88,942(-) |
| AFUB_095410 | 3.591552059 | 5.237553168 | 1.389063074 | 3.7705654 | 0.000162878 | 0.00030926 | | AFUB_095410 | unspecified product | N/A | N/A | ncRNA gene | | | | | | 72 | | | 0 | 0 | N/A (orthology r | N/A | | N/A | | | scf_000009_A_fumigatus_A1163:707,073..707,144(+) |
| AFUB_046180 | 229.0781034 | 5.292321497 | 0.276181283 | 19.16249154 | 7.61E-82 | 1.92E-80 | | AFUB_046180 | Protein of unknown function | Necrosis inducing protein | Necrosis inducing protein | protein coding gene | | | obsolete pathogenesis | | | 1299 | 297 | MVSLNLPKLAL | 328 | 1 | OG6_112505 | PF05630 | B0Y0H2 | IPR008701 | | 0 | scf_000003_A_fumigatus_A1163:3,575,011..3,576,309(+) |
| AFUB_026010 | 7.687050359 | 5.328971733 | 1.280058515 | 4.163068852 | 3.14E-05 | 6.40E-05 | | AFUB_026010 | Protein of unknown function | N/A | N/A | protein coding gene | | | | | | 2430 | 694 | | 69 | 0 | OG6_455279 | N/A | B0XS57 | N/A | | 0 | scf_000002_A_fumigatus_A1163:2,586,181..2,588,610(+) |
| AFUB_045770 | 7.835037111 | 5.359461318 | 1.279080808 | 4.190088135 | 2.79E-05 | 5.71E-05 | | AFUB_045770 | Protein of unknown function | Ribonuclease E inhibitor RraA/ | Ribonuclease E inhibitor Rra | protein coding gene | | GO:0008168 | methylation | | methyltransferase activity | 1423 | 465 | | 530 | 2 | OG6_100381 | PF03737 | B0Y064 | IPR005493;IPR036704 | | 0 | scf_000003_A_fumigatus_A1163:3,466,124..3,467,546(-) |
| AFUB_075760 | 3.914782988 | 5.360762964 | 1.366621092 | 3.922640296 | 8.76E-05 | 0.000171327 | | AFUB_075760 | Has domain(s) with predicted role in trans | Major facilitator superfamily | Tetracycline resistance prote | protein coding gene | GO:0016021;G( | GO:0022857 | gliotoxin biosynt | integral compon | transmembrane transport | 1629 | 542 | | 23 | 0 | OG6_454139 | PF07690 | B0Y815 | IPR001411;IPR011701;IPR0362 | | 13 | scf_000006_A_fumigatus_A1163:1,067,161..1,068,789(+) |
| AFUB_071760 | 44.32438576 | 5.381170991 | 0.633782473 | 8.490564535 | 2.06E-17 | 8.58E-17 | | AFUB_071760 | Has domain(s) with predicted methyltransf | Methyltransferase domain 25 | S-adenosyl-L-methionine-de | protein coding gene | | GO:0008168 | | | methyltransferase activity | 1024 | 282 | | 332 | 1 | OG6_111949 | PF13649 | B0Y587 | IPR029063;IPR041698 | | 0 | scf_000005_A_fumigatus_A1163:2,529,523..2,530,546(-) |
| AFUB_076860 | 3.994713462 | 5.381761027 | 1.376752587 | 3.909025541 | 9.27E-05 | 0.000180908 | | AFUB_076860 | Has domain(s) with predicted nucleotide b | N/A | Short-chain dehydrogenase/ | protein coding gene | | GO:0016491 | | | oxidoreductase activity | 1059 | 301 | | 14 | 0 | OG6_453918 | PF13561 | B0Y8C5 | IPR002347;IPR020904;IPR0362 | | 0 | scf_000006_A_fumigatus_A1163:1,359,582..1,360,640(+) |
| AFUB_095110 | 14.99053835 | 5.390135113 | 1.089236 | 4.94854661 | 7.48E-07 | 1.73E-06 | | AFUB_095110 | Protein of unknown function | N/A | Class I glutamine amidotrans | protein coding gene | | | | | | 954 | 317 | | 446 | 3 | OG6_102842 | N/A | B0YDC8 | IPR029062 | | 0 | scf_000009_A_fumigatus_A1163:623,875..624,828(-) |
| AFUB_075750 | 4.077930746 | 5.41883646 | 1.362023233 | 3.978520137 | 6.93E-05 | 0.000136985 | | AFUB_075750 | Ortholog(s) have role in gliotoxin biosynth | N/A | Gamma-glutamylcyclotransf | protein coding gene | | GO:0003839 | gliotoxin biosynthetic process | | gamma-glutamylcyclotran | 872 | 273 | | 199 | 1 | OG6_113302 | N/A | B0Y814 | IPR017939 | 2.3.2.4 (Transfe | 0 | scf_000006_A_fumigatus_A1163:1,065,701..1,066,572(+) |
| AFUB_064140 | 155.1286181 | 5.454229089 | 0.348533356 | 15.6490878 | 3.37E-55 | 4.72E-54 | | AFUB_064140 | Has domain(s) with predicted oxidoreduct | FMN-dependent dehydrogenas | FMN-dependent dehydroger | protein coding gene | | GO:0010181;G( | obsolete oxidation-reduction pro( | | FMN binding;catalytic acti | 1426 | 436 | | 104 | 0 | OG6_137795 | PF01070 | | IPR000262;IPR012133 | | 0 | scf_000005_A_fumigatus_A1163:374,979..376,404(-) |
| AFUB_013240 | 380.2997512 | 5.456682035 | 0.231248303 | 23.59663599 | 4.17E-123 | 2.41E-121 | | AFUB_013240 | Has domain(s) with predicted zinc ion binc | Zinc finger C2H2-type | Zinc finger C2H2-type;Zinc fi | protein coding gene | GO:0005634 | GO:0003677;GO:0008270 | | nucleus | DNA binding;zinc ion bind | 4235 | 729 | | 165 | 0 | OG6_137712 | PF00096 | B0XRM9 | IPR013087;IPR036236 | | 0 | scf_000001_A_fumigatus_A1163:3,736,511..3,740,745(+) |
| AFUB_075770 | 4.417721271 | 5.53187943 | 1.344144326 | 4.115539769 | 3.86E-05 | 7.81E-05 | | AFUB_075770 | Ortholog(s) have intracellular localization | N/A | S-adenosyl-L-methionine-de | protein coding gene | GO:0005622 | GO:0008168 | methylation | intracellular ana | methyltransferase activity | 965 | 282 | | 3418 | 5 | OG6_103977 | PF13489 | B0Y816 | IPR029063 | 2.1.1.- (Methyltr | 0 | scf_000006_A_fumigatus_A1163:1,069,179..1,070,143(-) |

### Table S5

| | | | | | | | | | | | | | | | | | | | | | | | | | | | | |
|---|---|---|---|---|---|---|---|---|---|---|---|---|---|---|---|---|---|---|---|---|---|---|---|---|---|---|---|---|
| **AFUB_088060** | 1868.489917 | 5.578556855 | 0.132740258 | 42.02611114 | 0 | 0 | | AFUB_088060 | Has domain(s) with predicted transporter a | Proton-dependent oligopeptide | Proton-dependent oligopepti | protein coding gene | GO:0016021;G | GO:0022857;G | transmembrane | integral compon | transmembrane transport | 2341 | 578 | 1048 | 2 | OG6_100318 | PF00854 | B0YBW5 | IPR000109;IPR036259 | | 11 | scf_000008_A_fumigatus_A1163:388,161..390,501(+) |
| **AFUB_036290** | 1327.780628 | 5.625872805 | 0.146245501 | 38.46868973 | 0 | 0 | | AFUB_036290 | Ortholog(s) have role in secondary metab | Major facilitator superfamily | Major facilitator superfamily;I | protein coding gene | GO:0016021;G | GO:0022857 | cellular respons | integral compon | transmembrane transport | 1991 | 449 | 2612 | 9 | OG6_100246 | PF07690 | B0XWL0 | IPR011701;IPR | 2.8.2.11 (Galact | 12 | scf_000003_A_fumigatus_A1163:634,552..636,542(-) |
| **AFUB_071810** | 9.429936744 | 5.632364476 | 1.262631492 | 4.46081419 | 8.16E-06 | 1.75E-05 | | AFUB_071810 | Ortholog(s) have role in asperthecin biosy | Metallo-beta-lactamase | Metallo-beta-lactamase;Ribc | protein coding gene | | GO:0016787 | asperthecin biosynthetic process | | hydrolase activity | 1341 | 320 | 377 | 3 | OG6_101772 | PF00753 | B0Y592 | IPR001279;IPR | 3.1.2.- (Thiolest | 0 | scf_000005_A_fumigatus_A1163:2,543,260..2,544,600(+) |
| **AFUB_025330** | 4.780225703 | 5.653941134 | 1.364356442 | 4.144035209 | 3.41E-05 | 6.93E-05 | | AFUB_025330 | Protein of unknown function | Cation transporter | Cation transporter | protein coding gene | GO:0016021 | GO:0008324 | cation transport; | integral compon | cation transmembrane tra | 479 | 89 | 1 | 0 | OG6_453971 | PF02386 | B0XRZ0 | IPR003445 | | 0 | scf_000002_A_fumigatus_A1163:2,400,086..2,400,564(+) |
| **AFUB_017530** | 39.75528856 | 5.819080836 | 0.755983844 | 7.69736137 | 1.39E-14 | 5.11E-14 | | AFUB_017530 | Has domain(s) with predicted sequence-s | Zn(2)-C6 fungal-type DNA-binc | Zn(2)-C6 fungal-type DNA-b | protein coding gene | GO:0042025;G | GO:0003677;G | regulation of tra | host cell nucleus | DNA binding;DNA-binding | 1254 | 417 | 219 | 2 | OG6_166201 | PF00172;PF11 | B0XST4 | IPR001138;IPR021858;IPR0368 | | 0 | scf_000002_A_fumigatus_A1163:93,239..94,492(-) |
| **AFUB_044870** | 10.91019104 | 5.849738199 | 1.248228063 | 4.686433811 | 2.78E-06 | 6.18E-06 | | AFUB_044870 | Has domain(s) with predicted acetyltransfe | Hexapeptide repeat;Maltose/ga | Hexapeptide repeat;Trimeric | protein coding gene | | GO:0005525;G | GDP-mannose biosynthetic proc | | GTP binding;acetyltransfe | 744 | 215 | 694 | 5 | OG6_101136 | PF00132;PF124 | B0XZQ5 | IPR001451;IPR | 2.3.1.- (Transfer | 0 | scf_000003_A_fumigatus_A1163:3,211,178..3,211,921(-) |
| **AFUB_085550** | 11.36322831 | 5.904177654 | 1.248920839 | 4.727423443 | 2.27E-06 | 5.10E-06 | | AFUB_085550 | Ortholog(s) have lipase, carboxylic ester h | Alpha/beta hydrolase fold-3 | Alpha/beta hydrolase fold-3; | protein coding gene | | GO:0052689;G | metabolic process | | carboxylic ester hydrolase | 972 | 323 | 2599 | 8 | OG6_100160 | PF07859 | B0YAR3 | IPR013094;IPR | 2.3.1.- (Transfer | 0 | scf_000007_A_fumigatus_A1163:1,707,280..1,708,251(-) |
| **AFUB_075780** | 12.02862744 | 5.994676524 | 1.242011967 | 4.826585158 | 1.39E-06 | 3.15E-06 | | AFUB_075780 | Has domain(s) with predicted monooxyge | Cytochrome P450 | Cytochrome P450;Cytochror | protein coding gene | GO:0016021 | GO:0009055;G | obsolete oxidati | integral compon | electron transfer activity;h | 1572 | 504 | 1912 | 4 | OG6_103200 | PF00067 | B0Y817 | IPR001128;IPR | 1.14.14.- (With r | 1 | scf_000006_A_fumigatus_A1163:1,070,597..1,072,168(-) |
| **AFUB_100620** | 12.52465455 | 6.046729958 | 1.248157632 | 4.844524285 | 1.27E-06 | 2.89E-06 | | AFUB_100620 | Has domain(s) with predicted oxidoreduct | TauD/TfdA-like domain | TauD/TfdA-like domain | protein coding gene | | GO:0051213;G | obsolete oxidation-reduction pro | | dioxygenase activity;oxido | 1151 | 350 | 534 | 2 | OG6_105576 | PF02668 | B0YEV6 | IPR003819 | 1.14.11.17 (Tau | 0 | scf_000011_A_fumigatus_A1163:31,045..32,195(-) |
| **AFUB_000090** | 6.325224325 | 6.048791036 | 1.29600586 | 4.667255932 | 3.05E-06 | 6.77E-06 | | AFUB_000090 | Has domain(s) with predicted 2-dehydropa | Ketopantoate reductase, N-terr | 6-phosphogluconate dehydr | protein coding gene | | GO:0008677;G | obsolete oxidation-reduction pro | | 2-dehydropantoate 2-redu | 1497 | 346 | 1055 | 5 | OG6_102045 | PF02558;PF085 | B0XMF7 | IPR008927;IPR | 1.1.1.169 (2-deh | 0 | scf_000001_A_fumigatus_A1163:22,791..24,287(+) |
| **AFUB_046450** | 164.9636583 | 6.072924094 | 0.407407915 | 14.90624965 | 3.00E-50 | 3.73E-49 | | AFUB_046450 | Protein of unknown function | N/A | N/A | protein coding gene | | | | | | 371 | 77 | 16 | 0 | OG6_453833 | N/A | B0Y0J6 | N/A | | 0 | scf_000003_A_fumigatus_A1163:3,636,651..3,637,021(+) |
| **AFUB_068510** | 6.547244625 | 6.094379666 | 1.296478358 | 4.700718396 | 2.59E-06 | 5.77E-06 | | AFUB_068510 | Protein of unknown function | N/A | N/A | protein coding gene | | | | | | 808 | 175 | 1 | 0 | OG6_454862 | N/A | B0Y6X9 | N/A | | 0 | scf_000005_A_fumigatus_A1163:1,586,765..1,587,572(-) |
| **AFUB_036240** | 72.79899251 | 6.112077828 | 0.613873458 | 9.956576143 | 2.36E-23 | 1.25E-22 | | AFUB_036240 | Ortholog(s) have role in fumiquinazoline C | FAD linked oxidase, N-terminal | FAD linked oxidase, N-termi | protein coding gene | GO:0005576;G | GO:0071949;G | asperfuranone b | extracellular reg | FAD binding;UDP-N-acety | 1567 | 468 | 83 | 0 | OG6_166372 | PF01565 | B0XWK5 | IPR006094;IPR | 1.5.3.6 ((R)-6-hy | 0 | scf_000003_A_fumigatus_A1163:618,814..620,380(+) |
| **AFUB_021250** | 6.763357979 | 6.147278226 | 1.303853376 | 4.714700547 | 2.42E-06 | 5.41E-06 | | AFUB_021250 | Protein of unknown function | N/A | N/A | protein coding gene | | | | | | 150 | 49 | 2 | 0 | OG6_454157 | N/A | B0XV09 | N/A | | 0 | scf_000002_A_fumigatus_A1163:1,149,427..1,149,576(-) |
| **AFUB_071750** | 7.471559881 | 6.286774642 | 1.27972778 | 4.912587459 | 8.99E-07 | 2.06E-06 | | AFUB_071750 | Ortholog(s) have role in xanthone-contain | Questin oxidase-like | Questin oxidase-like | protein coding gene | | GO:0016491 | monodictyphenone biosynthetic | | oxidoreductase activity | 1338 | 445 | 535 | 3 | OG6_109991 | PF14027 | B0Y586 | IPR025337 | 1.14.13.43 (Que | 0 | scf_000005_A_fumigatus_A1163:2,528,025..2,529,362(+) |
| **AFUB_017260** | 7377.604911 | 6.35150245 | 0.1074405 | 59.11646383 | 0 | 0 | | AFUB_017260 | Protein of unknown function | Amino acid transporter, transm | Amino acid transporter, trans | protein coding gene | GO:0016021 | | | integral component of membrane | | 1927 | 457 | 614 | 5 | OG6_124686 | PF01490 | B0XSQ8 | IPR013057 | | 11 | scf_000002_A_fumigatus_A1163:26,512..28,438(+) |
| **AFUB_036280** | 274.4958351 | 6.368555892 | 0.348606446 | 18.26861195 | 1.47E-74 | 3.11E-73 | | AFUB_036280 | Ortholog(s) have role in secondary metab | O-methyltransferase domain | O-methyltransferase domain | protein coding gene | | GO:0008171 | methylation;secondary metabolit | | O-methyltransferase activ | 1586 | 433 | 515 | 4 | OG6_102258 | PF00891 | B0XWK9 | IPR001077;IPR | 2.1.1.- (Methyltr | 0 | scf_000003_A_fumigatus_A1163:632,735..634,320(+) |
| **AFUB_025480** | 247.6071849 | 6.379969418 | 0.368803778 | 17.29908909 | 4.78E-67 | 8.69E-66 | | AFUB_025480 | Protein of unknown function | Phytanoyl-CoA dioxygenase | Phytanoyl-CoA dioxygenase | protein coding gene | | GO:0051213 | | | dioxygenase activity | 1329 | 442 | 170 | 0 | OG6_133096 | PF05721 | B0XS05 | IPR008775 | | 0 | scf_000002_A_fumigatus_A1163:2,441,559..2,442,887(-) |
| **AFUB_022090** | 4999.153759 | 6.388512689 | 0.112592723 | 56.74001405 | 0 | 0 | AoxA | AFUB_022090 | Ortholog(s) have alternative oxidase activi | Alternative oxidase | Alternative oxidase | protein coding gene | GO:0016021;G | GO:0009916;G | alternative respi | integral compon | alternative oxidase activity | 1178 | 352 | 363 | 0 | OG6_103139 | PF01786 | B0XVF7 | IPR002680 | 1.10.3.11 (Ubiqu | 2 | scf_000002_A_fumigatus_A1163:1,378,107..1,379,284(+) |
| **AFUB_099310** | 862.1787365 | 6.432533892 | 0.214840061 | 29.94103544 | 5.76E-197 | 9.24E-195 | | AFUB_099310 | Has domain(s) with predicted glutamate-a | Glutamine synthetase, catalytic | Glutamine synthetase, cataly | protein coding gene | | GO:0003824;G | nitrogen compound metabolic pr | | catalytic activity;glutamate | 1974 | 535 | 566 | 4 | OG6_100656 | PF00120 | | IPR008146;IPR | 6.3.1.2 (Glutami | 1 | scf_000010_A_fumigatus_A1163:331,469..333,442(-) |
| **AFUB_099340** | 595.1951744 | 6.447239033 | 0.249006191 | 25.89188251 | 8.22E-148 | 6.66E-146 | | AFUB_099340 | Has domain(s) with predicted nucleotide b | N/A | Short-chain dehydrogenase/ | protein coding gene | | GO:0016491 | metabolic process | | oxidoreductase activity | 954 | 277 | 1041 | 3 | OG6_101626 | PF13561 | B0YEI9 | IPR002347;IPR | 1.1.1.- (With NA | 0 | scf_000010_A_fumigatus_A1163:336,954..337,907(+) |
| **AFUB_049210** | 8.531835792 | 6.480857283 | 1.266843513 | 5.115752038 | 3.12E-07 | 7.44E-07 | | AFUB_049210 | Protein of unknown function | N/A | N/A | protein coding gene | | | | | | 225 | 74 | 1 | 0 | OG6_454104 | N/A | B0Y1A2 | N/A | | 0 | scf_000004_A_fumigatus_A1163:135,011..135,235(-) |
| **AFUB_048500** | 8.548395387 | 6.48287879 | 1.267306811 | 5.115476956 | 3.13E-07 | 7.45E-07 | | AFUB_048500 | Has domain(s) with predicted protein bindi | Ankyrin repeat-containing dom | Ankyrin repeat;Ankyrin repe | protein coding gene | | GO:0005515 | | | protein binding | 1765 | 546 | 10 | 0 | OG6_454901 | PF12796 | B0XXJ0 | IPR002110;IPR020683;IPR0367 | | 0 | scf_000003_A_fumigatus_A1163:4,192,858..4,194,622(+) |
| **AFUB_035700** | 18.00343373 | 6.583858492 | 1.22193003 | 5.38808142 | 7.12E-08 | 1.78E-07 | | AFUB_035700 | Protein of unknown function | Alpha/beta hydrolase fold-1 | Alpha/beta hydrolase fold-1; | protein coding gene | | | | | | 1080 | 359 | 15 | 0 | OG6_455054 | PF12697 | B0XWF2 | IPR000073;IPR029058 | | 0 | scf_000003_A_fumigatus_A1163:491,471..492,550(-) |
| **AFUB_099320** | 747.7456189 | 6.602981727 | 0.235366517 | 28.05404023 | 3.57E-173 | 4.04E-171 | | AFUB_099320 | Protein of unknown function | Glutamine amidotransferase | Glutamine amidotransferase | protein coding gene | | | glutamine metabolic process | | | 912 | 258 | 446 | 3 | OG6_102842 | PF00117 | B0YEI7 | IPR017926;IPR | 2.4.2.- (Pentosy | 0 | scf_000010_A_fumigatus_A1163:333,623..334,534(+) |
| **AFUB_077870** | 75.56886132 | 6.761488374 | 0.743579164 | 9.093165464 | 9.62E-20 | 4.44E-19 | | AFUB_077870 | Protein of unknown function | N/A | N/A | protein coding gene | GO:0016021 | | | integral component of membrane | | 1128 | 375 | 28 | 0 | OG6_454242 | N/A | B0Y8M4 | N/A | | 1 | scf_000006_A_fumigatus_A1163:1,648,669..1,649,796(-) |
| **AFUB_088580** | 534.7018403 | 6.775620575 | 0.288706109 | 23.46892003 | 8.47E-122 | 4.83E-120 | | AFUB_088580 | Has domain(s) with predicted heme bindin | Indoleamine 2,3-dioxygenase | Indoleamine 2,3-dioxygenas | protein coding gene | | GO:0051213;G | tryptophan catabolic process to k | | dioxygenase activity;heme | 1920 | 496 | 189 | 0 | OG6_115958 | PF01231 | B0YC17 | IPR000898;IPR | 1.13.11.11 (Tryp | 0 | scf_000008_A_fumigatus_A1163:539,704..541,623(-) |
| **AFUB_092830** | 15.43849512 | 7.333404094 | 1.230320383 | 5.960564577 | 2.51E-09 | 6.90E-09 | | AFUB_092830 | Has domain(s) with predicted nucleic acid | Helicase, C-terminal | Helicase, C-terminal;Helicas | protein coding gene | | GO:0005524;GO:0004386;GO:0016787;GO:000 | | | ATP binding;helicase acti | 3205 | 706 | 61 | 0 | OG6_137722 | PF00271 | B0YBH5 | IPR001650;IPR014001;IPR0274 | | 0 | scf_000008_A_fumigatus_A1163:1,837,874..1,841,078(-) |
| **AFUB_061410** | 58.1188495 | 7.385059558 | 1.038146918 | 7.113694057 | 1.13E-12 | 3.78E-12 | | AFUB_061410 | Protein of unknown function | Uncharacterised protein family | Uncharacterised protein fam | protein coding gene | | | | | | 381 | 126 | 101 | 0 | OG6_107843 | PF01906 | B0Y263 | IPR002765;IPR035439 | | 0 | scf_000004_A_fumigatus_A1163:3,505,509..3,505,889(+) |
| **AFUB_075800** | 17.20413058 | 7.493113346 | 1.223923091 | 6.122209312 | 9.23E-10 | 2.59E-09 | | AFUB_075800 | Protein of unknown function | Acetylaranotin biosynthesis clu | Acetylaranotin biosynthesis | protein coding gene | | | | | | 617 | 179 | 24 | 0 | OG6_454025 | PF08982 | B0Y819 | IPR015075 | | 0 | scf_000006_A_fumigatus_A1163:1,073,865..1,074,481(+) |
| **AFUB_018360** | 640.4379109 | 7.678320724 | 0.352156445 | 21.80372058 | 2.14E-105 | 8.92E-104 | | AFUB_018360 | Has domain(s) with predicted NAD binding | D-isomer specific 2-hydroxyaci | D-isomer specific 2-hydroxya | protein coding gene | | GO:0051287;G | metabolic process;obsolete oxid | | NAD binding;obsolete cof | 1112 | 350 | 242 | 0 | OG6_143663 | PF00389;PF028 | B0XT83 | IPR006139;IPR | 1.1.1.- (With NA | 0 | scf_000002_A_fumigatus_A1163:297,040..298,151(-) |
| **AFUB_035570** | 206.8318684 | 7.897749034 | 0.657239599 | 12.01654472 | 2.91E-33 | 2.20E-32 | | AFUB_035570 | Protein of unknown function | Cupin 2, conserved barrel | RmlC-like cupin domain sup | protein coding gene | | | | | | 474 | 127 | 123 | 0 | OG6_129963 | PF07883 | B0Y0D0 | IPR011051;IPR013096 | | 0 | scf_000003_A_fumigatus_A1163:464,088..464,561(+) |
| **AFUB_017540** | 64.82660309 | 8.440846022 | 1.192017648 | 7.081141822 | 1.43E-12 | 4.76E-12 | | AFUB_017540 | Protein of unknown function | NADP-dependent oxidoreducta | NADP-dependent oxidoredu | protein coding gene | | | | | | 1300 | 329 | 4 | 0 | OG6_455333 | PF00248 | B0XST5 | IPR023210;IPR036812 | | 0 | scf_000002_A_fumigatus_A1163:94,901..96,200(+) |
| **AFUB_097280** | 338.5368459 | 9.348515678 | 0.839660068 | 11.13369092 | 8.60E-29 | 5.59E-28 | | AFUB_097280 | Protein of unknown function | Xylose isomerase-like, TIM bar | Xylose isomerase-like, TIM b | protein coding gene | | GO:0051213 | | | dioxygenase activity | 1599 | 388 | 66 | 0 | OG6_454128 | PF01261 | B0YDZ0 | IPR013022;IPR | 4.2.1.- (Hydro-ly | 0 | scf_000009_A_fumigatus_A1163:1,265,727..1,267,325(-) |
| **AFUB_034150** | 141.4917306 | 9.568697737 | 1.186257603 | 8.066289915 | 7.25E-16 | 2.81E-15 | | AFUB_034150 | Protein of unknown function | Glyoxalase/fosfomycin resistan | Glyoxalase/fosfomycin resist | protein coding gene | | | | | | 509 | 153 | 91 | 0 | OG6_129479 | PF00903 | B0XZK1 | IPR004360;IPR029068 | | 0 | scf_000003_A_fumigatus_A1163:90,628..91,136(-) |
| **AFUB_026030** | 10292.78265 | 9.75280634 | 0.193503369 | 50.40122244 | 0 | 0 | | AFUB_026030 | Has domain(s) with predicted iron ion bind | Inositol oxygenase | Inositol oxygenase | protein coding gene | GO:0005737 | GO:0050113;G | inositol catabolic | cytoplasm | inositol oxygenase activity | 1155 | 315 | 289 | 0 | OG6_104704 | PF05153 | B0XS59 | IPR007828 | 1.13.99.1 (Inosit | 0 | scf_000002_A_fumigatus_A1163:2,591,242..2,592,396(+) |
